# Empirical measures of mutational effects define neutral models of regulatory evolution in *Saccharomyces cerevisiae*

**DOI:** 10.1101/551804

**Authors:** Andrea Hodgins-Davis, Fabien Duveau, Elizabeth Walker, Patricia J Wittkopp

**Author notes:** Corresponding author: Dr. Patricia J Wittkopp, 1105 N University Ave, Ann Arbor, MI, 48109 USA.

## Abstract

Understanding how phenotypes evolve requires disentangling the effects of mutation generating new variation from the effects of selection filtering it. Evolutionary models frequently assume that mutation introduces phenotypic variation symmetrically around the population mean, yet few studies have tested this assumption by deeply sampling the distributions of mutational effects for particular traits. Here, we examine distributions of mutational effects for gene expression in the budding yeast *Saccharomyces cerevisiae* by measuring the effects of thousands of point mutations introduced randomly throughout the genome. We find that the distributions of mutational effects differ for the 10 genes surveyed and violate the assumption of normality. For example, all ten distributions of mutational effects included more mutations with large effects than expected for normally distributed phenotypes. In addition, some genes also showed asymmetries in their distribution of mutational effects, with new mutations more likely to increase than decrease the gene’s expression or vice versa. Neutral models of regulatory evolution that take these empirically determined distributions into account suggest that neutral processes may explain more expression variation within natural populations than currently appreciated.

**Significance statement:** New mutations tend to arise randomly throughout the genome, but their phenotypic effects are often not random. This disconnect results from interactions among genes that define the genotype-phenotype map. The structure of this map is poorly known and different for each trait, making it challenging to predict the distribution of mutational effects for specific phenotypes. Empirical measures of the distribution of mutational effects are thus necessary to understand how traits can change in the absence of natural selection. In this work, we define such distributions for expression of ten genes in *S. cerevisiae* and show that they predict greater neutral expression divergence than commonly used models of phenotypic evolution.

## Introduction

Variation in gene expression is widespread within and between species (1, 2). This variation reflects the joint action of mutation introducing new phenotypic variation and selection filtering variants based on their fitness effects. When genes exhibit differences in the rate at which they accumulate expression variation over time, selection pressure that varies among genes is often invoked to explain variability in expression divergence (3, 4). However, variability in the effects of new mutations on gene expression may also affect evolutionary outcomes by biasing the variety of expression phenotypes available for selection and drift to act upon (5, 6). Identifying such biases in mutational effects is challenging because the effects of mutation are confounded with the effects of selection in natural populations.

One way to isolate the effects of new mutations on gene expression is to perform a mutation accumulation experiment, which allows new mutations to accumulate in the near absence of natural selection (7). By comparing expression among evolved lines at the end of a period of mutation accumulation, such experiments have been used to estimate mutational variance (*V*_*m*_) for gene expression, which is the variance in a gene’s expression due to spontaneous mutations each generation. *V*_*m*_ for gene expression has been estimated genome-wide for model organisms including budding yeast (8), nematode worms (9, 10), Drosophilid flies (11–13). *V*_*m*_ differs among genes (14), with a median genome-wide *V*_*m*_ for gene expression estimated to be in the range of 10^-5^ to 10^-4^ for each species (8, 9, 11, 15). Differences in expression *V*_*m*_ observed among genes in these studies reveal differences in the potential for a gene’s expression to change over time. For example, in yeast, genes with more transcriptional regulators (as estimated from transcriptional profiling of gene deletion strains) tended to have higher *V*_*m*_ for expression than genes with fewer transcriptional regulators (8), suggesting that differences among regulatory networks can influence changes in gene expression due to new mutations.

Although these mutation accumulation studies offer a global view of transcriptional changes across the genome, they provide a very limited view of the distribution of mutational effects for any single gene because the number of spontaneous mutations sampled in each study is low. For example, the 12.1Mb yeast genome has a point mutation rate in the range of 10^-9^ to 10^-10^ per base pair per cell division, suggesting that only four point mutations occur on average every 1000 cell divisions. Consequently, even ambitious MA experiments that capture 2-5 thousand generations of spontaneous mutations are expected to survey fewer than two dozen point mutations per line. Mutagenesis studies, by contrast, can sample mutations affecting expression of a given gene much more deeply, typically trading off breadth of information across the genome for more focused and comprehensive descriptions of distributions of mutational effects for single genes. Thus far, such single-gene studies have focused primarily on mutations in *cis*-acting sequences controlling a gene’s expression, such as promoters (16). For example, massively parallel reporter gene approaches have been used to describe the effects of thousands of mutations in promoters and enhancers on gene expression from diverse organisms including viruses, bacteria, yeast, and metazoans (e.g. 17–20). However, these *cis-*acting sequences are only one part of the mutational target for gene expression (21). Regions of the genome encoding or regulating *trans*-acting factors that interact with *cis*-acting sequences, either directly or indirectly, can also harbor mutations that affect gene expression (22). This *trans-*mutational target size is expected to be much larger than the *cis*-mutational target size (23) and can show different biases in the effects on expression (24).

Here, we use genome-wide mutagenesis to deeply sample and compare the effects of new mutations on expression of ten focal genes. We observe differences in distributions of mutational effects among these genes that are only partially captured by quantifying variance of mutational distributions (*V*_*m,*_). In particular, we also observed differences in higher moments of these distributions, including the extent of asymmetry described by skewness and the frequency of mutations with extreme effects on expression related to kurtosis. Consistent with these observations, we find that all ten distributions of mutational effects for gene expression are non-normal with heavy tails (i.e., they contain more extreme events than a normal distribution). By using these empirically determined distributions of mutational effects to parametrize neutral models of gene expression evolution, we show that dramatic differences in expression divergence can occur among genes even in the absence of selection. In other words, we find that distributions of mutational effects for gene expression are (i) more complex than frequently assumed, (ii) different among genes, and (iii) able to introduce biases in the direction of neutral evolution. These observations suggest that failing to account for mutational biases may underestimate the role of neutral evolution in expression divergence.

## Results and Discussion

To compare the effects of mutations on gene expression driven by promoters from different genes, we selected ten promoters from *S. cerevisae*: *GPD1, OST1, PFY1, RNR1, RNR2, STM1, TDH1, TDH2, TDH3*, and *VMA7*. These promoters vary for properties previously hypothesized to affect the mutability of gene expression, including expression noise (25), nucleosome occupancy (25), and number of mismatches to a canonical TATA box (8) (Table S1). This set of genes includes three paralogs, TDH1, TDH2, and TDH3, and two genes acting in the same molecular complex, RNR1 and RNR2. All of these promoters drive expression at levels that can be reliably detected by flow cytometry. For each gene, we cloned the promoter sequence upstream of the coding sequence of a yellow fluorescent protein (YFP) and inserted the resulting reporter gene into the *S. cerevisiae* genome at the *ho* locus. YFP expression level from these reporter genes is therefore expected to measure effects of mutations in the *cis*-acting promoter sequence as well as all *trans*-acting regulators of the gene from which the promoter was derived.

To assess the impact of new mutations on expression driven by each promoter, we exposed cells carrying each reporter gene to a low dose of ethyl methanesulfonate (EMS) (Figure 1a), which is a chemical mutagen that primarily introduces G->A and C-> T point mutations (26). While these mutations are a subset of the types of changes that arise spontaneously, they are the most common type of point mutation observed in mutation accumulation lines (27, 28) and the most common type of single nucleotide polymorphism segregating in natural populations of *S. cerevisiae* (29). Using a canvanine resistance assay (30, 31), we estimated that the EMS conditions used introduced ∼29 mutations per cell (95% percentiles: 24-39). Following mutagenesis, we isolated single cells from each of the mutagenized populations randomly with respect to the YFP fluorescence level using fluorescence activated cell sorting (FACS) (Figure 1a). Each of these sorted cells was grown clonally in four replicate populations. YFP fluorescence levels were then estimated based on at least ∼ 12 thousand events captured with flow cytometry from each replicate population (Supplementary Methods, Table S2a). These YFP fluorescence levels were used to estimate YFP mRNA expression levels as in (32) (Figure 1a). In parallel, for each promoter, populations of un-mutagenized cells were subjected to a sham treatment that was identical to the mutagenesis protocol except for exposure to EMS. Ultimately, we captured 148-254 genotypes with unique sets of mutations for each promoter (median: 214, Table S2b) as well as 44-62 genotypes isolated from each sham population (median: 55, Table S2b). For each of these genotypes, we calculated median YFP expression for each of the replicates and then calculated the mean of these medians to represent the expression level for that genotype. Comparing the levels of these means of medians for the sham genotypes among promoters showed differences in the YFP expression driven by each promoter (Figure 1b).

**Figure 1.**
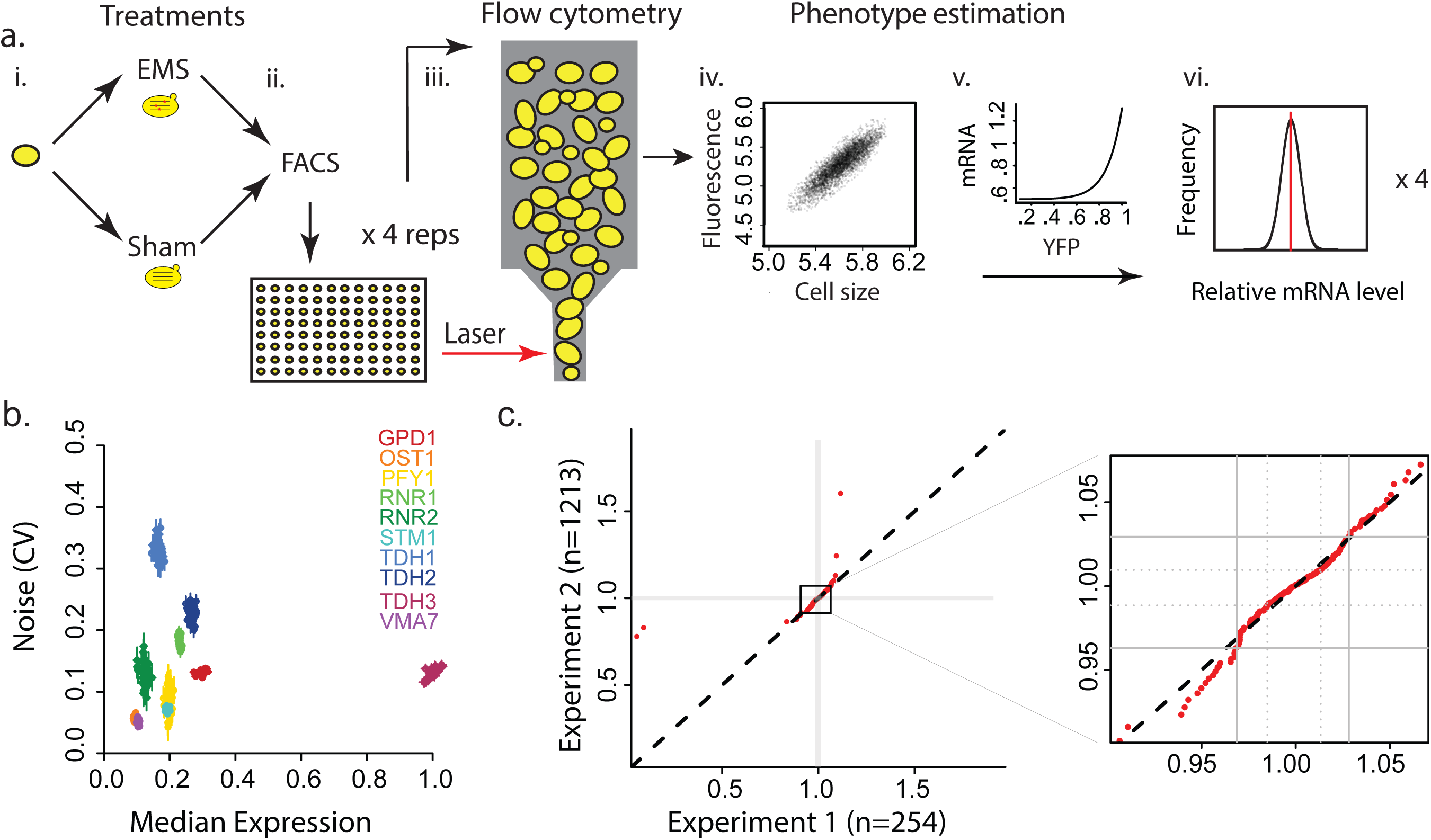
Experimental overview. (a) A schematic describing the experimental design is shown: i. Saturated cultures of cells possessing a promoter sequence driving the YFP reporter were divided in samples for mutagenesis and sham treatments. ii. After mutagenesis and recovery, single cells were isolated using fluorescence activated cell sorting (FACS). iii. Colonies arising from single cells were cultured in quadruplicate and scored using flow cytometry. iv. YFP expression was estimated (BD Accuri C6) as the ratio of fluorescence (log FL1-A: 488 nm laser, 530-30 nm filter) and cell size (log FSC-A). v. YFP fluorescence was converted to estimated mRNA level, adjusting for known non-linearity between YFP fluorescence and mRNA abundance (74). vi. Median mRNA concentrations were calculated for each individual replicate, and then the mean and standard deviation of all replicates were used to characterize expression level for each genotype. (b) Median expression levels (x axis) and expression noise (y axis) are shown for 40 independent unmutagenized genotypes per promoter. Data points represent the mean of 4 replicate flow cytometry measurements per genotype scaled relative to the activity of the *TDH3* promoter. Expression noise was measured as the coefficient of variation (CV) among cells measured within each replicate, and then CV estimates were averaged among replicates. Error bars represent 95% confidence intervals for each genotype. (c) Quantile-quantile plot comparing the distribution of mutational effects for two independent mutagenesis experiments for the TDH3 promoter (x axis: Experiment 1, n=254, y axis: Experiment 2, n=1213) are shown. The second panel enlarges the area corresponding to the central 95% density of both experiments (solid lines: 10^th^ and 90^th^ quantiles, dotted lines: 25^th^ and 75^th^ quantiles). The dashed line on the diagonal represents the hypothesis that the two samples are drawn from the same distribution. A non-parametric Anderson-Darling test fails to reject the null hypothesis that these two samples come from a common population (AD criterion=0.76056, p-value = 0.29).

For each promoter, the collection of EMS-treated genotypes (each of which had a unique set of mutations) was used to estimate a gene-specific distribution of mutational effects on gene expression. With an average of 29 mutations per genotype, we estimate that these distributions reflect the effects of approximately 4000 to 7200 individual mutations distributed across the genome for each promoter analyzed. Mutation accumulation lines in *S. cerevisiae* suggest that ∼10^-3^ spontaneous point mutations arise in each cell each generation (27), implying that at least 2.1 million generations of mutation accumulation would be required to assess the effects of a similar number of mutations for each promoter. For *TDH3*, the distribution of mutational effects observed based on the 254 mutagenized genotypes described was similar to the distribution of mutational effects inferred using an independent collection of >1200 mutagenized genotypes carrying the same *TDH3* reporter gene (24) (Figure 1c). This similarity suggests that the sample sizes used in this study provide reasonable approximations of the underlying mutational distributions.

### Distributions of mutational effects differ in skewness, kurtosis, and dispersion

To determine how mutations alter the expression of each reporter gene, we compared the distributions of expression levels between mutagenized and unmutagenized (sham) genotypes for each promoter (Figure 2). For all promoters, the distribution of expression levels for sham genotypes was symmetrical around the median, whereas, for some promoters, the distribution of expression levels for mutagenized genotypes was asymmetrical (Table S3a,b). This asymmetry suggests that these promoters have biases in the direction of expression changes caused by new mutations. For example, we observed significantly more mutagenized cells with increased than decreased expression for the STM1 promoter relative to the sham median (Figure 2f, n_inc_ = 138, n_dec_ = 83, two-sided exact binomial test, q = 0.002). The *RNR1* promoter showed the opposite pattern with more mutagenized cells showing decreased than increased expression (Figure 2d, n_inc_ = 93, n_dec_ = 132, q = 0.05, test results for all promoters available in Table S2a). In addition, three promoters exhibited departures from symmetry resulting from differences in the magnitude of increased or decreased expression relative to the sham median (Figure 2k; Table S2b): the *TDH1* and *STM1* promoters both exhibited larger increases in expression than decreases (permutation test, TDH1: *P* = 0.006; *STM1*: *P* = 0.031), and RNR2 exhibited larger decreases than increases (*P* = 0.013).

**Figure 2.**
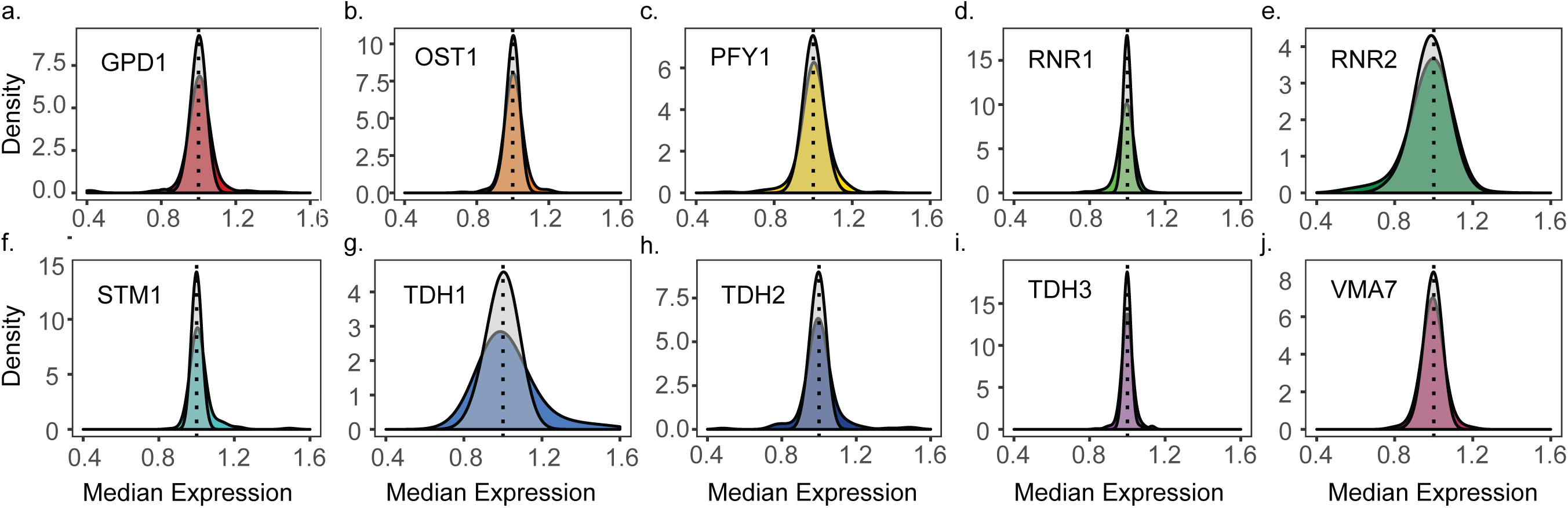
Distributions of expression levels observed for mutagenized and unmutagenized genotypes. (a-j) Distributions of expression among mutagenized (colors) and sham-treated genotypes (overlaid gray transparency) for each promoter. Expression levels (x axis) in both mutant and sham populations are expressed relative to the median of each wild-type expression level (dotted line). The frequency of phenotypes in each population is summarized by promoter as a smoothed density curve (y axis). Data for *TDH3* in (i) was previously published in (24). Sample sizes are listed in Table S3.

To infer distributions of mutational effects from expression observed for mutagenized genotypes, we used variability in the corresponding sham genotypes to control for non-genetic sources of variation in expression. Specifically, for each promoter, we scaled expression of each genotype by the variability in the sham genotypes for that promoter, subtracting the median sham phenotype and then dividing by the standard deviation of sham phenotypes to convert to Z-scores (Figure S1). In this Z-score scale, one unit of change corresponds to a change in expression equivalent to one standard deviation in the population of sham genotypes. We found that at least 60% of mutagenized genotypes (range: 60-86%) showed expression Z-scores within two standard deviations of the sham median for all promoters, suggesting that most new mutations have small effects on a gene’s expression (Figure 3a, Table S4). Prior studies have suggested that non-genetic variability in gene expression can predict mutational effects (8); however, we did not observe this relationship for these ten genes: the variance in Z-scores among EMS-treated cells was not significantly correlated with the variance of expression levels in the corresponding sham population (Spearman’s rho = - 0.27, *P* = 0.448).

**Figure 3.**
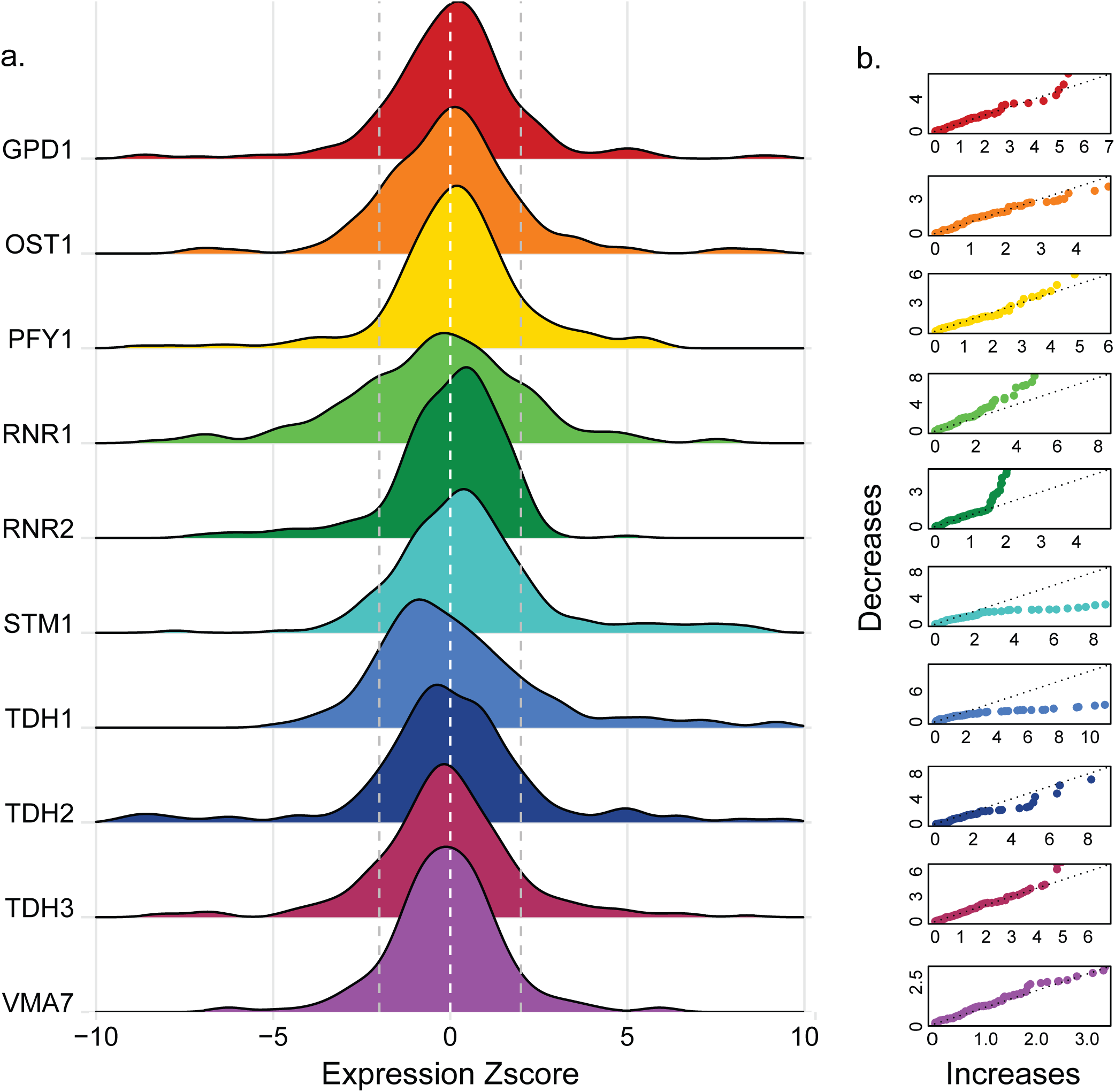
Distributions of mutational effects. (a) Stacked distributions of mutational effects among mutagenized populations. Mutational effects (x axis) are expressed as Z-scores. Dashed lines note the sham median and 2 standard deviations above and below the sham median. (b) The relative frequency of expression changes for a continuous range of magnitudes above and below the sham median are represented as quantile-quantile plots of the magnitude of increases in expression (x axis) by the magnitude of decreases in expression (y axis) for each promoter compared to the median sham phenotype. The dashed line on the diagonal represents the hypothesis that mutational effects generate increases and decreases from the sham median with equal frequency at all effect sizes.

**Figure 4.**
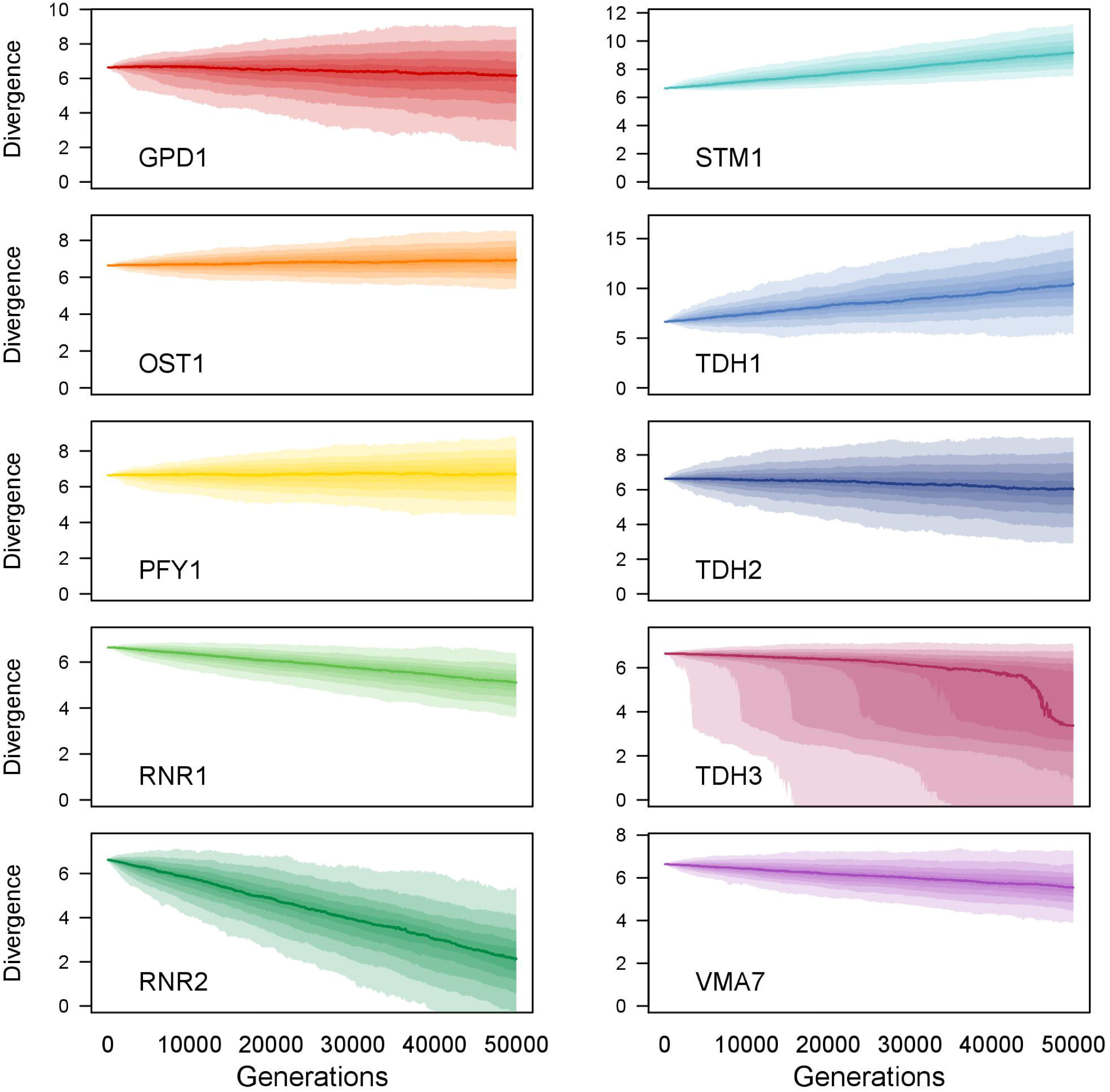
Neutral changes in gene expression predicted by distributions of mutational effects. Evolutionary simulations based on empirical mutational distributions for populations of 1000 individuals evolving in the absence of selection for fifty thousand generations are shown. Population divergence at each generation (x axis) was summarized by the mean population phenotype represented on a log2 scale (y axis). Shaded areas represent the 95% credible intervals for the mean population phenotype at each generation among 500 independent evolutionary trajectories. The darkest line represents the grand median of all independent simulations and lighter shading moving away from the grand median represents quantiles of replicate populations in increments of 10. All populations evolved from a starting population with a mean of log_2_(100), which equals 6.64.

The distributions of Z-scores for EMS-treated genotypes violated normality for all ten promoters (Shapiro-Wilks test, Table S5). To more fully describe the shapes of these distributions, we calculated robust summary statistics that are less sensitive to outliers than the traditional measures used to estimate moments of a distribution (33–35) (Figure S2-S5). Comparing the shapes of distributions of mutational effects among promoters, we observed differences in the centrality (median), dispersion (median-averaged deviation or MAD), skewness (medcouple or MC), and relative frequency of mutations with extreme effects (left/right medcouple or LMC/RMC; Table S4). A principal component analysis (PCA) of summary statistics describing these properties found that the first principal component explained 36% of the variance among promoters and primarily captured skewness and the frequency of extreme increases in expression (Figure S7a and d). A second, independent, principal component explained 30% of the variance and was strongly influenced by median and dispersion (Figure S7b and e). Finally, the third principal component explained 24% of the variance and was influenced by both extreme decreases in expression and dispersion (Figure S7c and f). Because differences in symmetry among promoters dominated these contrasts, we chose to more directly examine skewness for a range of effect sizes using quantile-quantile plots (QQ plots) comparing the magnitude of increases to the magnitude of decreases moving away from the median for each promoter (Figure 3b). By illustrating biases in the direction and magnitude of mutational effects as departures from the 45-degree line, these plots highlight differences among promoters such as the asymmetries described above for STM1, RNR1, TDH1, and RNR2.

Directly comparing the distributions of Z-scores between promoters, we detect 13 of 45 pairwise comparisons in which these distributions of mutational effects differed significantly between promoters (Anderson-Darling test, *P* < 0.05 with BH correction for multiple tests, Figure S6, Table S6). Seven of these 13 cases involved *RNR1*, which possessed an especially unique distribution of mutational effects due to a wide dispersion of Z-scores among mutagenized strains and an overall negative skew, including a small decrease in median compared to its sham population. The distribution of mutational effects for *STM1* was significantly different from five other promoters including *RNR1,* exhibiting biases in the opposite direction from the *RNR1* distribution: *STM1* also showed broad dispersion, but exhibited an overall positive skew with more large effect increases in expression than any other promoter, including a small increase in median compared to its sham population. Other distributions of mutational effects that showed pairwise differences from more than one other promoter included *TDH1* and *RNR2*, which were significantly different from each other and two other promoter distributions (*TDH1*: *RNR1, VMA7*; *RNR2*: *RNR1, STM1*). Like *STM1, TDH1* exhibited a positive overall skew with more density in the right tail of its distribution of mutational effects, but without the right shift in median. *RNR2* was distinct in showing the most humped (aka platykurtic) distribution with the least density in the extreme tails. This RNR2 distribution also showed a slight shift in median towards increased expression despite an overall negative skew.

### Relationships between promoter properties and distributions of mutational effects

As described above, the promoters included in this study were chosen because they vary for properties hypothesized to influence the mutability of gene expression. To determine whether any of these properties might explain the differences in distributions of mutational effects that we observed, we tested for evidence of a significant relationship between the robust summary statistics describing the empirical distributions of mutational effects and the following seven gene properties: (i) expression level of the native gene, (ii) expression noise for the native gene, (iii) presence of a canonical TATA box in the gene’s promoter, (iv) number of *trans*-regulatory factors annotated in YEASTRACT for the gene, (v) density of nucleosome occupancy in the gene’s promoter region, (vi) presence of a duplicate gene in the yeast genome, and (vii) fitness of strains homozygous for a deletion of the gene in rich media. We observed no statistically significant relationship between either dispersion (breadth) measured as MAD or skewness measured as MC and any of the seven properties tested after correction for multiple tests (Figure S8 a-n), but note that our power was limited by the number of genes analyzed. Future studies deeply sampling the effects of mutations on many more genes are needed to better understand how properties of promoters, or the regulatory networks they are embedded in, affect gene-specific distributions of mutational effects for gene expression.

### Predicting neutral expression divergence using distributions of mutational effects

In the absence of empirical data describing the distribution of mutational effects for a specific trait, quantitative genetics models often make the simplifying assumption that the distribution of mutational effects is normally distributed. This assumption is based on the idea that quantitative traits are generally controlled by many loci with small effects (36). If traits are controlled by relatively few loci and/or loci of large effect, as sometimes seems to be the case for gene expression (22, 37), the distribution of mutational effects may be particularly likely to violate normality. Our studies support this observation for gene expression phenotypes, and studies of mutational effects for morphological traits (largely in Drosophila) have also tended to produce non-normal (leptokurtic) distributions of mutational effects with heavy tails (38–46), suggesting they might have similar genetic architecture. Theoretical work has shown that ignoring non-normality of distributions of mutational effects can cause evolutionary models to produce misleading inferences (47–50), but the sparseness of empirical data describing distributions of mutational effects has limited our ability to assess the magnitude of errors caused by these assumptions. To address this knowledge gap, we used our empirical distributions of mutational effects to parametrize models of neutral regulatory evolution for 10 genes and then contrasted the expression divergence predicted by these models with the expression divergence predicted by a more conventional model of regulatory evolution based on normal distributions of mutational effects.

For each promoter, we founded a population of 1000 individuals with expression levels drawn randomly with replacement from the distribution of expression levels for that promoter’s sham population. Each individual had a probability of mutating equal to the spontaneous mutation rate observed in a *S. cerevisiae* mutation accumulation study (1.67 x 10^-10^ per nucleotide per generation (27)), resulting in a new mutation arising in 2 individuals each generation on average. The effect of each mutation was determined by randomly sampling with replacement from the distribution of mutational effects for that promoter and multiplying this effect by the individual’s original expression level, making the simplifying assumption that the distributions of mutational effects stays constant over time. We then randomly sampled 1000 individuals with replacement to populate the next generation. This procedure was repeated for 50,000 generations, calculating mean expression level within the population at each generation. 500 independent simulations were run for each promoter to determine the variation in simulated mean expression levels at each generation.

At the end of 50,000 generations, expression divergence differed dramatically among promoters (Figure a-j). Promoters with a strong positive mutational skew in the distribution of mutational effects like STM1 and TDH1 exhibited large increases in median population expression levels across 500 independent evolutionary trajectories, while promoters with a strong negative mutational skew like RNR1 and RNR2 showed large declines in median population expression levels. Promoters with more symmetric mutational distributions, for example VMA7 and OST1 (Table S4), exhibited less median expression divergence from the original expression level. The TDH3 mutational distribution was also symmetric, but included a few mutants with low expression relative to the rest of the population that caused large step-like decreases in expression when sampled; excluding the 5 lowest values or sampling from a larger collection mutant phenotypes resulted in much more symmetric evolution of TDH3 expression (Figure S9ab). As expected, differences in the distributions of mutational effects among genes were responsible for these outcomes: grand median expression at generation 50,000 was jointly predicted by skewness and weight in the extreme negative tail of the promoter-specific mutational distributions (grand median expression (log transformed to improve normality) at generation 50K ∼ MC + LMC, *F*_2,7_ = 20.15, *P* = 0.001). Similar results were observed using simulations with a population size of 100 instead of 1000 individuals (Figure S9c-l).

We then simulated changes in expression expected for each gene under the more commonly used model of random walks in phenotype space described by Brownian motion. In this model, mutational effects were drawn from a normal distribution centered on the starting expression level of the unmutagenized promoter with variance equal to the variance observed for the empirically-derived distribution of mutational effects for that promoter. We again examined the population means of 500 independent simulations after 50,000 generations. We found that the Brownian motion simulations showed less overall divergence from the starting point than simulations using the empirical distributions of mutational effects, although the extent of difference between the two predictions varied among promoters (Figure 5a, mean expression (log transformed) at generation 50K ∼ Promoter*Model Type: F _*19,9980*_ = 616, *P* < 2.2 x 10^-16^ with significant interaction identified by ANOVA F-test *P* < 2.2 x 10^-16^, see Table S7a). This variation in projected phenotypes from the two types of neutral models could be partially explained by differences in the effects of distribution skewness (MC), dispersion (MAD), and weight in the extreme negative tails on model outcomes (Figure 5b, Table S7b).

**Figure 5.**
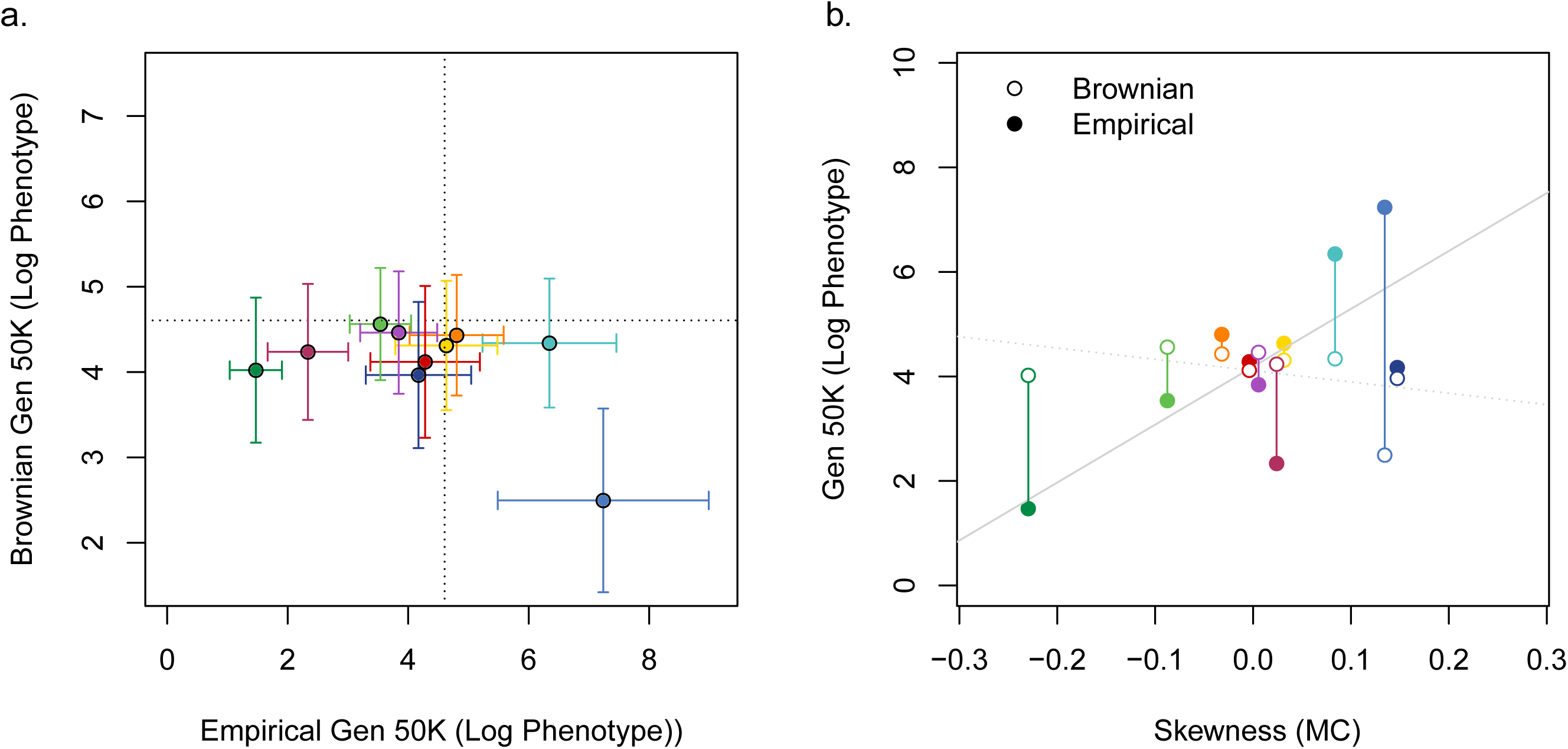
Contrasting neutral models of regulatory evolution. (a) Evolutionary divergence predicted after 50,000 generations under neutral simulations with mutational effects drawn from either a model sampling from the full empirical distributions of mutational effects (x axis) or a Brownian motion model assuming a normal distribution parametrized with the variance of the empirical mutational distributions (y axis). Points represent the grand median expression level among independent evolutionary trajectories for each gene at generation 50,000 under each model on a log scale. Error bars represent 95% credible intervals among 500 replicate populations. (b) Differences in projected expression levels after 50,000 generations of neutral evolution under different mutational models are compared to the observed skewness in the distribution of mutational effects (MC: x axis). Specifically, the grand median phenotype at generation 50,000 is shown on a log scale (y axis) for simulations using the normal (Brownian, open circles) and fully empirical (filled circles) distributions of mutational effects. Skewness predicts simulation results for the empirical (Log Grand Median at Generation 50K ∼ MC, F_1,8_ = 8.64, *P* = 0.018, solid gray line), but not Brownian (Log Grand Median at generation 50K ∼ MC, F_1,8_ = 1.53, *P* = 0.25, dotted gray line) mutational model. Colored bars between points illustrate the differences in simulation predictions when the assumption of normality is relaxed.

To determine how well empirically derived distributions of mutational effects might estimate neutral trait evolution, we compared properties of gene-specific distributions of mutational effects to levels of polymorphism seen for that gene among 22 natural isolates of *S. cerevisiae* (46) because polymorphism is often assumed to primarily reflect neutral processes (45). We observed a significant positive relationship between the degree of dispersion (MAD) in the mutational distribution and expression polymorphism measured as the expression variance among the 22 natural isolates (Figure S10, ExpVar ∼ MAD, *F*_*1,8*_ = 8.18, *P* = 0.021); however, this relationship was driven primarily by *TDH1*, which was an outlier for MAD with respect to the 10 other promoters (Figure S4). Excluding *TDH1* reduced the strength of this correlation and resulted in a p-value that was not statistically significant (ExpVar_noTDH1_ ∼ MAD_noTDH1_, *F*_*1,7*_ = 1.122, *P* = 0.325, dotted line). Skewness of the distribution of mutational effects also failed to significantly predict polymorphism (ExpVar ∼ MC, F = 0.58, *P* =); however, in both cases, we note that our power to detect such relationships is limited by the number of genes studied. Testing for relationships between the effects of mutation and polymorphism more robustly will require similarly deep sampling of mutational effects for many more genes in the yeast genome.

### Modeling distributions of mutational effects and the evolution of gene expression

One of the benefits of assuming a normal distribution of mutational effects is that it simplifies modeling by allowing draws from a well-known continuous distribution rather than a collection of discrete empirical data-points. We therefore sought to identify continuous probability distributions that reflect the shape of the observed empirical distributions of mutational effects better than normal distributions. Distributions of mutational effects for leaf traits in Arabidopsis have previously been described using the family of distributions known as LaPlace distributions (46), also known as double exponential distributions, which have fatter tails than the normal distribution and can be specified in both symmetric (2-parameter) and asymmetric (3-parameter) forms. To determine whether LaPlace distributions fit distributions of mutational effects for gene expression better than normal distributions, we used maximum likelihood to optimize parameters for Gaussian, symmetric LaPlace, and asymmetric LaPlace distributions and then used Bayesian Information Criteria (BIC) to identify the best fitting distribution for each promoter. For all ten promoters, LaPlace distributions were better supported than a normal distribution for the observed distribution of mutational effects (Table S8ab). The VMA7 promoter exhibited similar levels of support for symmetric and asymmetric LaPlace distributions, whereas all other promoters were best described by asymmetric LaPlace distributions. These observations suggest that LaPlace distributions may provide more realistic distributions of mutational effects than normal distributions. They also encourage further investigation into models of regulatory evolution that relax the common theoretical assumption that all loci have equal effects (47–49) and favor models of phenotypic evolution that allow for a high variance in mutational effects (51).

## Conclusion

By studying the effects of thousands of new mutations on expression of individual genes, we have shown how distributions of mutational effects for gene expression differ among genes. Differences observed in the direction and magnitude of mutational effects suggest that some genes may exhibit underlying biases in the expression variation available to selection. In addition, large changes in gene expression were more common than predicted by a normal distribution. For most genes, a null model of neutral expression divergence based on sampling mutations from these distributions predicted greater expression divergence than commonly used quantitative genetic models, suggesting that neutral evolution might explain more variability in gene expression within and between species than often assumed. Challenges for the future include (1) deeply characterizing the distribution of mutational effects for more genes, (2) determining how distributions of mutational effects vary among genetic backgrounds due to epistasis, and (3) identifying features of regulatory networks that can be used to predict a particular gene’s propensity for mutations of a certain effect. Because gene expression is a critical step in the conversion of genotypes to phenotypes, addressing these issues will improve our understanding of the evolutionary processes that generate, maintain, and control variation in complex traits more generally.

## Materials and Methods

More detailed information on the materials and methods used in this study are provided in SI Appendix, Materials and Methods.

### Promoter selection

Promoters from the *GPD1, OST1, PFY1, RNR1, RNR2, STM1, TDH1, TDH2, TDH3*, and *VMA7* genes were selected to represent a diverse range of properties expected to impact mutational variability (8, 18), including expression noise (52), presence of a TATA box motif (53), variation in nucleosome occupancy (54), mutational variance in MA studies, and fitness of homozygous gene deletion strains (55) (Table S1). Maximizing sensitivity for downstream flow cytometry required that all promoters drive relatively high expression, therefore all promoters selected were among the top 15% of highly expressed *S. cerevisiae* genes (56).

### Strain creation

To assay promoter activity, a construct consisting of the promoter region of each focal gene (defined as the intergenic sequence between the start codon of the focal gene and next upstream gene) followed by the Venus YFP coding sequence, the CYC1 terminator and an independently transcribed KanMX4 drug resistance was integrated at the HO locus of a BY4724-derivative strain. Constructs were generated through tailed PCR and transformed via homologous recombination into a strain with a ho::URA3-YFP allele. The genetic background of this strain carried the alleles RME1(ins-308A); TAO3(1493Q) (57) and SAL1; CAT5(91M); MIP1(661T) (58), which decrease petite frequency relative to the alleles of the ancestral BY4724. Data reported include previously published results for TDH3 (24) re-analyzed in a common framework with the nine additional promoters reported here for the first time.

### Mutagenesis

To sample the genome-wide effects of point mutations on promoter activity, we performed random mutagenesis of strains carrying all promoter constructs. Mutagenesis was executed as in (24) using the chemical mutagen ethylmethanosulfonate (EMS), which introduces G/C to A/T point mutations randomly throughout the genome (see SI Appendix for details). Based on canvanine resistance assays performed for *P-TDH1-YFP*, we estimated that ∼29 mutations were introduced per cell with the EMS conditions used (95% percentiles: 24-39), consistent with (24, 59). Sham-treated controls including both a promoter-matched genotype and a *P-TDH3-YFP* construct were maintained in parallel and treated identically with the exception of EMS exposure. Following mutagenesis, single cells from EMS- and sham-treated populations were isolated via FACS and recovered on YPD agar (2% dextrose, 1% yeast extract, 2% peptone, 2% agar) for 48-72 hours at 30 degrees C. Viability of isolated cells was significantly impacted by treatment condition, but not by the genetic background or mutagenesis assay performed (glm quasibinomial models: (Viability ∼ Condition) vs (Viability ∼ Condition + Assay), F test, F_47,55_ = 0.931, *P* = 0.5002; (Viability ∼ Condition) vs (Viability ∼ Condition*Assay), F test, F_15_,_55_ = 0.5656, *P* = 0.9242).

After cell isolation, colonies were transferred from agar to liquid YPD and grown to saturation with shaking, typically 20-24 hours at 30 deg C. Cultures were then used i) to preserve genotypes as glycerol stocks and ii) for initiating cultures to analyze fluorescence. Cultures for analyzing fluorescence were spotted on YPD agar, and 20-24 plate control strains for estimating random experimental effects were interspersed into each row and column. After an additional 48-72 hours growth, colonies were transferred to liquid YPD in 96-well deepwell culture plates, grown for 20-24 hours to saturation with shaking, and scored for fluorescence. A minimum of 4 replicate assays were performed for each plate.

### Phenotyping & data processing

To characterize promoter expression levels, YFP fluorescence driven by the promoter of interest was quantified for all sham-, EMS-treated and plate control genotypes. Fluorescence data was collected on a BD Accuri C6 (488 nm laser and 530/30 optical filter) coupled with an IntelliCyt HyperCyt autosampler. Cultures were diluted in 1x phosphate buffered saline (PBS) to approximately 10^6^ cells/mL directly before scoring. 48 samples were collected per instrument run with gentle vortexing to aerate and re-suspend cells between runs. Separation of run FSC files into well FSC files was performed automatically by Hyperview software (Intellicyt) and manually checked.

Using tools from the flowCore and flowClust libraries (60, 61) and custom scripts, flow cytometry data was analyzed to remove non-cellular debris, events where double cells passed the detector, extreme outliers in cell size or YFP, and correlation between cell size and YFP expression (see SI Materials and Methods). Additionally, because fixed PMT voltages on the Accuri C6 produce non-linearity between fluorophore concentration and fluorescence intensity level (62), this study follows (32) in using a standard curve determined by quantifying RNA abundance via pyrosequencing and fluorescence intensity via flow cytometry for the same samples to scale mRNA abundance estimates appropriately (see SI Materials and Methods). These procedures were performed on a single cell basis for all events that passed quality control thresholds in each sample well. Individual samples were then summarized by calculating median YFP RNA abundance and coefficient of variation (CV estimated as median-averaged deviation/median). Cell size was also summarized as median FSC and FSC MAD. A number of samples were excluded at this stage for phenotypes consistent with high levels of bacterial contamination (small cell size and no YFP expression) or contamination with *P-TDH3* sham controls (YFP expression at the median of P-TDH3 sham for a non P-TDH3 genotype). Any samples with fewer than 1000 single cells passing quality control filters were excluded from analysis. The median and minimum number of cells analyzed per sample are listed by promoter in Table S2.

To account for technical variability across plates, YFP mRNA abundance and FSC metrics for each sample were then normalized to remove random effects due to technical noise arising among instrument runs, plate row position, or plate column position. The power to perform these normalizations came from inclusion of 20-24 control strains in each experimental 96-well plate. Initial experiments were performed using a *P-TDH3-YFP* construct in control positions as in previous work, but when contrasts showed that controls provided more robust correction when matched to the fluorescence phenotype of the construct being corrected, subsequent experiments matched the genetic background of controls to the promoters tested.

To summarize phenotypes estimated for each genotype collected, means and standard deviations were calculated for independent measurements of population medians and CVs across replicated samples. Any individual replicate that was more than 4 MAD outside of other estimates for the same genotype was called as an outlier and excluded from further analysis. Only genotypes with at least 3 independent replicates passing all quality control filters were included in downstream data analysis. These stringent quality control procedures resulted in some differences in the total number of genotypes represented across conditions for different promoter constructs (Table S2b).

### Characterization of mutational spectra

Statistical analyses to characterize mutational spectra across promoters were performed in R (63). Scripts are provided in the supplement (upon publication).

To characterize asymmetry in distributions of expression levels, genotypes were divided into groups with expression greater than (increases) and less than (decreases) the median sham phenotype. We used a binomial test (binom.test with BH multiple test correction) to determine whether increases or decreases were more frequent for each expression distribution and a permutation test (custom function, see SI) to determine whether the mean absolute magnitudes of increases and decreases differed from one another.

The observation that sham-treated genotypes differed in their variability among promoters (Figure S1a) lead us to calculate a Z-score as a metric for capturing the increase in variability due to EMS treatment alone across promoters. To calculate mutational Z-scores, the median of sham phenotype for each promoter was subtracted from each genotype’s YFP expression value and the resulting quantity was divided by the standard deviation among the sham genotypes for that promoter. The resulting metric was centered on 0 and expressed in units representing standard deviations among un-mutated individuals expressing a matched promoter construct (Figure S1b).

To describe differences in the shapes of these distributions of mutational effects on this Z-score scale, we explored a variety of metrics. Summary statistics like sample mean, variance (or standard deviation), skewness, and kurtosis are commonly used to describe distributions, however, these measures have been shown to be particularly vulnerable to influence by outliers (35). More robust statistical measures can be used to describe distribution shape while controlling the impact of outliers in situations where sample size is limiting (33). Here we apply the median-averaged deviation or MAD to characterize distribution breadth in place of standard deviation (64), medcouple to characterize distribution bias in place of skewness (65), and left and right medcouple to characterize the location of distribution tail weight in place of kurtosis. By down-sampling an earlier mutagenesis experiment incorporating more than 1200 mutagenized genotypes, we illustrated that sample median, median-averaged deviation (MAD), medcouple (MC), and left and right medcouple (LMC, RMC) provide more robust and repeatable characterizations of distribution shape (Figure S2-S5). Then, to identify the combination of variables explaining differences between mutational distributions of different promoters, we performed a principle components analysis (66) on robust estimators of moments extracted from Z-score distributions across promoters (Figure S7).

Promoter-specific mutational distributions of Z-scores were visualized by generating stacked density plots using ggplot2 (67). To test for differences in the shapes of distributions of mutational effects between promoters, we applied the non-parametric Anderson-Darling (AD) k-sample test (68, 69) to identify pairwise differences between different promoter mutational distributions, applying the BH procedure to control the false discovery rate in these multiple pairwise tests at 5%.

### Correlation of promoter and population parameters with mutational summary statistics

Promoter properties were collected from the literature (53, 55, 70–72). Linear models predicting summary statistics MC and MAD independently were tested including all promoter property correlates as additive effects. Given the small number of genes involved, the relationship to promoter properties was explored both for continuous metrics and by dividing continuous data into categories of low and high values around the median. A process of model simplification was used to identify predictors explaining variation in MAD or MC, and a BH multiple test correction was performed. Population polymorphism quantified as variance in expression among 22 natural isolates (73) was also tested for a significant relationship with MAD and MC.

### Evolutionary simulations

To illustrate the consequences of the mutational distributions reported here for evolutionary predictions under neutrality, we simulated evolution of an asexual population of individuals randomly sampling mutations impacting the focal promoter and tracked the trajectory of the mean population phenotype over time. For each promoter, populations were initiated by sampling a starting phenotype for each individual (n=1000) from a smoothed version of the sham-treated population. Each generation each individual mutated with a probability determined based on the average estimate of per-generation rate of point mutations (∼1.67*10^10^ bp/generations) detected in mutation accumulation studies (27) multiplied by the *S. cerevisiae* genome size (1.25 x 10^7^ bp). When individuals mutated, they drew a mutational effect size from the distribution of expression levels estimated for EMS-treated genotypes and multiplied their current phenotype by that effect size. Individuals were randomly selected for inclusion in the population each generation. Simulations ran for 50,000 generations and 500 replicate simulations were performed for each promoter. To contrast these results with more typical evolutionary predictions based on an assumption of normally distributed mutational effects, we also ran versions of the simulation from each promoter that were identical except that the mutational effects were drawn from a normal distribution with mean of 0 and variance based on variance among all EMS-treated genotypes for the promoter. For each promoter, evolutionary trajectories were described by the distribution of population means for each generation among independent replicates, and total divergence within each simulation was summarized by the distribution of population means at generation 50,000.

To ask whether differences among models and among genes altered the neutral divergence predicted, we fit a linear model predicting the median phenotype of each replicate population after 50,000 generations based on promoter identity and model identity (Log Median Expression at Generation 50K ∼ Promoter * Model Type) and use an ANOVA F-test to assess fit of full and reduced models. To show how differences in distribution shape related to changes in evolutionary predictions under the two models, we summarized population mean expression among all replicates with a median and then predicted this median expression for each gene based on robust summary statistics calculated for each distribution of mutational effects (Log Grand Median Generation 50K ∼ MAD + MC + LMC + RMC). We performed this procedure separately for the Brownian and full empirical models, and performed model simplification dropping variables with no explanatory power to identify the summary statistics predicting expression divergence in each case.

### Distribution fitting

To identify the family of probability distributions best fitting the empirically-defined distributions of mutational effects, maximum likelihood estimation was performed to identify parameters and log-likelihoods for the Gaussian, symmetric LaPlace, and asymmetric LaPlace distributions given the empirical data. This procedure was performed independently for the EMS-treated populations of each promoter. BIC values were calculated for each fit to identify the best supported model while appropriately penalizing for differences in number of parameters among the distributions (2 for Gaussian and symmetric LaPlace, 3 for asymmetric LaPlace). ΔBIC values were calculated for each fit by subtracting the BIC of the model with the lowest BIC values from all others. We took ΔBIC values greater than 10 as a signal of poor support for a given model.

### Access to Data and Analysis Scripts

All raw flow cytometry data have been uploaded to FlowRepository (FR-FCM-ZYUW) and will be made available on publication.

Supplementary File 1: Primers used to generate and sequence confirm reporter constructs

Supplementary File 2: R code for processing raw .fcs files, normalizing phenotypes by plate controls, filtering outliers, and calculating mean phenotypes by promoter

Supplementary File 3: Layout spreadsheet with experimental metadata linking .fcs files to samples

Supplementary File 4: R code used to contrast mean phenotypes among promoters

Supplementary File 5: R code for evolutionary simulations

Supplementary File 6: Processed data file including mean phenotypes

## Acknowledgements

The authors thank current and past members of the Wittkopp Lab for discussions informing this work, Mark Hill and Brian Metzger provided helpful feedback on the manuscript draft, and the University of Michigan Center for Chemical Genomics and Flow Cytometry Core for access to flow cytometry equipment. We gratefully acknowledge the following funding sources that supported this work: NIH F32 GM115198 to AHD, EMBO ALTF 1114-2012 to FD, and NSF MCB-1021398, NIH 1R35GM118073, and NIH R01 GM108826 to PJW.

## Supplementary Table Legends

**Table S1. Properties of the promoters analyzed.** Promoter properties (mean expression, expression CV, presence or absence of TATA box, nucleosome occupancy score, and fitness of a strain homozygous for a deletion of the gene) are summarized from references (25, 53–55, 70).

**Table S2. Summary of sampling depths for conditions and promoters**. (a) Summary of flow cytometry sampling depth by condition. The number of flow cytometry events collected per sample by condition in unprocessed and processed data are provided as median and 95% intervals for the number of events measured from each well, and the minimum number of events describing any single well after quality control and data processing is provided for each condition. Counts and phenotypes for every sample at each processing step are provided in Supplementary File 1. (b) The number of genotypes used to describe expression distributions by promoter and condition after all data filtering.

**Table S3. Statistics summarizing tests for symmetry of distributions of expression values.** (a) Exact binomial test for differences in the frequency of EMS-treated genotypes showing median expression levels less than and greater than the median sham expression levels. Statistics reported for both sham and EMS populations include the probability of observing a decrease in expression among genotypes and the FDR-adjusted p-value for the test. (b) Permutation test for differences in mean magnitude of decreases versus increases. Statistics reported for both sham and EMS populations include the mean difference in magnitude between increases and decreases in expression from the median for 10 thousand samples matched in sample size, and the p-value testing the probability that the difference between increases and decreases was larger than a random sample of measured differences in expression from the median.

**Table S4. Summary statistics for the distribution of mutational effects for each promoter.** Summary statistics reported include sample sizes, median, median-averaged deviation (MAD), skewness as characterized by medcouple (MC), weight in left extreme tails as characterized by left medcouple (LMC), and weight in right extreme tails as characterized by right medcouple (RMC) on both the native scale (scaled relative to 0 expression and the sham median at 1) and on a Z-score scale (substracting the sham median and dividing by the sham MAD). The proportion of EMS-treated genotypes falling within 1-, 2-, and 3 standard deviations of the sham median are listed.

**Table S5. Results of the Shapiro-Wilks test for normality for distributions of mutational effects**. The tests were performed on the EMS-treated genotypes promoters expressed on a Z score scale and the test statistic and FDR-adjusted p-value are reported.

**Table S6. Results of the K-sample Anderson-Darling (AD) non-parametric test for similarity between mutational distributions.** The AD test was used to test the null hypothesis that each pair of mutational distributions expressed on a Z-score scale was drawn from the same underlying true distribution. This test of distribution similarity identified pairwise differences among distributions of mutational effects shown in Figure S6 (red: pairwise differences at *P* < 0.05). Test statistic and FDR-adjusted P-values are reported for each pairwise comparison.

**Table S7**. **Results predicting population divergence in evolutionary simulations based on promoter identity, mutational model, or the shape of distributions of mutational effects.** (a) Mean population divergence was log transformed to improve normality for statistical analyses. Population divergence in evolutionary simulations across all 500 trajectories for all promoters was best predicted by a model including an interaction between the promoter identity and mutational model type (ANOVA test of the full interaction vs a reduced additive model, *F* = 828, *P* <2.2 x 10^-16^). Regression coefficients and post-hoc *P*-values for the multiple regression test (Log Mean Population Phenotype after 50K Generations ∼ Promoter * Model Type) are reported in Table S7. (b) Comparisons of the most parsimonious models predicting grand median expression levels for both Brownian motion simulations and for simulations drawing mutational effects from the full empirical distribution from robust summary statistics are reported including models, regression coefficients, and post-hoc *P*-values.

**Table S8. Support for fits of different probability distributions to the empirical distributions of mutational effects.** The fit of different probability distributions was assessed based on Bayesian Information Criterion difference scores (δBIC) describing support for Gaussian, symmetric LaPlace, and asymmetric LaPlace model parametrizations. δBIC difference scores are expressed relative to the model with the lowest log-likelhood (highlighted with shading in this table). Models separated by 10Parametrizations of model with the most support including choice of model, minus log-likelihood, parameter estimates and BIC calculated.

## Supplemental Figure Legends

**Figure S1. Effects of converting expression distributions to Z-scores.** Differences in expression variation among un-mutagenized sham populations on the (a) median expression scale and (b) mutational Zscore scale represented as violin plots of smoothed density values are shown.

**Figure S2. Comparing robustness of metrics that estimate the central tendency of a distribution inferred using different sample sizes**. Density distributions of means (left) and medians (right) calculated from 1000 replicate samples of 50, 100, 200, 400, 600, 800, and 1000 values drawn from the same underlying distribution are shown. Dotted lines show the value of each parameter calculated from all 1200 datapoints in the distribution from which the samples were drawn. These data show that the median provides a more reliable measure of the central tendency of the underlying distribution, especially for smaller sample sizes.

**Figure S3. Comparing robustness of metrics that estimate dispersion of a distribution inferred using different sample sizes**. Density distributions of standard deviation (StDev, left) and median-averaged deviation (MAD, right) calculated from 1000 replicate samples of 50, 100, 200, 400, 600, 800, and 1000 values drawn from the same underlying distribution are shown. Dotted lines show the value of each parameter calculated from all 1200 datapoints in the distribution from which the samples were drawn. These data show that the MAD provides a more reliable measure of the dispersion of the underlying distribution, especially for smaller sample sizes.

**Figure S4. Comparing robustness of metrics that estimate asymmetry of a distribution inferred using different sample sizes**. Density distributions of skewness (left) and Medcouple (right) calculated from 1000 replicate samples of 50, 100, 200, 400, 600, 800, and 1000 values drawn from the same underlying distribution are shown. Dotted lines show the value of each parameter calculated from all 1200 datapoints in the distribution from which the samples were drawn. These data show that Medcouple provides a more reliable measure of the asymmetry of the underlying distribution, especially for smaller sample sizes.

**Figure S5. Comparing robustness of metrics that estimate weight in the extreme tails in a distribution inferred using different sample sizes**. Density distributions of kurtosis (left), medcouple for the left half of the distribution (LMC, middle), and medcouple for the right half of the distribution (RMC, right) calculated from 1000 replicate samples of 50, 100, 200, 400, 600, 800, and 1000 values drawn from the same underlying distribution are shown. Dotted lines show the value of each parameter calculated from all 1200 datapoints in the distribution from which the samples were drawn. Note that LMC and RMC measure distribution weight in the peak vs tails of a distribution independently on either side of the median, while kurtosis combines this information in a single measure. LMC and RMC each provide a more self-consistent estimate at every sample size than kurtosis, which shows a multimodal distribution of estimates that increases with sample size..

**Figure S6. Comparing distributions of mutational effects among promoters.** Quantile-quantile plots contrasting mutational distributions on a Z-score scale for each pair of promoters are shown. The dashed diagonal line represents the hypothesis that samples were drawn from the same underlying mutational distribution. Pairwise comparisons achieving significance (*P*<0.05 on non-parametric Anderson-Darling test with BH correction, Table S6) are plotted with red points.

**Figure S7. Principal Components Analysis (PCA) comparing shapes of mutational distributions.** (a-c) PCA factor scores for different promoters and (d-f) component loadings for summary statistics comparing PC1 vs PC2, PC2 vs PC3, and PC1 vs PC3 are shown. (g-i) Stacked distributions of mutational effects ordered by promoter scores on PC1 (g), PC2 (h), and PC3 (i) are shown.

**Figure S8. Do promoter properties predict distributions of mutational effects?** Dispersion (MAD, a-g) and asymmetry (MC, h-m) of distributions of mutational effects are compared for promoters with high and low native expression level (a,h), high and low expression CV (b,i), the absence (TATA-) or presence (TATA+) of a canonical TATA box (c,j), high and low numbers of annotated regulators (d,k), the absence (N) or presence (Y) of duplicate in the yeast genome (e,l), nucleosome occupancy (f,m), and competitive fitness of a strain with homozygous deletion of the gene of interest (g,n). Binary categories of continuous variables like expression level, CV, or numbers of regulators were formed by grouping genes with values above or below the median. Continuous metrics were also explored for quantifying TATA box (# of differences from a canonical TATA sequence) and nucleosome occupancy, but results were consistent whether continuous and categorical metrics of promoter properties were used (data not shown). The black dotted lines in panels f, g, m, and n show the best fit regression lines, though no relationships were identified that remained statistically significant at a *P* < 0.05 level after correction for multiple testing..

**Figure S9. Neutral changes in gene expression predicted by distributions of mutational effects.** (a,b) Evolutionary simulations based on empirical mutational distributions for the gene TDH3 using different samples of mutants to estimate mutational distributions for populations of 1000 individuals evolving for 50,000 generations are shown, including (a) a subset of the TDH3 mutants analyzed in this manuscript but excluding the mutants with 5 lowest expression levels, and (b) a larger sample of 1400 mutants described in (24). (c-l) Evolutionary simulations based on empirical mutational distributions for populations of 100 individuals evolving in the absence of selection for fifty thousand generations are shown. Population divergence at each generation (x axis) was summarized by the mean population phenotype represented on a log2 scale (y axis). Shaded areas represent the 95% credible intervals for the mean population phenotype at each generation among 500 independent evolutionary trajectories. The darkest line represents the grand median of all independent simulations and lighter shading moving away from the grand median represents quantiles of replicate populations in increments of 10. All populations evolved from a starting population with a mean of log_2_(100), which equals 6.64.

**Figure S10. Comparing mutational variability to levels of expression polymorphism in *S. cerevisiae*.** Correlation between variability of mutational distributions (MAD) and expression polymorphism among 22 natural isolates of *S. cerevisiae* described by (73) for the 10 genes examined in this study, characterized as variance in expression level among natural isolates. The positive relationship between mutational variability (MAD) and expression polymorphism (ExpVar ∼ MAD, F_1,8_ = 8.18, *P* = 0.021, solid line) is highly influenced by the outlier TDH1 (ExpVar_noTDH1_ ∼ MAD_noTDH1_, F_1,7_ = 1.122, *P* = 0.325, dotted line).

