## Supplemental tables, figures, files, and methods for "Empirical measures of mutational effects define neutral models of regulatory evolution in *Saccharomyces cerevisiae*": SI Appendix Materials and Methods 01302018.docx

### Creation of reporter constructs

Reporter constructs were inserted at the *ho* locus in a S288C-derived haploid MATα genetic background described in (1). Briefly, this strain contains alleles of SAL1, CAT5 and MIP1 that decrease the penetrance of the petite phenotype (2) and alleles of RME1 and TAO3 that increase sporulation efficiency in diploids (3). To create the different reporter constructs, we first PCR amplified the full promoter sequence of each gene to be studied from the end of the previous coding sequence to the start codon of the gene of interest. This PCR was performed using oligonucleotide primers with tails possessing a minimum of 40 bp of homology with the 5’ region of the *ho* locus and 40 bp of homology into the coding sequence of YFP (sequences available in Supplementary File 1). The resulting construct was transformed into a strain of the genetic background described containing a ho::pCORE-UH-YFP-Kan allele and counter-selected on 5-FOA with screening for YFP fluorescence (4). Successful transformation was confirmed with sequencing of the full locus. A minimum of two independent transformants per reporter construct were retained and screened for YFP fluorescence on the BD Accuri C6 to confirm consistent effects on expression among independent transformants.

### Mutagenesis

To sample the genome-wide effects of point mutations on promoter activity, we performed random mutagenesis of strains carrying all promoter constructs. Mutagenesis was executed as in (1) using the chemical mutagen ethylmethanosulfonate (EMS). EMS introduces G/C to A/T point mutations randomly throughout the genome, paralleling the most frequent spontaneously occurring single nucleotide mutation in sequenced mutation accumulation lines (5, 6). Briefly, strains to mutagenize were thawed from -80 °C on YPG agar (10 g/L yeast extract, 20 g/L peptone, 5% vol/vol glycerol, 20 g/L agar) for at least 48 hours incubated at 30 °C, then used to inoculate 5 mL cultures of YPD (10 g/L yeast extract, 20 g/L peptone, 20 g/L dextrose) incubated for 20-24 h at 30 °C on a rotor for mixing. To perform mutagenesis, a 2 mL aliquot of saturated culture (~10^7^ cells/mL) was washed twice in 5 ml of H2O, and then suspended in 2 mL of sodium phosphate buffer (0.1 M, pH 7) before division into two 1 mL aliquots in 15 mL microcentrifuge tubes. In total, 10 μl of EMS (99%; Acros Organics) was added to one sample (EMS treatment), but not the other (sham treatment), and both samples were sealed with parafilm and incubated at room temperature on a rotor for mixing. After 45 minutes of incubation, both samples were treated with 1 mL of EMS-neutralizing sodium thiosulfate (5%). Cells were pelleted and suspended again in 1 ml of sodium thiosulfate (5%), twice in 1 ml of H2O, and finally suspended in 1 ml of YPD. Cells were diluted 1 in 32 (0.125 ml cell suspension in 3.875 ml YPD) in a 15-ml culture tube and grown overnight to saturation at 30 °C on a rotor for mixing, when a 1 in 32 dilution was repeated and cells were incubated to saturation again for an additional 24 h. This dilution series was estimated to produce ∼10 generations of recovery after EMS treatment. In addition to EMS and sham-treated samples, an additional sham treatment of the promoter construct P-TDH3-YFP was performed in parallel with EMS and sham treatments of all other promoter constructs as a common control. Mutagenesis and sorting was performed in batches of two promoter constructs (EMS and sham treatments) and one P-TDH3-YFP sham treatment at a time.

### Cell isolation & culture for scoring

 After recovery, genotypes were isolated as single cells on YPD agar plates in a 96-position array using FACS (BD FACS Aria III, University of Michigan Flow Cytometry Core). To perform sorting, all samples were diluted in PBS buffer: 0.4 mL of saturated culture (∼ 10^7^ cells/mL) in 2 mL. Cells were sorted at a flow rate of ∼2,000 cells/second. Debris and cell doublets were discriminated based on forward scatter and width in the FACSDiva software and excluded from the population sorted. In the initial round of experiments for each promoter, populations of EMS-treated (n=240), sham-treated (n=64), and P-TDH3-YFP sham treated (n=64) cells sorted were selected randomly with respect to fluorescence phenotype. All treatment conditions for a single promoter construct were represented on each sorted plate and evenly distributed across rows and columns. Eight plates were sorted per promoter on to solid YPD agar plates. In addition, extra cells from each condition were isolated for replacing any missing positions due to failures in sorting or cell death. After sorting, cells were grown into colonies by incubating plates for 48-72 h at 30° C. Survival was scored and any missing colonies were noted. Using a V&P Scientific pin tool, colonies were transferred from plates to a 96-well deepwell culture plate containing 0.4 mL YPD per well and a sterile 3 mm glass bead to provide greater mixing and aeration on shaking . Plates were grown with shaking at 220 rpm to saturation, typically 20-24 hours. An additional 2mL culture of an ancestral wild type (wt) strain and a non-fluorescent control were also grown to saturation on a rotor. After growth to saturation, cultures in 96-well plates were then used i) to preserve genotypes as glycerol stocks at -80° C,  and using a pintool ii) to inoculate YPG plates for initiating cultures to analyze fluorescence . The saturated wt strain was then distributed across a clean 96-well plate at control positions in each row and column and then, using a pin tool, spotted into empty positions onto the YPG agar plates already inoculated with sham- and EMS-treated cells. Colonies on YPG plates were grown at 30° C. After 48-72 hours growth, colonies were transferred with a pin tool to 0.4 mL YPD in 96-well deepwell culture plates. These plates were grown for 20-24 hours to saturation and scored for fluorescence.  A minimum of 4 replicates were inoculated for each unique plate.

### Flow cytometry data collection, processing, & analysis

To characterize promoter expression levels, YFP fluorescence driven by the promoter of interest was quantified for all sham-, EMS-treated and control genotypes. Fluorescence data was collected on a BD Accuri C6 (488 nm laser and 530/30 optical filter) coupled with an IntelliCyt HyperCyt autosampler. After 20-24 hours growth in YPD, saturated cultures were diluted in 0.4 mL filter-sterilized 1x phosphate buffered saline (PBS, pH=7) to approximately 10^6^ cells/mL directly before scoring. Imposing only a lowpass filter to remove small particles (FSC-H< 5 x 10^4^), roughly 20,000 events were collected per culture in runs of 4 rows to avoid exceeding the instrument limit of 1M events/file. Plates were mixed thoroughly with gentle vortexing (Fisherbrand vortexer setting 3-5) immediately before each scoring run to aerate cultures and avoid cell settling. Separation of run FSC files into well FSC files was performed automatically by Hyperview software (Intellicyt) and manually checked.

To process flow cytometry data, we used R packages (7) *flowCore* (8) and *flowClust* (9) with custom modifications based on similar analyses performed in (1, 10, 11). Scripts are provided in Supplementary File 2 with metadata in Supplementary File 3. Briefly, fluorescence and cell size ratios were used to distinguish cell populations of interest from debris and doublets and to calculate a single cell fluorescence metric scaled by cell size (log(YFP)/log(FSC.Area)). Events were retained if: i) they fell within a a set of broadly inclusive hard gates (4 < log(FSC.Area) < 6.5; 4.85 < log (FSC.Height) < 6.8; 0.87 <  log(FSC.Area) / log (FSC.Height) <0.94; 20  < Width < 80), ii) they clustered with other single cells in a PCA used to distinguish singlets from doublets based on a combination of log (FSC.Area) and log (FSC.Area) / log(FSC.Height) variables, and iii) they fell within 4 median averaged deviations or MAD (a more robust measure of standard deviation) of the population median. Samples with fewer than 1000 events after all filtering steps were discarded. log(YFP)/log(FSC.Area) fluorescence metrics were rotated around a log(FSC.Area) centroid as in (1) to eliminate any lingering effect of cell size . Additionally, because fixed PMT voltages on the Accuri C6 produce non-linearity between fluorophore concentration and fluorescence intensity level (12) , this study follows (11) in using a standard curve determined by quantifying RNA abundance via pyrosequencing and fluorescence intensity via flow cytometry for the same samples to scale mRNA abundance estimates appropriately.  The function used to perform this calibration was  log(*y*) = *a* log(*x*) + *b* where *x* represents fluorescence intensity level and *y* represents fluorophore concentration (12). Using the empirical relationship of YFP intensity to mRNA abundance to derive parameters  *a* = 10.469 and *b* = -9.586  in this calibration function, we were able to estimate quantitative changes in YFP mRNA abundance across a range of YFP intensity levels (11). These procedures were performed on a single cell basis for all events that passed quality control thresholds in each sample well. Individual samples were then summarized by calculating median YFP RNA abundance and by calculating different estimates of between cell variability including YFP RNA abundance median-averaged deviation (MAD, a robust estimator for standard deviation), coefficient of variation (CV as MAD/median), and Fano factor (Fano as MAD^2^/median). Cell size was also summarized as median log FSC.Area and MAD log FSC.Area.  A number of samples were excluded at this stage for phenotypes consistent with high levels of bacterial contamination (tiny cell size and no YFP expression) or contamination with P-TDH3 sham controls (YFP expression at the median of P-TDH3 sham for a non P-TDH3 genotype).

To account for technical variability across plates, YFP mRNA abundance and FSC metrics for each sample were then normalized to remove random effects due to technical noise arising among instrument runs, plate row position, or plate column position. The power to perform these normalizations came from inclusion of 20-24 control strains in each experimental 96-well plate. Initial experiments were performed using a P-TDH3-YFP construct in control positions as in previous work, but when contrasts showed that wt controls provided more robust correction when matched to the fluorescence phenotype of the construct being corrected, subsequent experiments matched the genetic background of controls to the promoters tested (see Supplementary File 6 for control genotypes by experiment).

After normalization, initial YFP phenotypes were calculated on two different scales: i. a promoter-specific YFP scale was calculated by subtracting a non-fluorescent control and dividing by the median of all sham-treated controls with the same promoter construct, and ii. a TDH3 scale was calculated by subtracting a non-fluorescent control and dividing by the median of all sham-treated controls with a P-TDH3-YFP construct in the same experiment. As a result, samples can be expressed on a scale where 0 represents no expression of the YFP construct and 1 represents YFP expression relative to an un-mutated promoter construct or on a scale relative to an un-mutated P-TDH3 construct for comparison across promoters. Figure 1b shows results on the TDH3 scale and otherwise all results are presented on the promoter-specific scale (or a transformation of the promoter-specific scale to Z-scores as detailed below).

To summarize phenotypes estimated for each genotype collected, means and standard deviations were calculated for independent measurements across replicated samples (normalized YFP abundance summarized by medians, MAD, CV, Fano for both promoter-specific and P-TDH3-specific scales). Any replicate that was more than 4 MAD outside of other estimates for the same genotype was called as an outlier and excluded from further analysis. Only genotypes with at least 3 independent replicates passing all quality control filters were included in downstream data analysis. These stringent quality control procedures resulted in some differences in the total number of genotypes represented across conditions for different promoter constructs (Table S3).

### Characterization of mutational spectra

Statistical analyses to characterize mutational spectra across promoters were performed in R.  Scripts are provided in Supplementary File 4 and processed data are provided in Supplementary File 6.

We visualized promoter-specific mutational distributions by generating histograms, density plots, and violin plots by condition for each promoter using base R (7) and tidyverse ggplot2 (13). The observation that sham-treated genotypes differed in their variability among promoters lead us to calculate a Z-score as a metric for capturing the increase in variability due to EMS treatment across promoters. To calculate mutational Z-scores, the mean of each genotype in the promoter-specific YFP scale had the median of all the promoter-matched sham-treated genotypes subtracted from it and the resulting quantity was divided by the MAD among sham genotypes. The resulting metric was centered on 0 and expressed in units representing median-averaged deviation among un-mutated individuals expressing a matched promoter construct. Z-score distributions were visually examined using quantile-quantile (QQ) plots, including QQ-plots comparing distributions across promoters as well as well QQ-plots comparing the magnitude of expression changes increasing and decreasing from the median of the sham within each promoter.

To test for departures from symmetry, we performed a) binomial tests contrasting the number of EMS-treated genotypes with phenotypes greater and less than the median sham phenotype and b) permutation tests contrasting the magnitude of differences from the median sham phenotype among EMS-treated genotypes with phenotypes greater and less than the median sham phenotype. To test for differences in the shapes of distributions of mutational effects between promoters, we applied the non-parametric Anderson-Darling (AD) k-sample test (14) from the *kSamples* package (15) to identify pairwise differences between different promoter mutational distributions by assessing goodness-of-fit between two empirical cumulative distribution functions at a time. We applied the Benjamini-Hochberg procedure to control the false discovery rate in these multiple pairwise tests at 5%.

Differences in distribution shape were quantitatively described by calculating summary statistics describing distribution centrality, dispersion, symmetry, and tail weight. Classic summary statistics describing these features of distributions include mean, standard deviation, skewness, and kurtosis respectively, however, these moments are particularly vulnerable to the presence of outliers even at moderate sample sizes (16). Robust summary statistics including median, median-averaged deviation, medcouple, and left and right medcouple are formulated to capture similar information about distribution shape while decreasing the influence of outliers (17). We confirmed that these robust summary statistics provided more precise estimates in this context by resampling 1200 mutagenized genotypes carrying the P-TDH3-YFP construct generated in a previous experiment at a variety of sample sizes, calculating summary statistics on the population using the *e1071* package (18) and *robustbase* package (19), and repeating this sampling 1000 times to generate distributions of summary statistics estimated at different sample sizes. Robust summary statistics exhibited desirable behavior including unimodal distributions of estimates, less variance in estimation, and greater reliability at every sample size (Figures S2-S5), and are used throughout the remaining analyses. Principle components analysis of robust summary statistics with the *FactoMiner* package (20) was used to identify the combination of variable explaining differences between mutational distributions of different promoters.

### Evolutionary simulations

To illustrate the consequences of the mutational spectra reported here for evolutionary predictions under neutrality, we simulated evolution of an asexual population of individuals that randomly sample mutations impacting the focal promoter and tracked the trajectory of the mean population phenotype over time (Supplementary File 5). For each promoter, populations were initiated by sampling a starting phenotype for 1000 individuals from a smoothed version of the sham-treated population with an arbitrary mean phenotype value of 100. Each generation each individual mutated with a probability determined based on the average estimate of per-generation rate of point mutations (~1.67*10^10^ bp/generations) detected in mutation accumulation studies (6) multiplied by the *S. cerevisiae* genome size (1.25 x 10^7^ bp). When individuals mutated, they drew a mutational effect size from the distribution of EMS-treated genotypes for the focal promoter and multiplied their current phenotype by that mutational effect size.  Individuals were randomly selected for inclusion in the population each generation with replacement. Simulations ran for 50,000 generations and 500 replicate simulations were performed for each promoter. To contrast these results with more typical evolutionary predictions based on an assumption of normally distributed mutational effects, we also collected data for each promoter from a Brownian motion version of the simulation that was identical except that the mutational effects were drawn from a normal distribution with mean of 1 and variance based on variance among all EMS-treated genotypes for the promoter.
