## Supplemental tables, figures, files, and methods for "Empirical measures of mutational effects define neutral models of regulatory evolution in *Saccharomyces cerevisiae*": Supp Figs Merged 02152019.pdf

Supplementary Figure 1

a.

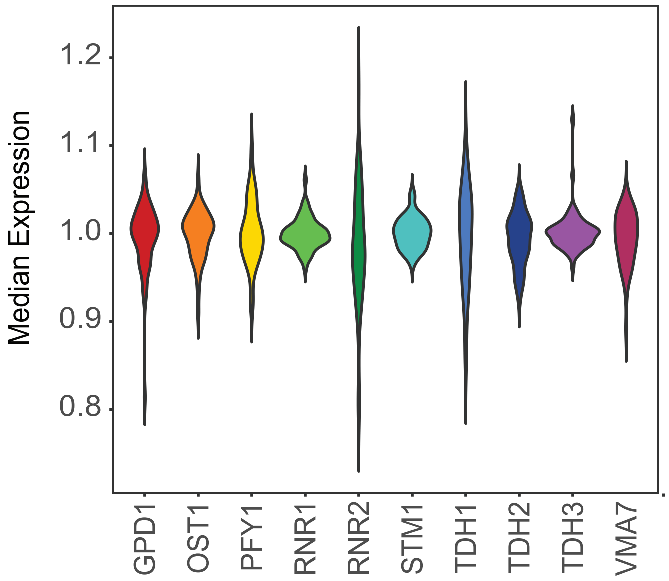


b.

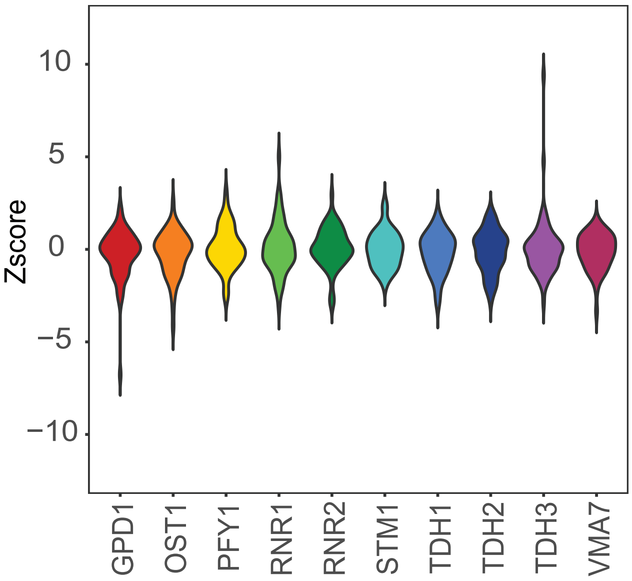

Supplementary Figure 2

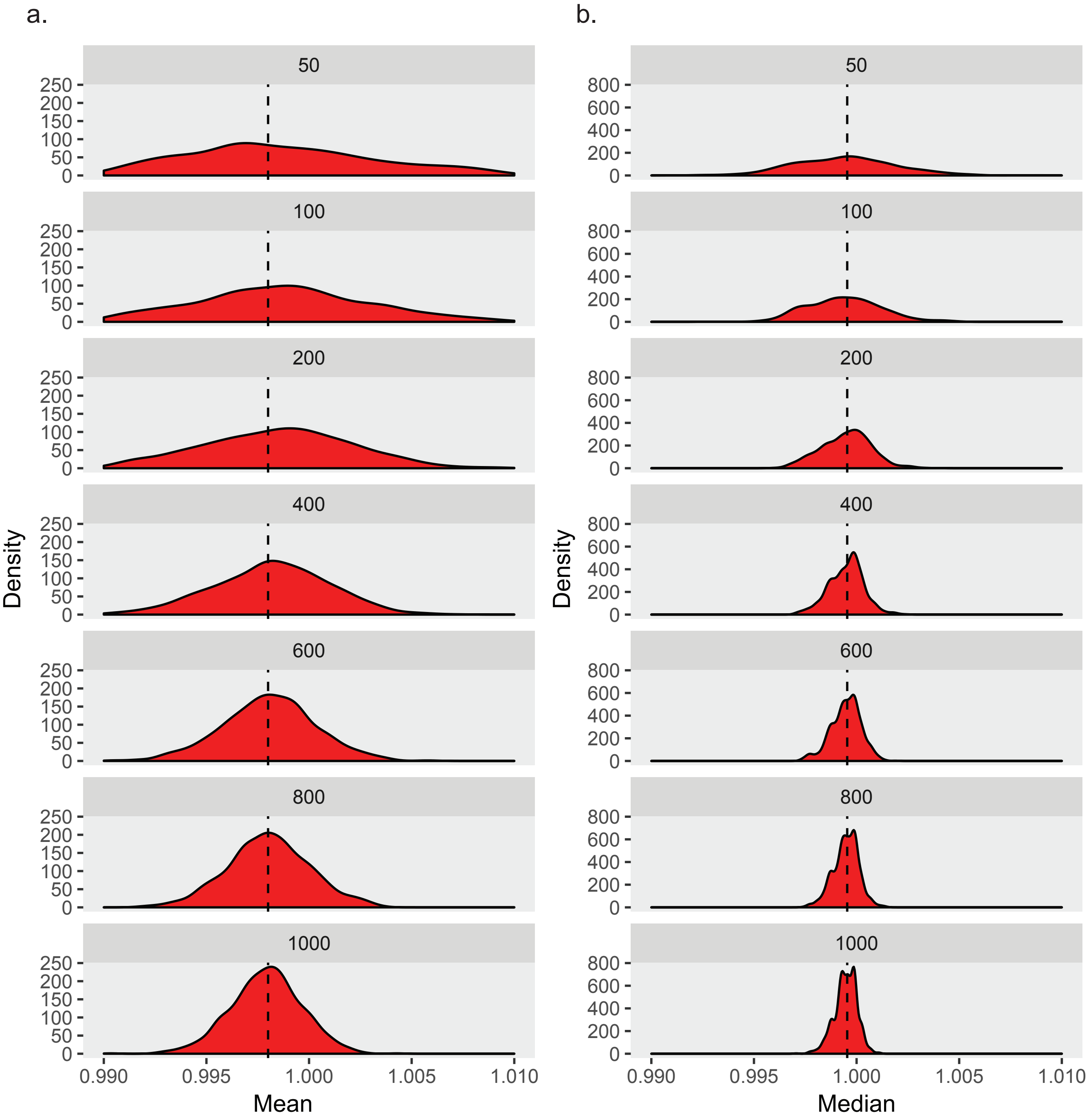

Supplementary Figure 3

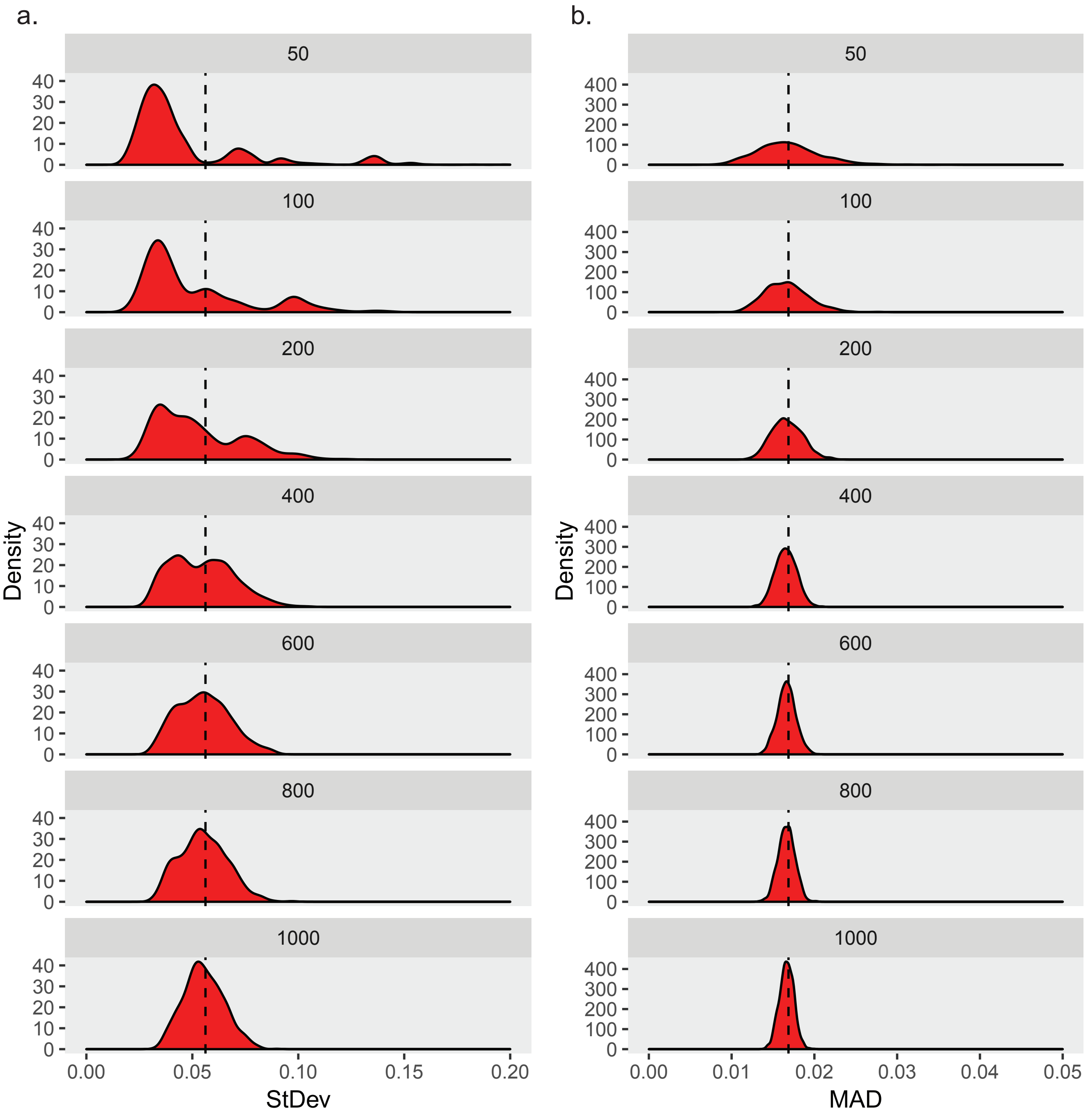

Supplementary Figure 4

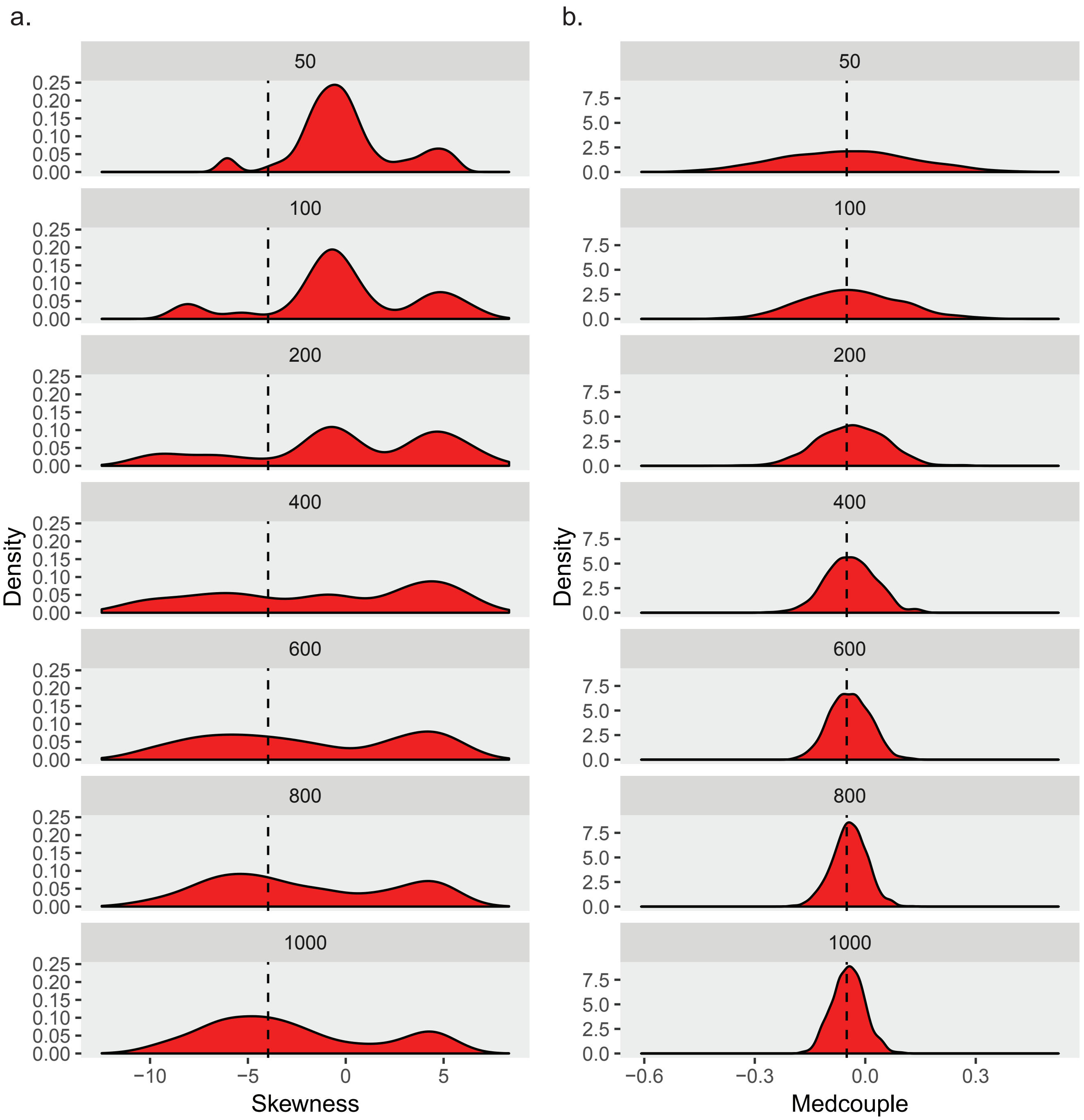

Supplementary Figure 5

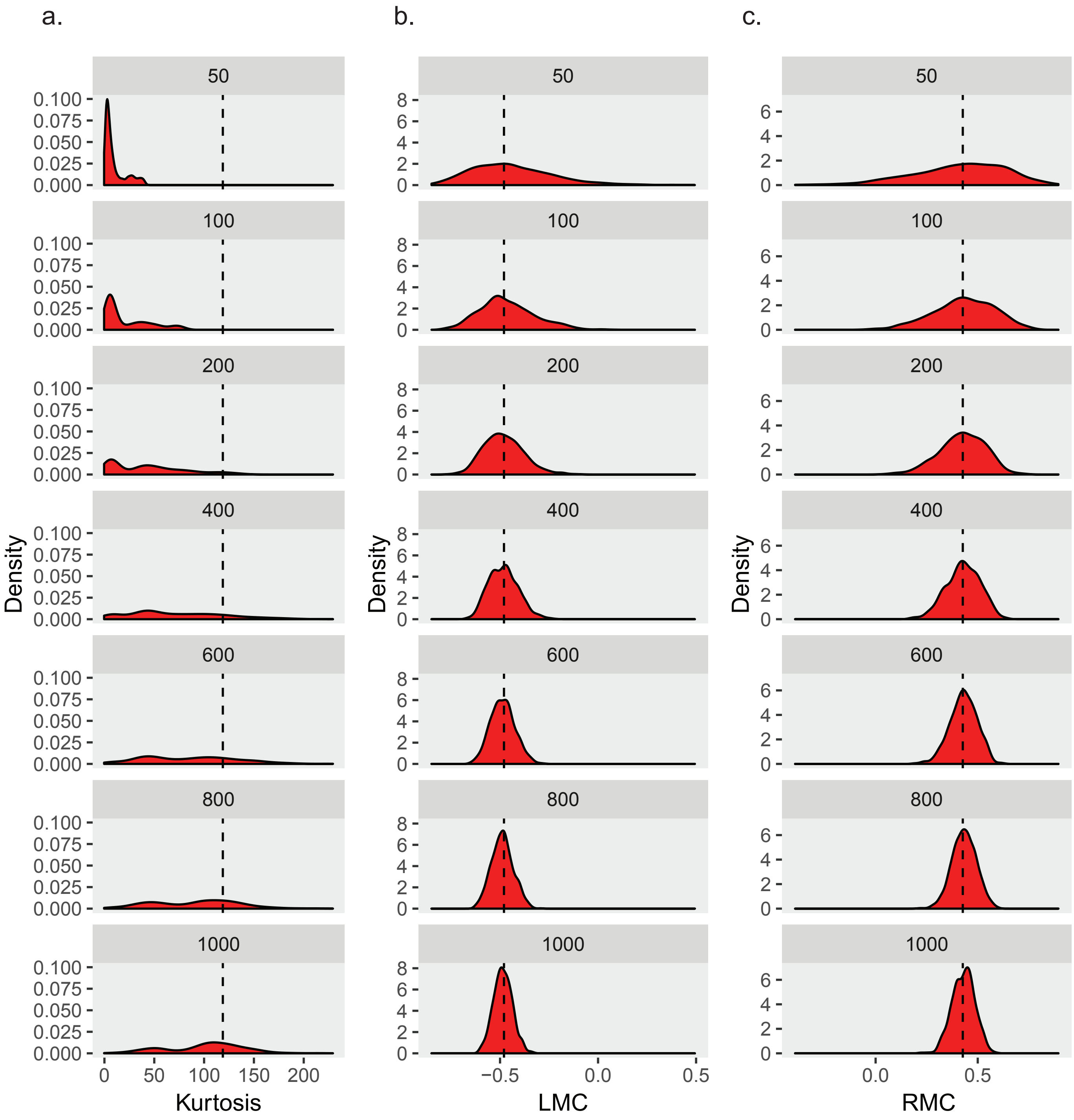

Supplementary Figure S6

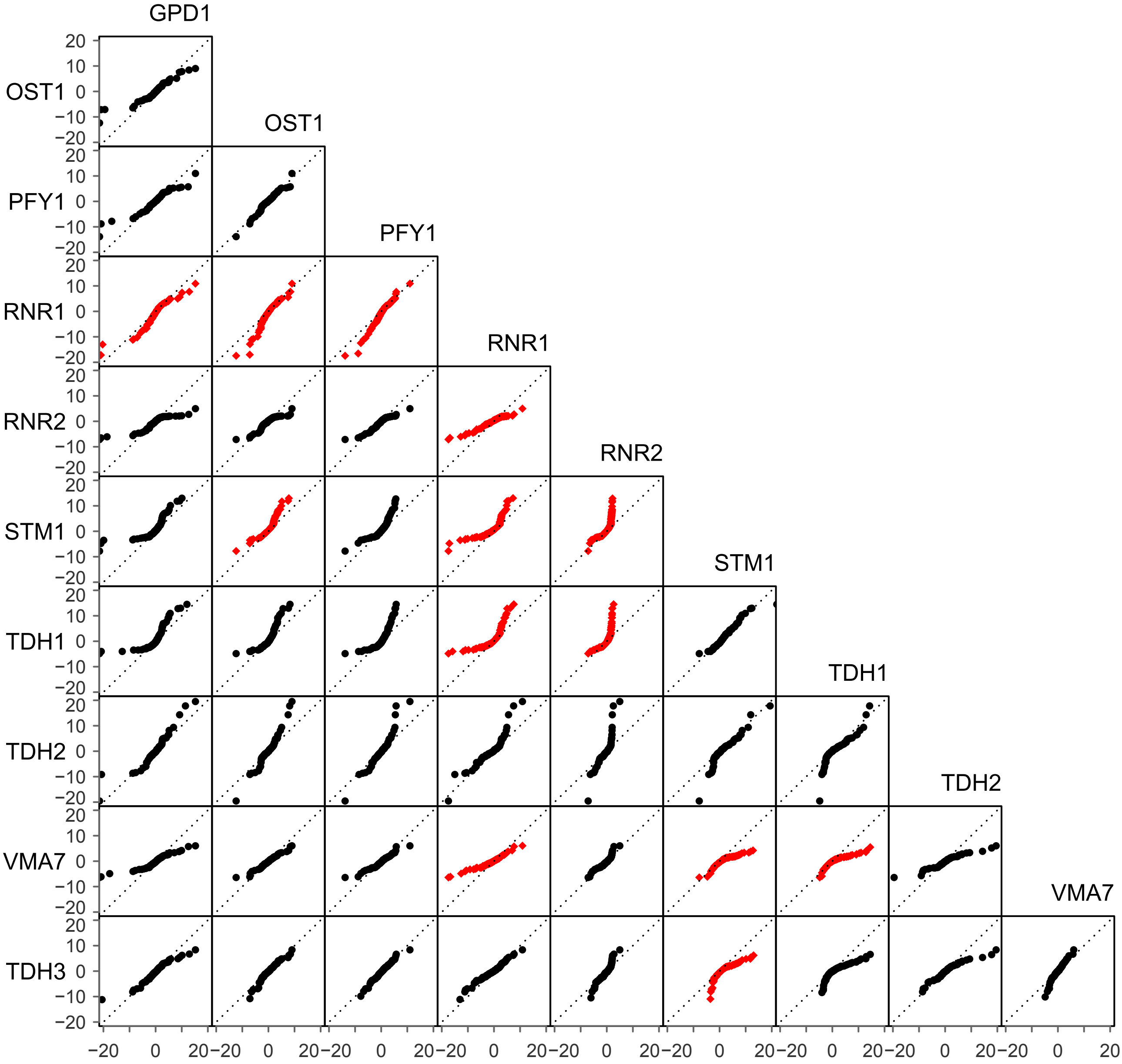

Supplementary Figure 7

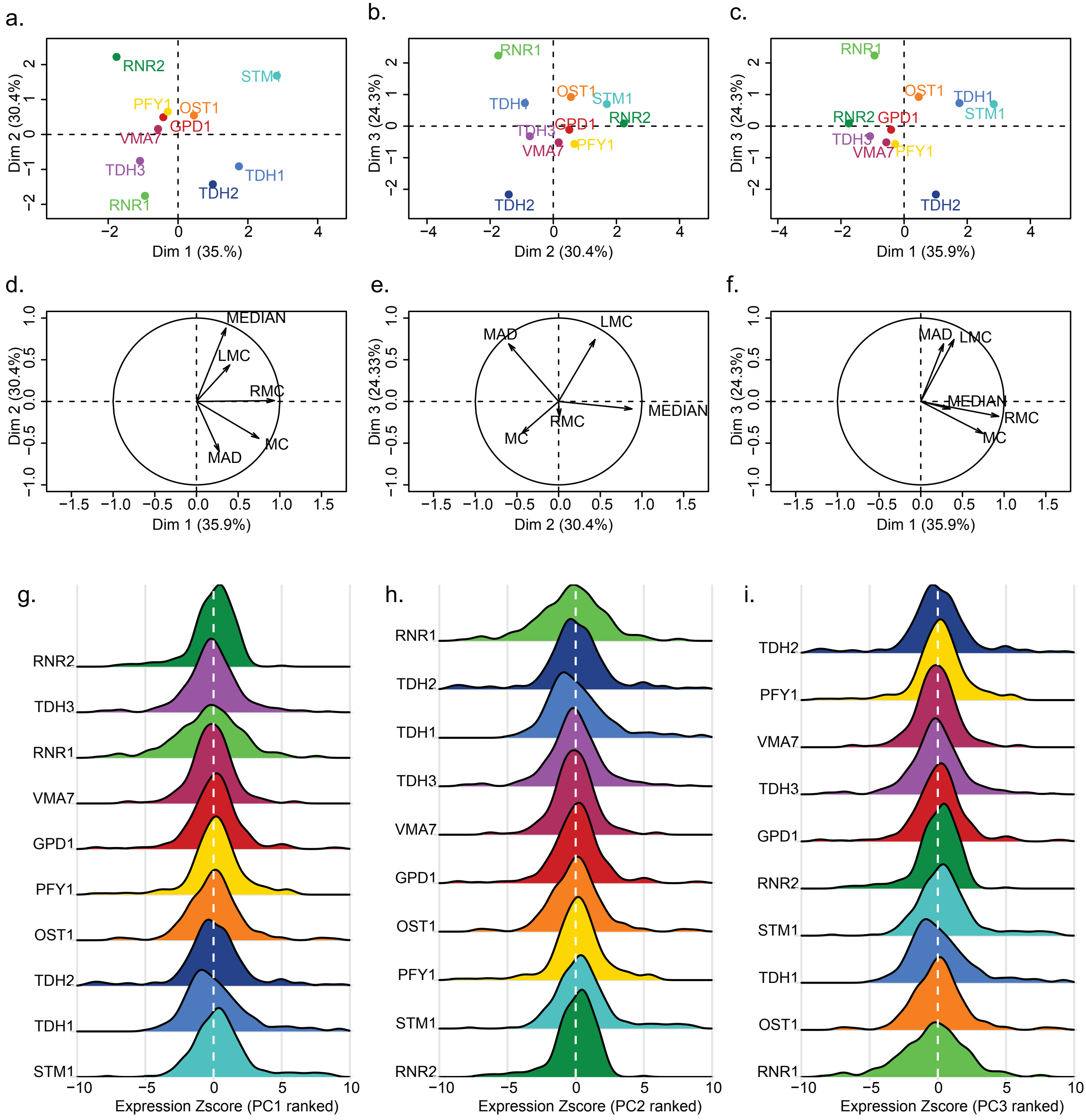

Supplementary Figure 8

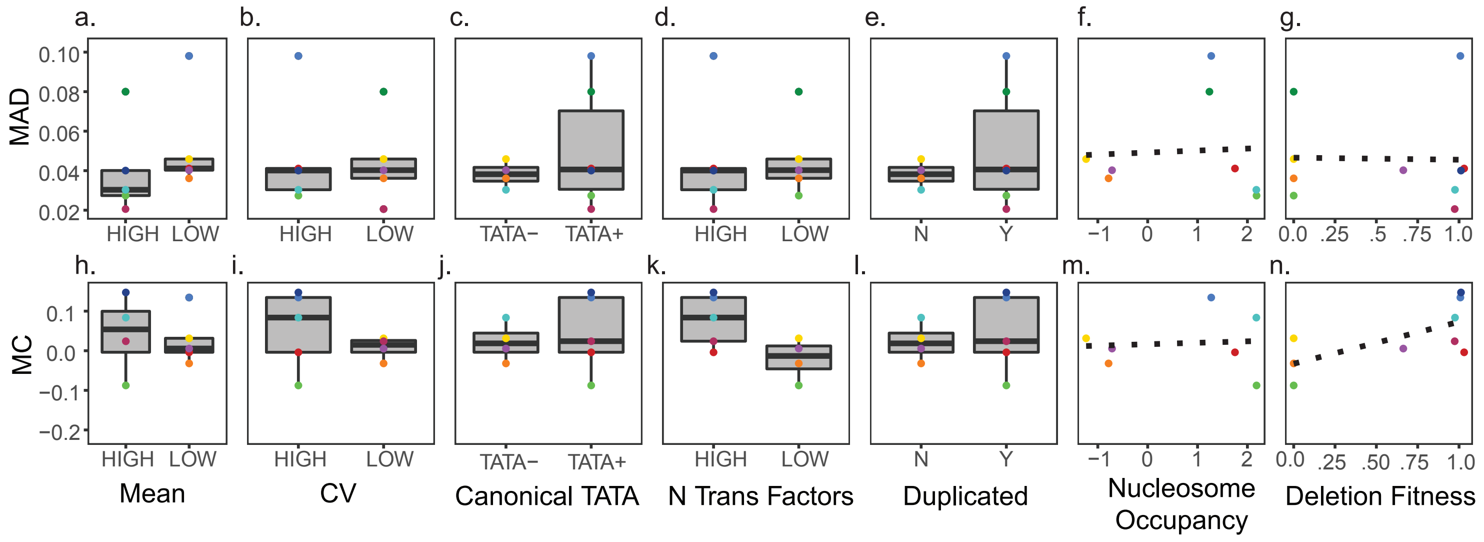

Supplementary Figure 9

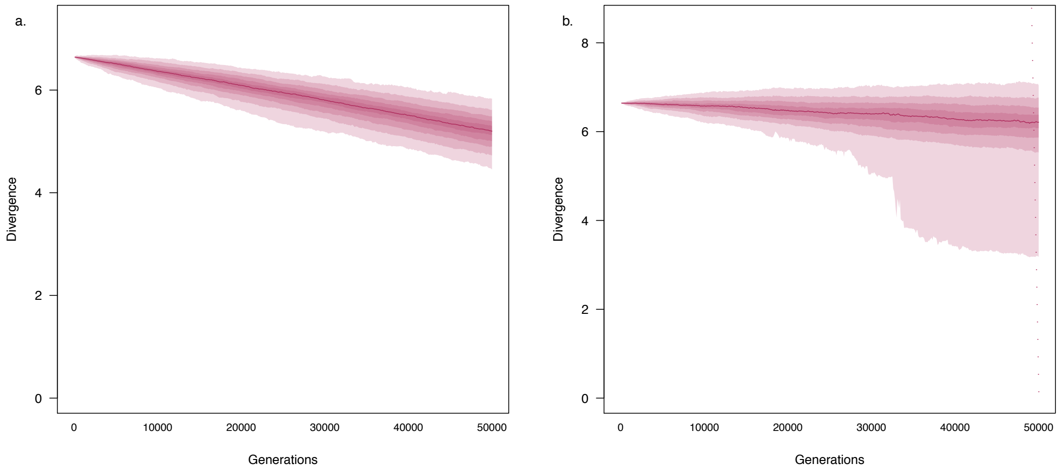

Supplementary Figure 9

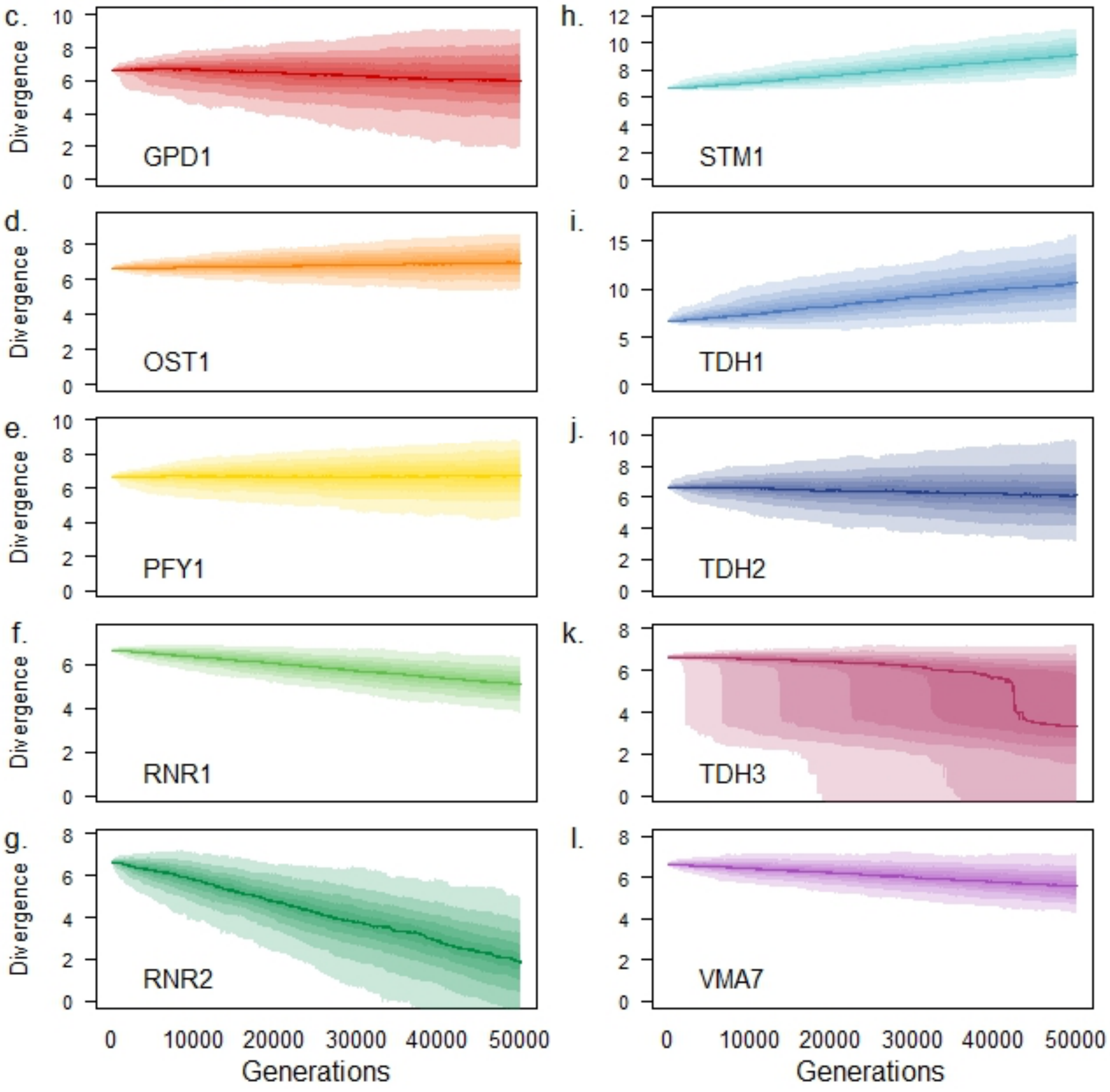

Supplementary Figure 10

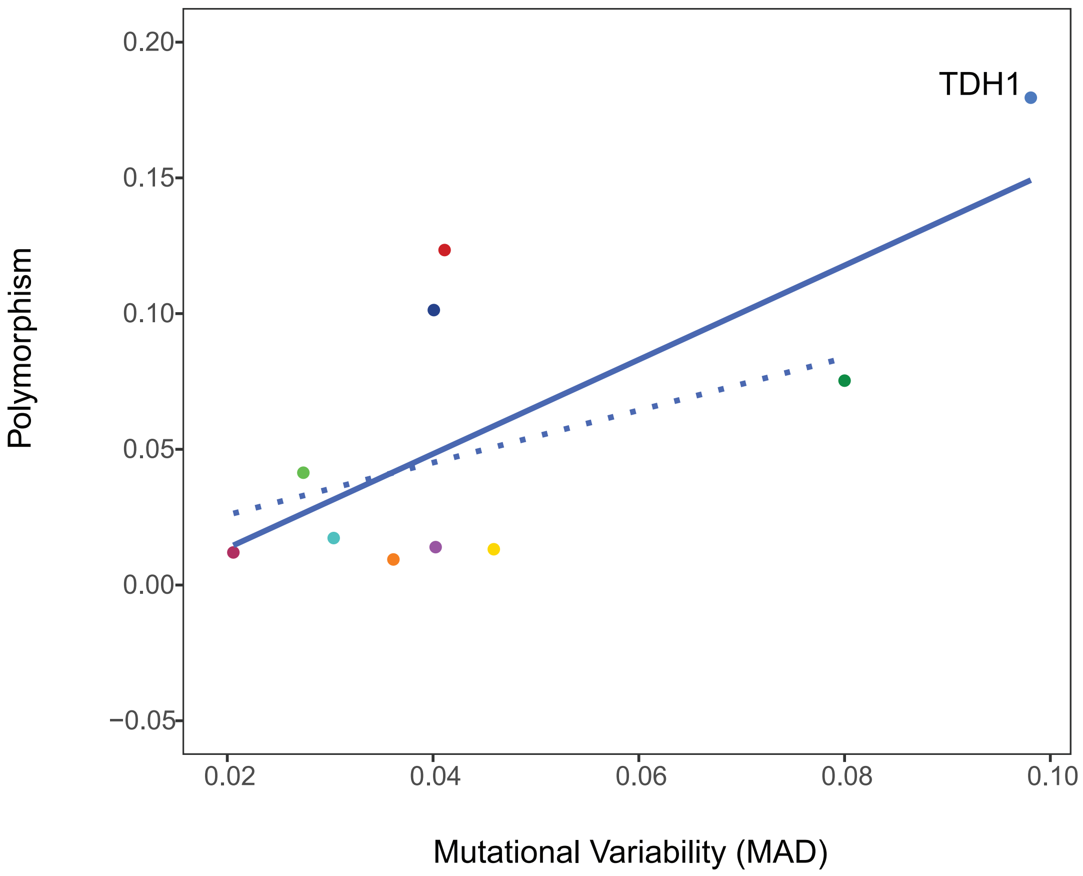
