## Supplemental tables, figures, files, and methods for "Empirical measures of mutational effects define neutral models of regulatory evolution in *Saccharomyces cerevisiae*": Supplementary File 2.R

```
#######################################################################
# Hodgins-Davis, Duveau, Walker, and Wittkopp. 2019.  
#
# Supplementary File 2
# includes R functions for processing .fcs data.
#
# To run: evaluate in the custom functions (1-5) below first, 
# then execute commands for cleaning data for each promoter 
# that begin at line 622. Requires input files with metadata 
# and columns for path to .fcs files (Supplementary File 3). 
#
# Custom functions are similar to those published in Metzger, Duveau et al 2016 MBE for:
# 1. Rotating YFP phenotypes around a centroid to eliminate effect of cell size differences
# 2. Establishing hard gates for removing cellular debris and doublets (draw gates liberally here to allow adaptive cluster finding procedures to work best)
# 3. Cleaning .fcs data
# 4. Generating quality control plots for examining contrast between original and cleaned files
# 5. Performing log transformation
#
# To run these files, you will need to replace the placeholder PATH with the appropriate directory for your filesystem.
# Similarly, you will need to ensure that the column containing the path to the .fcs files in Supplementary Files 3 is
# edited appropriately for the names and locations of the files downloaded from FlowRepository.
#
# Note for anyone interested in these analyses, I've stripped out many of the plots 
# originally created in the course of analyses as sanity checks to avoid creating dozens to hundreds of 
# automatically generated files that could clutter up somone's attempt to reproduce only what we've reported.
# I'm leaving examples of these kinds of sanity check for the first promoter (GPD1) below. If you want a
# other expampels, feel free to email andrea(dot)hodgins(dot)davis(at)gmail(dot)com asking for code for more 
# exploratory analyses.


#Clear memory
rm(list=ls())
options(warn=-1)


  ##############################
  ### 1-ROTATION OF FCS DATA ###
  ##############################
   
ROT <- function(x,Rotation){
	Result <- Rotation%*%x
	return(Result)
}

#--------------------------------------------------------------------------------------------------------------------------------#

  ###############################
  ### 2-HARD GATE CALIBRATION ###
  ###############################

GATE.CALIB <- function(x,logFSC.A.MIN,logFSC.A.MAX,logFSC.H.MIN,logFSC.H.MAX,FSC.A_FSC.H.MIN,FSC.A_FSC.H.MAX,Width.MIN,Width.MAX) {

	if (missing(logFSC.A.MIN)) {logFSC.A.MIN <- 4.7}
	if (missing(logFSC.A.MAX)) {logFSC.A.MAX <- 6.5}
	if (missing(logFSC.H.MIN)) {logFSC.H.MIN <- 4.85}
	if (missing(logFSC.H.MAX)) {logFSC.H.MAX <- 6.8}
	if (missing(FSC.A_FSC.H.MIN)) {FSC.A_FSC.H.MIN <- 0.87}
	if (missing(FSC.A_FSC.H.MAX)) {FSC.A_FSC.H.MAX <- 0.94}
	if (missing(Width.MIN)) {Width.MIN <- 40}
	if (missing(Width.MAX)) {Width.MAX <- 80}
	
	Merge.Frame <- read.FCS(x,transformation=FALSE,alter.names=TRUE)
		
	##############################	
	##Log transformation of data##
	##############################
	
	Merge.Frame <- transform(Merge.Frame,`logFSC.A`=logTrans(`FSC.A`))
	Merge.Frame <- transform(Merge.Frame,`logFSC.H`=logTrans(`FSC.H`))
	Merge.Frame <- transform(Merge.Frame,`logFL1.A`=logTrans(`FL1.A`))
	Merge.Frame <- transform(Merge.Frame,`logFL1.H`=logTrans(`FL1.H`))
	
	####################################
	##Calculate phenotypes of interest##
	####################################
	
	Data.Fluo <- as.data.frame(exprs(Merge.Frame))
	Data.Fluo[Data.Fluo == 0] <- 1
	
	Phenotype3 <- Data.Fluo[,"logFL1.A"]/Data.Fluo[,"logFSC.A"]
	Phenotype3 <- as.matrix(Phenotype3)
	colnames(Phenotype3) <- "FL1/FSC"
	Merge.Frame <- cbind2(Merge.Frame, Phenotype3)
	
	Phenotype4 <- (Data.Fluo[,"logFSC.A"])/(Data.Fluo[,"logFSC.H"])
	Phenotype4 <- as.matrix(Phenotype4)
	colnames(Phenotype4) <- "FSC.A/FSC.H"
	Merge.Frame <- cbind2(Merge.Frame, Phenotype4)		
	
	PlotAll <- as.data.frame(exprs(Merge.Frame))
	
	
	#############################
	###Quick plot to set gates###
	#############################
	
	# #Quick plot to set gates
	windows()
	par(mfrow=c(2,2))
	
	plot(PlotAll[,"Width"],PlotAll[,"logFSC.A"],pch=20,cex=0.5,col="#00000022",xlim=c(20,100),ylim=c(4,7))
	abline(v=Width.MIN)
	abline(v=Width.MAX)
	abline(h=logFSC.A.MIN)
	abline(h=logFSC.A.MAX)
	
	plot(PlotAll[,"logFSC.A"],PlotAll[,"logFL1.A"],pch=20,cex=0.5,col="#00000022",xlim=c(4,7),ylim=c(1.5,6.5))
	abline(v=logFSC.A.MIN)
	abline(v=logFSC.A.MAX)
	
	plot(PlotAll[,"logFSC.H"],PlotAll[,"logFSC.A"],pch=20,cex=0.4,col="#00000022",xlim=c(3,8),ylim=c(4,7))
	abline(v=logFSC.H.MIN)
	abline(v=logFSC.H.MAX)
	abline(h=logFSC.A.MIN)
	abline(h=logFSC.A.MAX)
	
	plot(PlotAll[,"logFSC.A"],PlotAll[,"FSC.A/FSC.H"],pch=20,cex=0.4,col="#00000022",xlim=c(3,8))
	abline(v=logFSC.A.MIN)
	abline(v=logFSC.A.MAX)
	abline(h=FSC.A_FSC.H.MIN)
	abline(h=FSC.A_FSC.H.MAX)
	
	
	
	OUTPUT <- c(logFSC.A.MIN,logFSC.A.MAX,logFSC.H.MIN,logFSC.H.MAX,FSC.A_FSC.H.MIN,FSC.A_FSC.H.MAX,Width.MIN,Width.MAX)
	names(OUTPUT) <- c("logFSC.A.MIN","logFSC.A.MAX","logFSC.H.MIN","logFSC.H.MAX","FSC.A_FSC.H.MIN","FSC.A_FSC.H.MAX","Width.MIN","Width.MAX")
	
	return(OUTPUT)
}


#--------------------------------------------------------------------------------------------------------------------------------#

#######################
### 3-CLEANING FCS DATA ###
#######################

#This function accepts: 
#   1. one row at a time of a data frame containing metadata and the path to a raw .fcs file (in column "FILENAMES") and
#   2. a set of hard gates defined as the output of the GATE.CALIB function.
#
#The script reads each .fcs file in and summarizes the collective fluorescence and cell size phenotypes 
#   for the sample, and the number of events used to estimate them before and after a number of filtering
#   steps used to separate events including cellular debris and cell doublets from single cell measurements.
#   The end of the code also performs a cetroid rotation of the cleaned data to remove effects of cell size on 
#   fluorescence estimates, and a correction for non-linearity of YFP fluorescence measured on the BD Accuri.
#
#The output of the script includes:
#   1. a smaller .txt data file that includes only the retained population of events after cleaning and 
#   phenotypes estimated for each .fcs file analyzed
#   2. a summary data frame returned in the script where the function was called with a row summarizing fluorescence 
#   and cell size phenotypes and the number of events used to estimate them for all .fcs files analyzed
#   3. currently, every 10 files analyzed, the script will also plot a series of contrasts of which subpopulations are 
#   retained and discarded in the 10th .fcs file analyzed as a .pdf to allow spot-checking as analysis proceeds

CLEANING <- function(x,GATES) {
   
  Merge.Frame <- read.FCS(x["FILENAMES"],transformation=FALSE,alter.names=TRUE)
  #for troubleshooting: Merge.Frame <- read.FCS("Path/Filename.fcs",transformation=FALSE,alter.names=TRUE)

  OUTPUT <- data.frame(matrix(nrow=1,ncol=length(c("COUNTS.INITIAL",	"COUNTS.GATES",	"COUNTS.SINGLES",	"COUNTS.FINAL", "FSC.KURTOSIS",	"WIDTH",	"FSC.MEDIAN.INITIAL",	"FSC.MAD.INITIAL",	"FL1.MEDIAN.INITIAL",	"FL1.MAD.INITIAL",	"YFP.MEDIAN.INITIAL",	"YFP.MAD.INITIAL", "YFP.SD.INITIAL",	"INTERCEPT.INITIAL",	"SLOPE.INITIAL", "THETA", "YFP.MEDIAN.ROT","YFP.MAD.ROT","YFP.SD.ROT",	"FSC.MEDIAN.FINAL",	"FSC.MAD.FINAL",	"YFP.MEDIAN.FINAL",	"YFP.MAD.FINAL", "YFP.SD.FINAL","log.YFP.MEDIAN","log.YFP.MAD","log.YFP.SD")
  )))
  colnames(OUTPUT) <- c("COUNTS.INITIAL",	"COUNTS.GATES",	"COUNTS.SINGLES",	"COUNTS.FINAL", "FSC.KURTOSIS",	"WIDTH",	"FSC.MEDIAN.INITIAL",	"FSC.MAD.INITIAL",	"FL1.MEDIAN.INITIAL",	"FL1.MAD.INITIAL",	"YFP.MEDIAN.INITIAL",	"YFP.MAD.INITIAL", "YFP.SD.INITIAL",	"INTERCEPT.INITIAL",	"SLOPE.INITIAL", "THETA", "YFP.MEDIAN.ROT","YFP.MAD.ROT","YFP.SD.ROT",	"FSC.MEDIAN.FINAL",	"FSC.MAD.FINAL",	"YFP.MEDIAN.FINAL",	"YFP.MAD.FINAL", "YFP.SD.FINAL","log.YFP.MEDIAN","log.YFP.MAD","log.YFP.SD")

  OUTPUT["COUNTS.INITIAL"] <- nrow(exprs(Merge.Frame))
  
  
  ##############################	
  ##Log transformation of data##
  ##############################
  
  Start.exp <- exprs(Merge.Frame)
  Start.exp[,"FL1.A"] <- Start.exp[,"FL1.A"] + 10
  
  Merge.Frame <- new("flowFrame",Start.exp)
  Merge.Frame <- transform(Merge.Frame,`logFSC.A`=logTrans(`FSC.A`))
  Merge.Frame <- transform(Merge.Frame,`logFSC.H`=logTrans(`FSC.H`))
  Merge.Frame <- transform(Merge.Frame,`logFL1.A`=logTrans(`FL1.A`))
  Merge.Frame <- transform(Merge.Frame,`logFL1.H`=logTrans(`FL1.H`))
  
  Data.Fluo <- as.data.frame(exprs(Merge.Frame))
  Data.Fluo[Data.Fluo == 0] <- 1	
  
  CONTAM <- sum(Data.Fluo$logFSC.A<4.5)/nrow(Data.Fluo)
  
  #if (OUTPUT["COUNTS.INITIAL"] > 1000 && x["SKIP"] == "NO" && CONTAM < 0.5)	{
  #"SKIP" VARIABLE IS A NEW COLUMN IN THE TEMPLATE FILE. All samples to be analyzed should have value = "NO" and all samples to be skipped should have value "YES"
  #It may be useful to incorporate a filter for contamination (logFSC.A<5) here.
  
  if (OUTPUT["COUNTS.INITIAL"] > 2000 && CONTAM < 0.5)	{
    ####################################
    ##Calculate phenotypes of interest##
    ####################################
    
    
    Phenotype3 <- Data.Fluo[,"logFL1.A"]/Data.Fluo[,"logFSC.A"]
    Phenotype3 <- as.matrix(Phenotype3)
    colnames(Phenotype3) <- "YFP.INITIAL"
    Merge.Frame <- cbind2(Merge.Frame, Phenotype3)
    
    Phenotype4 <- (Data.Fluo[,"logFSC.A"])/(Data.Fluo[,"logFSC.H"])
    Phenotype4 <- as.matrix(Phenotype4)
    colnames(Phenotype4) <- "FSC.A/FSC.H"
    Merge.Frame <- cbind2(Merge.Frame, Phenotype4)		
    
    PlotAll <- as.data.frame(exprs(Merge.Frame))	
    
    ############
    #Hard Gates# 
    ############
    
    rectGate <- rectangleGate(filterId="Noise Removal","logFSC.A"=c(GATES["logFSC.A.MIN"],GATES["logFSC.A.MAX"]), "logFSC.H"=c(GATES["logFSC.H.MIN"],GATES["logFSC.H.MAX"]), "FSC.A/FSC.H"=c(GATES["FSC.A_FSC.H.MIN"],GATES["FSC.A_FSC.H.MAX"]),"Width"=c(GATES["Width.MIN"],GATES["Width.MAX"]),"YFP.INITIAL"=c(min(PlotAll[,"YFP.INITIAL"]),max(PlotAll[,"YFP.INITIAL"])),"FL1.A"=c(11,max(PlotAll[,"FL1.A"])))
    
    Hard.Gates <- Subset(Merge.Frame, rectGate)
    Hard.Gates.exp <- exprs(Hard.Gates)
    
    OUTPUT["COUNTS.GATES"] <- nrow(Hard.Gates.exp)
    
    ###############################
    #Determine lower threshold for logFSC.A#
    ###############################
    
    DENSITY <- density(Hard.Gates.exp[,"logFSC.A"])
    
    TABLE <- as.data.frame(cbind(DENSITY$x,DENSITY$y))
    colnames(TABLE) <- c("logFSC.A","Density")
    
    TABLE <- subset(TABLE, Density > 0.2)
    
    for (i in 2:(nrow(TABLE)-1))
    {
      if (TABLE[i,"Density"] > TABLE[i-1,"Density"] & TABLE[i,"Density"] > TABLE[i+1,"Density"])
      {
        TABLE[i,"PEAK"] <- "YES"
      } else {
        TABLE[i,"PEAK"] <- "NO"
      }
    }
    
    PEAKS <- subset(TABLE, PEAK == "YES")
    
    FSC.LOW <- max(PEAKS[,"logFSC.A"]) - 0.6
    FSC.HIGH <- max(PEAKS[,"logFSC.A"]) + 0.4
    
    rectGate <- rectangleGate(filterId="Noise Removal","logFSC.A"=c(FSC.LOW,FSC.HIGH), "logFSC.H"=c(GATES["logFSC.H.MIN"],GATES["logFSC.H.MAX"]), "FSC.A/FSC.H"=c(GATES["FSC.A_FSC.H.MIN"],GATES["FSC.A_FSC.H.MAX"]),"Width"=c(GATES["Width.MIN"],GATES["Width.MAX"]),"YFP.INITIAL"=c(min(PlotAll[,"YFP.INITIAL"]),max(PlotAll[,"YFP.INITIAL"])),"FL1.A"=c(11,max(PlotAll[,"FL1.A"])))
    
    Hard.Gates <- Subset(Merge.Frame, rectGate)
    Hard.Gates.exp <- exprs(Hard.Gates)
    
    ################
    #Doublet Hard Gates# 
    ################
    
    Doublet.Model <- PCAgrid(cbind(Hard.Gates.exp[,"logFSC.A"],Hard.Gates.exp[,"FSC.A/FSC.H"]),k=2,method="sd",scores=TRUE,center="median")
    
    Scores <- Doublet.Model$scores
    
    Distri <- normalmixEM2comp(Scores[,2],sigsqrd=c(0.0022,0.0068)^2,mu=c(-0.0013,0.0086),lambda=c(0.56,0.44))
    
    Lambda <- Distri$lambda
    Mu <- Distri$mu
    Sigma <- Distri$sigma 
    
    Order <- c(which(Mu == min(Mu)),which(Mu == max(Mu)))
    
    Lambda <- Lambda[Order]
    Mu <- Mu[Order]
    Sigma <- Sigma[Order] 
    
    #Find good cluster
    f <- function(x) dnorm(x,m=Mu[1],sd=Sigma[1])*Lambda[1]-dnorm(x,m=Mu[2],sd=Sigma[2])*Lambda[2]
    Threshold <- try(uniroot(f,interval=c(Mu[1],Mu[2]+Sigma[2]))$root)
    
    
    #Remove big cells based on FSC.A/FSC.H
    Position <- which(Scores[,2] < Threshold)
    Doublet.Gates.exp <- Hard.Gates.exp[Position,]
    Doublet.Gates <- new("flowFrame",Doublet.Gates.exp)
    
    #Remove cells with extreme FSC.A 
    DOUBLETS <- Doublet.Gates.exp[,"logFSC.A"]
    
    MEDIAN <- median(DOUBLETS)
    MAD <- mad(DOUBLETS)
    LOW <- MEDIAN - 2*MAD
    HIGH <- MEDIAN + 2*MAD
    
    NEW.MEDIAN <- MEDIAN
    OLD.MEDIAN <- median(DOUBLETS[which(DOUBLETS > LOW & DOUBLETS < HIGH)])
    
    while (abs(NEW.MEDIAN-OLD.MEDIAN) > 0.001)
    {
      NEW.DOUBLETS <- DOUBLETS[which(DOUBLETS > LOW & DOUBLETS < HIGH)]
      OLD.MEDIAN <- NEW.MEDIAN
      NEW.MEDIAN <- median(NEW.DOUBLETS)
      NEW.MAD <- mad(NEW.DOUBLETS)
      
      LOW <- NEW.MEDIAN - 2*NEW.MAD
      HIGH <- NEW.MEDIAN + 2*NEW.MAD
    }
    
    rectGate <- rectangleGate(filterId="Outliers logFSC.A","logFSC.A"=c(LOW,HIGH))
    
    Final.Doublets <- Subset(Doublet.Gates, rectGate)
    
    OUTPUT["FSC.KURTOSIS"] <- kurtosis(DOUBLETS)
    
		#################
		#Singles Cluster#
		#################
		
		Doublet.filter <- flowClust(Final.Doublets,varNames=c("logFSC.H","logFSC.A"),K=1,B=50,min.count=1000,nu.est=2,trans=0,seed=10,z.cutoff=0,level=0.9,tol=1e-4)
		Well.pop <- split(Final.Doublets,Doublet.filter,population=list(sc1=1))
		Well.Doublet <- Well.pop$sc1
		
		Doublets.exp <- as.data.frame(exprs(Well.Doublet))
						
		OUTPUT["COUNTS.SINGLES"] <- nrow(Doublets.exp)

		####################
		#Fluo DEBRIS filter#
		####################
		
		if (OUTPUT["COUNTS.SINGLES"]>1000 & ((as.numeric(quantile(Doublets.exp[,"YFP.INITIAL"],probs=.95))-as.numeric(quantile(Doublets.exp[,"YFP.INITIAL"],probs=.05)))/as.numeric(quantile(Doublets.exp[,"YFP.INITIAL"],probs=.95)) > 0.1 )) {
  
		   Distri <- normalmixEM(Doublets.exp[,"YFP.INITIAL"],mu=c(as.numeric(quantile(Doublets.exp[,"YFP.INITIAL"],probs=.05)),median(Doublets.exp[,"YFP.INITIAL"])),lambda=c(0.1,0.9))
	
		    Lambda <- Distri$lambda
	  	  Mu <- Distri$mu
		    Sigma <- Distri$sigma 
		
		    Order <- c(which(Mu == min(Mu)),which(Mu == max(Mu)))
		
		    Lambda <- Lambda[Order]
		    Mu <- Mu[Order]
		    Sigma <- Sigma[Order] 
		
		    #Good cluster - solves for intercept
		    f <- function(x) dnorm(x,m=Mu[1],sd=Sigma[1])*Lambda[1]-dnorm(x,m=Mu[2],sd=Sigma[2])*Lambda[2]
		    Threshold <- try(uniroot(f,interval=c(Mu[1],Mu[2]+Sigma[2]))$root)
		
		 if (class(Threshold)=="try-error")
		 {
		 Threshold <- 0
		 }
		
		  #Remove cell debris based on fluorescence
		    if((Lambda[2]-Lambda[1] >.05) | Threshold == 0) {
	   	  Position <- which(Doublets.exp[,"YFP.INITIAL"] > Threshold)
		    } else {Position <- which(Doublets.exp[,"YFP.INITIAL"] < Threshold)}
		
		}else {Debris.Gates.exp<-Doublets.exp}
		
		Debris.Gates.exp <- Doublets.exp[Position,]
		Debris.Gates <- new("flowFrame",as.matrix(Debris.Gates.exp))

		   #windows()
		  #par(mfrow=c(1,2))
		  # plot.mixEM(Distri, 2, breaks=500)
		  # abline(v=Threshold,col="purple")
		  # plot(Doublets.exp[,"logFSC.A"],Doublets.exp[,"YFP.INITIAL"],pch=20,cex=0.5,col="#00000022",xlim=c(4.5,6),ylim=c(0.3,1))
		  # points(Debris.Gates.exp[,"logFSC.A"],Debris.Gates.exp[,"YFP.INITIAL"],pch=20,col="#FF000099")


		#REMOVE OUTLIERS FL1/FSC
		FLUO <- Debris.Gates.exp[,"YFP.INITIAL"]

		MEDIAN <- median(FLUO,na.rm=TRUE)
		MAD <- mad(FLUO,na.rm=TRUE)
		LOW <- MEDIAN - 4*MAD
		HIGH <- MEDIAN + 4*MAD
		NEW.MEDIAN <- MEDIAN
		OLD.MEDIAN <- median(FLUO[which(FLUO > LOW & FLUO < HIGH)])

		
		while (abs(NEW.MEDIAN-OLD.MEDIAN) > 0.001)
		{
			NEW.FLUO <- FLUO[which(FLUO > LOW & FLUO < HIGH)]
			OLD.MEDIAN <- NEW.MEDIAN
			NEW.MEDIAN <- median(NEW.FLUO)
			NEW.MAD <- mad(NEW.FLUO)

			LOW <- NEW.MEDIAN - 4*NEW.MAD
			HIGH <- NEW.MEDIAN + 4*NEW.MAD
		}

		rectGate <- rectangleGate(filterId="Outliers FL1/FSC Removal","YFP.INITIAL"=c(LOW,HIGH))

		Hard.Fluo <- Subset(Debris.Gates, rectGate)

		Gate.Fluo <- flowClust(Hard.Fluo, varNames=c("logFSC.A","logFL1.A"),K=1,B=50,min.count=1000,nu.est=1,trans=0,z.cutoff=0.5,seed=10,tol=1e-5,nu=1.5,level=0.98)
		Well.pop <- split(Hard.Fluo,Gate.Fluo,population=list(sc1=1))
		Well.Fluo <- Well.pop$sc1
		Fluo.exp <- as.data.frame(exprs(Well.Fluo))

		#SAVE DATA
		OUTPUT["CONTAM"] <- CONTAM
		OUTPUT["COUNTS.FINAL"] <- nrow(Fluo.exp)
		OUTPUT["WIDTH"] <- median(Fluo.exp[,"Width"])
		OUTPUT["FSC.MEDIAN.INITIAL"] <- median(Fluo.exp[,"logFSC.A"])
		OUTPUT["FSC.MAD.INITIAL"] <- mad(Fluo.exp[,"logFSC.A"])
		OUTPUT["FL1.MEDIAN.INITIAL"] <- median(Fluo.exp[,"logFL1.A"])
		OUTPUT["FL1.MAD.INITIAL"] <- mad(Fluo.exp[,"logFL1.A"])	
		OUTPUT["YFP.MEDIAN.INITIAL"] <- median(Fluo.exp[,"YFP.INITIAL"])
		OUTPUT["YFP.MAD.INITIAL"] <- mad(Fluo.exp[,"YFP.INITIAL"])
		OUTPUT["YFP.SD.INITIAL"] <- sd(Fluo.exp[,"YFP.INITIAL"])
		
		
		###############################################################
		#####Remove correlation between logFSC.A and FL1.A/FSC.A#######
		###############################################################
		
		#1-Define orthogonal regression
		Intercept <- c()
		Slope <- c()
		Theta <- c()
		
		Fluo.Model <- PCAgrid(cbind(Fluo.exp[,"logFSC.A"],Fluo.exp[,"YFP.INITIAL"]),k=2,method="sd",scores=FALSE,center="median")
		
		#2-Center of rotation
		x.center <- Fluo.Model$center[1]
		y.center <- Fluo.Model$center[2]
				
		#3-Initial Intercept and Slope
		Slope[1] <- Fluo.Model$loadings[2,1] / Fluo.Model$loadings[1,1]
		Intercept[1] <- Fluo.Model$center[2] - Slope[1]*Fluo.Model$center[1]
				
		#4-Calculate angle of rotation
		a <- c(x.center-0,y.center-Intercept[1]) #Vector from Intercept to Centroid
		b <- c(x.center-0,y.center-y.center) #Vector with slope 0 through Centroid
				
		Theta[1] <- acos(sum(a*b)/(sqrt(sum(a*a))*sqrt(sum(b*b)))) #Angle between 2 vectors
				
		if (Slope[1] < 0)
		{
			Theta[1] <- -Theta[1]
		}		
				
		#5-Define rotation matrix
		Rotation <- matrix(c(cos(Theta[1]),-sin(Theta[1]),sin(Theta[1]),cos(Theta[1])),ncol=2,nrow=2)
	
		#6-Transform Data
		Coord <- t(as.matrix(Fluo.exp[,c("logFSC.A","YFP.INITIAL")]))
		
		Coord[1,] <- Coord[1,] - x.center
		Coord[2,] <- Coord[2,] - y.center
		
		Result <- ROT(x=Coord,Rotation=Rotation)
		
		Result[1,] <- Result[1,] + x.center
		Result[2,] <- Result[2,] + y.center
		
		#7-Keep record of rotated values
		Fluo.exp[,"FSC.FINAL"] <- Result[1,]
		Fluo.exp[,"YFP.ROT"] <- Result[2,]
		

		
		###########################################################################
		#Apply correction for linear relation between fluorescence and mRNA levels#
		###########################################################################
		
		#Using the equation relating FL1/FSC to RNA levels from Wang and Gaigalas 2011 (J Res Natl Inst Stand Technol) to perform the translation below.  
		#Equation constants were identified by using paired pyro & Accuri data.
		#Equation features extra constant -0.05 that improves fit of equation to pyro & Accuri data
				
		REF <- 0.905811693	#scaling relationship, not essential
		NEG <- 0.519116913	
				
		for (j in 1:nrow(Fluo.exp))
		{
			#Fluo.exp[j,"YFP.FINAL"] <- (exp((Fluo.exp[j,"YFP.ROT"]-0.905274742)*log(10)/0.294448097) - 0.05)

			Fluo.exp[j,"YFP.FINAL"] <- (exp((Fluo.exp[j,"YFP.ROT"]-0.905274742)*log(10)/0.294448097) - 0.05) * (REF - NEG) + NEG
		}
	
		OUTPUT["INTERCEPT.INITIAL"] <- Intercept[1]
		OUTPUT["SLOPE.INITIAL"] <- Slope[1]
		OUTPUT["THETA"] <- Theta[1]
		OUTPUT["YFP.MEDIAN.ROT"] <- median(Fluo.exp[,"YFP.ROT"])
		OUTPUT["YFP.MAD.ROT"] <- mad(Fluo.exp[,"YFP.ROT"])
		OUTPUT["YFP.SD.ROT"] <- sd(Fluo.exp[,"YFP.ROT"])
		OUTPUT["FSC.MEDIAN.FINAL"] <- median(Fluo.exp[,"FSC.FINAL"])
		OUTPUT["FSC.MAD.FINAL"] <- mad(Fluo.exp[,"FSC.FINAL"])	
		OUTPUT["YFP.MEDIAN.FINAL"] <- median(Fluo.exp[,"YFP.FINAL"])
		OUTPUT["YFP.MAD.FINAL"] <- mad(Fluo.exp[,"YFP.FINAL"])	
		OUTPUT["YFP.SD.FINAL"] <- sd(Fluo.exp[,"YFP.FINAL"])	
		OUTPUT["YFP.CV.FINAL"] <- sd(Fluo.exp[,"YFP.FINAL"])	
		VALID <-Fluo.exp[Fluo.exp$YFP.FINAL>0,]
		VALID <-VALID[complete.cases(VALID),]
		OUTPUT["N.log.YFP.MEDIAN"] <- nrow(VALID)
		OUTPUT["log.YFP.MEDIAN"] <- median(log(VALID[,"YFP.FINAL"]))
		OUTPUT["log.YFP.MAD"] <- mad(log(VALID[,"YFP.FINAL"]))	
		OUTPUT["log.YFP.SD"] <- sd(log(VALID[,"YFP.FINAL"]))	
		
	
		
		x["COUNTER"]
		print(x["COUNTER"])
		
		#Save clean data with rotation correction
		Data <- Fluo.exp[,c("Width","Time","logFSC.A","logFSC.H","logFL1.A","logFL1.H","YFP.INITIAL","YFP.ROT","FSC.FINAL","YFP.FINAL")]
		write.table(Data,file=paste("CLEAN.DATA/",x["ASSAY"],"_",x["PLATE"],"_",x["REP"],"_",x["POSITION"],"_",x["CONDITION"],"_DAY",x["DAY"],"_DCv34.txt",sep=""),row.names=FALSE,col.names=TRUE,sep="\t",quote=FALSE)
		
		if(as.numeric(x["COUNTER"])%%10 == 0 & nrow(Fluo.exp) > 1000){
		  
		  PNAMES<-paste("CLEANING.PLOTS/PlotFilters_",x["ASSAY"],x["PLATE"],".",x["POSITION"],"_",x["CONDITION"],"_",x["REP"],"_",x["DAY"],"_DEBRISCLEAN.pdf",sep="")
		  
		  pdf(file=PNAMES)
		  
		  SAM<-sample.int(nrow(Fluo.exp),1000)
		  par(mfrow=c(2,2))
		
	  	plot(Hard.Gates.exp[SAM,"logFSC.A"],Hard.Gates.exp[SAM,"logFSC.H"],pch=20,col="#00000066")
	  	points(Doublets.exp[SAM,"logFSC.A"],Doublets.exp[SAM,"logFSC.H"],pch=20,col="#FF000099")
		
	  	plot(Hard.Gates.exp[SAM,"logFSC.A"],Hard.Gates.exp[SAM,"YFP.INITIAL"],pch=20,cex=0.5,col="#00000022",xlim=c(4.5,6),ylim=c(0.3,1))
	  	points(Debris.Gates.exp[SAM,"logFSC.A"],Debris.Gates.exp[SAM,"YFP.INITIAL"],pch=20,col="#FF000099")
		
	  	plot(Doublets.exp[SAM,"logFSC.A"],Doublets.exp[SAM,"logFL1.A"],pch=20,col="#00000066",xlim=c(5,6),ylim=c(1,6))
	  	points(Fluo.exp[SAM,"logFSC.A"],Fluo.exp[SAM,"logFL1.A"],pch=20,col="#FF000099")		
	  	
	  	hist(Fluo.exp[,"YFP.FINAL"],breaks=50,xlab="FL1/FSC")
		
	  	dev.off()
		}
		return(OUTPUT)	
		
	} else {
		OUTPUT[] <- NA
		OUTPUT["COUNTS.INITIAL"] <- nrow(exprs(Merge.Frame))
		
		Data <- as.data.frame(matrix(data=NA,ncol=10))
		colnames(Data) <- c("Width","Time","logFSC.A","logFSC.H","logFL1.A","logFL1.H","YFP.INITIAL","YFP.ROT","FSC.FINAL","YFP.FINAL")
		write.table(Data,file=paste("CLEAN.DATA/",x["ASSAY"],"_",x["PLATE"],"_",x["REP"],"_",x["POSITION"],"_",x["CONDITION"],"_DAY",x["DAY"],"_DCv34.txt",sep=""),row.names=FALSE,col.names=TRUE,sep="\t",quote=FALSE)
		
		return(OUTPUT)
	}
	
}


#--------------------------------------------------------------------------------------------------------------------------------#

  ###########################
### 4-QUALITY CONTROL PLOTS ###
  ###########################


FCS.PLOT <- function(x) {
	
	INITIAL.DATA <- read.FCS(x["INITIAL"],transformation=FALSE,alter.names=TRUE)
	CLEAN.DATA <-  read.table(x["CLEAN"],header=TRUE,as.is=TRUE)

	if ( nrow(CLEAN.DATA) > 750)	{

	##############################	
	##Log transformation of data##
	##############################
		
		Merge.Frame <- transform(INITIAL.DATA,`logFSC.A`=logTrans(`FSC.A`))
		Merge.Frame <- transform(Merge.Frame,`logFSC.H`=logTrans(`FSC.H`))
		Merge.Frame <- transform(Merge.Frame,`logFL1.A`=logTrans(`FL1.A`))
		Merge.Frame <- transform(Merge.Frame,`logFL1.H`=logTrans(`FL1.H`))
		
	####################################
	##Calculate phenotypes of interest##
	####################################
		
		Data.Fluo <- as.data.frame(exprs(Merge.Frame))
		Data.Fluo[Data.Fluo == 0] <- 1
		
		Phenotype3 <- Data.Fluo[,"logFL1.A"]/Data.Fluo[,"logFSC.A"]
		Phenotype3 <- as.matrix(Phenotype3)
		colnames(Phenotype3) <- "YFP.INITIAL"
		Merge.Frame <- cbind2(Merge.Frame, Phenotype3)
		
		Phenotype4 <- (Data.Fluo[,"logFSC.A"])/(Data.Fluo[,"logFSC.H"])
		Phenotype4 <- as.matrix(Phenotype4)
		colnames(Phenotype4) <- "FSC.A/FSC.H"
		Merge.Frame <- cbind2(Merge.Frame, Phenotype4)		
		
		PLOT.ALL <- as.data.frame(exprs(Merge.Frame))

		NAME <- paste(x["ASSAY"],".",x["PLATE"],".",x["POSITION"],"_",x["CONDITION"],"_",x["REP"],"_DAY",x["DAY"],sep="")
	

		pdf(paste("CLEANING.PLOTS/",NAME,"_summary.pdf",sep=""))

	par(mfrow=c(2,2))

	plot(PLOT.ALL$logFSC.A,PLOT.ALL$logFSC.H,pch=20,cex=0.5,col="#00000033",xlim=c(4.3,6.5),ylim=c(5,7),xlab="logFSC.A",ylab="logFSC.H",main=NAME)
	points(CLEAN.DATA$logFSC.A,CLEAN.DATA$logFSC.H,pch=20,cex=0.5,col="#FF000066",main=NAME)

	plot(PLOT.ALL$logFSC.A,PLOT.ALL$logFL1.A,pch=20,cex=0.5,col="#00000033",xlim=c(4.3,6.5),ylim=c(1.5,6.5),xlab="logFSC.A",ylab="logFL1.A",main=NAME)
	points(CLEAN.DATA$logFSC.A,CLEAN.DATA$logFL1.A,pch=20,cex=0.5,col="#FF000066",main=NAME)

	plot(CLEAN.DATA$logFSC.A,CLEAN.DATA$YFP.INITIAL,pch=20,cex=0.5,col="#FF0000AA",xlab="logFSC.A",ylab="FL1/FSC",main=NAME)
	points(CLEAN.DATA$FSC.FINAL,CLEAN.DATA$YFP.ROT,pch=20,cex=0.5,col="#00FF00AA",main=NAME)
	abline(lm(CLEAN.DATA$YFP.INITIAL~CLEAN.DATA$logFSC.A),lty=2,col="darkred",lwd=2.5)
	abline(lm(CLEAN.DATA$YFP.ROT~CLEAN.DATA$FSC.FINAL),lty=2,col="darkgreen",lwd=2.5)
	
	hist(CLEAN.DATA$YFP.FINAL,breaks=50,main=NAME,xlab="FL1/FSC")

	dev.off()	

	}
	
}


###########################
### 5- LOG TRANSFORMATION ###
###########################


logTrans <- logTransform(transformationId="log10-transformation",logbase=10,r=1,d=1)

#--------------------------------------------------------------------------------------------------------------------------------#

################################################
# Analysis of each promoter attached below.
#
#


##########
#  GPD1  #
##########


#Clear memory
rm(list=ls())
options(warn=-1)


#####################
#LOADING LIBRARIES#
#####################

library(flowCore)
library(flowClust)
library(flowViz)
library(plotrix)
library(nlme)
library(MethComp)
library(outliers)
library(pcaPP)
library(reshape2)
library(MASS)
library(ggplot2)
library(Hmisc)
library(fBasics)
library(lawstat)
library(fitdistrplus)
library(mixtools)
library(vioplot)
library(gplots)
library(RColorBrewer)
library(calibrate)
library(e1071)

box <- graphics::box


###############
#  CLEAN DATA #
###############

#Set working directory
parent.dir <- PATH
setwd(parent.dir)

#Load experiment setup
SETUP <- read.csv("Template_GPD1_V2_10102016.csv",header=TRUE,as.is=TRUE)
TESTNAMES <- list.files("PATH/GPD1/",pattern=".fcs",full.name=TRUE,recursive=TRUE,include.dirs=TRUE) #This should direct to folder containing .fcs files to be analyzed
SETUP<-subset(SETUP, SETUP$FILENAMES %in% TESTNAMES, drop = TRUE)
SETUP[,"COUNTER"] <- c(1:nrow(SETUP))

#Determine Hard Gates
GATES <- GATE.CALIB(SETUP$FILENAMES[1])

#Clean Data
Output <- apply(SETUP,1,CLEANING,GATES=GATES)
OUTPUT <- as.data.frame(Output[[1]])
OUTPUT[2:nrow(SETUP),1:ncol(OUTPUT)] <- NA
for (i in 2:nrow(SETUP))
{
  if (is.null(Output[[i]]))
  {} 
  else {
    OUTPUT[i,] <- Output[[i]]
  }
}

write.table(OUTPUT,"GPD1_Experiment.Output_V2_DCv34.txt",row.names=FALSE,sep="\t")

CLEAN <- cbind.data.frame(SETUP[,1:(ncol(SETUP)-2)],OUTPUT)

write.table(CLEAN,"GPD1_Clean.Data_V2_10112016_DCv34.txt",row.names=FALSE,sep="\t")

################################################################
#     PLATE CORRECTION & OUTLIER REMOVAL
################################################################

#Clear memory
rm(list=ls())
options(warn=-1)

#####################
#LOADING LIBRARIES#
#####################
library(flowCore)
library(flowClust)
library(flowViz)
library(pcaPP)
library(mixtools)
library(plyr)
library(MASS)
box <- graphics::box
options(warn=-1)

####################################
#Quality Control and Corrections#
####################################

parent.dir <- PATH
setwd(parent.dir)

DATA.TYPE <- c(
  "factor","factor","factor","factor","factor","factor","factor","factor","factor","factor","factor","factor","factor","factor","factor",
  "integer","integer","integer","integer",
  "numeric","numeric","numeric","numeric","numeric","numeric","numeric","numeric","numeric","numeric","numeric","numeric","numeric","numeric","numeric","numeric","numeric","numeric","numeric","numeric","numeric","numeric","numeric","numeric","numeric",
  "integer")

###Read in data sets
CLEAN.DATA <- read.table("GPD1_Clean.Data_V2_10112016_DCv34.txt",colClasses=DATA.TYPE,header=TRUE, sep="\t")

###Remove samples with NA
CLEAN.DATA <- CLEAN.DATA[complete.cases(CLEAN.DATA),]

##Translate into log scale for appropriate correction
CLEAN.DATA[,"log.RNA.MEDIAN"] <- log((CLEAN.DATA[,"YFP.MEDIAN.FINAL"]) + 0.05)
CLEAN.DATA[,"log.RNA.MAD"] <- CLEAN.DATA[,"YFP.MAD.FINAL"] / (CLEAN.DATA[,"YFP.MEDIAN.FINAL"] + 0.05)
CLEAN.DATA[,"log.RNA.SD"] <- CLEAN.DATA[,"YFP.SD.FINAL"] / (CLEAN.DATA[,"YFP.MEDIAN.FINAL"] + 0.05)

###Remove samples with less than 1000 events
CLEAN.DATA <- subset(CLEAN.DATA,CLEAN.DATA$COUNTS.FINAL > 1000)

####Add variable to ID whether control is own promoter or TDH3
CLEAN.DATA[,"CTRL.GENO"]<-as.factor("GPD1")

###Select desired data / Drop known bad samples
pdf(file="Plate Correction control plots GPD1 10112016 DCv34.pdf")

#windows()
plot(CLEAN.DATA$YFP.MEDIAN.FINAL[CLEAN.DATA$CONDITION=="TDH3.SHAM"]~CLEAN.DATA$PLATE[CLEAN.DATA$CONDITION=="TDH3.SHAM"],col="red", main="Control genotypes visualized",xlab="Plate",ylab="YFP Median Final")
boxplot(CLEAN.DATA$YFP.MEDIAN.FINAL[CLEAN.DATA$CONDITION=="CTRL"]~CLEAN.DATA$PLATE[CLEAN.DATA$CONDITION=="CTRL"],col="black",fill="lightgray",add=TRUE)
points(CLEAN.DATA$YFP.MEDIAN.FINAL[CLEAN.DATA$CONDITION=="SHAM"]~CLEAN.DATA$PLATE[CLEAN.DATA$CONDITION=="SHAM"],col="blue",add=TRUE)
legend("left",c("Plate controls","TDH3 Sham","Sham"),text.col=c("black","red","blue"),bty="n")

par(mfrow=c(1,2))
plot(CLEAN.DATA$FSC.MEDIAN.FINAL[CLEAN.DATA$CONDITION=="CTRL"],CLEAN.DATA$YFP.MEDIAN.FINAL[CLEAN.DATA$CONDITION=="CTRL"],col="black",ylim=c(0.5,1.1),xlim=c(5,6),pch=20,main="Plate Controls",xlab="Cell Size (FSC Median)",ylab="YFP")
abline(v=5.35,lty=3)
abline(h=.53,lty=3)

plot(CLEAN.DATA$FSC.MEDIAN.FINAL[CLEAN.DATA$CONDITION=="SHAM"],CLEAN.DATA$YFP.MEDIAN.FINAL[CLEAN.DATA$CONDITION=="SHAM"],col="blue",ylim=c(0.5,1.1),xlim=c(5,6),pch=20,main="Data",xlab="Cell Size (FSC Median)",ylab="YFP")
points(CLEAN.DATA$FSC.MEDIAN.FINAL[CLEAN.DATA$CONDITION=="TDH3.SHAM"],CLEAN.DATA$YFP.MEDIAN.FINAL[CLEAN.DATA$CONDITION=="TDH3.SHAM"],col="black",pch=20)
points(CLEAN.DATA$FSC.MEDIAN.FINAL[CLEAN.DATA$CONDITION=="EMS"],CLEAN.DATA$YFP.MEDIAN.FINAL[CLEAN.DATA$CONDITION=="EMS"],col="red",pch=20)
points(CLEAN.DATA$FSC.MEDIAN.FINAL[CLEAN.DATA$CONDITION=="EMS.LOW"],CLEAN.DATA$YFP.MEDIAN.FINAL[CLEAN.DATA$CONDITION=="EMS.LOW"],col="orange",pch=20)
points(CLEAN.DATA$FSC.MEDIAN.FINAL[CLEAN.DATA$CONDITION=="EMS.HIGH"],CLEAN.DATA$YFP.MEDIAN.FINAL[CLEAN.DATA$CONDITION=="EMS.HIGH"],col="orange",pch=20)
points(CLEAN.DATA$FSC.MEDIAN.FINAL[CLEAN.DATA$CONDITION=="NULL"],CLEAN.DATA$YFP.MEDIAN.FINAL[CLEAN.DATA$CONDITION=="NULL"],col="green",pch=20)
points(CLEAN.DATA$FSC.MEDIAN.FINAL[CLEAN.DATA$CONDITION=="SHAM"],CLEAN.DATA$YFP.MEDIAN.FINAL[CLEAN.DATA$CONDITION=="SHAM"],col="blue",pch=20)

abline(v=5.4,lty=3)
abline(h=.53,lty=3)
abline(h=0.7,lty=3)
legend("topleft",c("Sham","Random EMS","EMS Tails","Null","TDH3 Sham"),text.col=c("blue","red","orange","green","black"),bty="n")

CLEAN.DATA$CONTAM.CHECK<-"NONE"
CLEAN.DATA$CONTAM.CHECK[(CLEAN.DATA$CONDITION=="EMS"|CLEAN.DATA$CONDITION=="SHAM"|CLEAN.DATA$CONDITION=="EMS.LOW"|CLEAN.DATA$CONDITION=="EMS.HIGH") & (CLEAN.DATA$YFP.MEDIAN.FINAL<.53)]<-"BACT.CONTAM.CHECK"
CLEAN.DATA$CONTAM.CHECK[(CLEAN.DATA$CONDITION=="EMS"|CLEAN.DATA$CONDITION=="SHAM"|CLEAN.DATA$CONDITION=="EMS.LOW"|CLEAN.DATA$CONDITION=="EMS.HIGH") & (CLEAN.DATA$YFP.MEDIAN.FINAL>0.8)]<-"TDH3.CONTAM.CHECK"
CLEAN.DATA$CONTAM.CHECK[(CLEAN.DATA$CONDITION=="TDH3.SHAM" & CLEAN.DATA$YFP.MEDIAN.FINAL<0.8)]<-"EXP.SHAM.CONTAM.CHECK"
CLEAN.DATA$CONTAM.CHECK[(CLEAN.DATA$CONDITION=="NULL" & CLEAN.DATA$YFP.MEDIAN.FINAL>0.53)]<-"EXP.NULL.CONTAM.CHECK"


###Seperate controls and remove FSC outliers
CONTROL <- subset(CLEAN.DATA,CLEAN.DATA$CONDITION == "CTRL")
FSC.CORRECT <- lm(CONTROL$FSC.MEDIAN.FINAL ~ 0 + CONTROL$FLOW.RUN)
RESID <- FSC.CORRECT$resid
CLEAN.DATA <- droplevels(CLEAN.DATA[-which(CLEAN.DATA$CONDITION == "CTRL")[abs(RESID) > 3*mad(RESID)],])
CONTROL <- droplevels(subset(CLEAN.DATA,CLEAN.DATA$CONDITION == "CTRL"))

INITIAL.CORRECT <- lm(CONTROL$log.RNA.MEDIAN ~ 0 + CONTROL$FLOW.RUN)
RESID <- INITIAL.CORRECT$resid
CLEAN.DATA<- droplevels(CLEAN.DATA[-which(CLEAN.DATA$CONDITION == "CTRL")[abs(RESID) > 3*mad(RESID)],])
CONTROL <- droplevels(subset(CLEAN.DATA,CLEAN.DATA$CONDITION == "CTRL"))


#Remove any flow runs with insufficient # of control positions to perform full plate correction
`%notin%` <- function(x,y) !(x %in% y)
RESULT<-count(CONTROL$FLOW.RUN)
RESULT.MUT<-count(CLEAN.DATA$FLOW.RUN)
EXCLUDE1<-(RESULT.MUT[which(RESULT.MUT[,1] %notin% RESULT[,1]),1])
EXCLUDE2<-(RESULT[which(RESULT[,2] <5),1])
CLEAN.DATA<-droplevels(CLEAN.DATA[CLEAN.DATA$FLOW.RUN %notin% EXCLUDE1,])
CLEAN.DATA<-droplevels(CLEAN.DATA[CLEAN.DATA$FLOW.RUN %notin% EXCLUDE2,])
CONTROL <- subset(CLEAN.DATA,CLEAN.DATA$CONDITION == "CTRL")


###Corrections for YFP median and MAD
#Correct Median
#fit<- lm(log.RNA.MEDIAN~FLOW.RUN+REP+STACK.ORDER+STACK.ORDER:FLOW.RUN+REP+ROW+COLUMN+BLOCK+BLOCK:ROW+ROW:COLUMN+STACK.ORDER:BLOCK,data=CONTROL)
#step <- stepAIC(fit, direction="both")
log.RNA.MEDIAN.CORRECT <- lm(log.RNA.MEDIAN ~ FLOW.RUN + COLUMN + ROW + ROW:COLUMN, data=CONTROL)
CLEAN.DATA[,"log.RNA.MEDIAN.CORRECT"] <- CLEAN.DATA$log.RNA.MEDIAN - predict(log.RNA.MEDIAN.CORRECT, CLEAN.DATA) + mean(CONTROL$log.RNA.MEDIAN)
CONTROL <- subset(CLEAN.DATA,CLEAN.DATA$CONDITION == "CTRL")

#Correct sd
#fit<- lm(log.RNA.SD~FLOW.RUN+REP+STACK.ORDER+STACK.ORDER:FLOW.RUN+REP+ROW+COLUMN+BLOCK+BLOCK:ROW+ROW:COLUMN+STACK.ORDER:BLOCK,data=CONTROL)
#step <- step(fit, direction="both")
log.RNA.SD.CORRECT <- lm(log.RNA.SD ~  FLOW.RUN + ROW + COLUMN+ ROW:COLUMN,data=CONTROL)
CLEAN.DATA[,"log.RNA.SD.CORRECT"] <- CLEAN.DATA$log.RNA.SD - predict(log.RNA.SD.CORRECT , CLEAN.DATA) + mean(CONTROL$log.RNA.SD)
CONTROL <- subset(CLEAN.DATA,CLEAN.DATA$CONDITION == "CTRL")

#Correct mad
#fit<- lm(log.RNA.MAD~FLOW.RUN+REP+STACK.ORDER+STACK.ORDER:FLOW.RUN+REP+ROW+COLUMN+BLOCK+BLOCK:ROW+ROW:COLUMN+STACK.ORDER:BLOCK,data=CONTROL)
#step <- step(fit, direction="both")
log.RNA.MAD.CORRECT <- lm(log.RNA.MAD ~  FLOW.RUN + ROW + COLUMN+ ROW:COLUMN,data=CONTROL)
CLEAN.DATA[,"log.RNA.MAD.CORRECT"] <- CLEAN.DATA$log.RNA.MAD - predict(log.RNA.MAD.CORRECT , CLEAN.DATA) + mean(CONTROL$log.RNA.MAD)
CONTROL <- subset(CLEAN.DATA,CLEAN.DATA$CONDITION == "CTRL")

#Correct FSC
#fit<- lm(FSC.MEDIAN.FINAL~FLOW.RUN+REP+STACK.ORDER+STACK.ORDER:FLOW.RUN+REP+ROW+COLUMN+BLOCK+BLOCK:ROW+ROW:COLUMN+STACK.ORDER:BLOCK,data=CONTROL)
#step <- stepAIC(fit, direction="both")
FSC.CORRECT <- lm(FSC.MEDIAN.FINAL ~  FLOW.RUN + ROW + COLUMN,data=CONTROL)
CLEAN.DATA[,"FSC.CORRECT"] <- CLEAN.DATA$FSC.MEDIAN.FINAL - predict(FSC.CORRECT, CLEAN.DATA) + mean(CONTROL$FSC.MEDIAN.FINAL)
CONTROL <- subset(CLEAN.DATA,CLEAN.DATA$CONDITION == "CTRL")

par(mfrow=c(2,3))
plot(CLEAN.DATA$FSC.MEDIAN.FINAL[CLEAN.DATA$CONDITION=="CTRL"],CLEAN.DATA$log.RNA.MEDIAN[CLEAN.DATA$CONDITION=="CTRL"],col="black",main="Before Correction",ylim=c(-0.6,0.3),xlim=c(5,6),pch=20,xlab="Plate Controls: FSC Median Final",ylab="Plate Controls: YFP (log RNA Median)")
points(CLEAN.DATA$FSC.MEDIAN.FINAL[CLEAN.DATA$CONDITION=="NULL"],CLEAN.DATA$log.RNA.MEDIAN[CLEAN.DATA$CONDITION=="NULL"],col="green",pch=20)
legend("topleft",c(paste("Plate Controls=",as.character(CLEAN.DATA$CTRL.GENO[1])),"Null"),text.col=c("black","green"),bty="n")

plot(CLEAN.DATA$FSC.MEDIAN.FINAL[CLEAN.DATA$CONDITION=="CTRL"],CLEAN.DATA$log.RNA.MEDIAN.CORRECT[CLEAN.DATA$CONDITION=="CTRL"],col="black",main="YFP Corrected",ylim=c(-0.6,0.3),xlim=c(5,6),pch=20,xlab="Plate Controls: FSC Median Final",ylab="Plate Controls: YFP Corrected (log RNA Median)")
points(CLEAN.DATA$FSC.MEDIAN.FINAL[CLEAN.DATA$CONDITION=="NULL"],CLEAN.DATA$log.RNA.MEDIAN.CORRECT[CLEAN.DATA$CONDITION=="NULL"],col="green",pch=20)

plot(CLEAN.DATA$FSC.CORRECT[CLEAN.DATA$CONDITION=="CTRL"],CLEAN.DATA$log.RNA.MEDIAN.CORRECT[CLEAN.DATA$CONDITION=="CTRL"],col="black",main="FSC & YFP Corrected",ylim=c(-0.6,0.3),xlim=c(5,6),pch=20,xlab="Plate Controls: FSC Median Corrected",ylab="Plate Controls: YFP Corrected (log RNA Median)")
points(CLEAN.DATA$FSC.CORRECT[CLEAN.DATA$CONDITION=="NULL"],CLEAN.DATA$log.RNA.MEDIAN.CORRECT[CLEAN.DATA$CONDITION=="NULL"],col="green",pch=20)

plot(CLEAN.DATA$FSC.MEDIAN.FINAL[CLEAN.DATA$CONDITION=="SHAM"],CLEAN.DATA$log.RNA.MEDIAN[CLEAN.DATA$CONDITION=="SHAM"],col="blue",ylim=c(-0.6,0.3),xlim=c(5,6),pch=20,xlab="FSC Median Final",ylab="YFP (log RNA Median)")
points(CLEAN.DATA$FSC.MEDIAN.FINAL[CLEAN.DATA$CONDITION=="EMS"],CLEAN.DATA$log.RNA.MEDIAN[CLEAN.DATA$CONDITION=="EMS"],col="red",pch=20)
points(CLEAN.DATA$FSC.MEDIAN.FINAL[CLEAN.DATA$CONDITION=="EMS.LOW"],CLEAN.DATA$log.RNA.MEDIAN[CLEAN.DATA$CONDITION=="EMS.LOW"],col="orange",pch=20)
points(CLEAN.DATA$FSC.MEDIAN.FINAL[CLEAN.DATA$CONDITION=="EMS.HIGH"],CLEAN.DATA$log.RNA.MEDIAN[CLEAN.DATA$CONDITION=="EMS.HIGH"],col="orange",pch=20)
points(CLEAN.DATA$FSC.MEDIAN.FINAL[CLEAN.DATA$CONDITION=="NULL"],CLEAN.DATA$log.RNA.MEDIAN[CLEAN.DATA$CONDITION=="NULL"],col="green",pch=20)
points(CLEAN.DATA$FSC.MEDIAN.FINAL[CLEAN.DATA$CONDITION=="SHAM"],CLEAN.DATA$log.RNA.MEDIAN[CLEAN.DATA$CONDITION=="SHAM"],col="blue",pch=20)
legend("topleft",c("Sham","Random EMS","EMS Tails","Null"),text.col=c("blue","red","orange","green"),bty="n")

plot(CLEAN.DATA$FSC.MEDIAN.FINAL[CLEAN.DATA$CONDITION=="SHAM"],CLEAN.DATA$log.RNA.MEDIAN.CORRECT[CLEAN.DATA$CONDITION=="SHAM"],col="blue",ylim=c(-0.6,0.3),xlim=c(5,6),pch=20,xlab="FSC Median Final",ylab="YFP Corrected (log RNA Median)")
points(CLEAN.DATA$FSC.MEDIAN.FINAL[CLEAN.DATA$CONDITION=="EMS"],CLEAN.DATA$log.RNA.MEDIAN.CORRECT[CLEAN.DATA$CONDITION=="EMS"],col="red",pch=20)
points(CLEAN.DATA$FSC.MEDIAN.FINAL[CLEAN.DATA$CONDITION=="EMS.LOW"],CLEAN.DATA$log.RNA.MEDIAN.CORRECT[CLEAN.DATA$CONDITION=="EMS.LOW"],col="orange",pch=20)
points(CLEAN.DATA$FSC.MEDIAN.FINAL[CLEAN.DATA$CONDITION=="EMS.HIGH"],CLEAN.DATA$log.RNA.MEDIAN.CORRECT[CLEAN.DATA$CONDITION=="EMS.HIGH"],col="orange",pch=20)
points(CLEAN.DATA$FSC.MEDIAN.FINAL[CLEAN.DATA$CONDITION=="NULL"],CLEAN.DATA$log.RNA.MEDIAN.CORRECT[CLEAN.DATA$CONDITION=="NULL"],col="green",pch=20)
points(CLEAN.DATA$FSC.MEDIAN.FINAL[CLEAN.DATA$CONDITION=="SHAM"],CLEAN.DATA$log.RNA.MEDIAN.CORRECT[CLEAN.DATA$CONDITION=="SHAM"],col="blue",pch=20)

plot(CLEAN.DATA$FSC.CORRECT[CLEAN.DATA$CONDITION=="SHAM"],CLEAN.DATA$log.RNA.MEDIAN.CORRECT[CLEAN.DATA$CONDITION=="SHAM"],col="blue",ylim=c(-0.6,0.3),xlim=c(5,6),pch=20,xlab="FSC Median Corrected",ylab="YFP Corrected(log RNA Median)")
points(CLEAN.DATA$FSC.CORRECT[CLEAN.DATA$CONDITION=="EMS"],CLEAN.DATA$log.RNA.MEDIAN.CORRECT[CLEAN.DATA$CONDITION=="EMS"],col="red",pch=20)
points(CLEAN.DATA$FSC.CORRECT[CLEAN.DATA$CONDITION=="EMS.LOW"],CLEAN.DATA$log.RNA.MEDIAN.CORRECT[CLEAN.DATA$CONDITION=="EMS.LOW"],col="orange",pch=20)
points(CLEAN.DATA$FSC.CORRECT[CLEAN.DATA$CONDITION=="EMS.HIGH"],CLEAN.DATA$log.RNA.MEDIAN.CORRECT[CLEAN.DATA$CONDITION=="EMS.HIGH"],col="orange",pch=20)
points(CLEAN.DATA$FSC.CORRECT[CLEAN.DATA$CONDITION=="NULL"],CLEAN.DATA$log.RNA.MEDIAN.CORRECT[CLEAN.DATA$CONDITION=="NULL"],col="green",pch=20)
points(CLEAN.DATA$FSC.CORRECT[CLEAN.DATA$CONDITION=="SHAM"],CLEAN.DATA$log.RNA.MEDIAN.CORRECT[CLEAN.DATA$CONDITION=="SHAM"],col="blue",pch=20)

#Back translate to linear scale
CLEAN.DATA[,"YFP.MEDIAN.CORRECT"] <- (exp(CLEAN.DATA[,"log.RNA.MEDIAN.CORRECT"]) - 0.05)
CLEAN.DATA[,"YFP.SD.CORRECT"] <- CLEAN.DATA[,"log.RNA.SD.CORRECT"] * CLEAN.DATA[,"YFP.MEDIAN.CORRECT"]
CLEAN.DATA[,"YFP.MAD.CORRECT"] <- CLEAN.DATA[,"log.RNA.MAD.CORRECT"] * CLEAN.DATA[,"YFP.MEDIAN.CORRECT"]


#PERFORM FABIEN'S OUTLIER CORRECTION FILTERING OUT ANY REPLICATE OUTSIDE OF 4*MAD

#Remove outliers
for (i in 1:nrow(CLEAN.DATA))
{
  CUR <- subset(CLEAN.DATA, STRAIN == CLEAN.DATA[i,"STRAIN"])
  if (CLEAN.DATA[i,"YFP.MEDIAN.CORRECT"] > median(CUR$YFP.MEDIAN.CORRECT) + 4*mad(CUR$YFP.MEDIAN.CORRECT) | CLEAN.DATA[i,"YFP.MEDIAN.CORRECT"] < median(CUR$YFP.MEDIAN.CORRECT) - 4*mad(CUR$YFP.MEDIAN.CORRECT)) {
    CLEAN.DATA[i,"MEDIAN.OUTLIER"] <- "YES"
  } else {
    CLEAN.DATA[i,"MEDIAN.OUTLIER"] <- "NO"
  }
  if (CLEAN.DATA[i,"YFP.SD.CORRECT"] > median(CUR$YFP.SD.CORRECT) + 4*mad(CUR$YFP.SD.CORRECT) | CLEAN.DATA[i,"YFP.SD.CORRECT"] < median(CUR$YFP.SD.CORRECT) - 4*mad(CUR$YFP.SD.CORRECT)) {
    CLEAN.DATA[i,"SD.OUTLIER"] <- "YES"
  } else {
    CLEAN.DATA[i,"SD.OUTLIER"] <- "NO"
  }
  if (CLEAN.DATA[i,"FSC.CORRECT"] > median(CUR$FSC.CORRECT) + 4*mad(CUR$FSC.CORRECT) | CLEAN.DATA[i,"FSC.CORRECT"] < median(CUR$FSC.CORRECT) - 4*mad(CUR$FSC.CORRECT)) {
    CLEAN.DATA[i,"FSC.OUTLIER"] <- "YES"
  } else {
    CLEAN.DATA[i,"FSC.OUTLIER"] <- "NO"
  }	
}


CLEAN.DATA <- subset(CLEAN.DATA, MEDIAN.OUTLIER == "NO" & SD.OUTLIER == "NO" & FSC.OUTLIER == "NO")

#Write processed data to file
write.table(CLEAN.DATA,"GPD1.CLEAN.CORRECT_10112016_DCv34.csv",sep=",",quote=FALSE,row.names=FALSE)

########################################################################
#                          Control plots                               #
#                                                                      #
# These plots will be removed for brevity for the following promoters  #
#                                                                      #
########################################################################

CONTROL <- droplevels(subset(CLEAN.DATA,CLEAN.DATA$CONDITION == "CTRL"))
SHAM <- droplevels(subset(CLEAN.DATA,CLEAN.DATA$CONDITION == "SHAM"))
TDH3.SHAM <- droplevels(subset(CLEAN.DATA,CLEAN.DATA$CONDITION == "TDH3.SHAM"))


#YFP MEDIAN
par(mfrow = c(2,1))
plot(CONTROL$YFP.MEDIAN.FINAL         ~ CONTROL$FLOW.RUN,       xlab = "Run", ylab = "YFP.MEAN", pch = 19, cex = 0.15, ylim=c(0.6,0.8), main="Median YFP by flow run - Before correction")
plot(CONTROL$YFP.MEDIAN.CORRECT ~ CONTROL$FLOW.RUN,       xlab = "Run", ylab = "YFP.MEAN", pch = 19, cex = 0.15, ylim=c(0.4,0.8), main="After correction")
par(mfrow = c(2,1))
plot(SHAM$YFP.MEDIAN.FINAL         ~ SHAM$FLOW.RUN,       xlab = "Run", ylab = "YFP.MEAN", pch = 19, cex = 0.15, ylim=c(0.6,0.8), main=" SHAM Median YFP by flow run - Before correction")
plot(SHAM$YFP.MEDIAN.CORRECT ~ SHAM$FLOW.RUN,       xlab = "Run", ylab = "YFP.MEAN", pch = 19, cex = 0.15, ylim=c(0.6,0.8), main="After correction")
par(mfrow = c(2,1))
plot(TDH3.SHAM$YFP.MEDIAN.FINAL         ~ TDH3.SHAM$FLOW.RUN,       xlab = "Run", ylab = "YFP.MEAN", pch = 19, cex = 0.15, ylim=c(0.8,1.1), main="TDH3 SHAM Median YFP by flow run - Before correction")
plot(TDH3.SHAM$YFP.MEDIAN.CORRECT ~ TDH3.SHAM$FLOW.RUN,       xlab = "Run", ylab = "YFP.MEAN", pch = 19, cex = 0.15, ylim=c(0.8,1.1), main="After correction")


par(mfrow = c(2,1))
plot(CONTROL$YFP.MEDIAN.FINAL         ~ CONTROL$ROW,       xlab = "ROW", ylab = "YFP.MEAN", pch = 19, cex = 0.15, ylim=c(0.6,0.8), main="Median YFP by row - Before correction")
plot(CONTROL$YFP.MEDIAN.CORRECT ~ CONTROL$ROW,       xlab = "ROW", ylab = "YFP.MEAN", pch = 19, cex = 0.15, ylim=c(0.6,0.8), main="After correction")

par(mfrow = c(2,1))
plot(CONTROL$YFP.MEDIAN.FINAL         ~ CONTROL$COLUMN,       xlab = "COLUMN", ylab = "YFP.MEAN", pch = 19, cex = 0.15,  ylim=c(0.6,0.8),main="Median YFP by column - Before correction")
plot(CONTROL$YFP.MEDIAN.CORRECT ~ CONTROL$COLUMN,       xlab = "COLUMN", ylab = "YFP.MEAN", pch = 19, cex = 0.15,  ylim=c(0.6,0.8),main="After correction")

par(mfrow = c(2,1))
plot(CONTROL$YFP.MEDIAN.FINAL         ~ CONTROL$PLATE,       xlab = "Plate", ylab = "YFP.MEAN", pch = 19, cex = 0.15, ylim=c(0.6,0.8), main="Median YFP by plate - Before correction")
plot(CONTROL$YFP.MEDIAN.CORRECT ~ CONTROL$PLATE,       xlab = "Plate", ylab = "YFP.MEAN", pch = 19, cex = 0.15, ylim=c(0.6,0.8), main="After correction")

par(mfrow = c(2,1))
plot(CONTROL$YFP.MEDIAN.FINAL         ~ CONTROL$REP,       xlab = "REPLICATE", ylab = "YFP.MEAN", pch = 19, cex = 0.15, ylim=c(0.6,0.8), main="Median YFP by replicate plate - Before correction")
plot(CONTROL$YFP.MEDIAN.CORRECT ~ CONTROL$REP,       xlab = "REPLICATE", ylab = "YFP.MEAN", pch = 19, cex = 0.15, ylim=c(0.6,0.8), main="After correction")

par(mfrow = c(2,1))
plot(CONTROL$YFP.MEDIAN.FINAL         ~ CONTROL$POSITION,       xlab = "POSITION", ylab = "YFP.MEAN", pch = 19, cex = 0.15, ylim=c(0.6,0.8), main="Median YFP by position - Before correction")
plot(CONTROL$YFP.MEDIAN.CORRECT ~ CONTROL$POSITION,       xlab = "POSITION", ylab = "YFP.MEAN", pch = 19, cex = 0.15, ylim=c(0.6,0.8), main="After correction")
par(mfrow = c(2,1))
plot(SHAM$YFP.MEDIAN.FINAL         ~ SHAM$POSITION,       xlab = "POSITION", ylab = "YFP.MEAN", pch = 19, cex = 0.15, ylim=c(0.6,0.8), main="SHAM Median YFP by position - Before correction")
plot(SHAM$YFP.MEDIAN.CORRECT ~ SHAM$POSITION,       xlab = "POSITION", ylab = "YFP.MEAN", pch = 19, cex = 0.15, ylim=c(0.6,0.8), main="After correction")
par(mfrow = c(2,1))
plot(TDH3.SHAM$YFP.MEDIAN.FINAL         ~ TDH3.SHAM$POSITION,       xlab = "POSITION", ylab = "YFP.MEAN", pch = 19, cex = 0.15, ylim=c(0.8,1.1), main="TDH3.SHAM Median YFP by position - Before correction")
plot(TDH3.SHAM$YFP.MEDIAN.CORRECT ~ TDH3.SHAM$POSITION,       xlab = "POSITION", ylab = "YFP.MEAN", pch = 19, cex = 0.15, ylim=c(0.8,1.1), main="After correction")

#YFP SD
par(mfrow = c(2,1))
plot(CONTROL$YFP.SD.FINAL         ~ CONTROL$FLOW.RUN,       xlab = "Run", ylab = "YFP.SD", pch = 19, cex = 0.15, ylim=c(0,0.04), main="YFP.SD by flow run - Before correction")
plot(CONTROL$YFP.SD.CORRECT ~ CONTROL$FLOW.RUN,       xlab = "Run", ylab = "YFP.SD", pch = 19, cex = 0.15,ylim=c(0,0.04), main="After correction")
par(mfrow = c(2,1))
plot(SHAM$YFP.SD.FINAL         ~ SHAM$FLOW.RUN,       xlab = "Run", ylab = "YFP.SD", pch = 19, cex = 0.15, ylim=c(0,0.04), main="SHAM YFP.SD by flow run - Before correction")
plot(SHAM$YFP.SD.CORRECT ~ SHAM$FLOW.RUN,       xlab = "Run", ylab = "YFP.SD", pch = 19, cex = 0.15,ylim=c(0,0.04), main="After correction")
par(mfrow = c(2,1))
plot(TDH3.SHAM$YFP.SD.FINAL         ~ TDH3.SHAM$FLOW.RUN,       xlab = "Run", ylab = "YFP.SD", pch = 19, cex = 0.15, ylim=c(0,0.08), main="TDH3SHAM YFP.SD by flow run - Before correction")
plot(TDH3.SHAM$YFP.SD.CORRECT ~ TDH3.SHAM$FLOW.RUN,       xlab = "Run", ylab = "YFP.SD", pch = 19, cex = 0.15,ylim=c(0,0.08), main="After correction")

par(mfrow = c(2,1))
plot(CONTROL$YFP.SD.FINAL         ~ CONTROL$ROW,       xlab = "ROW", ylab = "YFP.SD", pch = 19, cex = 0.15, ylim=c(0,0.04), main="YFP.SD by row - Before correction")
plot(CONTROL$YFP.SD.CORRECT ~ CONTROL$ROW,       xlab = "ROW", ylab = "YFP.SD", pch = 19, cex = 0.15, ylim=c(0,0.04), main="After correction")

par(mfrow = c(2,1))
plot(CONTROL$YFP.SD.FINAL         ~ CONTROL$COLUMN,       xlab = "COLUMN", ylab = "YFP.SD", pch = 19, cex = 0.15, ylim=c(0,0.04), main="YFP.SD by column - Before correction")
plot(CONTROL$YFP.SD.CORRECT ~ CONTROL$COLUMN,       xlab = "COLUMN", ylab = "YFP.SD", pch = 19, cex = 0.15, ylim=c(0,0.04), main="After correction")

par(mfrow = c(2,1))
plot(CONTROL$YFP.SD.FINAL         ~ CONTROL$PLATE,       xlab = "Plate", ylab = "YFP.SD", pch = 19, cex = 0.15, ylim=c(0,0.04), main="YFP.SD by plate - Before correction")
plot(CONTROL$YFP.SD.CORRECT ~ CONTROL$PLATE,       xlab = "Plate", ylab = "YFP.SD", pch = 19, cex = 0.15, ylim=c(0,0.04), main="After correction")

par(mfrow = c(2,1))
plot(CONTROL$YFP.SD.FINAL         ~ CONTROL$REP,       xlab = "REPLICATE", ylab = "YFP.SD", pch = 19, cex = 0.15, ylim=c(0,0.04), main="YFP.SD by replicate plate - Before correction")
plot(CONTROL$YFP.SD.CORRECT ~ CONTROL$REP,       xlab = "REPLICATE", ylab = "YFP.SD", pch = 19, cex = 0.15, ylim=c(0,0.04), main="After correction")

par(mfrow = c(2,1))
plot(CONTROL$YFP.SD.FINAL         ~ CONTROL$POSITION,       xlab = "POSITION", ylab = "YFP.SD", pch = 19, cex = 0.15, ylim=c(0,0.04), main="YFP.SD by position - Before correction")
plot(CONTROL$YFP.SD.CORRECT ~ CONTROL$POSITION,       xlab = "POSITION", ylab = "YFP.SD", pch = 19, cex = 0.15, ylim=c(0,0.04), main="After correction")
par(mfrow = c(2,1))
plot(SHAM$YFP.SD.FINAL         ~ SHAM$POSITION,       xlab = "POSITION", ylab = "YFP.SD", pch = 19, cex = 0.15, ylim=c(0,0.04), main="YFP.SD by position - Before correction")
plot(SHAM$YFP.SD.CORRECT ~ SHAM$POSITION,       xlab = "POSITION", ylab = "YFP.SD", pch = 19, cex = 0.15, ylim=c(0,0.04), main="After correction")

par(mfrow = c(2,1))
plot(CONTROL$YFP.SD.FINAL         ~ CONTROL$POSITION,       xlab = "POSITION", ylab = "YFP.SD", pch = 19, cex = 0.15, ylim=c(0,0.04), main="YFP.SD by position - Before correction")
plot(CONTROL$YFP.SD.CORRECT ~ CONTROL$POSITION,       xlab = "POSITION", ylab = "YFP.SD", pch = 19, cex = 0.15, ylim=c(0,0.04), main="After correction")
par(mfrow = c(2,1))
plot(SHAM$YFP.SD.FINAL         ~ SHAM$POSITION,       xlab = "POSITION", ylab = "YFP.SD", pch = 19, cex = 0.15, ylim=c(0,0.04), main="SHAM YFP.SD by position - Before correction")
plot(SHAM$YFP.SD.CORRECT ~ SHAM$POSITION,       xlab = "POSITION", ylab = "YFP.SD", pch = 19, cex = 0.15, ylim=c(0,0.04), main="After correction")
par(mfrow = c(2,1))
plot(TDH3.SHAM$YFP.SD.FINAL         ~ TDH3.SHAM$POSITION,       xlab = "POSITION", ylab = "YFP.SD", pch = 19, cex = 0.15, ylim=c(0,0.08), main="TDH3.SHAM YFP.SD by position - Before correction")
plot(TDH3.SHAM$YFP.SD.CORRECT ~ TDH3.SHAM$POSITION,       xlab = "POSITION", ylab = "YFP.SD", pch = 19, cex = 0.15, ylim=c(0,0.08), main="After correction")


#YFP MAD
par(mfrow = c(2,1))
plot(CONTROL$YFP.MAD.FINAL         ~ CONTROL$FLOW.RUN,       xlab = "Run", ylab = "YFP.MAD", pch = 19, cex = 0.15, ylim=c(0,0.04), main="YFP.MAD by flow run - Before correction")
plot(CONTROL$YFP.MAD.CORRECT ~ CONTROL$FLOW.RUN,       xlab = "Run", ylab = "YFP.MAD", pch = 19, cex = 0.15,ylim=c(0,0.04), main="After correction")
par(mfrow = c(2,1))
plot(SHAM$YFP.MAD.FINAL         ~ SHAM$FLOW.RUN,       xlab = "Run", ylab = "YFP.MAD", pch = 19, cex = 0.15, ylim=c(0,0.04), main="SHAM YFP.MAD by flow run - Before correction")
plot(SHAM$YFP.MAD.CORRECT ~ SHAM$FLOW.RUN,       xlab = "Run", ylab = "YFP.MAD", pch = 19, cex = 0.15,ylim=c(0,0.04), main="After correction")

par(mfrow = c(2,1))
plot(CONTROL$YFP.MAD.FINAL         ~ CONTROL$ROW,       xlab = "ROW", ylab = "YFP.MAD", pch = 19, cex = 0.15, ylim=c(0,0.04), main="YFP.MAD by row - Before correction")
plot(CONTROL$YFP.MAD.CORRECT ~ CONTROL$ROW,       xlab = "ROW", ylab = "YFP.MAD", pch = 19, cex = 0.15, ylim=c(0,0.04), main="After correction")

par(mfrow = c(2,1))
plot(CONTROL$YFP.MAD.FINAL         ~ CONTROL$COLUMN,       xlab = "COLUMN", ylab = "YFP.MAD", pch = 19, cex = 0.15, ylim=c(0,0.04), main="YFP.MAD by column - Before correction")
plot(CONTROL$YFP.MAD.CORRECT ~ CONTROL$COLUMN,       xlab = "COLUMN", ylab = "YFP.MAD", pch = 19, cex = 0.15, ylim=c(0,0.04), main="After correction")

par(mfrow = c(2,1))
plot(CONTROL$YFP.MAD.FINAL         ~ CONTROL$PLATE,       xlab = "Plate", ylab = "YFP.MAD", pch = 19, cex = 0.15, ylim=c(0,0.04), main="YFP.MAD by plate - Before correction")
plot(CONTROL$YFP.MAD.CORRECT ~ CONTROL$PLATE,       xlab = "Plate", ylab = "YFP.MAD", pch = 19, cex = 0.15, ylim=c(0,0.04), main="After correction")

par(mfrow = c(2,1))
plot(CONTROL$YFP.MAD.FINAL         ~ CONTROL$REP,       xlab = "REPLICATE", ylab = "YFP.MAD", pch = 19, cex = 0.15, ylim=c(0,0.04), main="YFP.MAD by replicate plate - Before correction")
plot(CONTROL$YFP.MAD.CORRECT ~ CONTROL$REP,       xlab = "REPLICATE", ylab = "YFP.MAD", pch = 19, cex = 0.15, ylim=c(0,0.04), main="After correction")

par(mfrow = c(2,1))
plot(CONTROL$YFP.MAD.FINAL         ~ CONTROL$POSITION,       xlab = "POSITION", ylab = "YFP.MAD", pch = 19, cex = 0.15, ylim=c(0,0.04), main="YFP.MAD by position - Before correction")
plot(CONTROL$YFP.MAD.CORRECT ~ CONTROL$POSITION,       xlab = "POSITION", ylab = "YFP.MAD", pch = 19, cex = 0.15, ylim=c(0,0.04), main="After correction")
par(mfrow = c(2,1))
plot(SHAM$YFP.MAD.FINAL         ~ SHAM$POSITION,       xlab = "POSITION", ylab = "YFP.MAD", pch = 19, cex = 0.15, ylim=c(0,0.04), main="SHAM YFP.MAD by position - Before correction")
plot(SHAM$YFP.MAD.CORRECT ~ SHAM$POSITION,       xlab = "POSITION", ylab = "YFP.MAD", pch = 19, cex = 0.15, ylim=c(0,0.04), main="After correction")
par(mfrow = c(2,1))
plot(TDH3.SHAM$YFP.MAD.FINAL         ~ TDH3.SHAM$POSITION,       xlab = "POSITION", ylab = "YFP.MAD", pch = 19, cex = 0.15, ylim=c(0,0.09), main="TDH3.YFP.MAD by position - Before correction")
plot(TDH3.SHAM$YFP.MAD.CORRECT ~ TDH3.SHAM$POSITION,       xlab = "POSITION", ylab = "YFP.MAD", pch = 19, cex = 0.15, ylim=c(0,0.09), main="After correction")


#FSC MEDIAN
par(mfrow = c(2,1))
plot(CONTROL$FSC.MEDIAN.FINAL         ~ CONTROL$FLOW.RUN,       xlab = "Plate", ylab = "FSC.MEDIAN", pch = 19, cex = 0.15, ylim=c(5.2,5.9), main="FSC by flow run - Before correction")
plot(CONTROL$FSC.CORRECT ~ CONTROL$FLOW.RUN,       xlab = "Plate", ylab = "FSC.MEDIAN", pch = 19, cex = 0.15, ylim=c(5.2,5.9), main="After correction")
par(mfrow = c(2,1))
plot(SHAM$FSC.MEDIAN.FINAL         ~ SHAM$FLOW.RUN,       xlab = "Plate", ylab = "FSC.MEDIAN", pch = 19, cex = 0.15, ylim=c(5.2,5.9), main="FSC by flow run - Before correction")
plot(SHAM$FSC.CORRECT ~ SHAM$FLOW.RUN,       xlab = "Plate", ylab = "FSC.MEDIAN", pch = 19, cex = 0.15, ylim=c(5.2,5.9), main="After correction")


par(mfrow = c(2,1))
plot(CONTROL$FSC.MEDIAN.FINAL         ~ CONTROL$ROW,       xlab = "ROW", ylab = "FSC.MEDIAN", pch = 19, cex = 0.15, ylim=c(5.2,5.9), main="FSC by row - Before correction")
plot(CONTROL$FSC.CORRECT ~ CONTROL$ROW,       xlab = "ROW", ylab = "FSC.MEDIAN", pch = 19, cex = 0.15, ylim=c(5.2,5.9), main="After correction")

par(mfrow = c(2,1))
plot(CONTROL$FSC.MEDIAN.FINAL         ~ CONTROL$COLUMN,       xlab = "COLUMN", ylab = "FSC.MEDIAN", pch = 19, cex = 0.15, ylim=c(5.2,5.9), main="FSC by column - Before correction")
plot(CONTROL$FSC.CORRECT ~ CONTROL$COLUMN,       xlab = "COLUMN", ylab = "FSC.MEDIAN", pch = 19, cex = 0.15, ylim=c(5.2,5.9), main="After correction")

par(mfrow = c(2,1))
plot(CONTROL$FSC.MEDIAN.FINAL         ~ CONTROL$PLATE,       xlab = "Plate", ylab = "FSC.MEDIAN", pch = 19, ylim=c(5.2,5.9), cex = 0.15, main="FSC by plate - Before correction")
plot(CONTROL$FSC.CORRECT ~ CONTROL$PLATE,       xlab = "Plate", ylab = "FSC.MEDIAN", pch = 19, cex = 0.15, ylim=c(5.2,5.9), main="After correction")

par(mfrow = c(2,1))
plot(CONTROL$FSC.MEDIAN.FINAL         ~ CONTROL$REP,       xlab = "REPLICATE", ylab = "FSC.MEDIAN", pch = 19, cex = 0.15, ylim=c(5.2,5.9), main="FSC by replicate plate - Before correction")
plot(CONTROL$FSC.CORRECT ~ CONTROL$REP,       xlab = "REPLICATE", ylab = "FSC.MEDIAN", pch = 19, cex = 0.15, ylim=c(5.2,5.9), main="After correction")

par(mfrow = c(2,1))
plot(CONTROL$FSC.MEDIAN.FINAL         ~ CONTROL$POSITION,       xlab = "POSITION", ylab = "FSC.MEDIAN", pch = 19, cex = 0.15, ylim=c(5.2,5.9), main="FSC by position - Before correction")
plot(CONTROL$FSC.CORRECT ~ CONTROL$POSITION,       xlab = "POSITION", ylab = "FSC.MEDIAN", pch = 19, cex = 0.15, ylim=c(5.2,5.9), main="After correction")
par(mfrow = c(2,1))
plot(SHAM$FSC.MEDIAN.FINAL         ~ SHAM$POSITION,       xlab = "POSITION", ylab = "FSC.MEDIAN", pch = 19, cex = 0.15, ylim=c(5.2,5.9), main="SHAM FSC by position - Before correction")
plot(SHAM$FSC.CORRECT ~ SHAM$POSITION,       xlab = "POSITION", ylab = "FSC.MEDIAN", pch = 19, cex = 0.15, ylim=c(5.2,5.9), main="After correction")
par(mfrow = c(2,1))
plot(TDH3.SHAM$FSC.MEDIAN.FINAL         ~ TDH3.SHAM$POSITION,       xlab = "POSITION", ylab = "FSC.MEDIAN", pch = 19, cex = 0.15, ylim=c(5.2,5.9), main="TDH3 SHAM FSC by position - Before correction")
plot(TDH3.SHAM$FSC.CORRECT ~ TDH3.SHAM$POSITION,       xlab = "POSITION", ylab = "FSC.MEDIAN", pch = 19, cex = 0.15, ylim=c(5.2,5.9), main="After correction")
dev.off()


###############################################################################
#
#           Strain Estimates
#
###############################################################################

# Clear memory
rm(list=ls())
options(warn=-1)

setwd("E:/wittkopp data/processed data/RNA scale analyses/DCv34")

#GPD1
CLEAN.DATA <- read.csv("GPD1.CLEAN.CORRECT_10112016_DCv34.csv", header=TRUE)

#Calculate mean and noise for each sample relative to controls
TDH3.SHAM <- subset(CLEAN.DATA, CLEAN.DATA$CONDITION == "TDH3.SHAM")
NEG <- subset(CLEAN.DATA, CLEAN.DATA$CONDITION == "NULL")
WT.SHAM <- subset(CLEAN.DATA, CLEAN.DATA$CONDITION == "SHAM")

#ABSOLUTE EXPRESSION SCALE
CLEAN.DATA[,"YFP.MEDIAN.ABS"] <- (CLEAN.DATA$YFP.MEDIAN.CORRECT - median(NEG$YFP.MEDIAN.CORRECT))
CLEAN.DATA[,"YFP.SD.ABS"]   <- CLEAN.DATA$YFP.SD.CORRECT - median(NEG$YFP.SD.CORRECT)
CLEAN.DATA[,"YFP.MAD.ABS"]   <- CLEAN.DATA$YFP.MAD.CORRECT - median(NEG$YFP.SD.CORRECT)

#Re-calculate mean and noise for each sample relative to controls
TDH3.SHAM <- subset(CLEAN.DATA, CLEAN.DATA$CONDITION == "TDH3.SHAM")
NEG <- subset(CLEAN.DATA, CLEAN.DATA$CONDITION == "NULL")
WT.SHAM <- subset(CLEAN.DATA, CLEAN.DATA$CONDITION == "SHAM")

#PROMOTER-SPECIFIC SCALING
CLEAN.DATA[,"YFP.MEDIAN.WT"] <- CLEAN.DATA$YFP.MEDIAN.ABS/median(WT.SHAM$YFP.MEDIAN.ABS)
CLEAN.DATA[,"YFP.SD.WT"]   <- CLEAN.DATA$YFP.SD.ABS/median(WT.SHAM$YFP.MEDIAN.ABS)
CLEAN.DATA[,"YFP.MAD.WT"]   <- CLEAN.DATA$YFP.MAD.ABS/median(WT.SHAM$YFP.MEDIAN.ABS)

#RELATIVE TO TDH3 EXPRESSION
CLEAN.DATA[,"YFP.MEDIAN.TDH3"] <- CLEAN.DATA$YFP.MEDIAN.ABS/median(TDH3.SHAM$YFP.MEDIAN.ABS)
CLEAN.DATA[,"YFP.SD.TDH3"]   <- CLEAN.DATA$YFP.SD.ABS/median(TDH3.SHAM$YFP.MEDIAN.ABS)
CLEAN.DATA[,"YFP.MAD.TDH3"]   <- CLEAN.DATA$YFP.MAD.ABS/median(TDH3.SHAM$YFP.MEDIAN.ABS)

#Re-calculate mean and noise for each sample relative to controls
TDH3.SHAM <- subset(CLEAN.DATA, CLEAN.DATA$CONDITION == "TDH3.SHAM")
TDH3.SHAM<-subset(TDH3.SHAM,TDH3.SHAM$YFP.MEDIAN.FINAL>0.7)
NEG <- subset(CLEAN.DATA, CLEAN.DATA$CONDITION == "NULL")
WT.SHAM <- subset(CLEAN.DATA, CLEAN.DATA$CONDITION == "SHAM")

#CALCULATE NOISE METRICS
CLEAN.DATA[,"YFP.CV.WT"]   <- CLEAN.DATA$YFP.SD.WT/CLEAN.DATA$YFP.MEDIAN.WT
CLEAN.DATA[,"YFP.CV.TDH3"]   <- CLEAN.DATA$YFP.SD.TDH3/CLEAN.DATA$YFP.MEDIAN.TDH3
CLEAN.DATA[,"YFP.FANO.WT"]   <- CLEAN.DATA$YFP.SD.WT^2/CLEAN.DATA$YFP.MEDIAN.WT
CLEAN.DATA[,"YFP.FANO.TDH3"]   <- CLEAN.DATA$YFP.SD.TDH3^2/CLEAN.DATA$YFP.MEDIAN.TDH3


#Re-calculate mean and noise for each sample relative to controls
TDH3.SHAM <- subset(CLEAN.DATA, CLEAN.DATA$CONDITION == "TDH3.SHAM")
TDH3.SHAM<-subset(TDH3.SHAM,TDH3.SHAM$YFP.MEDIAN.FINAL>0.7)
NEG <- subset(CLEAN.DATA, CLEAN.DATA$CONDITION == "NULL")
WT.SHAM <- subset(CLEAN.DATA, CLEAN.DATA$CONDITION == "SHAM")

#CALCULATE RELATIVE NOISE METRICS
CLEAN.DATA[,"YFP.SD.WT.REL"]   <- CLEAN.DATA$YFP.SD.WT/median(WT.SHAM$YFP.SD.WT)
CLEAN.DATA[,"YFP.SD.TDH3.REL"]   <- CLEAN.DATA$YFP.SD.TDH3/median(WT.SHAM$YFP.SD.TDH3)
CLEAN.DATA[,"YFP.CV.WT.REL"]   <- CLEAN.DATA$YFP.SD.WT.REL/CLEAN.DATA$YFP.MEDIAN.WT
CLEAN.DATA[,"YFP.CV.TDH3.REL"]   <- CLEAN.DATA$YFP.SD.TDH3.REL/CLEAN.DATA$YFP.MEDIAN.TDH3
CLEAN.DATA[,"YFP.FANO.WT.REL"]   <- CLEAN.DATA$YFP.SD.WT.REL^2/CLEAN.DATA$YFP.MEDIAN.WT
CLEAN.DATA[,"YFP.FANO.TDH3.REL"]   <- CLEAN.DATA$YFP.SD.TDH3.REL^2/CLEAN.DATA$YFP.MEDIAN.TDH3


#Add NA for samples where CV or FANO are undefined.
for (i in 1:nrow(CLEAN.DATA))
{
  if (CLEAN.DATA[i,"YFP.MEDIAN.ABS"] < 0 | CLEAN.DATA[i,"YFP.SD.ABS"] < 0) {
    CLEAN.DATA[i,"YFP.MAD.WT"] <- NA
    CLEAN.DATA[i,"YFP.MAD.TDH3"] <- NA
    CLEAN.DATA[i,"YFP.CV.WT"] <- NA
    CLEAN.DATA[i,"YFP.CV.TDH3"] <- NA
    CLEAN.DATA[i,"YFP.CV.WT.REL"] <- NA
    CLEAN.DATA[i,"YFP.CV.TDH3.REL"] <- NA
    CLEAN.DATA[i,"YFP.FANO.WT"] <- NA
    CLEAN.DATA[i,"YFP.FANO.TDH3"] <- NA
    CLEAN.DATA[i,"YFP.FANO.WT.REL"] <- NA
    CLEAN.DATA[i,"YFP.FANO.TDH3.REL"] <- NA
    
  }
}

CLEAN.DATA <- droplevels(subset(CLEAN.DATA,CLEAN.DATA$CONDITION!="CTRL"))

write.table(CLEAN.DATA,"GPD1.CLEAN.ADJUST_10112016_DCv34.csv",sep=",",quote=FALSE,row.names=FALSE)

CLEAN.DATA <- read.csv("GPD1.CLEAN.ADJUST_10112016_DCv34.csv", header=TRUE)

GPD1.MEAN <- aggregate(cbind(YFP.MEDIAN.ABS,YFP.SD.ABS,YFP.MAD.ABS,YFP.MEDIAN.WT,YFP.SD.WT,YFP.MAD.WT,YFP.MEDIAN.TDH3,YFP.SD.TDH3,YFP.MAD.TDH3,YFP.CV.WT,YFP.FANO.WT,YFP.CV.TDH3,YFP.FANO.TDH3,YFP.SD.WT.REL,YFP.CV.WT.REL,YFP.FANO.WT.REL,YFP.SD.TDH3.REL,YFP.CV.TDH3.REL,YFP.FANO.TDH3.REL,FSC.CORRECT) ~ STRAIN + CONDITION + ASSAY + PLATE + CTRL.GENO, data=CLEAN.DATA, FUN = mean)
GPD1.SD <- aggregate(cbind(YFP.MEDIAN.ABS,YFP.SD.ABS,YFP.MAD.ABS,YFP.MEDIAN.WT,YFP.SD.WT,YFP.MAD.WT,YFP.MEDIAN.TDH3,YFP.SD.TDH3,YFP.MAD.TDH3,YFP.CV.WT,YFP.FANO.WT,YFP.CV.TDH3,YFP.FANO.TDH3,YFP.SD.WT.REL,YFP.CV.WT.REL,YFP.FANO.WT.REL,YFP.SD.TDH3.REL,YFP.CV.TDH3.REL,YFP.FANO.TDH3.REL,FSC.CORRECT) ~ STRAIN + CONDITION + ASSAY + PLATE + CTRL.GENO, data=CLEAN.DATA, FUN = sd)
GPD1.N <- aggregate(cbind(YFP.MEDIAN.ABS,YFP.SD.ABS,YFP.MAD.ABS,YFP.MEDIAN.WT,YFP.SD.WT,YFP.MAD.WT,YFP.MEDIAN.TDH3,YFP.SD.TDH3,YFP.MAD.TDH3,YFP.CV.WT,YFP.FANO.WT,YFP.CV.TDH3,YFP.FANO.TDH3,YFP.SD.WT.REL,YFP.CV.WT.REL,YFP.FANO.WT.REL,YFP.SD.TDH3.REL,YFP.CV.TDH3.REL,YFP.FANO.TDH3.REL,FSC.CORRECT) ~ STRAIN + CONDITION + ASSAY + PLATE + CTRL.GENO, data=CLEAN.DATA, FUN = length)

GPD1.MEAN  <- GPD1.MEAN[which(GPD1.N[,6] >= 3),]
GPD1.SD <- GPD1.SD[which(GPD1.N[,6] >= 3),]
GPD1.N <- GPD1.N[which(GPD1.N[,6] >= 3),]

GPD1 <- cbind.data.frame(GPD1.MEAN,GPD1.SD[,6:25],GPD1.N[,6])

colnames(GPD1) <- c("STRAIN","CONDITION","ASSAY","PLATE","CTRL.GENO",paste0("MEAN.",colnames(GPD1.MEAN[,6:25])),paste0("SD.",colnames(GPD1.SD[,6:25])),"N")

write.table(GPD1,"GPD1.SUMMARY.DATA_10112016_DCv34.txt",sep="\t",quote=FALSE,row.names=FALSE)

CONTAM.CHECK<-droplevels(subset(CLEAN.DATA,CLEAN.DATA$CONTAM.CHECK!="NONE"))

`%notin%` <- function(x,y) !(x %in% y)

if(all(CONTAM.CHECK$STRAIN %notin% GPD1$STRAIN)) {
  #All datapoints flagged as suspicious have been removed by outlier removal filters.
  print("Yay! No suspicious values detected. Proceed.")
}

if(any(CONTAM.CHECK$STRAIN %in% GPD1$STRAIN)) {
  CONTAM.CHECK<-CONTAM.CHECK[CONTAM.CHECK$STRAIN %in% CONTAM.CHECK$STRAIN[which(CONTAM.CHECK$STRAIN %in% GPD1$STRAIN)],]
  write.table(CONTAM.CHECK,"GPD1.CONTAM.CHECK_outliersexcl.csv",sep=",",quote=FALSE,row.names=FALSE)  
}


###############################
#HIERARCHICAL PERMUTATION TEST#
###############################

#Clear memory
rm(list=ls())
options(warn=-1)

library(permute)
library(Hmisc)

N.PERM <- 10000

#Permutation function

SHUFFLE <- function(STRAIN.LIST = STRAIN.LIST[[i]],WT.LIST = WT.LIST) {
  SAMPLE <- c(STRAIN.LIST,WT.LIST[[sample(1:length(WT.LIST),1)]])
  OUTPUT <- sample(SAMPLE,length(SAMPLE))
  DIST.SIM <- abs(mean(OUTPUT[1:length(STRAIN.LIST)]) - mean(OUTPUT[(length(STRAIN.LIST)+1):length(SAMPLE)]))
  DIST.OBS <- abs(mean(SAMPLE[1:length(STRAIN.LIST)]) - mean(SAMPLE[(length(STRAIN.LIST)+1):length(SAMPLE)]))
  
  DIFF <- DIST.OBS - DIST.SIM
  
  return(DIFF)
}

parent.dir <- PATH
setwd(parent.dir)

####GPD1 Analysis date 10112016#########################################################################

ALL.REP <- read.csv("GPD1.CLEAN.ADJUST_10112016_DCv34.csv",header=TRUE)
ALL.REP <- subset(ALL.REP, MEDIAN.OUTLIER == "NO" & SD.OUTLIER == "NO" & FSC.OUTLIER == "NO")

SUMMARY <- read.table("GPD1.SUMMARY.DATA_10112016_DCv34.txt",header=TRUE)

#SUMMARY <- droplevels(subset(SUMMARY, STRAIN != "1139"))

WT <- subset(ALL.REP, CONDITION == "SHAM")
LOW.FSC <- median(WT$FSC.CORRECT) - 4*mad(WT$FSC.CORRECT)
HIGH.FSC <- median(WT$FSC.CORRECT) + 4*mad(WT$FSC.CORRECT)
LOW.FLUO <- median(WT$YFP.MEDIAN.CORRECT) - 4*mad(WT$YFP.MEDIAN.CORRECT)
HIGH.FLUO <- median(WT$YFP.MEDIAN.CORRECT) + 4*mad(WT$YFP.MEDIAN.CORRECT)
WT <- subset(WT, FSC.CORRECT > LOW.FSC & FSC.CORRECT < HIGH.FSC & YFP.MEDIAN.CORRECT > LOW.FLUO & YFP.MEDIAN.CORRECT < HIGH.FLUO)

#MEDIAN

#Create list of WT phenotype grouped by strain

N.WT <- length(unique(WT$STRAIN))
WT.LIST <- vector('list',N.WT)
for (i in 1:N.WT)
{
  CUR.WT <- subset(WT, STRAIN == unique(WT$STRAIN)[i])
  WT.LIST[[i]] <- CUR.WT$YFP.MEDIAN.CORRECT 
}


#Create list of all strain phenotypes grouped by strain

SUMMARY <- SUMMARY[order(SUMMARY$STRAIN),]
ALL.REP <- droplevels(subset(ALL.REP,STRAIN %in% as.character(SUMMARY$STRAIN)))
ALL.REP <- ALL.REP[order(ALL.REP$STRAIN),]
N.STRAIN <- length(unique(ALL.REP$STRAIN))
STRAIN.LIST <- vector('list',N.STRAIN)
for (i in 1:N.STRAIN)
{
  CUR.STRAIN <- subset(ALL.REP, STRAIN == unique(ALL.REP$STRAIN)[i])
  STRAIN.LIST[[i]] <- CUR.STRAIN$YFP.MEDIAN.CORRECT
}


for (i in 1:length(STRAIN.LIST))
{
  DIFF <- sapply(1:N.PERM,FUN=function(x)	SHUFFLE(STRAIN.LIST = STRAIN.LIST[[i]],WT.LIST = WT.LIST))
  P.VAL <- (length(which(DIFF < 0)) + 1) / (length(which(DIFF < 0)) + length(which(DIFF > 0)) + 1) 	
  SUMMARY[i,"P.VAL.MEDIAN"] <- P.VAL
  print(round(i/length(STRAIN.LIST)*100,1))
}

#NOISE: CV

#Create list of WT phenotype grouped by strain

N.WT <- length(unique(WT$STRAIN))
WT.LIST <- vector('list',N.WT)
for (i in 1:N.WT)
{
  CUR.WT <- subset(WT, STRAIN == unique(WT$STRAIN)[i])
  WT.LIST[[i]] <- CUR.WT$YFP.CV.WT
}


#Create list of all strain phenotypes grouped by strain

SUMMARY <- SUMMARY[order(SUMMARY$STRAIN),]
ALL.REP <- droplevels(subset(ALL.REP,STRAIN %in% as.character(SUMMARY$STRAIN)))
ALL.REP <- ALL.REP[order(ALL.REP$STRAIN),]
N.STRAIN <- length(unique(ALL.REP$STRAIN))
STRAIN.LIST <- vector('list',N.STRAIN)
for (i in 1:N.STRAIN)
{
  CUR.STRAIN <- subset(ALL.REP, STRAIN == unique(ALL.REP$STRAIN)[i])
  STRAIN.LIST[[i]] <- CUR.STRAIN$YFP.CV.WT
}


for (i in 1:length(STRAIN.LIST))
{
  DIFF <- sapply(1:N.PERM,FUN=function(x)	SHUFFLE(STRAIN.LIST = STRAIN.LIST[[i]],WT.LIST = WT.LIST))
  P.VAL <- (length(which(DIFF < 0))+1) / (length(which(DIFF < 0)) + length(which(DIFF > 0)) + 1) 	
  SUMMARY[i,"P.VAL.CV.WT"] <- P.VAL
  print(round(i/length(STRAIN.LIST)*100,1))
}

#NOISE: FANO

#Create list of WT phenotype grouped by strain

N.WT <- length(unique(WT$STRAIN))
WT.LIST <- vector('list',N.WT)
for (i in 1:N.WT)
{
  CUR.WT <- subset(WT, STRAIN == unique(WT$STRAIN)[i])
  WT.LIST[[i]] <- CUR.WT$YFP.FANO.WT
}


#Create list of all strain phenotypes grouped by strain

SUMMARY <- SUMMARY[order(SUMMARY$STRAIN),]
ALL.REP <- droplevels(subset(ALL.REP,STRAIN %in% as.character(SUMMARY$STRAIN)))
ALL.REP <- ALL.REP[order(ALL.REP$STRAIN),]
N.STRAIN <- length(unique(ALL.REP$STRAIN))
STRAIN.LIST <- vector('list',N.STRAIN)
for (i in 1:N.STRAIN)
{
  CUR.STRAIN <- subset(ALL.REP, STRAIN == unique(ALL.REP$STRAIN)[i])
  STRAIN.LIST[[i]] <- CUR.STRAIN$YFP.FANO.WT
}


for (i in 1:length(STRAIN.LIST))
{
  DIFF <- sapply(1:N.PERM,FUN=function(x)	SHUFFLE(STRAIN.LIST = STRAIN.LIST[[i]],WT.LIST = WT.LIST))
  P.VAL <- (length(which(DIFF < 0))+1) / (length(which(DIFF < 0)) + length(which(DIFF > 0)) + 1) 	
  SUMMARY[i,"P.VAL.FANO.WT"] <- P.VAL
  print(round(i/length(STRAIN.LIST)*100,1))
}

#FSC MEDIAN

#Create list of WT phenotype grouped by strain

N.WT <- length(unique(WT$STRAIN))
WT.LIST <- vector('list',N.WT)
for (i in 1:N.WT)
{
  CUR.WT <- subset(WT, STRAIN == unique(WT$STRAIN)[i])
  WT.LIST[[i]] <- CUR.WT$FSC.CORRECT
}


#Create list of all strain phenotypes grouped by strain

SUMMARY <- SUMMARY[order(SUMMARY$STRAIN),]
ALL.REP <- droplevels(subset(ALL.REP,STRAIN %in% as.character(SUMMARY$STRAIN)))
ALL.REP <- ALL.REP[order(ALL.REP$STRAIN),]
N.STRAIN <- length(unique(ALL.REP$STRAIN))
STRAIN.LIST <- vector('list',N.STRAIN)
for (i in 1:N.STRAIN)
{
  CUR.STRAIN <- subset(ALL.REP, STRAIN == unique(ALL.REP$STRAIN)[i])
  STRAIN.LIST[[i]] <- CUR.STRAIN$FSC.CORRECT
}


for (i in 1:length(STRAIN.LIST))
{
  DIFF <- sapply(1:N.PERM,FUN=function(x)	SHUFFLE(STRAIN.LIST = STRAIN.LIST[[i]],WT.LIST = WT.LIST))
  P.VAL <- (length(which(DIFF < 0))+1) / (length(which(DIFF < 0)) + length(which(DIFF > 0)) + 1) 	
  SUMMARY[i,"P.VAL.FSC"] <- P.VAL
  print(round(i/length(STRAIN.LIST)*100,1))
}

write.table(SUMMARY,"GPD1.SUMMARY.DCv34.Pvals.txt",sep="\t",quote=FALSE,row.names=FALSE)

#_________________________________________________________________________________________________________________________

#######################
#   OST1
#######################


#Clear memory
rm(list=ls())
options(warn=-1)


#####################
#  LOADING LIBRARIES#
#####################

library(flowCore)
library(flowClust)
library(flowViz)
library(plotrix)
library(nlme)
library(MethComp)
library(outliers)
library(pcaPP)

library(reshape2)
library(MASS)
library(ggplot2)
library(Hmisc)
library(fBasics)
library(lawstat)
library(fitdistrplus)
library(mixtools)
library(vioplot)
library(gplots)
library(RColorBrewer)
library(calibrate)
library(e1071)

box <- graphics::box


#Set working directory
parent.dir <- PATH
setwd(parent.dir)

#Load experiment setup
SETUP <- read.csv("PATH/Template_OST1_10102016.csv",header=TRUE,as.is=TRUE)
TESTNAMES <- list.files("PATH/OST1",pattern=".fcs",full.name=TRUE,recursive=TRUE,include.dirs=TRUE)
SETUP<-subset(SETUP, SETUP$FILENAMES %in% TESTNAMES, drop = TRUE)
SETUP[,"COUNTER"] <- c(1:nrow(SETUP))

#Determine Hard Gates
GATES <- GATE.CALIB(SETUP$FILENAMES[1])

#Clean Data
Output <- apply(SETUP,1,CLEANING,GATES=GATES)
OUTPUT <- as.data.frame(Output[[1]])
OUTPUT[2:nrow(SETUP),1:ncol(OUTPUT)] <- NA
for (i in 2:nrow(SETUP))
{
  if (is.null(Output[[i]]))
  {} 
  else {
    OUTPUT[i,] <- Output[[i]]
  }
}

write.table(OUTPUT,"OST1_Experiment.Output_DCv34.txt",row.names=FALSE,sep="\t")

CLEAN <- cbind.data.frame(SETUP[,1:(ncol(SETUP)-2)],OUTPUT)

write.table(CLEAN,"OST1_Clean.Data_10112016_DCv34.txt",row.names=FALSE,sep="\t")
################################################################
#     PLATE CORRECTION & OUTLIER REMOVAL
################################################################

#Clear memory
rm(list=ls())
options(warn=-1)

#####################
#1-LOADING LIBRARIES#
#####################
library(flowCore)
library(flowClust)
library(flowViz)
library(pcaPP)
library(mixtools)
library(plyr)
library(MASS)
box <- graphics::box
options(warn=-1)

####################################
#2- Quality Control and Corrections#
####################################

parent.dir <- PATH
setwd(parent.dir)

DATA.TYPE <- c(
  "factor","factor","factor","factor","factor","factor","factor","factor","factor","factor","factor","factor","factor","factor","factor",
  "integer","integer","integer","integer",
  "numeric","numeric","numeric","numeric","numeric","numeric","numeric","numeric","numeric","numeric","numeric","numeric","numeric","numeric","numeric","numeric","numeric","numeric","numeric","numeric","numeric","numeric","numeric","numeric","numeric",
  "integer")

###Read in data sets
CLEAN.DATA <- read.table("OST1_Clean.Data_10112016_DCv34.txt",colClasses=DATA.TYPE,header=TRUE, sep="\t")

###Remove samples with NA
CLEAN.DATA <- CLEAN.DATA[complete.cases(CLEAN.DATA),]

##Translate into log scale for appropriate correction
CLEAN.DATA[,"log.RNA.MEDIAN"] <- log((CLEAN.DATA[,"YFP.MEDIAN.FINAL"]) + 0.05)
CLEAN.DATA[,"log.RNA.MAD"] <- CLEAN.DATA[,"YFP.MAD.FINAL"] / (CLEAN.DATA[,"YFP.MEDIAN.FINAL"] + 0.05)
CLEAN.DATA[,"log.RNA.SD"] <- CLEAN.DATA[,"YFP.SD.FINAL"] / (CLEAN.DATA[,"YFP.MEDIAN.FINAL"] + 0.05)

###Remove samples with less than 1000 events
CLEAN.DATA <- subset(CLEAN.DATA,CLEAN.DATA$COUNTS.FINAL > 1000)

####Add variable to ID whether control is own promoter or TDH3
CLEAN.DATA[,"CTRL.GENO"]<-as.factor("OST1")

CLEAN.DATA$CONTAM.CHECK<-"NONE"
CLEAN.DATA$CONTAM.CHECK[(CLEAN.DATA$CONDITION=="EMS"|CLEAN.DATA$CONDITION=="SHAM"|CLEAN.DATA$CONDITION=="EMS.LOW"|CLEAN.DATA$CONDITION=="EMS.HIGH") & (CLEAN.DATA$YFP.MEDIAN.FINAL<.53)]<-"BACT.CONTAM.CHECK"
CLEAN.DATA$CONTAM.CHECK[(CLEAN.DATA$CONDITION=="EMS"|CLEAN.DATA$CONDITION=="SHAM"|CLEAN.DATA$CONDITION=="EMS.LOW"|CLEAN.DATA$CONDITION=="EMS.HIGH") & (CLEAN.DATA$YFP.MEDIAN.FINAL>0.75)]<-"TDH3.CONTAM.CHECK"
CLEAN.DATA$CONTAM.CHECK[(CLEAN.DATA$CONDITION=="TDH3.SHAM" & CLEAN.DATA$YFP.MEDIAN.FINAL<0.8)]<-"EXP.SHAM.CONTAM.CHECK"
CLEAN.DATA$CONTAM.CHECK[(CLEAN.DATA$CONDITION=="NULL" & CLEAN.DATA$YFP.MEDIAN.FINAL>0.53)]<-"EXP.NULL.CONTAM.CHECK"


###Seperate controls and remove FSC outliers
CONTROL <- subset(CLEAN.DATA,CLEAN.DATA$CONDITION == "CTRL")
FSC.CORRECT <- lm(CONTROL$FSC.MEDIAN.FINAL ~ 0 + CONTROL$FLOW.RUN)
RESID <- FSC.CORRECT$resid
CLEAN.DATA <- droplevels(CLEAN.DATA[-which(CLEAN.DATA$CONDITION == "CTRL")[abs(RESID) > 3*mad(RESID)],])
CONTROL <- droplevels(subset(CLEAN.DATA,CLEAN.DATA$CONDITION == "CTRL"))

INITIAL.CORRECT <- lm(CONTROL$log.RNA.MEDIAN ~ 0 + CONTROL$FLOW.RUN)
RESID <- INITIAL.CORRECT$resid
CLEAN.DATA<- droplevels(CLEAN.DATA[-which(CLEAN.DATA$CONDITION == "CTRL")[abs(RESID) > 3*mad(RESID)],])
CONTROL <- droplevels(subset(CLEAN.DATA,CLEAN.DATA$CONDITION == "CTRL"))

#Remove any flow runs with insufficient # of control positions to perform full plate correction
`%notin%` <- function(x,y) !(x %in% y)
RESULT<-count(CONTROL$FLOW.RUN)
RESULT.MUT<-count(CLEAN.DATA$FLOW.RUN)
EXCLUDE1<-(RESULT.MUT[which(RESULT.MUT[,1] %notin% RESULT[,1]),1])
EXCLUDE2<-(RESULT[which(RESULT[,2] <5),1])
CLEAN.DATA<-droplevels(CLEAN.DATA[CLEAN.DATA$FLOW.RUN %notin% EXCLUDE1,])
CLEAN.DATA<-droplevels(CLEAN.DATA[CLEAN.DATA$FLOW.RUN %notin% EXCLUDE2,])
CONTROL <- subset(CLEAN.DATA,CLEAN.DATA$CONDITION == "CTRL")


###Corrections for YFP median and MAD
#Correct Median
#fit<- lm(log.RNA.MEDIAN~FLOW.RUN+REP+STACK.ORDER+STACK.ORDER:FLOW.RUN+REP+ROW+COLUMN+BLOCK+BLOCK:ROW+ROW:COLUMN+STACK.ORDER:BLOCK,data=CONTROL)
#step <- stepAIC(fit, direction="both")
log.RNA.MEDIAN.CORRECT <- lm(log.RNA.MEDIAN ~ FLOW.RUN + COLUMN + ROW + ROW:COLUMN, data=CONTROL)
CLEAN.DATA[,"log.RNA.MEDIAN.CORRECT"] <- CLEAN.DATA$log.RNA.MEDIAN - predict(log.RNA.MEDIAN.CORRECT, CLEAN.DATA) + mean(CONTROL$log.RNA.MEDIAN)
CONTROL <- subset(CLEAN.DATA,CLEAN.DATA$CONDITION == "CTRL")

#Correct sd
#fit<- lm(log.RNA.SD~FLOW.RUN+REP+STACK.ORDER+STACK.ORDER:FLOW.RUN+REP+ROW+COLUMN+BLOCK+BLOCK:ROW+ROW:COLUMN+STACK.ORDER:BLOCK,data=CONTROL)
#step <- step(fit, direction="both")
log.RNA.SD.CORRECT <- lm(log.RNA.SD ~  FLOW.RUN + ROW + COLUMN + ROW:COLUMN,data=CONTROL)
CLEAN.DATA[,"log.RNA.SD.CORRECT"] <- CLEAN.DATA$log.RNA.SD - predict(log.RNA.SD.CORRECT , CLEAN.DATA) + mean(CONTROL$log.RNA.SD)
CONTROL <- subset(CLEAN.DATA,CLEAN.DATA$CONDITION == "CTRL")

#Correct mad
#fit<- lm(log.RNA.MAD~FLOW.RUN+REP+STACK.ORDER+STACK.ORDER:FLOW.RUN+REP+ROW+COLUMN+BLOCK+BLOCK:ROW+ROW:COLUMN+STACK.ORDER:BLOCK,data=CONTROL)
#step <- step(fit, direction="both")
log.RNA.MAD.CORRECT <- lm(log.RNA.MAD ~  FLOW.RUN + ROW + COLUMN + ROW:COLUMN,data=CONTROL)
CLEAN.DATA[,"log.RNA.MAD.CORRECT"] <- CLEAN.DATA$log.RNA.MAD - predict(log.RNA.MAD.CORRECT , CLEAN.DATA) + mean(CONTROL$log.RNA.MAD)
CONTROL <- subset(CLEAN.DATA,CLEAN.DATA$CONDITION == "CTRL")

#Correct FSC
#fit<- lm(FSC.MEDIAN.FINAL~FLOW.RUN+REP+STACK.ORDER+STACK.ORDER:FLOW.RUN+REP+ROW+COLUMN+BLOCK+BLOCK:ROW+ROW:COLUMN+STACK.ORDER:BLOCK,data=CONTROL)
#step <- stepAIC(fit, direction="both")
FSC.CORRECT <- lm(FSC.MEDIAN.FINAL ~  FLOW.RUN + ROW + COLUMN + ROW:COLUMN,data=CONTROL)
CLEAN.DATA[,"FSC.CORRECT"] <- CLEAN.DATA$FSC.MEDIAN.FINAL - predict(FSC.CORRECT, CLEAN.DATA) + mean(CONTROL$FSC.MEDIAN.FINAL)
CONTROL <- subset(CLEAN.DATA,CLEAN.DATA$CONDITION == "CTRL")

#Back translate to linear scale
CLEAN.DATA[,"YFP.MEDIAN.CORRECT"] <- (exp(CLEAN.DATA[,"log.RNA.MEDIAN.CORRECT"]) - 0.05)
CLEAN.DATA[,"YFP.SD.CORRECT"] <- CLEAN.DATA[,"log.RNA.SD.CORRECT"] * CLEAN.DATA[,"YFP.MEDIAN.CORRECT"]
CLEAN.DATA[,"YFP.MAD.CORRECT"] <- CLEAN.DATA[,"log.RNA.MAD.CORRECT"] * CLEAN.DATA[,"YFP.MEDIAN.CORRECT"]


#PERFORM FABIEN'S OUTLIER CORRECTION FILTERING OUT ANY REPLICATE OUTSIDE OF 4*MAD

#Remove outliers
for (i in 1:nrow(CLEAN.DATA))
{
  CUR <- subset(CLEAN.DATA, STRAIN == CLEAN.DATA[i,"STRAIN"])
  if (CLEAN.DATA[i,"YFP.MEDIAN.CORRECT"] > median(CUR$YFP.MEDIAN.CORRECT) + 4*mad(CUR$YFP.MEDIAN.CORRECT) | CLEAN.DATA[i,"YFP.MEDIAN.CORRECT"] < median(CUR$YFP.MEDIAN.CORRECT) - 4*mad(CUR$YFP.MEDIAN.CORRECT)) {
    CLEAN.DATA[i,"MEDIAN.OUTLIER"] <- "YES"
  } else {
    CLEAN.DATA[i,"MEDIAN.OUTLIER"] <- "NO"
  }
  if (CLEAN.DATA[i,"YFP.SD.CORRECT"] > median(CUR$YFP.SD.CORRECT) + 4*mad(CUR$YFP.SD.CORRECT) | CLEAN.DATA[i,"YFP.SD.CORRECT"] < median(CUR$YFP.SD.CORRECT) - 4*mad(CUR$YFP.SD.CORRECT)) {
    CLEAN.DATA[i,"SD.OUTLIER"] <- "YES"
  } else {
    CLEAN.DATA[i,"SD.OUTLIER"] <- "NO"
  }
  if (CLEAN.DATA[i,"FSC.CORRECT"] > median(CUR$FSC.CORRECT) + 4*mad(CUR$FSC.CORRECT) | CLEAN.DATA[i,"FSC.CORRECT"] < median(CUR$FSC.CORRECT) - 4*mad(CUR$FSC.CORRECT)) {
    CLEAN.DATA[i,"FSC.OUTLIER"] <- "YES"
  } else {
    CLEAN.DATA[i,"FSC.OUTLIER"] <- "NO"
  }	
}

CLEAN.DATA <- subset(CLEAN.DATA, MEDIAN.OUTLIER == "NO" & SD.OUTLIER == "NO" & FSC.OUTLIER == "NO")

#Write processed data to file
write.table(CLEAN.DATA,"OST1.CLEAN.CORRECT_10112016_DCv34.csv",sep=",",quote=FALSE,row.names=FALSE)

#####################
#5 - Control plots  #
#####################

CONTROL <- droplevels(subset(CLEAN.DATA,CLEAN.DATA$CONDITION == "CTRL"))
SHAM <- droplevels(subset(CLEAN.DATA,CLEAN.DATA$CONDITION == "SHAM"))
TDH3.SHAM <- droplevels(subset(CLEAN.DATA,CLEAN.DATA$CONDITION == "TDH3.SHAM"))

#removed for brevity

################################################################################
#
#         Strain Estimates
#
################################################################################

#Clear memory
rm(list=ls())
options(warn=-1)

setwd(PATH)

#OST1
CLEAN.DATA <- read.csv("OST1.CLEAN.CORRECT_10112016_DCv34.csv", header=TRUE)

#Calculate mean and noise for each sample relative to controls
TDH3.SHAM <- subset(CLEAN.DATA, CLEAN.DATA$CONDITION == "TDH3.SHAM")
NEG <- subset(CLEAN.DATA, CLEAN.DATA$CONDITION == "NULL")
WT.SHAM <- subset(CLEAN.DATA, CLEAN.DATA$CONDITION == "SHAM")

#ABSOLUTE EXPRESSION SCALE
CLEAN.DATA[,"YFP.MEDIAN.ABS"] <- (CLEAN.DATA$YFP.MEDIAN.CORRECT - median(NEG$YFP.MEDIAN.CORRECT))
CLEAN.DATA[,"YFP.SD.ABS"]   <- CLEAN.DATA$YFP.SD.CORRECT - median(NEG$YFP.SD.CORRECT)
CLEAN.DATA[,"YFP.MAD.ABS"]   <- CLEAN.DATA$YFP.MAD.CORRECT - median(NEG$YFP.SD.CORRECT)

#Re-calculate mean and noise for each sample relative to controls
TDH3.SHAM <- subset(CLEAN.DATA, CLEAN.DATA$CONDITION == "TDH3.SHAM")
NEG <- subset(CLEAN.DATA, CLEAN.DATA$CONDITION == "NULL")
WT.SHAM <- subset(CLEAN.DATA, CLEAN.DATA$CONDITION == "SHAM")

#PROMOTER-SPECIFIC SCALING
CLEAN.DATA[,"YFP.MEDIAN.WT"] <- CLEAN.DATA$YFP.MEDIAN.ABS/median(WT.SHAM$YFP.MEDIAN.ABS)
CLEAN.DATA[,"YFP.SD.WT"]   <- CLEAN.DATA$YFP.SD.ABS/median(WT.SHAM$YFP.MEDIAN.ABS)
CLEAN.DATA[,"YFP.MAD.WT"]   <- CLEAN.DATA$YFP.MAD.ABS/median(WT.SHAM$YFP.MEDIAN.ABS)

#RELATIVE TO TDH3 EXPRESSION
CLEAN.DATA[,"YFP.MEDIAN.TDH3"] <- CLEAN.DATA$YFP.MEDIAN.ABS/median(TDH3.SHAM$YFP.MEDIAN.ABS)
CLEAN.DATA[,"YFP.SD.TDH3"]   <- CLEAN.DATA$YFP.SD.ABS/median(TDH3.SHAM$YFP.MEDIAN.ABS)
CLEAN.DATA[,"YFP.MAD.TDH3"]   <- CLEAN.DATA$YFP.MAD.ABS/median(TDH3.SHAM$YFP.MEDIAN.ABS)

#Re-calculate mean and noise for each sample relative to controls
TDH3.SHAM <- subset(CLEAN.DATA, CLEAN.DATA$CONDITION == "TDH3.SHAM")
NEG <- subset(CLEAN.DATA, CLEAN.DATA$CONDITION == "NULL")
WT.SHAM <- subset(CLEAN.DATA, CLEAN.DATA$CONDITION == "SHAM")

#CALCULATE NOISE METRICS
CLEAN.DATA[,"YFP.CV.WT"]   <- CLEAN.DATA$YFP.SD.WT/CLEAN.DATA$YFP.MEDIAN.WT
CLEAN.DATA[,"YFP.CV.TDH3"]   <- CLEAN.DATA$YFP.SD.TDH3/CLEAN.DATA$YFP.MEDIAN.TDH3
CLEAN.DATA[,"YFP.FANO.WT"]   <- CLEAN.DATA$YFP.SD.WT^2/CLEAN.DATA$YFP.MEDIAN.WT
CLEAN.DATA[,"YFP.FANO.TDH3"]   <- CLEAN.DATA$YFP.SD.TDH3^2/CLEAN.DATA$YFP.MEDIAN.TDH3


#Re-calculate mean and noise for each sample relative to controls
TDH3.SHAM <- subset(CLEAN.DATA, CLEAN.DATA$CONDITION == "TDH3.SHAM")
NEG <- subset(CLEAN.DATA, CLEAN.DATA$CONDITION == "NULL")
WT.SHAM <- subset(CLEAN.DATA, CLEAN.DATA$CONDITION == "SHAM")

#CALCULATE RELATIVE NOISE METRICS
CLEAN.DATA[,"YFP.SD.WT.REL"]   <- CLEAN.DATA$YFP.SD.WT/median(WT.SHAM$YFP.SD.WT)
CLEAN.DATA[,"YFP.SD.TDH3.REL"]   <- CLEAN.DATA$YFP.SD.TDH3/median(WT.SHAM$YFP.SD.TDH3)
CLEAN.DATA[,"YFP.CV.WT.REL"]   <- CLEAN.DATA$YFP.SD.WT.REL/CLEAN.DATA$YFP.MEDIAN.WT
CLEAN.DATA[,"YFP.CV.TDH3.REL"]   <- CLEAN.DATA$YFP.SD.TDH3.REL/CLEAN.DATA$YFP.MEDIAN.TDH3
CLEAN.DATA[,"YFP.FANO.WT.REL"]   <- CLEAN.DATA$YFP.SD.WT.REL^2/CLEAN.DATA$YFP.MEDIAN.WT
CLEAN.DATA[,"YFP.FANO.TDH3.REL"]   <- CLEAN.DATA$YFP.SD.TDH3.REL^2/CLEAN.DATA$YFP.MEDIAN.TDH3


#Add NA for samples where CV or FANO are undefined.
for (i in 1:nrow(CLEAN.DATA))
{
  if (CLEAN.DATA[i,"YFP.MEDIAN.ABS"] < 0 | CLEAN.DATA[i,"YFP.SD.ABS"] < 0) {
    CLEAN.DATA[i,"YFP.MAD.WT"] <- NA
    CLEAN.DATA[i,"YFP.MAD.TDH3"] <- NA
    CLEAN.DATA[i,"YFP.CV.WT"] <- NA
    CLEAN.DATA[i,"YFP.CV.TDH3"] <- NA
    CLEAN.DATA[i,"YFP.CV.WT.REL"] <- NA
    CLEAN.DATA[i,"YFP.CV.TDH3.REL"] <- NA
    CLEAN.DATA[i,"YFP.FANO.WT"] <- NA
    CLEAN.DATA[i,"YFP.FANO.TDH3"] <- NA
    CLEAN.DATA[i,"YFP.FANO.WT.REL"] <- NA
    CLEAN.DATA[i,"YFP.FANO.TDH3.REL"] <- NA
    
  }
}

CLEAN.DATA <- droplevels(subset(CLEAN.DATA,CLEAN.DATA$CONDITION!="CTRL"))

write.table(CLEAN.DATA,"OST1.CLEAN.ADJUST_10112016_DCv34.csv",sep=",",quote=FALSE,row.names=FALSE)

CLEAN.DATA <- read.csv("OST1.CLEAN.ADJUST_10112016_DCv34.csv", header=TRUE)

OST1.MEAN <- aggregate(cbind(YFP.MEDIAN.ABS,YFP.SD.ABS,YFP.MAD.ABS,YFP.MEDIAN.WT,YFP.SD.WT,YFP.MAD.WT,YFP.MEDIAN.TDH3,YFP.SD.TDH3,YFP.MAD.TDH3,YFP.CV.WT,YFP.FANO.WT,YFP.CV.TDH3,YFP.FANO.TDH3,YFP.SD.WT.REL,YFP.CV.WT.REL,YFP.FANO.WT.REL,YFP.SD.TDH3.REL,YFP.CV.TDH3.REL,YFP.FANO.TDH3.REL,FSC.CORRECT) ~ STRAIN + CONDITION + ASSAY + PLATE + CTRL.GENO, data=CLEAN.DATA, FUN = mean)
OST1.SD <- aggregate(cbind(YFP.MEDIAN.ABS,YFP.SD.ABS,YFP.MAD.ABS,YFP.MEDIAN.WT,YFP.SD.WT,YFP.MAD.WT,YFP.MEDIAN.TDH3,YFP.SD.TDH3,YFP.MAD.TDH3,YFP.CV.WT,YFP.FANO.WT,YFP.CV.TDH3,YFP.FANO.TDH3,YFP.SD.WT.REL,YFP.CV.WT.REL,YFP.FANO.WT.REL,YFP.SD.TDH3.REL,YFP.CV.TDH3.REL,YFP.FANO.TDH3.REL,FSC.CORRECT) ~ STRAIN + CONDITION + ASSAY + PLATE + CTRL.GENO, data=CLEAN.DATA, FUN = sd)
OST1.N <- aggregate(cbind(YFP.MEDIAN.ABS,YFP.SD.ABS,YFP.MAD.ABS,YFP.MEDIAN.WT,YFP.SD.WT,YFP.MAD.WT,YFP.MEDIAN.TDH3,YFP.SD.TDH3,YFP.MAD.TDH3,YFP.CV.WT,YFP.FANO.WT,YFP.CV.TDH3,YFP.FANO.TDH3,YFP.SD.WT.REL,YFP.CV.WT.REL,YFP.FANO.WT.REL,YFP.SD.TDH3.REL,YFP.CV.TDH3.REL,YFP.FANO.TDH3.REL,FSC.CORRECT) ~ STRAIN + CONDITION + ASSAY + PLATE + CTRL.GENO, data=CLEAN.DATA, FUN = length)

OST1.MEAN  <- OST1.MEAN[which(OST1.N[,6] >= 3),]
OST1.SD <- OST1.SD[which(OST1.N[,6] >= 3),]
OST1.N <- OST1.N[which(OST1.N[,6] >= 3),]

OST1 <- cbind.data.frame(OST1.MEAN,OST1.SD[,6:25],OST1.N[,6])

colnames(OST1) <- c("STRAIN","CONDITION","ASSAY","PLATE","CTRL.GENO",paste0("MEAN.",colnames(OST1.MEAN[,6:25])),paste0("SD.",colnames(OST1.SD[,6:25])),"N")

write.table(OST1,"OST1.SUMMARY.DATA_10112016_DCv34.txt",sep="\t",quote=FALSE,row.names=FALSE)

`%notin%` <- function(x,y) !(x %in% y)

CONTAM.CHECK<-droplevels(subset(CLEAN.DATA,CLEAN.DATA$CONTAM.CHECK!="NONE"))

if(all(CONTAM.CHECK$STRAIN %notin% OST1$STRAIN)) {
  #All datapoints flagged as suspicious have been removed by outlier removal filters.
  print("Yay! No suspicious values detected. Proceed.")
}

if(any(CONTAM.CHECK$STRAIN %in% OST1$STRAIN)) {
  CONTAM.CHECK<-CONTAM.CHECK[CONTAM.CHECK$STRAIN %in% CONTAM.CHECK$STRAIN[which(CONTAM.CHECK$STRAIN %in% OST1$STRAIN)],]
  write.table(CONTAM.CHECK,"OST1.CONTAM.CHECK_outliersexcl.csv",sep=",",quote=FALSE,row.names=FALSE)  
}


###############################
#HIERARCHICAL PERMUTATION TEST#
###############################

#Clear memory
rm(list=ls())
options(warn=-1)

library(permute)
library(Hmisc)

N.PERM <- 10000

#Permutation function

SHUFFLE <- function(STRAIN.LIST = STRAIN.LIST[[i]],WT.LIST = WT.LIST) {
  SAMPLE <- c(STRAIN.LIST,WT.LIST[[sample(1:length(WT.LIST),1)]])
  OUTPUT <- sample(SAMPLE,length(SAMPLE))
  DIST.SIM <- abs(mean(OUTPUT[1:length(STRAIN.LIST)]) - mean(OUTPUT[(length(STRAIN.LIST)+1):length(SAMPLE)]))
  DIST.OBS <- abs(mean(SAMPLE[1:length(STRAIN.LIST)]) - mean(SAMPLE[(length(STRAIN.LIST)+1):length(SAMPLE)]))
  
  DIFF <- DIST.OBS - DIST.SIM
  
  return(DIFF)
}

parent.dir <- PATH
setwd(parent.dir)

####OST1 Analysis date 10112016#########################################################################

ALL.REP <- read.csv("OST1.CLEAN.ADJUST_10112016_DCv34.csv",header=TRUE)
ALL.REP <- subset(ALL.REP, MEDIAN.OUTLIER == "NO" & SD.OUTLIER == "NO" & FSC.OUTLIER == "NO")

SUMMARY <- read.table("OST1.SUMMARY.DATA_10112016_DCv34.txt",header=TRUE)

#SUMMARY <- droplevels(subset(SUMMARY, STRAIN != "1139"))

WT <- subset(ALL.REP, CONDITION == "SHAM")
LOW.FSC <- median(WT$FSC.CORRECT) - 4*mad(WT$FSC.CORRECT)
HIGH.FSC <- median(WT$FSC.CORRECT) + 4*mad(WT$FSC.CORRECT)
LOW.FLUO <- median(WT$YFP.MEDIAN.CORRECT) - 4*mad(WT$YFP.MEDIAN.CORRECT)
HIGH.FLUO <- median(WT$YFP.MEDIAN.CORRECT) + 4*mad(WT$YFP.MEDIAN.CORRECT)
WT <- subset(WT, FSC.CORRECT > LOW.FSC & FSC.CORRECT < HIGH.FSC & YFP.MEDIAN.CORRECT > LOW.FLUO & YFP.MEDIAN.CORRECT < HIGH.FLUO)

#MEDIAN

#Create list of WT phenotype grouped by strain

N.WT <- length(unique(WT$STRAIN))
WT.LIST <- vector('list',N.WT)
for (i in 1:N.WT)
{
  CUR.WT <- subset(WT, STRAIN == unique(WT$STRAIN)[i])
  WT.LIST[[i]] <- CUR.WT$YFP.MEDIAN.CORRECT 
}


#Create list of all strain phenotypes grouped by strain

SUMMARY <- SUMMARY[order(SUMMARY$STRAIN),]
ALL.REP <- droplevels(subset(ALL.REP,STRAIN %in% as.character(SUMMARY$STRAIN)))
ALL.REP <- ALL.REP[order(ALL.REP$STRAIN),]
N.STRAIN <- length(unique(ALL.REP$STRAIN))
STRAIN.LIST <- vector('list',N.STRAIN)
for (i in 1:N.STRAIN)
{
  CUR.STRAIN <- subset(ALL.REP, STRAIN == unique(ALL.REP$STRAIN)[i])
  STRAIN.LIST[[i]] <- CUR.STRAIN$YFP.MEDIAN.CORRECT
}


for (i in 1:length(STRAIN.LIST))
{
  DIFF <- sapply(1:N.PERM,FUN=function(x)	SHUFFLE(STRAIN.LIST = STRAIN.LIST[[i]],WT.LIST = WT.LIST))
  P.VAL <- (length(which(DIFF < 0)) + 1) / (length(which(DIFF < 0)) + length(which(DIFF > 0)) + 1) 	
  SUMMARY[i,"P.VAL.MEDIAN"] <- P.VAL
  print(round(i/length(STRAIN.LIST)*100,1))
}

#NOISE: CV

#Create list of WT phenotype grouped by strain

N.WT <- length(unique(WT$STRAIN))
WT.LIST <- vector('list',N.WT)
for (i in 1:N.WT)
{
  CUR.WT <- subset(WT, STRAIN == unique(WT$STRAIN)[i])
  WT.LIST[[i]] <- CUR.WT$YFP.CV.WT
}


#Create list of all strain phenotypes grouped by strain

SUMMARY <- SUMMARY[order(SUMMARY$STRAIN),]
ALL.REP <- droplevels(subset(ALL.REP,STRAIN %in% as.character(SUMMARY$STRAIN)))
ALL.REP <- ALL.REP[order(ALL.REP$STRAIN),]
N.STRAIN <- length(unique(ALL.REP$STRAIN))
STRAIN.LIST <- vector('list',N.STRAIN)
for (i in 1:N.STRAIN)
{
  CUR.STRAIN <- subset(ALL.REP, STRAIN == unique(ALL.REP$STRAIN)[i])
  STRAIN.LIST[[i]] <- CUR.STRAIN$YFP.CV.WT
}


for (i in 1:length(STRAIN.LIST))
{
  DIFF <- sapply(1:N.PERM,FUN=function(x)	SHUFFLE(STRAIN.LIST = STRAIN.LIST[[i]],WT.LIST = WT.LIST))
  P.VAL <- (length(which(DIFF < 0))+1) / (length(which(DIFF < 0)) + length(which(DIFF > 0)) + 1) 	
  SUMMARY[i,"P.VAL.CV.WT"] <- P.VAL
  print(round(i/length(STRAIN.LIST)*100,1))
}

#NOISE: FANO

#Create list of WT phenotype grouped by strain

N.WT <- length(unique(WT$STRAIN))
WT.LIST <- vector('list',N.WT)
for (i in 1:N.WT)
{
  CUR.WT <- subset(WT, STRAIN == unique(WT$STRAIN)[i])
  WT.LIST[[i]] <- CUR.WT$YFP.FANO.WT
}


#Create list of all strain phenotypes grouped by strain

SUMMARY <- SUMMARY[order(SUMMARY$STRAIN),]
ALL.REP <- droplevels(subset(ALL.REP,STRAIN %in% as.character(SUMMARY$STRAIN)))
ALL.REP <- ALL.REP[order(ALL.REP$STRAIN),]
N.STRAIN <- length(unique(ALL.REP$STRAIN))
STRAIN.LIST <- vector('list',N.STRAIN)
for (i in 1:N.STRAIN)
{
  CUR.STRAIN <- subset(ALL.REP, STRAIN == unique(ALL.REP$STRAIN)[i])
  STRAIN.LIST[[i]] <- CUR.STRAIN$YFP.FANO.WT
}


for (i in 1:length(STRAIN.LIST))
{
  DIFF <- sapply(1:N.PERM,FUN=function(x)	SHUFFLE(STRAIN.LIST = STRAIN.LIST[[i]],WT.LIST = WT.LIST))
  P.VAL <- (length(which(DIFF < 0))+1) / (length(which(DIFF < 0)) + length(which(DIFF > 0)) + 1) 	
  SUMMARY[i,"P.VAL.FANO.WT"] <- P.VAL
  print(round(i/length(STRAIN.LIST)*100,1))
}

#FSC MEDIAN

#Create list of WT phenotype grouped by strain

N.WT <- length(unique(WT$STRAIN))
WT.LIST <- vector('list',N.WT)
for (i in 1:N.WT)
{
  CUR.WT <- subset(WT, STRAIN == unique(WT$STRAIN)[i])
  WT.LIST[[i]] <- CUR.WT$FSC.CORRECT
}


#Create list of all strain phenotypes grouped by strain

SUMMARY <- SUMMARY[order(SUMMARY$STRAIN),]
ALL.REP <- droplevels(subset(ALL.REP,STRAIN %in% as.character(SUMMARY$STRAIN)))
ALL.REP <- ALL.REP[order(ALL.REP$STRAIN),]
N.STRAIN <- length(unique(ALL.REP$STRAIN))
STRAIN.LIST <- vector('list',N.STRAIN)
for (i in 1:N.STRAIN)
{
  CUR.STRAIN <- subset(ALL.REP, STRAIN == unique(ALL.REP$STRAIN)[i])
  STRAIN.LIST[[i]] <- CUR.STRAIN$FSC.CORRECT
}


for (i in 1:length(STRAIN.LIST))
{
  DIFF <- sapply(1:N.PERM,FUN=function(x)	SHUFFLE(STRAIN.LIST = STRAIN.LIST[[i]],WT.LIST = WT.LIST))
  P.VAL <- (length(which(DIFF < 0))+1) / (length(which(DIFF < 0)) + length(which(DIFF > 0)) + 1) 	
  SUMMARY[i,"P.VAL.FSC"] <- P.VAL
  print(round(i/length(STRAIN.LIST)*100,1))
}

write.table(SUMMARY,"OST1.SUMMARY.DCv34.Pvals.txt",sep="\t",quote=FALSE,row.names=FALSE)


###################
#    PFY1         #
###################


#Clear memory
rm(list=ls())
options(warn=-1)

#####################
#1-LOADING LIBRARIES#
#####################
library(flowCore)
library(flowClust)
library(flowViz)
library(pcaPP)
library(mixtools)
library(plyr)
library(MASS)
box <- graphics::box
options(warn=-1)

####################################
#2- Quality Control and Corrections#
####################################

parent.dir <- PATH
setwd(parent.dir)

DATA.TYPE <- c(
  "factor","factor","factor","factor","factor","factor","factor","factor","factor","factor","factor","factor","factor","factor","factor",
  "integer","integer","integer","integer",
  "numeric","numeric","numeric","numeric","numeric","numeric","numeric","numeric","numeric","numeric","numeric","numeric","numeric","numeric","numeric","numeric","numeric","numeric","numeric","numeric","numeric","numeric","numeric","numeric","numeric",
  "integer")

###Read in data sets
CLEAN.DATA <- read.table("PFY1_Clean.Data_10112016_DCv34.txt",colClasses=DATA.TYPE,header=TRUE, sep="\t")

###Remove samples with NA
CLEAN.DATA <- CLEAN.DATA[complete.cases(CLEAN.DATA),]

##Translate into log scale for appropriate correction
CLEAN.DATA[,"log.RNA.MEDIAN"] <- log((CLEAN.DATA[,"YFP.MEDIAN.FINAL"]) + 0.05)
CLEAN.DATA[,"log.RNA.MAD"] <- CLEAN.DATA[,"YFP.MAD.FINAL"] / (CLEAN.DATA[,"YFP.MEDIAN.FINAL"] + 0.05)
CLEAN.DATA[,"log.RNA.SD"] <- CLEAN.DATA[,"YFP.SD.FINAL"] / (CLEAN.DATA[,"YFP.MEDIAN.FINAL"] + 0.05)

###Remove samples with less than 1000 events
CLEAN.DATA <- subset(CLEAN.DATA,CLEAN.DATA$COUNTS.FINAL > 1000)

####Add variable to ID whether control is own promoter or TDH3
CLEAN.DATA[,"CTRL.GENO"]<-as.factor("PFY1")

#ID any suspicious phenotypes
CLEAN.DATA$CONTAM.CHECK<-"NONE"
CLEAN.DATA$CONTAM.CHECK[(CLEAN.DATA$CONDITION=="EMS"|CLEAN.DATA$CONDITION=="SHAM"|CLEAN.DATA$CONDITION=="EMS.LOW"|CLEAN.DATA$CONDITION=="EMS.HIGH") & (CLEAN.DATA$YFP.MEDIAN.FINAL<.56)]<-"BACT.CONTAM.CHECK"
CLEAN.DATA$CONTAM.CHECK[(CLEAN.DATA$CONDITION=="EMS"|CLEAN.DATA$CONDITION=="SHAM"|CLEAN.DATA$CONDITION=="EMS.LOW"|CLEAN.DATA$CONDITION=="EMS.HIGH") & (CLEAN.DATA$YFP.MEDIAN.FINAL>0.8)]<-"TDH3.CONTAM.CHECK"
CLEAN.DATA$CONTAM.CHECK[(CLEAN.DATA$CONDITION=="TDH3.SHAM" & CLEAN.DATA$YFP.MEDIAN.FINAL<0.8)]<-"EXP.SHAM.CONTAM.CHECK"
CLEAN.DATA$CONTAM.CHECK[(CLEAN.DATA$CONDITION=="NULL" & CLEAN.DATA$YFP.MEDIAN.FINAL>0.53)]<-"EXP.NULL.CONTAM.CHECK"

#Previously removed at this step:
#CLEAN.DATA<-droplevels(subset(CLEAN.DATA,CLEAN.DATA$STRAIN %nin% c("PFY1.P7.A5","PFY1.SHAM.P7.C11","PFY1.P7.E12","PFY1.P7.F7","PFY1.P7.H1")))
#CLEAN.DATA<-droplevels(subset(CLEAN.DATA,!(CLEAN.DATA$PLATE=="P7" & CLEAN.DATA$WELL=="G10")))

###Seperate controls and remove FSC outliers

CONTROL <- subset(CLEAN.DATA,CLEAN.DATA$CONDITION == "CTRL")
FSC.CORRECT <- lm(CONTROL$FSC.MEDIAN.FINAL ~ 0 + CONTROL$FLOW.RUN)
RESID <- FSC.CORRECT$resid
CLEAN.DATA <- droplevels(CLEAN.DATA[-which(CLEAN.DATA$CONDITION == "CTRL")[abs(RESID) > 3*mad(RESID)],])
CONTROL <- droplevels(subset(CLEAN.DATA,CLEAN.DATA$CONDITION == "CTRL"))

INITIAL.CORRECT <- lm(CONTROL$log.RNA.MEDIAN ~ 0 + CONTROL$FLOW.RUN)
RESID <- INITIAL.CORRECT$resid
CLEAN.DATA<- droplevels(CLEAN.DATA[-which(CLEAN.DATA$CONDITION == "CTRL")[abs(RESID) > 3*mad(RESID)],])
CONTROL <- droplevels(subset(CLEAN.DATA,CLEAN.DATA$CONDITION == "CTRL"))

`%notin%` <- function(x,y) !(x %in% y)
RESULT<-count(CONTROL$FLOW.RUN)
RESULT.MUT<-count(CLEAN.DATA$FLOW.RUN)
EXCLUDE1<-(RESULT.MUT[which(RESULT.MUT[,1] %notin% RESULT[,1]),1])
EXCLUDE2<-(RESULT[which(RESULT[,2] <5),1])
CLEAN.DATA<-droplevels(CLEAN.DATA[CLEAN.DATA$FLOW.RUN %notin% EXCLUDE1,])
CLEAN.DATA<-droplevels(CLEAN.DATA[CLEAN.DATA$FLOW.RUN %notin% EXCLUDE2,])
CONTROL <- subset(CLEAN.DATA,CLEAN.DATA$CONDITION == "CTRL")

###Corrections for YFP median and MAD
#Correct Median
#fit<- lm(log.RNA.MEDIAN~FLOW.RUN+REP+STACK.ORDER+STACK.ORDER:FLOW.RUN+REP+ROW+COLUMN+BLOCK+BLOCK:ROW+ROW:COLUMN+STACK.ORDER:BLOCK,data=CONTROL)
#step <- stepAIC(fit, direction="both")
log.RNA.MEDIAN.CORRECT <- lm(log.RNA.MEDIAN ~ FLOW.RUN + COLUMN + ROW + ROW:COLUMN, data=CONTROL)
CLEAN.DATA[,"log.RNA.MEDIAN.CORRECT"] <- CLEAN.DATA$log.RNA.MEDIAN - predict(log.RNA.MEDIAN.CORRECT, CLEAN.DATA) + mean(CONTROL$log.RNA.MEDIAN)
CONTROL <- subset(CLEAN.DATA,CLEAN.DATA$CONDITION == "CTRL")

#Correct sd - NOT HAPPY THAT WE SEE STACK.ORDER BLOCK AND REP COME UP SIG
#fit<- lm(log.RNA.SD~FLOW.RUN+REP+STACK.ORDER+STACK.ORDER:FLOW.RUN+REP+ROW+COLUMN+BLOCK+BLOCK:ROW+ROW:COLUMN+STACK.ORDER:BLOCK,data=CONTROL)
#step <- step(fit, direction="both")
log.RNA.SD.CORRECT <- lm(log.RNA.SD ~  FLOW.RUN + ROW + COLUMN,data=CONTROL)
CLEAN.DATA[,"log.RNA.SD.CORRECT"] <- CLEAN.DATA$log.RNA.SD - predict(log.RNA.SD.CORRECT , CLEAN.DATA) + mean(CONTROL$log.RNA.SD)
CONTROL <- subset(CLEAN.DATA,CLEAN.DATA$CONDITION == "CTRL")

#Correct mad
#fit<- lm(log.RNA.MAD~FLOW.RUN+REP+STACK.ORDER+STACK.ORDER:FLOW.RUN+REP+ROW+COLUMN+BLOCK+BLOCK:ROW+ROW:COLUMN+STACK.ORDER:BLOCK,data=CONTROL)
#step <- step(fit, direction="both")
log.RNA.MAD.CORRECT <- lm(log.RNA.MAD ~  FLOW.RUN + ROW + COLUMN,data=CONTROL)
CLEAN.DATA[,"log.RNA.MAD.CORRECT"] <- CLEAN.DATA$log.RNA.MAD - predict(log.RNA.MAD.CORRECT , CLEAN.DATA) + mean(CONTROL$log.RNA.MAD)
CONTROL <- subset(CLEAN.DATA,CLEAN.DATA$CONDITION == "CTRL")

#Correct FSC
#fit<- lm(FSC.MEDIAN.FINAL~FLOW.RUN+REP+STACK.ORDER+STACK.ORDER:FLOW.RUN+REP+ROW+COLUMN+BLOCK+BLOCK:ROW+ROW:COLUMN+STACK.ORDER:BLOCK,data=CONTROL)
#step <- stepAIC(fit, direction="both")
FSC.CORRECT <- lm(FSC.MEDIAN.FINAL ~  FLOW.RUN + ROW + COLUMN + ROW:COLUMN,data=CONTROL)
CLEAN.DATA[,"FSC.CORRECT"] <- CLEAN.DATA$FSC.MEDIAN.FINAL - predict(FSC.CORRECT, CLEAN.DATA) + mean(CONTROL$FSC.MEDIAN.FINAL)
CONTROL <- subset(CLEAN.DATA,CLEAN.DATA$CONDITION == "CTRL")


#Back translate to linear scale
CLEAN.DATA[,"YFP.MEDIAN.CORRECT"] <- (exp(CLEAN.DATA[,"log.RNA.MEDIAN.CORRECT"]) - 0.05)
CLEAN.DATA[,"YFP.SD.CORRECT"] <- CLEAN.DATA[,"log.RNA.SD.CORRECT"] * CLEAN.DATA[,"YFP.MEDIAN.CORRECT"]
CLEAN.DATA[,"YFP.MAD.CORRECT"] <- CLEAN.DATA[,"log.RNA.MAD.CORRECT"] * CLEAN.DATA[,"YFP.MEDIAN.CORRECT"]


#PERFORM FABIEN'S OUTLIER CORRECTION FILTERING OUT ANY REPLICATE OUTSIDE OF 4*MAD

#Remove outliers
for (i in 1:nrow(CLEAN.DATA))
{
  CUR <- subset(CLEAN.DATA, STRAIN == CLEAN.DATA[i,"STRAIN"])
  if (CLEAN.DATA[i,"YFP.MEDIAN.CORRECT"] > median(CUR$YFP.MEDIAN.CORRECT) + 4*mad(CUR$YFP.MEDIAN.CORRECT) | CLEAN.DATA[i,"YFP.MEDIAN.CORRECT"] < median(CUR$YFP.MEDIAN.CORRECT) - 4*mad(CUR$YFP.MEDIAN.CORRECT)) {
    CLEAN.DATA[i,"MEDIAN.OUTLIER"] <- "YES"
  } else {
    CLEAN.DATA[i,"MEDIAN.OUTLIER"] <- "NO"
  }
  if (CLEAN.DATA[i,"YFP.SD.CORRECT"] > median(CUR$YFP.SD.CORRECT) + 4*mad(CUR$YFP.SD.CORRECT) | CLEAN.DATA[i,"YFP.SD.CORRECT"] < median(CUR$YFP.SD.CORRECT) - 4*mad(CUR$YFP.SD.CORRECT)) {
    CLEAN.DATA[i,"SD.OUTLIER"] <- "YES"
  } else {
    CLEAN.DATA[i,"SD.OUTLIER"] <- "NO"
  }
  if (CLEAN.DATA[i,"FSC.CORRECT"] > median(CUR$FSC.CORRECT) + 4*mad(CUR$FSC.CORRECT) | CLEAN.DATA[i,"FSC.CORRECT"] < median(CUR$FSC.CORRECT) - 4*mad(CUR$FSC.CORRECT)) {
    CLEAN.DATA[i,"FSC.OUTLIER"] <- "YES"
  } else {
    CLEAN.DATA[i,"FSC.OUTLIER"] <- "NO"
  }	
}

CLEAN.DATA <- subset(CLEAN.DATA, MEDIAN.OUTLIER == "NO" & SD.OUTLIER == "NO" & FSC.OUTLIER == "NO")

#Write processed data to file
write.table(CLEAN.DATA,"PFY1.CLEAN.CORRECT_10112016_DCv34.csv",sep=",",quote=FALSE,row.names=FALSE)

#####################
#5 - Control plots  #
#####################

CONTROL <- droplevels(subset(CLEAN.DATA,CLEAN.DATA$CONDITION == "CTRL"))
SHAM <- droplevels(subset(CLEAN.DATA,CLEAN.DATA$CONDITION == "SHAM"))
TDH3.SHAM <- droplevels(subset(CLEAN.DATA,CLEAN.DATA$CONDITION == "TDH3.SHAM"))


###############################################################################
#
#                     Strain Estimates
#
###############################################################################

# #Clear memory
rm(list=ls())
options(warn=-1)

setwd(PATH)

#PFY1
CLEAN.DATA <- read.csv("PFY1.CLEAN.CORRECT_10112016_DCv34.csv", header=TRUE)

#Calculate mean and noise for each sample relative to controls
TDH3.SHAM <- subset(CLEAN.DATA, CLEAN.DATA$CONDITION == "TDH3.SHAM")
NEG <- subset(CLEAN.DATA, CLEAN.DATA$CONDITION == "NULL")
NEG<-subset(NEG,NEG$log.RNA.MEDIAN< -0.5)
WT.SHAM <- subset(CLEAN.DATA, CLEAN.DATA$CONDITION == "SHAM")

#ABSOLUTE EXPRESSION SCALE
CLEAN.DATA[,"YFP.MEDIAN.ABS"] <- (CLEAN.DATA$YFP.MEDIAN.CORRECT - median(NEG$YFP.MEDIAN.CORRECT))
CLEAN.DATA[,"YFP.SD.ABS"]   <- CLEAN.DATA$YFP.SD.CORRECT - median(NEG$YFP.SD.CORRECT)
CLEAN.DATA[,"YFP.MAD.ABS"]   <- CLEAN.DATA$YFP.MAD.CORRECT - median(NEG$YFP.SD.CORRECT)

#Re-calculate mean and noise for each sample relative to controls
TDH3.SHAM <- subset(CLEAN.DATA, CLEAN.DATA$CONDITION == "TDH3.SHAM")
NEG <- subset(CLEAN.DATA, CLEAN.DATA$CONDITION == "NULL")
NEG<-subset(NEG,NEG$log.RNA.MEDIAN< -0.5)
WT.SHAM <- subset(CLEAN.DATA, CLEAN.DATA$CONDITION == "SHAM")

#PROMOTER-SPECIFIC SCALING
CLEAN.DATA[,"YFP.MEDIAN.WT"] <- CLEAN.DATA$YFP.MEDIAN.ABS/median(WT.SHAM$YFP.MEDIAN.ABS)
CLEAN.DATA[,"YFP.SD.WT"]   <- CLEAN.DATA$YFP.SD.ABS/median(WT.SHAM$YFP.MEDIAN.ABS)
CLEAN.DATA[,"YFP.MAD.WT"]   <- CLEAN.DATA$YFP.MAD.ABS/median(WT.SHAM$YFP.MEDIAN.ABS)

#RELATIVE TO TDH3 EXPRESSION
CLEAN.DATA[,"YFP.MEDIAN.TDH3"] <- CLEAN.DATA$YFP.MEDIAN.ABS/median(TDH3.SHAM$YFP.MEDIAN.ABS)
CLEAN.DATA[,"YFP.SD.TDH3"]   <- CLEAN.DATA$YFP.SD.ABS/median(TDH3.SHAM$YFP.MEDIAN.ABS)
CLEAN.DATA[,"YFP.MAD.TDH3"]   <- CLEAN.DATA$YFP.MAD.ABS/median(TDH3.SHAM$YFP.MEDIAN.ABS)

#Re-calculate mean and noise for each sample relative to controls
TDH3.SHAM <- subset(CLEAN.DATA, CLEAN.DATA$CONDITION == "TDH3.SHAM")
NEG <- subset(CLEAN.DATA, CLEAN.DATA$CONDITION == "NULL")
WT.SHAM <- subset(CLEAN.DATA, CLEAN.DATA$CONDITION == "SHAM")

#CALCULATE NOISE METRICS
CLEAN.DATA[,"YFP.CV.WT"]   <- CLEAN.DATA$YFP.SD.WT/CLEAN.DATA$YFP.MEDIAN.WT
CLEAN.DATA[,"YFP.CV.TDH3"]   <- CLEAN.DATA$YFP.SD.TDH3/CLEAN.DATA$YFP.MEDIAN.TDH3
CLEAN.DATA[,"YFP.FANO.WT"]   <- CLEAN.DATA$YFP.SD.WT^2/CLEAN.DATA$YFP.MEDIAN.WT
CLEAN.DATA[,"YFP.FANO.TDH3"]   <- CLEAN.DATA$YFP.SD.TDH3^2/CLEAN.DATA$YFP.MEDIAN.TDH3


#Re-calculate mean and noise for each sample relative to controls
TDH3.SHAM <- subset(CLEAN.DATA, CLEAN.DATA$CONDITION == "TDH3.SHAM")
NEG <- subset(CLEAN.DATA, CLEAN.DATA$CONDITION == "NULL")
WT.SHAM <- subset(CLEAN.DATA, CLEAN.DATA$CONDITION == "SHAM")

#CALCULATE RELATIVE NOISE METRICS
CLEAN.DATA[,"YFP.SD.WT.REL"]   <- CLEAN.DATA$YFP.SD.WT/median(WT.SHAM$YFP.SD.WT)
CLEAN.DATA[,"YFP.SD.TDH3.REL"]   <- CLEAN.DATA$YFP.SD.TDH3/median(WT.SHAM$YFP.SD.TDH3)
CLEAN.DATA[,"YFP.CV.WT.REL"]   <- CLEAN.DATA$YFP.SD.WT.REL/CLEAN.DATA$YFP.MEDIAN.WT
CLEAN.DATA[,"YFP.CV.TDH3.REL"]   <- CLEAN.DATA$YFP.SD.TDH3.REL/CLEAN.DATA$YFP.MEDIAN.TDH3
CLEAN.DATA[,"YFP.FANO.WT.REL"]   <- CLEAN.DATA$YFP.SD.WT.REL^2/CLEAN.DATA$YFP.MEDIAN.WT
CLEAN.DATA[,"YFP.FANO.TDH3.REL"]   <- CLEAN.DATA$YFP.SD.TDH3.REL^2/CLEAN.DATA$YFP.MEDIAN.TDH3


#Add NA for samples where CV or FANO are undefined.
for (i in 1:nrow(CLEAN.DATA))
{
  if (CLEAN.DATA[i,"YFP.MEDIAN.ABS"] < 0 | CLEAN.DATA[i,"YFP.SD.ABS"] < 0) {
    CLEAN.DATA[i,"YFP.MAD.WT"] <- NA
    CLEAN.DATA[i,"YFP.MAD.TDH3"] <- NA
    CLEAN.DATA[i,"YFP.CV.WT"] <- NA
    CLEAN.DATA[i,"YFP.CV.TDH3"] <- NA
    CLEAN.DATA[i,"YFP.CV.WT.REL"] <- NA
    CLEAN.DATA[i,"YFP.CV.TDH3.REL"] <- NA
    CLEAN.DATA[i,"YFP.FANO.WT"] <- NA
    CLEAN.DATA[i,"YFP.FANO.TDH3"] <- NA
    CLEAN.DATA[i,"YFP.FANO.WT.REL"] <- NA
    CLEAN.DATA[i,"YFP.FANO.TDH3.REL"] <- NA
    
  }
}

CLEAN.DATA <- droplevels(subset(CLEAN.DATA,CLEAN.DATA$CONDITION!="CTRL"))

write.table(CLEAN.DATA,"PFY1.CLEAN.ADJUST_10112016_DCv34.csv",sep=",",quote=FALSE,row.names=FALSE)

CLEAN.DATA <- read.csv("PFY1.CLEAN.ADJUST_10112016_DCv34.csv", header=TRUE)

PFY1.MEAN <- aggregate(cbind(YFP.MEDIAN.ABS,YFP.SD.ABS,YFP.MAD.ABS,YFP.MEDIAN.WT,YFP.SD.WT,YFP.MAD.WT,YFP.MEDIAN.TDH3,YFP.SD.TDH3,YFP.MAD.TDH3,YFP.CV.WT,YFP.FANO.WT,YFP.CV.TDH3,YFP.FANO.TDH3,YFP.SD.WT.REL,YFP.CV.WT.REL,YFP.FANO.WT.REL,YFP.SD.TDH3.REL,YFP.CV.TDH3.REL,YFP.FANO.TDH3.REL,FSC.CORRECT) ~ STRAIN + CONDITION + ASSAY + PLATE + CTRL.GENO, data=CLEAN.DATA, FUN = mean)
PFY1.SD <- aggregate(cbind(YFP.MEDIAN.ABS,YFP.SD.ABS,YFP.MAD.ABS,YFP.MEDIAN.WT,YFP.SD.WT,YFP.MAD.WT,YFP.MEDIAN.TDH3,YFP.SD.TDH3,YFP.MAD.TDH3,YFP.CV.WT,YFP.FANO.WT,YFP.CV.TDH3,YFP.FANO.TDH3,YFP.SD.WT.REL,YFP.CV.WT.REL,YFP.FANO.WT.REL,YFP.SD.TDH3.REL,YFP.CV.TDH3.REL,YFP.FANO.TDH3.REL,FSC.CORRECT) ~ STRAIN + CONDITION + ASSAY + PLATE + CTRL.GENO, data=CLEAN.DATA, FUN = sd)
PFY1.N <- aggregate(cbind(YFP.MEDIAN.ABS,YFP.SD.ABS,YFP.MAD.ABS,YFP.MEDIAN.WT,YFP.SD.WT,YFP.MAD.WT,YFP.MEDIAN.TDH3,YFP.SD.TDH3,YFP.MAD.TDH3,YFP.CV.WT,YFP.FANO.WT,YFP.CV.TDH3,YFP.FANO.TDH3,YFP.SD.WT.REL,YFP.CV.WT.REL,YFP.FANO.WT.REL,YFP.SD.TDH3.REL,YFP.CV.TDH3.REL,YFP.FANO.TDH3.REL,FSC.CORRECT) ~ STRAIN + CONDITION + ASSAY + PLATE + CTRL.GENO, data=CLEAN.DATA, FUN = length)

PFY1.MEAN  <- PFY1.MEAN[which(PFY1.N[,6] >= 3),]
PFY1.SD <- PFY1.SD[which(PFY1.N[,6] >= 3),]
PFY1.N <- PFY1.N[which(PFY1.N[,6] >= 3),]

PFY1 <- cbind.data.frame(PFY1.MEAN,PFY1.SD[,6:25],PFY1.N[,6])

colnames(PFY1) <- c("STRAIN","CONDITION","ASSAY","PLATE","CTRL.GENO",paste0("MEAN.",colnames(PFY1.MEAN[,6:25])),paste0("SD.",colnames(PFY1.SD[,6:25])),"N")

write.table(PFY1,"PFY1.SUMMARY.DATA_10112016_DCv34.txt",sep="\t",quote=FALSE,row.names=FALSE)

`%notin%` <- function(x,y) !(x %in% y)

CONTAM.CHECK<-droplevels(subset(CLEAN.DATA,CLEAN.DATA$CONTAM.CHECK!="NONE"))

if(all(CONTAM.CHECK$STRAIN %notin% PFY1$STRAIN)) {
  #All datapoints flagged as suspicious have been removed by outlier removal filters.
  print("Yay! No suspicious values detected. Proceed.")
}

if(any(CONTAM.CHECK$STRAIN %in% PFY1$STRAIN)) {
  CONTAM.CHECK<-CONTAM.CHECK[CONTAM.CHECK$STRAIN %in% CONTAM.CHECK$STRAIN[which(CONTAM.CHECK$STRAIN %in% PFY1$STRAIN)],]
  write.table(CONTAM.CHECK,"PFY1.CONTAM.CHECK_outliersexcl.csv",sep=",",quote=FALSE,row.names=FALSE)  
}


###############################
#HIERARCHICAL PERMUTATION TEST#
###############################

#Clear memory
rm(list=ls())
options(warn=-1)

library(permute)
library(Hmisc)

N.PERM <- 10000

#Permutation function

SHUFFLE <- function(STRAIN.LIST = STRAIN.LIST[[i]],WT.LIST = WT.LIST) {
  SAMPLE <- c(STRAIN.LIST,WT.LIST[[sample(1:length(WT.LIST),1)]])
  OUTPUT <- sample(SAMPLE,length(SAMPLE))
  DIST.SIM <- abs(mean(OUTPUT[1:length(STRAIN.LIST)]) - mean(OUTPUT[(length(STRAIN.LIST)+1):length(SAMPLE)]))
  DIST.OBS <- abs(mean(SAMPLE[1:length(STRAIN.LIST)]) - mean(SAMPLE[(length(STRAIN.LIST)+1):length(SAMPLE)]))
  
  DIFF <- DIST.OBS - DIST.SIM
  
  return(DIFF)
}

parent.dir <- PATH
setwd(parent.dir)

####PFY1 Analysis date 10112016#########################################################################

ALL.REP <- read.csv("PFY1.CLEAN.ADJUST_10112016_DCv34.csv",header=TRUE)
ALL.REP <- subset(ALL.REP, MEDIAN.OUTLIER == "NO" & SD.OUTLIER == "NO" & FSC.OUTLIER == "NO")

SUMMARY <- read.table("PFY1.SUMMARY.DATA_10112016_DCv34.txt",header=TRUE)

#SUMMARY <- droplevels(subset(SUMMARY, STRAIN != "1139"))

WT <- subset(ALL.REP, CONDITION == "SHAM")
LOW.FSC <- median(WT$FSC.CORRECT) - 4*mad(WT$FSC.CORRECT)
HIGH.FSC <- median(WT$FSC.CORRECT) + 4*mad(WT$FSC.CORRECT)
LOW.FLUO <- median(WT$YFP.MEDIAN.CORRECT) - 4*mad(WT$YFP.MEDIAN.CORRECT)
HIGH.FLUO <- median(WT$YFP.MEDIAN.CORRECT) + 4*mad(WT$YFP.MEDIAN.CORRECT)
WT <- subset(WT, FSC.CORRECT > LOW.FSC & FSC.CORRECT < HIGH.FSC & YFP.MEDIAN.CORRECT > LOW.FLUO & YFP.MEDIAN.CORRECT < HIGH.FLUO)

#MEDIAN

#Create list of WT phenotype grouped by strain

N.WT <- length(unique(WT$STRAIN))
WT.LIST <- vector('list',N.WT)
for (i in 1:N.WT)
{
  CUR.WT <- subset(WT, STRAIN == unique(WT$STRAIN)[i])
  WT.LIST[[i]] <- CUR.WT$YFP.MEDIAN.CORRECT 
}


#Create list of all strain phenotypes grouped by strain

SUMMARY <- SUMMARY[order(SUMMARY$STRAIN),]
ALL.REP <- droplevels(subset(ALL.REP,STRAIN %in% as.character(SUMMARY$STRAIN)))
ALL.REP <- ALL.REP[order(ALL.REP$STRAIN),]
N.STRAIN <- length(unique(ALL.REP$STRAIN))
STRAIN.LIST <- vector('list',N.STRAIN)
for (i in 1:N.STRAIN)
{
  CUR.STRAIN <- subset(ALL.REP, STRAIN == unique(ALL.REP$STRAIN)[i])
  STRAIN.LIST[[i]] <- CUR.STRAIN$YFP.MEDIAN.CORRECT
}


for (i in 1:length(STRAIN.LIST))
{
  DIFF <- sapply(1:N.PERM,FUN=function(x)	SHUFFLE(STRAIN.LIST = STRAIN.LIST[[i]],WT.LIST = WT.LIST))
  P.VAL <- (length(which(DIFF < 0)) + 1) / (length(which(DIFF < 0)) + length(which(DIFF > 0)) + 1) 	
  SUMMARY[i,"P.VAL.MEDIAN"] <- P.VAL
  print(round(i/length(STRAIN.LIST)*100,1))
}

#NOISE: CV

#Create list of WT phenotype grouped by strain

N.WT <- length(unique(WT$STRAIN))
WT.LIST <- vector('list',N.WT)
for (i in 1:N.WT)
{
  CUR.WT <- subset(WT, STRAIN == unique(WT$STRAIN)[i])
  WT.LIST[[i]] <- CUR.WT$YFP.CV.WT
}


#Create list of all strain phenotypes grouped by strain

SUMMARY <- SUMMARY[order(SUMMARY$STRAIN),]
ALL.REP <- droplevels(subset(ALL.REP,STRAIN %in% as.character(SUMMARY$STRAIN)))
ALL.REP <- ALL.REP[order(ALL.REP$STRAIN),]
N.STRAIN <- length(unique(ALL.REP$STRAIN))
STRAIN.LIST <- vector('list',N.STRAIN)
for (i in 1:N.STRAIN)
{
  CUR.STRAIN <- subset(ALL.REP, STRAIN == unique(ALL.REP$STRAIN)[i])
  STRAIN.LIST[[i]] <- CUR.STRAIN$YFP.CV.WT
}


for (i in 1:length(STRAIN.LIST))
{
  DIFF <- sapply(1:N.PERM,FUN=function(x)	SHUFFLE(STRAIN.LIST = STRAIN.LIST[[i]],WT.LIST = WT.LIST))
  P.VAL <- (length(which(DIFF < 0))+1) / (length(which(DIFF < 0)) + length(which(DIFF > 0)) + 1) 	
  SUMMARY[i,"P.VAL.CV.WT"] <- P.VAL
  print(round(i/length(STRAIN.LIST)*100,1))
}

#NOISE: FANO

#Create list of WT phenotype grouped by strain

N.WT <- length(unique(WT$STRAIN))
WT.LIST <- vector('list',N.WT)
for (i in 1:N.WT)
{
  CUR.WT <- subset(WT, STRAIN == unique(WT$STRAIN)[i])
  WT.LIST[[i]] <- CUR.WT$YFP.FANO.WT
}


#Create list of all strain phenotypes grouped by strain

SUMMARY <- SUMMARY[order(SUMMARY$STRAIN),]
ALL.REP <- droplevels(subset(ALL.REP,STRAIN %in% as.character(SUMMARY$STRAIN)))
ALL.REP <- ALL.REP[order(ALL.REP$STRAIN),]
N.STRAIN <- length(unique(ALL.REP$STRAIN))
STRAIN.LIST <- vector('list',N.STRAIN)
for (i in 1:N.STRAIN)
{
  CUR.STRAIN <- subset(ALL.REP, STRAIN == unique(ALL.REP$STRAIN)[i])
  STRAIN.LIST[[i]] <- CUR.STRAIN$YFP.FANO.WT
}


for (i in 1:length(STRAIN.LIST))
{
  DIFF <- sapply(1:N.PERM,FUN=function(x)	SHUFFLE(STRAIN.LIST = STRAIN.LIST[[i]],WT.LIST = WT.LIST))
  P.VAL <- (length(which(DIFF < 0))+1) / (length(which(DIFF < 0)) + length(which(DIFF > 0)) + 1) 	
  SUMMARY[i,"P.VAL.FANO.WT"] <- P.VAL
  print(round(i/length(STRAIN.LIST)*100,1))
}

#FSC MEDIAN

#Create list of WT phenotype grouped by strain

N.WT <- length(unique(WT$STRAIN))
WT.LIST <- vector('list',N.WT)
for (i in 1:N.WT)
{
  CUR.WT <- subset(WT, STRAIN == unique(WT$STRAIN)[i])
  WT.LIST[[i]] <- CUR.WT$FSC.CORRECT
}


#Create list of all strain phenotypes grouped by strain

SUMMARY <- SUMMARY[order(SUMMARY$STRAIN),]
ALL.REP <- droplevels(subset(ALL.REP,STRAIN %in% as.character(SUMMARY$STRAIN)))
ALL.REP <- ALL.REP[order(ALL.REP$STRAIN),]
N.STRAIN <- length(unique(ALL.REP$STRAIN))
STRAIN.LIST <- vector('list',N.STRAIN)
for (i in 1:N.STRAIN)
{
  CUR.STRAIN <- subset(ALL.REP, STRAIN == unique(ALL.REP$STRAIN)[i])
  STRAIN.LIST[[i]] <- CUR.STRAIN$FSC.CORRECT
}


for (i in 1:length(STRAIN.LIST))
{
  DIFF <- sapply(1:N.PERM,FUN=function(x)	SHUFFLE(STRAIN.LIST = STRAIN.LIST[[i]],WT.LIST = WT.LIST))
  P.VAL <- (length(which(DIFF < 0))+1) / (length(which(DIFF < 0)) + length(which(DIFF > 0)) + 1) 	
  SUMMARY[i,"P.VAL.FSC"] <- P.VAL
  print(round(i/length(STRAIN.LIST)*100,1))
}

write.table(SUMMARY,"PFY1.SUMMARY.10112016.DCv34.Pvals.txt",sep="\t",quote=FALSE,row.names=FALSE)

#________________________________________________________________________________________________________________________

#######################
#         RNR1
#######################


########################################
#  CLEANING
########################################

#RNR1

#Clear memory
rm(list=ls())
options(warn=-1)

#Set working directory
parent.dir <- PATH
setwd(parent.dir)

#Load experiment setup
SETUP <- read.csv("E:/wittkopp data/raw data/RNR1/Template_RNR1_V2_10102016.csv",header=TRUE,as.is=TRUE)
TESTNAMES <- list.files("E:/wittkopp data/raw data/RNR1/Round2",pattern=".fcs",full.name=TRUE,recursive=TRUE,include.dirs=TRUE)
SETUP<-subset(SETUP, SETUP$FILENAMES %in% TESTNAMES, drop = TRUE)

#For troubleshooting
#SETUP<-SETUP[1:100,]


SETUP[,"COUNTER"] <- c(1:nrow(SETUP))

#Determine Hard Gates
GATES <- GATE.CALIB(SETUP$FILENAMES[1])

#Clean Data
Output <- apply(SETUP,1,CLEANING,GATES=GATES)
OUTPUT <- as.data.frame(Output[[1]])
OUTPUT[2:nrow(SETUP),1:ncol(OUTPUT)] <- NA
for (i in 2:nrow(SETUP))
{
  if (is.null(Output[[i]]))
  {} 
  else {
    OUTPUT[i,] <- Output[[i]]
  }
}


write.table(OUTPUT,"RNR1_Experiment.Output_DCv34.txt",row.names=FALSE,sep="\t")

CLEAN <- cbind.data.frame(SETUP[,1:(ncol(SETUP)-2)],OUTPUT)

write.table(CLEAN,"RNR1_Clean.Data_10112016_DCv34.txt",row.names=FALSE,sep="\t")

################################################################
#     PLATE CORRECTION & OUTLIER REMOVAL
################################################################

#Clear memory
rm(list=ls())
options(warn=-1)

#####################
#1-LOADING LIBRARIES#
#####################
library(flowCore)
library(flowClust)
library(flowViz)
library(pcaPP)
library(mixtools)
library(plyr)
library(MASS)
box <- graphics::box
options(warn=-1)

####################################
#2- Quality Control and Corrections#
####################################

parent.dir <- PATH
setwd(parent.dir)

DATA.TYPE <- c(
  "factor","factor","factor","factor","factor","factor","factor","factor","factor","factor","factor","factor","factor","factor","factor",
  "integer","integer","integer","integer",
  "numeric","numeric","numeric","numeric","numeric","numeric","numeric","numeric","numeric","numeric","numeric","numeric","numeric","numeric","numeric","numeric","numeric","numeric","numeric","numeric","numeric","numeric","numeric","numeric","numeric",
  "integer")

###Read in data sets
CLEAN.DATA <- read.table("RNR1_Clean.Data_10112016_DCv34.txt",colClasses=DATA.TYPE,header=TRUE, sep="\t")

###Remove samples with NA
CLEAN.DATA <- CLEAN.DATA[complete.cases(CLEAN.DATA),]

##Translate into log scale for appropriate correction
CLEAN.DATA[,"log.RNA.MEDIAN"] <- log((CLEAN.DATA[,"YFP.MEDIAN.FINAL"]) + 0.05)
CLEAN.DATA[,"log.RNA.MAD"] <- CLEAN.DATA[,"YFP.MAD.FINAL"] / (CLEAN.DATA[,"YFP.MEDIAN.FINAL"] + 0.05)
CLEAN.DATA[,"log.RNA.SD"] <- CLEAN.DATA[,"YFP.SD.FINAL"] / (CLEAN.DATA[,"YFP.MEDIAN.FINAL"] + 0.05)

###Remove samples with less than 1000 events
CLEAN.DATA <- subset(CLEAN.DATA,CLEAN.DATA$COUNTS.FINAL > 1000)

####Add variable to ID whether control is own promoter or TDH3
for (i in 1:nrow(CLEAN.DATA))
{
  CUR <- subset(CLEAN.DATA, CLEAN.DATA$ASSAY == CLEAN.DATA[i,"ASSAY"] & CLEAN.DATA$FLOW.RUN == CLEAN.DATA[i,"FLOW.RUN"])
  CUR.TDH3.SHAM<-subset(CUR, CONDITION == "TDH3.SHAM")
  CUR.CTRL<-subset(CUR, CONDITION == "CTRL")
  CUR.SHAM<-subset(CUR, CONDITION == "SHAM")
  
  if (nrow(CUR.TDH3.SHAM) == 0 | nrow(CUR.SHAM) == 0 | nrow(CUR.CTRL) == 0 ) {
    CLEAN.DATA[i,"CTRL.GENO"] <- "exclude"
  } else{
    TDH3.DIFF<-abs(median(CUR.TDH3.SHAM$log.RNA.MEDIAN)-median(CUR.CTRL$log.RNA.MEDIAN))
    OWN.DIFF<-abs(median(CUR.SHAM$log.RNA.MEDIAN)-median(CUR.CTRL$log.RNA.MEDIAN))
    
    if (TDH3.DIFF < OWN.DIFF) {
      CLEAN.DATA[i,"CTRL.GENO"] <- "TDH3"
    } else {
      CLEAN.DATA[i,"CTRL.GENO"] <- as.character(CLEAN.DATA[i,"ASSAY"])
    }
  }
}

CLEAN.DATA[,"CTRL.GENO"]<-as.factor(CLEAN.DATA[,"CTRL.GENO"])


###Select desired data / Drop known bad samples

CLEAN.DATA$CONTAM.CHECK<-"NONE"
CLEAN.DATA$CONTAM.CHECK[(CLEAN.DATA$CONDITION=="EMS"|CLEAN.DATA$CONDITION=="SHAM"|CLEAN.DATA$CONDITION=="EMS.LOW"|CLEAN.DATA$CONDITION=="EMS.HIGH") & (CLEAN.DATA$YFP.MEDIAN.FINAL<.56)]<-"BACT.CONTAM.CHECK"
CLEAN.DATA$CONTAM.CHECK[(CLEAN.DATA$CONDITION=="EMS"|CLEAN.DATA$CONDITION=="SHAM"|CLEAN.DATA$CONDITION=="EMS.LOW"|CLEAN.DATA$CONDITION=="EMS.HIGH") & (CLEAN.DATA$YFP.MEDIAN.FINAL>0.8)]<-"TDH3.CONTAM.CHECK"
CLEAN.DATA$CONTAM.CHECK[(CLEAN.DATA$CONDITION=="TDH3.SHAM" & CLEAN.DATA$YFP.MEDIAN.FINAL<0.8)]<-"EXP.SHAM.CONTAM.CHECK"
CLEAN.DATA$CONTAM.CHECK[(CLEAN.DATA$CONDITION=="NULL" & CLEAN.DATA$YFP.MEDIAN.FINAL>0.56)]<-"EXP.NULL.CONTAM.CHECK"


CLEAN.DATA.OWN.CTRL<-subset(CLEAN.DATA,CLEAN.DATA$FLOW.RUN %in% c(1:64))
CLEAN.DATA.TDH3.CTRL<-subset(CLEAN.DATA,CLEAN.DATA$FLOW.RUN %in% c(65:80))

###Seperate controls and remove FSC outliers
TDH3.CONTROL <- subset(CLEAN.DATA.TDH3.CTRL,CLEAN.DATA.TDH3.CTRL$CONDITION == "CTRL")
FSC.CORRECT <- lm(TDH3.CONTROL$FSC.MEDIAN.FINAL ~ 0 + TDH3.CONTROL$FLOW.RUN)
RESID <- FSC.CORRECT$resid
CLEAN.DATA.TDH3.CTRL <- droplevels(CLEAN.DATA.TDH3.CTRL[-which(CLEAN.DATA.TDH3.CTRL$CONDITION == "CTRL")[abs(RESID) > 3*mad(RESID)],])
TDH3.CONTROL <- droplevels(subset(CLEAN.DATA.TDH3.CTRL,CLEAN.DATA.TDH3.CTRL$CONDITION == "CTRL"))

INITIAL.CORRECT <- lm(TDH3.CONTROL$log.RNA.MEDIAN ~ 0 + TDH3.CONTROL$FLOW.RUN)
RESID <- INITIAL.CORRECT$resid
CLEAN.DATA.TDH3.CTRL<- droplevels(CLEAN.DATA.TDH3.CTRL[-which(CLEAN.DATA.TDH3.CTRL$CONDITION == "CTRL")[abs(RESID) > 3*mad(RESID)],])
TDH3.CONTROL <- droplevels(subset(CLEAN.DATA.TDH3.CTRL,CLEAN.DATA.TDH3.CTRL$CONDITION == "CTRL"))

`%notin%` <- function(x,y) !(x %in% y)
RESULT<-count(TDH3.CONTROL$FLOW.RUN)
RESULT.MUT<-count(CLEAN.DATA.TDH3.CTRL$FLOW.RUN)
EXCLUDE1<-(RESULT.MUT[which(RESULT.MUT[,1] %notin% RESULT[,1]),1])
EXCLUDE2<-(RESULT[which(RESULT[,2] <5),1])
CLEAN.DATA.TDH3.CTRL<-droplevels(CLEAN.DATA.TDH3.CTRL[CLEAN.DATA.TDH3.CTRL$FLOW.RUN %notin% EXCLUDE1,])
CLEAN.DATA.TDH3.CTRL<-droplevels(CLEAN.DATA.TDH3.CTRL[CLEAN.DATA.TDH3.CTRL$FLOW.RUN %notin% EXCLUDE2,])
TDH3.CONTROL <- subset(CLEAN.DATA.TDH3.CTRL,CLEAN.DATA.TDH3.CTRL$CONDITION == "CTRL")


###Corrections for YFP median and MAD
#Correct Median
#fit<- lm(log.RNA.MEDIAN~FLOW.RUN+REP+STACK.ORDER+STACK.ORDER:FLOW.RUN+REP+ROW+COLUMN+BLOCK+BLOCK:ROW+ROW:COLUMN+STACK.ORDER:BLOCK,data=TDH3.CONTROL)
#step <- stepAIC(fit, direction="both")
log.RNA.MEDIAN.CORRECT <- lm(log.RNA.MEDIAN ~ FLOW.RUN + COLUMN + ROW, data=TDH3.CONTROL)
CLEAN.DATA.TDH3.CTRL[,"log.RNA.MEDIAN.CORRECT"] <- CLEAN.DATA.TDH3.CTRL$log.RNA.MEDIAN - predict(log.RNA.MEDIAN.CORRECT, CLEAN.DATA.TDH3.CTRL) + mean(TDH3.CONTROL$log.RNA.MEDIAN)
TDH3.CONTROL <- subset(CLEAN.DATA.TDH3.CTRL,CLEAN.DATA.TDH3.CTRL$CONDITION == "CTRL")

#Correct sd
#fit<- lm(log.RNA.SD~FLOW.RUN+REP+STACK.ORDER+STACK.ORDER:FLOW.RUN+REP+ROW+COLUMN+BLOCK+BLOCK:ROW+ROW:COLUMN+STACK.ORDER:BLOCK,data=TDH3.CONTROL)
#step <- step(fit, direction="both")
log.RNA.SD.CORRECT <- lm(log.RNA.SD ~  FLOW.RUN + ROW + COLUMN,data=TDH3.CONTROL)
CLEAN.DATA.TDH3.CTRL[,"log.RNA.SD.CORRECT"] <- CLEAN.DATA.TDH3.CTRL$log.RNA.SD - predict(log.RNA.SD.CORRECT , CLEAN.DATA.TDH3.CTRL) + mean(TDH3.CONTROL$log.RNA.SD)
TDH3.CONTROL <- subset(CLEAN.DATA.TDH3.CTRL,CLEAN.DATA.TDH3.CTRL$CONDITION == "CTRL")

#Correct mad
#fit<- lm(log.RNA.MAD~FLOW.RUN+REP+STACK.ORDER+STACK.ORDER:FLOW.RUN+REP+ROW+COLUMN+BLOCK+BLOCK:ROW+ROW:COLUMN+STACK.ORDER:BLOCK,data=TDH3.CONTROL)
#step <- step(fit, direction="both")
log.RNA.MAD.CORRECT <- lm(log.RNA.MAD ~  FLOW.RUN + ROW + COLUMN,data=TDH3.CONTROL)
CLEAN.DATA.TDH3.CTRL[,"log.RNA.MAD.CORRECT"] <- CLEAN.DATA.TDH3.CTRL$log.RNA.MAD - predict(log.RNA.MAD.CORRECT , CLEAN.DATA.TDH3.CTRL) + mean(TDH3.CONTROL$log.RNA.MAD)
TDH3.CONTROL <- subset(CLEAN.DATA.TDH3.CTRL,CLEAN.DATA.TDH3.CTRL$CONDITION == "CTRL")

#Correct FSC
#fit<- lm(FSC.MEDIAN.FINAL~FLOW.RUN+REP+STACK.ORDER+STACK.ORDER:FLOW.RUN+REP+ROW+COLUMN+BLOCK+BLOCK:ROW+ROW:COLUMN+STACK.ORDER:BLOCK,data=TDH3.CONTROL)
#step <- stepAIC(fit, direction="both")
FSC.CORRECT <- lm(FSC.MEDIAN.FINAL ~  FLOW.RUN + ROW + COLUMN,data=TDH3.CONTROL)
CLEAN.DATA.TDH3.CTRL[,"FSC.CORRECT"] <- CLEAN.DATA.TDH3.CTRL$FSC.MEDIAN.FINAL - predict(FSC.CORRECT, CLEAN.DATA.TDH3.CTRL) + mean(TDH3.CONTROL$FSC.MEDIAN.FINAL)
TDH3.CONTROL <- subset(CLEAN.DATA.TDH3.CTRL,CLEAN.DATA.TDH3.CTRL$CONDITION == "CTRL")

#Back translate to linear scale
CLEAN.DATA.TDH3.CTRL[,"YFP.MEDIAN.CORRECT"] <- (exp(CLEAN.DATA.TDH3.CTRL[,"log.RNA.MEDIAN.CORRECT"]) - 0.05)
CLEAN.DATA.TDH3.CTRL[,"YFP.SD.CORRECT"] <- CLEAN.DATA.TDH3.CTRL[,"log.RNA.SD.CORRECT"] * CLEAN.DATA.TDH3.CTRL[,"YFP.MEDIAN.CORRECT"]
CLEAN.DATA.TDH3.CTRL[,"YFP.MAD.CORRECT"] <- CLEAN.DATA.TDH3.CTRL[,"log.RNA.MAD.CORRECT"] * CLEAN.DATA.TDH3.CTRL[,"YFP.MEDIAN.CORRECT"]


#PERFORM FABIEN'S OUTLIER CORRECTION FILTERING OUT ANY REPLICATE OUTSIDE OF 4*MAD

#Remove outliers
for (i in 1:nrow(CLEAN.DATA.TDH3.CTRL))
{
  CUR <- subset(CLEAN.DATA.TDH3.CTRL, STRAIN == CLEAN.DATA.TDH3.CTRL[i,"STRAIN"])
  if (CLEAN.DATA.TDH3.CTRL[i,"YFP.MEDIAN.CORRECT"] > median(CUR$YFP.MEDIAN.CORRECT) + 4*mad(CUR$YFP.MEDIAN.CORRECT) | CLEAN.DATA.TDH3.CTRL[i,"YFP.MEDIAN.CORRECT"] < median(CUR$YFP.MEDIAN.CORRECT) - 4*mad(CUR$YFP.MEDIAN.CORRECT)) {
    CLEAN.DATA.TDH3.CTRL[i,"MEDIAN.OUTLIER"] <- "YES"
  } else {
    CLEAN.DATA.TDH3.CTRL[i,"MEDIAN.OUTLIER"] <- "NO"
  }
  if (CLEAN.DATA.TDH3.CTRL[i,"YFP.SD.CORRECT"] > median(CUR$YFP.SD.CORRECT) + 4*mad(CUR$YFP.SD.CORRECT) | CLEAN.DATA.TDH3.CTRL[i,"YFP.SD.CORRECT"] < median(CUR$YFP.SD.CORRECT) - 4*mad(CUR$YFP.SD.CORRECT)) {
    CLEAN.DATA.TDH3.CTRL[i,"SD.OUTLIER"] <- "YES"
  } else {
    CLEAN.DATA.TDH3.CTRL[i,"SD.OUTLIER"] <- "NO"
  }
  if (CLEAN.DATA.TDH3.CTRL[i,"FSC.CORRECT"] > median(CUR$FSC.CORRECT) + 4*mad(CUR$FSC.CORRECT) | CLEAN.DATA.TDH3.CTRL[i,"FSC.CORRECT"] < median(CUR$FSC.CORRECT) - 4*mad(CUR$FSC.CORRECT)) {
    CLEAN.DATA.TDH3.CTRL[i,"FSC.OUTLIER"] <- "YES"
  } else {
    CLEAN.DATA.TDH3.CTRL[i,"FSC.OUTLIER"] <- "NO"
  }	
}

CLEAN.DATA.TDH3.CTRL <- subset(CLEAN.DATA.TDH3.CTRL, MEDIAN.OUTLIER == "NO" & SD.OUTLIER == "NO" & FSC.OUTLIER == "NO")

###Seperate controls and remove FSC outliers
OWN.CONTROL <- subset(CLEAN.DATA.OWN.CTRL,CLEAN.DATA.OWN.CTRL$CONDITION == "CTRL")
FSC.CORRECT <- lm(OWN.CONTROL$FSC.MEDIAN.FINAL ~ 0 + OWN.CONTROL$FLOW.RUN)
RESID <- FSC.CORRECT$resid
CLEAN.DATA.OWN.CTRL <- droplevels(CLEAN.DATA.OWN.CTRL[-which(CLEAN.DATA.OWN.CTRL$CONDITION == "CTRL")[abs(RESID) >3*mad(RESID)],])
OWN.CONTROL <- droplevels(subset(CLEAN.DATA.OWN.CTRL,CLEAN.DATA.OWN.CTRL$CONDITION == "CTRL"))

INITIAL.CORRECT <- lm(OWN.CONTROL$log.RNA.MEDIAN ~ 0 + OWN.CONTROL$FLOW.RUN)
RESID <- INITIAL.CORRECT$resid
CLEAN.DATA.OWN.CTRL<- droplevels(CLEAN.DATA.OWN.CTRL[-which(CLEAN.DATA.OWN.CTRL$CONDITION == "CTRL")[abs(RESID) > 3*mad(RESID)],])
OWN.CONTROL <- droplevels(subset(CLEAN.DATA.OWN.CTRL,CLEAN.DATA.OWN.CTRL$CONDITION == "CTRL"))

`%notin%` <- function(x,y) !(x %in% y)
RESULT<-count(OWN.CONTROL$FLOW.RUN)
RESULT.MUT<-count(CLEAN.DATA.OWN.CTRL$FLOW.RUN)
EXCLUDE1<-(RESULT.MUT[which(RESULT.MUT[,1] %notin% RESULT[,1]),1])
EXCLUDE2<-(RESULT[which(RESULT[,2] <5),1])
CLEAN.DATA.OWN.CTRL<-droplevels(CLEAN.DATA.OWN.CTRL[CLEAN.DATA.OWN.CTRL$FLOW.RUN %notin% EXCLUDE1,])
CLEAN.DATA.OWN.CTRL<-droplevels(CLEAN.DATA.OWN.CTRL[CLEAN.DATA.OWN.CTRL$FLOW.RUN %notin% EXCLUDE2,])
OWN.CONTROL <- subset(CLEAN.DATA.OWN.CTRL,CLEAN.DATA.OWN.CTRL$CONDITION == "CTRL")


###Corrections for YFP median and MAD
#Correct Median
#fit<- lm(log.RNA.MEDIAN~FLOW.RUN+REP+STACK.ORDER+STACK.ORDER:FLOW.RUN+REP+ROW+COLUMN+BLOCK+BLOCK:ROW+ROW:COLUMN+STACK.ORDER:BLOCK,data=OWN.CONTROL)
#step <- stepAIC(fit, direction="both")
log.RNA.MEDIAN.CORRECT <- lm(log.RNA.MEDIAN ~ FLOW.RUN + COLUMN + ROW, data=OWN.CONTROL)
CLEAN.DATA.OWN.CTRL[,"log.RNA.MEDIAN.CORRECT"] <- CLEAN.DATA.OWN.CTRL$log.RNA.MEDIAN - predict(log.RNA.MEDIAN.CORRECT, CLEAN.DATA.OWN.CTRL) + mean(OWN.CONTROL$log.RNA.MEDIAN)
OWN.CONTROL <- subset(CLEAN.DATA.OWN.CTRL,CLEAN.DATA.OWN.CTRL$CONDITION == "CTRL")

#Correct sd
#fit<- lm(log.RNA.SD~FLOW.RUN+REP+STACK.ORDER+STACK.ORDER:FLOW.RUN+REP+ROW+COLUMN+BLOCK+BLOCK:ROW+ROW:COLUMN+STACK.ORDER:BLOCK,data=OWN.CONTROL)
#step <- step(fit, direction="both")
log.RNA.SD.CORRECT <- lm(log.RNA.SD ~  FLOW.RUN + ROW + COLUMN + ROW:COLUMN,data=OWN.CONTROL)
CLEAN.DATA.OWN.CTRL[,"log.RNA.SD.CORRECT"] <- CLEAN.DATA.OWN.CTRL$log.RNA.SD - predict(log.RNA.SD.CORRECT , CLEAN.DATA.OWN.CTRL) + mean(OWN.CONTROL$log.RNA.SD)
OWN.CONTROL <- subset(CLEAN.DATA.OWN.CTRL,CLEAN.DATA.OWN.CTRL$CONDITION == "CTRL")

#Correct mad
#fit<- lm(log.RNA.MAD~FLOW.RUN+REP+STACK.ORDER+STACK.ORDER:FLOW.RUN+REP+ROW+COLUMN+BLOCK+BLOCK:ROW+ROW:COLUMN+STACK.ORDER:BLOCK,data=OWN.CONTROL)
#step <- step(fit, direction="both")
log.RNA.MAD.CORRECT <- lm(log.RNA.MAD ~  FLOW.RUN + ROW + COLUMN,data=OWN.CONTROL)
CLEAN.DATA.OWN.CTRL[,"log.RNA.MAD.CORRECT"] <- CLEAN.DATA.OWN.CTRL$log.RNA.MAD - predict(log.RNA.MAD.CORRECT , CLEAN.DATA.OWN.CTRL) + mean(OWN.CONTROL$log.RNA.MAD)
OWN.CONTROL <- subset(CLEAN.DATA.OWN.CTRL,CLEAN.DATA.OWN.CTRL$CONDITION == "CTRL")

#Correct FSC
#fit<- lm(FSC.MEDIAN.FINAL~FLOW.RUN+REP+STACK.ORDER+STACK.ORDER:FLOW.RUN+REP+ROW+COLUMN+BLOCK+BLOCK:ROW+ROW:COLUMN+STACK.ORDER:BLOCK,data=OWN.CONTROL)
#step <- stepAIC(fit, direction="both")
FSC.CORRECT <- lm(FSC.MEDIAN.FINAL ~  FLOW.RUN + ROW + COLUMN + ROW:COLUMN,data=OWN.CONTROL)
CLEAN.DATA.OWN.CTRL[,"FSC.CORRECT"] <- CLEAN.DATA.OWN.CTRL$FSC.MEDIAN.FINAL - predict(FSC.CORRECT, CLEAN.DATA.OWN.CTRL) + mean(OWN.CONTROL$FSC.MEDIAN.FINAL)
OWN.CONTROL <- subset(CLEAN.DATA.OWN.CTRL,CLEAN.DATA.OWN.CTRL$CONDITION == "CTRL")

#WORTH CHECKING THE SPECTRUM OF ANY STRAIN WITH FLUOR LESS THAN LOG.RNA.MEDIAN =-0.55 TO SEE WHETHER CONTAMINATION DOMINATED CELLS CAPTURED

#Back translate to linear scale
CLEAN.DATA.OWN.CTRL[,"YFP.MEDIAN.CORRECT"] <- (exp(CLEAN.DATA.OWN.CTRL[,"log.RNA.MEDIAN.CORRECT"]) - 0.05)
CLEAN.DATA.OWN.CTRL[,"YFP.SD.CORRECT"] <- CLEAN.DATA.OWN.CTRL[,"log.RNA.SD.CORRECT"] * CLEAN.DATA.OWN.CTRL[,"YFP.MEDIAN.CORRECT"]
CLEAN.DATA.OWN.CTRL[,"YFP.MAD.CORRECT"] <- CLEAN.DATA.OWN.CTRL[,"log.RNA.MAD.CORRECT"] * CLEAN.DATA.OWN.CTRL[,"YFP.MEDIAN.CORRECT"]


#PERFORM FABIEN'S OUTLIER CORRECTION FILTERING OUT ANY REPLICATE OUTSIDE OF 4*MAD

#Remove outliers
for (i in 1:nrow(CLEAN.DATA.OWN.CTRL))
{
  CUR <- subset(CLEAN.DATA.OWN.CTRL, STRAIN == CLEAN.DATA.OWN.CTRL[i,"STRAIN"])
  if (CLEAN.DATA.OWN.CTRL[i,"YFP.MEDIAN.CORRECT"] > median(CUR$YFP.MEDIAN.CORRECT) + 4*mad(CUR$YFP.MEDIAN.CORRECT) | CLEAN.DATA.OWN.CTRL[i,"YFP.MEDIAN.CORRECT"] < median(CUR$YFP.MEDIAN.CORRECT) - 4*mad(CUR$YFP.MEDIAN.CORRECT)) {
    CLEAN.DATA.OWN.CTRL[i,"MEDIAN.OUTLIER"] <- "YES"
  } else {
    CLEAN.DATA.OWN.CTRL[i,"MEDIAN.OUTLIER"] <- "NO"
  }
  if (CLEAN.DATA.OWN.CTRL[i,"YFP.SD.CORRECT"] > median(CUR$YFP.SD.CORRECT) + 4*mad(CUR$YFP.SD.CORRECT) | CLEAN.DATA.OWN.CTRL[i,"YFP.SD.CORRECT"] < median(CUR$YFP.SD.CORRECT) - 4*mad(CUR$YFP.SD.CORRECT)) {
    CLEAN.DATA.OWN.CTRL[i,"SD.OUTLIER"] <- "YES"
  } else {
    CLEAN.DATA.OWN.CTRL[i,"SD.OUTLIER"] <- "NO"
  }
  if (CLEAN.DATA.OWN.CTRL[i,"FSC.CORRECT"] > median(CUR$FSC.CORRECT) + 4*mad(CUR$FSC.CORRECT) | CLEAN.DATA.OWN.CTRL[i,"FSC.CORRECT"] < median(CUR$FSC.CORRECT) - 4*mad(CUR$FSC.CORRECT)) {
    CLEAN.DATA.OWN.CTRL[i,"FSC.OUTLIER"] <- "YES"
  } else {
    CLEAN.DATA.OWN.CTRL[i,"FSC.OUTLIER"] <- "NO"
  }	
}

CLEAN.DATA.OWN.CTRL <- subset(CLEAN.DATA.OWN.CTRL, MEDIAN.OUTLIER == "NO" & SD.OUTLIER == "NO" & FSC.OUTLIER == "NO")

#Write processed data to file
write.table(CLEAN.DATA.OWN.CTRL,"RNR1.CLEAN.CORRECT_10112016_RNR1ctrl_DCv34.csv",sep=",",quote=FALSE,row.names=FALSE)
write.table(CLEAN.DATA.TDH3.CTRL,"RNR1.CLEAN.CORRECT_10112016_TDH3ctrl_DCv34.csv",sep=",",quote=FALSE,row.names=FALSE)

#####################
#5 - Control plots  #
#####################


##############################################################################
#
#                  Strain Estimates
#
##############################################################################

#Clear memory
rm(list=ls())
options(warn=-1)

setwd("E:/wittkopp data/processed data/RNA scale analyses/DCv34")

#RNR1
CLEAN.DATA.TDH3.CTRL <- read.csv("RNR1.CLEAN.CORRECT_10112016_TDH3ctrl_DCv34.csv", header=TRUE)

#Calculate mean and noise for each sample relative to controls
TDH3.SHAM <- subset(CLEAN.DATA.TDH3.CTRL, CLEAN.DATA.TDH3.CTRL$CONDITION == "TDH3.SHAM")
TDH3.SHAM<-subset(TDH3.SHAM, TDH3.SHAM$YFP.MEDIAN.FINAL>0.8)
NEG <- subset(CLEAN.DATA.TDH3.CTRL, CLEAN.DATA.TDH3.CTRL$CONDITION == "NULL")
WT.SHAM <- subset(CLEAN.DATA.TDH3.CTRL, CLEAN.DATA.TDH3.CTRL$CONDITION == "SHAM")
WT.SHAM<-subset(WT.SHAM, (WT.SHAM$YFP.MEDIAN.FINAL<0.8)&(WT.SHAM$YFP.MEDIAN.FINAL>0.55))
#ABSOLUTE EXPRESSION SCALE
CLEAN.DATA.TDH3.CTRL[,"YFP.MEDIAN.ABS"] <- (CLEAN.DATA.TDH3.CTRL$YFP.MEDIAN.CORRECT - median(NEG$YFP.MEDIAN.CORRECT))
CLEAN.DATA.TDH3.CTRL[,"YFP.SD.ABS"]   <- CLEAN.DATA.TDH3.CTRL$YFP.SD.CORRECT - median(NEG$YFP.SD.CORRECT)
CLEAN.DATA.TDH3.CTRL[,"YFP.MAD.ABS"]   <- CLEAN.DATA.TDH3.CTRL$YFP.MAD.CORRECT - median(NEG$YFP.SD.CORRECT)

#Re-calculate mean and noise for each sample relative to controls
TDH3.SHAM <- subset(CLEAN.DATA.TDH3.CTRL, CLEAN.DATA.TDH3.CTRL$CONDITION == "TDH3.SHAM")
NEG <- subset(CLEAN.DATA.TDH3.CTRL, CLEAN.DATA.TDH3.CTRL$CONDITION == "NULL")
WT.SHAM <- subset(CLEAN.DATA.TDH3.CTRL, CLEAN.DATA.TDH3.CTRL$CONDITION == "SHAM")

#PROMOTER-SPECIFIC SCALING
CLEAN.DATA.TDH3.CTRL[,"YFP.MEDIAN.WT"] <- CLEAN.DATA.TDH3.CTRL$YFP.MEDIAN.ABS/median(WT.SHAM$YFP.MEDIAN.ABS)
CLEAN.DATA.TDH3.CTRL[,"YFP.SD.WT"]   <- CLEAN.DATA.TDH3.CTRL$YFP.SD.ABS/median(WT.SHAM$YFP.MEDIAN.ABS)
CLEAN.DATA.TDH3.CTRL[,"YFP.MAD.WT"]   <- CLEAN.DATA.TDH3.CTRL$YFP.MAD.ABS/median(WT.SHAM$YFP.MEDIAN.ABS)

#RELATIVE TO TDH3 EXPRESSION
CLEAN.DATA.TDH3.CTRL[,"YFP.MEDIAN.TDH3"] <- CLEAN.DATA.TDH3.CTRL$YFP.MEDIAN.ABS/median(TDH3.SHAM$YFP.MEDIAN.ABS)
CLEAN.DATA.TDH3.CTRL[,"YFP.SD.TDH3"]   <- CLEAN.DATA.TDH3.CTRL$YFP.SD.ABS/median(TDH3.SHAM$YFP.MEDIAN.ABS)
CLEAN.DATA.TDH3.CTRL[,"YFP.MAD.TDH3"]   <- CLEAN.DATA.TDH3.CTRL$YFP.MAD.ABS/median(TDH3.SHAM$YFP.MEDIAN.ABS)

#Re-calculate mean and noise for each sample relative to controls
TDH3.SHAM <- subset(CLEAN.DATA.TDH3.CTRL, CLEAN.DATA.TDH3.CTRL$CONDITION == "TDH3.SHAM")
NEG <- subset(CLEAN.DATA.TDH3.CTRL, CLEAN.DATA.TDH3.CTRL$CONDITION == "NULL")
WT.SHAM <- subset(CLEAN.DATA.TDH3.CTRL, CLEAN.DATA.TDH3.CTRL$CONDITION == "SHAM")

#CALCULATE NOISE METRICS
CLEAN.DATA.TDH3.CTRL[,"YFP.CV.WT"]   <- CLEAN.DATA.TDH3.CTRL$YFP.SD.WT/CLEAN.DATA.TDH3.CTRL$YFP.MEDIAN.WT
CLEAN.DATA.TDH3.CTRL[,"YFP.CV.TDH3"]   <- CLEAN.DATA.TDH3.CTRL$YFP.SD.TDH3/CLEAN.DATA.TDH3.CTRL$YFP.MEDIAN.TDH3
CLEAN.DATA.TDH3.CTRL[,"YFP.FANO.WT"]   <- CLEAN.DATA.TDH3.CTRL$YFP.SD.WT^2/CLEAN.DATA.TDH3.CTRL$YFP.MEDIAN.WT
CLEAN.DATA.TDH3.CTRL[,"YFP.FANO.TDH3"]   <- CLEAN.DATA.TDH3.CTRL$YFP.SD.TDH3^2/CLEAN.DATA.TDH3.CTRL$YFP.MEDIAN.TDH3


#Re-calculate mean and noise for each sample relative to controls
TDH3.SHAM <- subset(CLEAN.DATA.TDH3.CTRL, CLEAN.DATA.TDH3.CTRL$CONDITION == "TDH3.SHAM")
NEG <- subset(CLEAN.DATA.TDH3.CTRL, CLEAN.DATA.TDH3.CTRL$CONDITION == "NULL")
WT.SHAM <- subset(CLEAN.DATA.TDH3.CTRL, CLEAN.DATA.TDH3.CTRL$CONDITION == "SHAM")

#CALCULATE RELATIVE NOISE METRICS
CLEAN.DATA.TDH3.CTRL[,"YFP.SD.WT.REL"]   <- CLEAN.DATA.TDH3.CTRL$YFP.SD.WT/median(WT.SHAM$YFP.SD.WT)
CLEAN.DATA.TDH3.CTRL[,"YFP.SD.TDH3.REL"]   <- CLEAN.DATA.TDH3.CTRL$YFP.SD.TDH3/median(WT.SHAM$YFP.SD.TDH3)
CLEAN.DATA.TDH3.CTRL[,"YFP.CV.WT.REL"]   <- CLEAN.DATA.TDH3.CTRL$YFP.SD.WT.REL/CLEAN.DATA.TDH3.CTRL$YFP.MEDIAN.WT
CLEAN.DATA.TDH3.CTRL[,"YFP.CV.TDH3.REL"]   <- CLEAN.DATA.TDH3.CTRL$YFP.SD.TDH3.REL/CLEAN.DATA.TDH3.CTRL$YFP.MEDIAN.TDH3
CLEAN.DATA.TDH3.CTRL[,"YFP.FANO.WT.REL"]   <- CLEAN.DATA.TDH3.CTRL$YFP.SD.WT.REL^2/CLEAN.DATA.TDH3.CTRL$YFP.MEDIAN.WT
CLEAN.DATA.TDH3.CTRL[,"YFP.FANO.TDH3.REL"]   <- CLEAN.DATA.TDH3.CTRL$YFP.SD.TDH3.REL^2/CLEAN.DATA.TDH3.CTRL$YFP.MEDIAN.TDH3


#Add NA for samples where CV or FANO are undefined.
for (i in 1:nrow(CLEAN.DATA))
{
  if (CLEAN.DATA.TDH3.CTRL[i,"YFP.MEDIAN.ABS"] < 0 | CLEAN.DATA.TDH3.CTRL[i,"YFP.SD.ABS"] < 0) {
    CLEAN.DATA.TDH3.CTRL[i,"YFP.MAD.WT"] <- NA
    CLEAN.DATA.TDH3.CTRL[i,"YFP.MAD.TDH3"] <- NA
    CLEAN.DATA.TDH3.CTRL[i,"YFP.CV.WT"] <- NA
    CLEAN.DATA.TDH3.CTRL[i,"YFP.CV.TDH3"] <- NA
    CLEAN.DATA.TDH3.CTRL[i,"YFP.CV.WT.REL"] <- NA
    CLEAN.DATA.TDH3.CTRL[i,"YFP.CV.TDH3.REL"] <- NA
    CLEAN.DATA.TDH3.CTRL[i,"YFP.FANO.WT"] <- NA
    CLEAN.DATA.TDH3.CTRL[i,"YFP.FANO.TDH3"] <- NA
    CLEAN.DATA.TDH3.CTRL[i,"YFP.FANO.WT.REL"] <- NA
    CLEAN.DATA.TDH3.CTRL[i,"YFP.FANO.TDH3.REL"] <- NA
    
  }
}

CLEAN.DATA.TDH3.CTRL <- droplevels(subset(CLEAN.DATA.TDH3.CTRL,CLEAN.DATA.TDH3.CTRL$CONDITION!="CTRL"))

write.table(CLEAN.DATA.TDH3.CTRL,"RNR1.CLEAN.ADJUST_10112016_TDH3ctrl_DCv34.csv",sep=",",quote=FALSE,row.names=FALSE)

CLEAN.DATA.TDH3.CTRL <- read.csv("RNR1.CLEAN.ADJUST_10112016_TDH3ctrl_DCv34.csv", header=TRUE)

RNR1.MEAN <- aggregate(cbind(YFP.MEDIAN.ABS,YFP.SD.ABS,YFP.MAD.ABS,YFP.MEDIAN.WT,YFP.SD.WT,YFP.MAD.WT,YFP.MEDIAN.TDH3,YFP.SD.TDH3,YFP.MAD.TDH3,YFP.CV.WT,YFP.FANO.WT,YFP.CV.TDH3,YFP.FANO.TDH3,YFP.SD.WT.REL,YFP.CV.WT.REL,YFP.FANO.WT.REL,YFP.SD.TDH3.REL,YFP.CV.TDH3.REL,YFP.FANO.TDH3.REL,FSC.CORRECT) ~ STRAIN + CONDITION + ASSAY + PLATE + CTRL.GENO, data=CLEAN.DATA.TDH3.CTRL, FUN = mean)
RNR1.SD <- aggregate(cbind(YFP.MEDIAN.ABS,YFP.SD.ABS,YFP.MAD.ABS,YFP.MEDIAN.WT,YFP.SD.WT,YFP.MAD.WT,YFP.MEDIAN.TDH3,YFP.SD.TDH3,YFP.MAD.TDH3,YFP.CV.WT,YFP.FANO.WT,YFP.CV.TDH3,YFP.FANO.TDH3,YFP.SD.WT.REL,YFP.CV.WT.REL,YFP.FANO.WT.REL,YFP.SD.TDH3.REL,YFP.CV.TDH3.REL,YFP.FANO.TDH3.REL,FSC.CORRECT) ~ STRAIN + CONDITION + ASSAY + PLATE + CTRL.GENO, data=CLEAN.DATA.TDH3.CTRL, FUN = sd)
RNR1.N <- aggregate(cbind(YFP.MEDIAN.ABS,YFP.SD.ABS,YFP.MAD.ABS,YFP.MEDIAN.WT,YFP.SD.WT,YFP.MAD.WT,YFP.MEDIAN.TDH3,YFP.SD.TDH3,YFP.MAD.TDH3,YFP.CV.WT,YFP.FANO.WT,YFP.CV.TDH3,YFP.FANO.TDH3,YFP.SD.WT.REL,YFP.CV.WT.REL,YFP.FANO.WT.REL,YFP.SD.TDH3.REL,YFP.CV.TDH3.REL,YFP.FANO.TDH3.REL,FSC.CORRECT) ~ STRAIN + CONDITION + ASSAY + PLATE + CTRL.GENO, data=CLEAN.DATA.TDH3.CTRL, FUN = length)

RNR1.MEAN  <- RNR1.MEAN[which(RNR1.N[,6] >= 3),]
RNR1.SD <- RNR1.SD[which(RNR1.N[,6] >= 3),]
RNR1.N <- RNR1.N[which(RNR1.N[,6] >= 3),]

RNR1 <- cbind.data.frame(RNR1.MEAN,RNR1.SD[,6:25],RNR1.N[,6])

colnames(RNR1) <- c("STRAIN","CONDITION","ASSAY","PLATE","CTRL.GENO",paste0("MEAN.",colnames(RNR1.MEAN[,6:25])),paste0("SD.",colnames(RNR1.SD[,6:25])),"N")

write.table(RNR1,"RNR1.SUMMARY.DATA_10112016_DCv34_TDH3ctrl.txt",sep="\t",quote=FALSE,row.names=FALSE)

`%notin%` <- function(x,y) !(x %in% y)

CONTAM.CHECK.TDH3.CTRL<-droplevels(subset(CLEAN.DATA.TDH3.CTRL,CLEAN.DATA.TDH3.CTRL$CONTAM.CHECK!="NONE"))

if(all(CONTAM.CHECK.TDH3.CTRL$STRAIN %notin% RNR1$STRAIN)) {
  #All datapoints flagged as suspicious have been removed by outlier removal filters.
  print("Yay! Suspicious values eliminated. Proceed.")
}

if(any(CONTAM.CHECK.TDH3.CTRL$STRAIN %in% RNR1$STRAIN)) {
  CONTAM.CHECK.TDH3.CTRL<-CONTAM.CHECK.TDH3.CTRL[CONTAM.CHECK.TDH3.CTRL$STRAIN %in% CONTAM.CHECK.TDH3.CTRL$STRAIN[which(CONTAM.CHECK.TDH3.CTRL$STRAIN %in% RNR1$STRAIN)],]
  write.table(CONTAM.CHECK.TDH3.CTRL,"RNR1.CONTAM.CHECK_outliersexcl.csv",sep=",",quote=FALSE,row.names=FALSE)  
}


# ##############################################################################
#
# #Strain Estimates
#
# #####################################################################################
# #Clear memory
rm(list=ls())
options(warn=-1)

setwd("E:/wittkopp data/processed data/RNA scale analyses/DCv34")

#RNR1
CLEAN.DATA.OWN.CTRL <- read.csv("RNR1.CLEAN.CORRECT_10112016_RNR1ctrl_DCv34.csv", header=TRUE)

#Calculate mean and noise for each sample relative to controls
TDH3.SHAM <- subset(CLEAN.DATA.OWN.CTRL, CLEAN.DATA.OWN.CTRL$CONDITION == "TDH3.SHAM")
TDH3.SHAM<-subset(TDH3.SHAM, TDH3.SHAM$YFP.MEDIAN.FINAL>0.8)
NEG <- subset(CLEAN.DATA.OWN.CTRL, CLEAN.DATA.OWN.CTRL$CONDITION == "NULL")
WT.SHAM <- subset(CLEAN.DATA.OWN.CTRL, CLEAN.DATA.OWN.CTRL$CONDITION == "SHAM")
WT.SHAM<-subset(WT.SHAM, (WT.SHAM$YFP.MEDIAN.FINAL<0.8)&(WT.SHAM$YFP.MEDIAN.FINAL>0.55))
#ABSOLUTE EXPRESSION SCALE
CLEAN.DATA.OWN.CTRL[,"YFP.MEDIAN.ABS"] <- (CLEAN.DATA.OWN.CTRL$YFP.MEDIAN.CORRECT - median(NEG$YFP.MEDIAN.CORRECT))
CLEAN.DATA.OWN.CTRL[,"YFP.SD.ABS"]   <- CLEAN.DATA.OWN.CTRL$YFP.SD.CORRECT - median(NEG$YFP.SD.CORRECT)
CLEAN.DATA.OWN.CTRL[,"YFP.MAD.ABS"]   <- CLEAN.DATA.OWN.CTRL$YFP.MAD.CORRECT - median(NEG$YFP.SD.CORRECT)

#Re-calculate mean and noise for each sample relative to controls
TDH3.SHAM <- subset(CLEAN.DATA.OWN.CTRL, CLEAN.DATA.OWN.CTRL$CONDITION == "TDH3.SHAM")
NEG <- subset(CLEAN.DATA.OWN.CTRL, CLEAN.DATA.OWN.CTRL$CONDITION == "NULL")
WT.SHAM <- subset(CLEAN.DATA.OWN.CTRL, CLEAN.DATA.OWN.CTRL$CONDITION == "SHAM")

#PROMOTER-SPECIFIC SCALING
CLEAN.DATA.OWN.CTRL[,"YFP.MEDIAN.WT"] <- CLEAN.DATA.OWN.CTRL$YFP.MEDIAN.ABS/median(WT.SHAM$YFP.MEDIAN.ABS)
CLEAN.DATA.OWN.CTRL[,"YFP.SD.WT"]   <- CLEAN.DATA.OWN.CTRL$YFP.SD.ABS/median(WT.SHAM$YFP.MEDIAN.ABS)
CLEAN.DATA.OWN.CTRL[,"YFP.MAD.WT"]   <- CLEAN.DATA.OWN.CTRL$YFP.MAD.ABS/median(WT.SHAM$YFP.MEDIAN.ABS)

#RELATIVE TO TDH3 EXPRESSION
CLEAN.DATA.OWN.CTRL[,"YFP.MEDIAN.TDH3"] <- CLEAN.DATA.OWN.CTRL$YFP.MEDIAN.ABS/median(TDH3.SHAM$YFP.MEDIAN.ABS)
CLEAN.DATA.OWN.CTRL[,"YFP.SD.TDH3"]   <- CLEAN.DATA.OWN.CTRL$YFP.SD.ABS/median(TDH3.SHAM$YFP.MEDIAN.ABS)
CLEAN.DATA.OWN.CTRL[,"YFP.MAD.TDH3"]   <- CLEAN.DATA.OWN.CTRL$YFP.MAD.ABS/median(TDH3.SHAM$YFP.MEDIAN.ABS)

#Re-calculate mean and noise for each sample relative to controls
TDH3.SHAM <- subset(CLEAN.DATA.OWN.CTRL, CLEAN.DATA.OWN.CTRL$CONDITION == "TDH3.SHAM")
NEG <- subset(CLEAN.DATA.OWN.CTRL, CLEAN.DATA.OWN.CTRL$CONDITION == "NULL")
WT.SHAM <- subset(CLEAN.DATA.OWN.CTRL, CLEAN.DATA.OWN.CTRL$CONDITION == "SHAM")

#CALCULATE NOISE METRICS
CLEAN.DATA.OWN.CTRL[,"YFP.CV.WT"]   <- CLEAN.DATA.OWN.CTRL$YFP.SD.WT/CLEAN.DATA.OWN.CTRL$YFP.MEDIAN.WT
CLEAN.DATA.OWN.CTRL[,"YFP.CV.TDH3"]   <- CLEAN.DATA.OWN.CTRL$YFP.SD.TDH3/CLEAN.DATA.OWN.CTRL$YFP.MEDIAN.TDH3
CLEAN.DATA.OWN.CTRL[,"YFP.FANO.WT"]   <- CLEAN.DATA.OWN.CTRL$YFP.SD.WT^2/CLEAN.DATA.OWN.CTRL$YFP.MEDIAN.WT
CLEAN.DATA.OWN.CTRL[,"YFP.FANO.TDH3"]   <- CLEAN.DATA.OWN.CTRL$YFP.SD.TDH3^2/CLEAN.DATA.OWN.CTRL$YFP.MEDIAN.TDH3


#Re-calculate mean and noise for each sample relative to controls
TDH3.SHAM <- subset(CLEAN.DATA.OWN.CTRL, CLEAN.DATA.OWN.CTRL$CONDITION == "TDH3.SHAM")
NEG <- subset(CLEAN.DATA.OWN.CTRL, CLEAN.DATA.OWN.CTRL$CONDITION == "NULL")
WT.SHAM <- subset(CLEAN.DATA.OWN.CTRL, CLEAN.DATA.OWN.CTRL$CONDITION == "SHAM")

#CALCULATE RELATIVE NOISE METRICS
CLEAN.DATA.OWN.CTRL[,"YFP.SD.WT.REL"]   <- CLEAN.DATA.OWN.CTRL$YFP.SD.WT/median(WT.SHAM$YFP.SD.WT)
CLEAN.DATA.OWN.CTRL[,"YFP.SD.TDH3.REL"]   <- CLEAN.DATA.OWN.CTRL$YFP.SD.TDH3/median(WT.SHAM$YFP.SD.TDH3)
CLEAN.DATA.OWN.CTRL[,"YFP.CV.WT.REL"]   <- CLEAN.DATA.OWN.CTRL$YFP.SD.WT.REL/CLEAN.DATA.OWN.CTRL$YFP.MEDIAN.WT
CLEAN.DATA.OWN.CTRL[,"YFP.CV.TDH3.REL"]   <- CLEAN.DATA.OWN.CTRL$YFP.SD.TDH3.REL/CLEAN.DATA.OWN.CTRL$YFP.MEDIAN.TDH3
CLEAN.DATA.OWN.CTRL[,"YFP.FANO.WT.REL"]   <- CLEAN.DATA.OWN.CTRL$YFP.SD.WT.REL^2/CLEAN.DATA.OWN.CTRL$YFP.MEDIAN.WT
CLEAN.DATA.OWN.CTRL[,"YFP.FANO.TDH3.REL"]   <- CLEAN.DATA.OWN.CTRL$YFP.SD.TDH3.REL^2/CLEAN.DATA.OWN.CTRL$YFP.MEDIAN.TDH3


#Add NA for samples where CV or FANO are undefined.
for (i in 1:nrow(CLEAN.DATA))
{
  if (CLEAN.DATA.OWN.CTRL[i,"YFP.MEDIAN.ABS"] < 0 | CLEAN.DATA.OWN.CTRL[i,"YFP.SD.ABS"] < 0) {
    CLEAN.DATA.OWN.CTRL[i,"YFP.MAD.WT"] <- NA
    CLEAN.DATA.OWN.CTRL[i,"YFP.MAD.TDH3"] <- NA
    CLEAN.DATA.OWN.CTRL[i,"YFP.CV.WT"] <- NA
    CLEAN.DATA.OWN.CTRL[i,"YFP.CV.TDH3"] <- NA
    CLEAN.DATA.OWN.CTRL[i,"YFP.CV.WT.REL"] <- NA
    CLEAN.DATA.OWN.CTRL[i,"YFP.CV.TDH3.REL"] <- NA
    CLEAN.DATA.OWN.CTRL[i,"YFP.FANO.WT"] <- NA
    CLEAN.DATA.OWN.CTRL[i,"YFP.FANO.TDH3"] <- NA
    CLEAN.DATA.OWN.CTRL[i,"YFP.FANO.WT.REL"] <- NA
    CLEAN.DATA.OWN.CTRL[i,"YFP.FANO.TDH3.REL"] <- NA
    
  }
}

CLEAN.DATA.OWN.CTRL <- droplevels(subset(CLEAN.DATA.OWN.CTRL,CLEAN.DATA.OWN.CTRL$CONDITION!="CTRL"))

write.table(CLEAN.DATA.OWN.CTRL,"RNR1.CLEAN.ADJUST_10112016_RNR1ctrl_DCv34.csv",sep=",",quote=FALSE,row.names=FALSE)

CLEAN.DATA.OWN.CTRL <- read.csv("RNR1.CLEAN.ADJUST_10112016_RNR1ctrl_DCv34.csv", header=TRUE)

RNR1.MEAN <- aggregate(cbind(YFP.MEDIAN.ABS,YFP.SD.ABS,YFP.MAD.ABS,YFP.MEDIAN.WT,YFP.SD.WT,YFP.MAD.WT,YFP.MEDIAN.TDH3,YFP.SD.TDH3,YFP.MAD.TDH3,YFP.CV.WT,YFP.FANO.WT,YFP.CV.TDH3,YFP.FANO.TDH3,YFP.SD.WT.REL,YFP.CV.WT.REL,YFP.FANO.WT.REL,YFP.SD.TDH3.REL,YFP.CV.TDH3.REL,YFP.FANO.TDH3.REL,FSC.CORRECT) ~ STRAIN + CONDITION + ASSAY + PLATE + CTRL.GENO, data=CLEAN.DATA.OWN.CTRL, FUN = mean)
RNR1.SD <- aggregate(cbind(YFP.MEDIAN.ABS,YFP.SD.ABS,YFP.MAD.ABS,YFP.MEDIAN.WT,YFP.SD.WT,YFP.MAD.WT,YFP.MEDIAN.TDH3,YFP.SD.TDH3,YFP.MAD.TDH3,YFP.CV.WT,YFP.FANO.WT,YFP.CV.TDH3,YFP.FANO.TDH3,YFP.SD.WT.REL,YFP.CV.WT.REL,YFP.FANO.WT.REL,YFP.SD.TDH3.REL,YFP.CV.TDH3.REL,YFP.FANO.TDH3.REL,FSC.CORRECT) ~ STRAIN + CONDITION + ASSAY + PLATE + CTRL.GENO, data=CLEAN.DATA.OWN.CTRL, FUN = sd)
RNR1.N <- aggregate(cbind(YFP.MEDIAN.ABS,YFP.SD.ABS,YFP.MAD.ABS,YFP.MEDIAN.WT,YFP.SD.WT,YFP.MAD.WT,YFP.MEDIAN.TDH3,YFP.SD.TDH3,YFP.MAD.TDH3,YFP.CV.WT,YFP.FANO.WT,YFP.CV.TDH3,YFP.FANO.TDH3,YFP.SD.WT.REL,YFP.CV.WT.REL,YFP.FANO.WT.REL,YFP.SD.TDH3.REL,YFP.CV.TDH3.REL,YFP.FANO.TDH3.REL,FSC.CORRECT) ~ STRAIN + CONDITION + ASSAY + PLATE + CTRL.GENO, data=CLEAN.DATA.OWN.CTRL, FUN = length)

RNR1.MEAN  <- RNR1.MEAN[which(RNR1.N[,6] >= 3),]
RNR1.SD <- RNR1.SD[which(RNR1.N[,6] >= 3),]
RNR1.N <- RNR1.N[which(RNR1.N[,6] >= 3),]

RNR1 <- cbind.data.frame(RNR1.MEAN,RNR1.SD[,6:25],RNR1.N[,6])

colnames(RNR1) <- c("STRAIN","CONDITION","ASSAY","PLATE","CTRL.GENO",paste0("MEAN.",colnames(RNR1.MEAN[,6:25])),paste0("SD.",colnames(RNR1.SD[,6:25])),"N")

write.table(RNR1,"RNR1.SUMMARY.DATA_10112016_DCv34_RNR1ctrl.txt",sep="\t",quote=FALSE,row.names=FALSE)

`%notin%` <- function(x,y) !(x %in% y)

CONTAM.CHECK.OWN.CTRL<-droplevels(subset(CLEAN.DATA.OWN.CTRL,CLEAN.DATA.OWN.CTRL$CONTAM.CHECK!="NONE"))

if(all(CONTAM.CHECK.OWN.CTRL$STRAIN %notin% RNR1$STRAIN)) {
  #All datapoints flagged as suspicious have been removed by outlier removal filters.
  print("Yay! Suspicious values eliminated. Proceed.")
}

if(any(CONTAM.CHECK.OWN.CTRL$STRAIN %in% RNR1$STRAIN)) {
  CONTAM.CHECK.OWN.CTRL<-CONTAM.CHECK.OWN.CTRL[CONTAM.CHECK.OWN.CTRL$STRAIN %in% CONTAM.CHECK.OWN.CTRL$STRAIN[which(CONTAM.CHECK.OWN.CTRL$STRAIN %in% RNR1$STRAIN)],]
  write.table(CONTAM.CHECK.OWN.CTRL,"RNR1.CONTAM.CHECK_outliersexcl.csv",sep=",",quote=FALSE,row.names=FALSE,append=TRUE)  
}


###############################
#HIERARCHICAL PERMUTATION TEST#
###############################

#Clear memory
rm(list=ls())
options(warn=-1)

library(permute)
library(Hmisc)

N.PERM <- 10000

#Permutation function

SHUFFLE <- function(STRAIN.LIST = STRAIN.LIST[[i]],WT.LIST = WT.LIST) {
  SAMPLE <- c(STRAIN.LIST,WT.LIST[[sample(1:length(WT.LIST),1)]])
  OUTPUT <- sample(SAMPLE,length(SAMPLE))
  DIST.SIM <- abs(mean(OUTPUT[1:length(STRAIN.LIST)]) - mean(OUTPUT[(length(STRAIN.LIST)+1):length(SAMPLE)]))
  DIST.OBS <- abs(mean(SAMPLE[1:length(STRAIN.LIST)]) - mean(SAMPLE[(length(STRAIN.LIST)+1):length(SAMPLE)]))
  
  DIFF <- DIST.OBS - DIST.SIM
  
  return(DIFF)
}

parent.dir <- PATH
setwd(parent.dir)

####RNR1 Analysis date 10112016#########################################################################

ALL.REP <- read.csv("RNR1.CLEAN.ADJUST_10112016_RNR1ctrl_DCv34.csv",header=TRUE)
ALL.REP <- subset(ALL.REP, MEDIAN.OUTLIER == "NO" & SD.OUTLIER == "NO" & FSC.OUTLIER == "NO")

SUMMARY <- read.table("RNR1.SUMMARY.DATA_10112016_DCv34_RNR1ctrl.txt",header=TRUE)

#SUMMARY <- droplevels(subset(SUMMARY, STRAIN != "1139"))

WT <- subset(ALL.REP, CONDITION == "SHAM")
LOW.FSC <- median(WT$FSC.CORRECT) - 4*mad(WT$FSC.CORRECT)
HIGH.FSC <- median(WT$FSC.CORRECT) + 4*mad(WT$FSC.CORRECT)
LOW.FLUO <- median(WT$YFP.MEDIAN.CORRECT) - 4*mad(WT$YFP.MEDIAN.CORRECT)
HIGH.FLUO <- median(WT$YFP.MEDIAN.CORRECT) + 4*mad(WT$YFP.MEDIAN.CORRECT)
WT <- subset(WT, FSC.CORRECT > LOW.FSC & FSC.CORRECT < HIGH.FSC & YFP.MEDIAN.CORRECT > LOW.FLUO & YFP.MEDIAN.CORRECT < HIGH.FLUO)

#MEDIAN

#Create list of WT phenotype grouped by strain

N.WT <- length(unique(WT$STRAIN))
WT.LIST <- vector('list',N.WT)
for (i in 1:N.WT)
{
  CUR.WT <- subset(WT, STRAIN == unique(WT$STRAIN)[i])
  WT.LIST[[i]] <- CUR.WT$YFP.MEDIAN.CORRECT 
}


#Create list of all strain phenotypes grouped by strain

SUMMARY <- SUMMARY[order(SUMMARY$STRAIN),]
ALL.REP <- droplevels(subset(ALL.REP,STRAIN %in% as.character(SUMMARY$STRAIN)))
ALL.REP <- ALL.REP[order(ALL.REP$STRAIN),]
N.STRAIN <- length(unique(ALL.REP$STRAIN))
STRAIN.LIST <- vector('list',N.STRAIN)
for (i in 1:N.STRAIN)
{
  CUR.STRAIN <- subset(ALL.REP, STRAIN == unique(ALL.REP$STRAIN)[i])
  STRAIN.LIST[[i]] <- CUR.STRAIN$YFP.MEDIAN.CORRECT
}


for (i in 1:length(STRAIN.LIST))
{
  DIFF <- sapply(1:N.PERM,FUN=function(x)	SHUFFLE(STRAIN.LIST = STRAIN.LIST[[i]],WT.LIST = WT.LIST))
  P.VAL <- (length(which(DIFF < 0)) + 1) / (length(which(DIFF < 0)) + length(which(DIFF > 0)) + 1) 	
  SUMMARY[i,"P.VAL.MEDIAN"] <- P.VAL
  print(round(i/length(STRAIN.LIST)*100,1))
}

#NOISE: CV

#Create list of WT phenotype grouped by strain

N.WT <- length(unique(WT$STRAIN))
WT.LIST <- vector('list',N.WT)
for (i in 1:N.WT)
{
  CUR.WT <- subset(WT, STRAIN == unique(WT$STRAIN)[i])
  WT.LIST[[i]] <- CUR.WT$YFP.CV.WT
}


#Create list of all strain phenotypes grouped by strain

SUMMARY <- SUMMARY[order(SUMMARY$STRAIN),]
ALL.REP <- droplevels(subset(ALL.REP,STRAIN %in% as.character(SUMMARY$STRAIN)))
ALL.REP <- ALL.REP[order(ALL.REP$STRAIN),]
N.STRAIN <- length(unique(ALL.REP$STRAIN))
STRAIN.LIST <- vector('list',N.STRAIN)
for (i in 1:N.STRAIN)
{
  CUR.STRAIN <- subset(ALL.REP, STRAIN == unique(ALL.REP$STRAIN)[i])
  STRAIN.LIST[[i]] <- CUR.STRAIN$YFP.CV.WT
}


for (i in 1:length(STRAIN.LIST))
{
  DIFF <- sapply(1:N.PERM,FUN=function(x)	SHUFFLE(STRAIN.LIST = STRAIN.LIST[[i]],WT.LIST = WT.LIST))
  P.VAL <- (length(which(DIFF < 0))+1) / (length(which(DIFF < 0)) + length(which(DIFF > 0)) + 1) 	
  SUMMARY[i,"P.VAL.CV.WT"] <- P.VAL
  print(round(i/length(STRAIN.LIST)*100,1))
}

#NOISE: FANO

#Create list of WT phenotype grouped by strain

N.WT <- length(unique(WT$STRAIN))
WT.LIST <- vector('list',N.WT)
for (i in 1:N.WT)
{
  CUR.WT <- subset(WT, STRAIN == unique(WT$STRAIN)[i])
  WT.LIST[[i]] <- CUR.WT$YFP.FANO.WT
}


#Create list of all strain phenotypes grouped by strain

SUMMARY <- SUMMARY[order(SUMMARY$STRAIN),]
ALL.REP <- droplevels(subset(ALL.REP,STRAIN %in% as.character(SUMMARY$STRAIN)))
ALL.REP <- ALL.REP[order(ALL.REP$STRAIN),]
N.STRAIN <- length(unique(ALL.REP$STRAIN))
STRAIN.LIST <- vector('list',N.STRAIN)
for (i in 1:N.STRAIN)
{
  CUR.STRAIN <- subset(ALL.REP, STRAIN == unique(ALL.REP$STRAIN)[i])
  STRAIN.LIST[[i]] <- CUR.STRAIN$YFP.FANO.WT
}


for (i in 1:length(STRAIN.LIST))
{
  DIFF <- sapply(1:N.PERM,FUN=function(x)	SHUFFLE(STRAIN.LIST = STRAIN.LIST[[i]],WT.LIST = WT.LIST))
  P.VAL <- (length(which(DIFF < 0))+1) / (length(which(DIFF < 0)) + length(which(DIFF > 0)) + 1) 	
  SUMMARY[i,"P.VAL.FANO.WT"] <- P.VAL
  print(round(i/length(STRAIN.LIST)*100,1))
}

#FSC MEDIAN

#Create list of WT phenotype grouped by strain

N.WT <- length(unique(WT$STRAIN))
WT.LIST <- vector('list',N.WT)
for (i in 1:N.WT)
{
  CUR.WT <- subset(WT, STRAIN == unique(WT$STRAIN)[i])
  WT.LIST[[i]] <- CUR.WT$FSC.CORRECT
}


#Create list of all strain phenotypes grouped by strain

SUMMARY <- SUMMARY[order(SUMMARY$STRAIN),]
ALL.REP <- droplevels(subset(ALL.REP,STRAIN %in% as.character(SUMMARY$STRAIN)))
ALL.REP <- ALL.REP[order(ALL.REP$STRAIN),]
N.STRAIN <- length(unique(ALL.REP$STRAIN))
STRAIN.LIST <- vector('list',N.STRAIN)
for (i in 1:N.STRAIN)
{
  CUR.STRAIN <- subset(ALL.REP, STRAIN == unique(ALL.REP$STRAIN)[i])
  STRAIN.LIST[[i]] <- CUR.STRAIN$FSC.CORRECT
}


for (i in 1:length(STRAIN.LIST))
{
  DIFF <- sapply(1:N.PERM,FUN=function(x)	SHUFFLE(STRAIN.LIST = STRAIN.LIST[[i]],WT.LIST = WT.LIST))
  P.VAL <- (length(which(DIFF < 0))+1) / (length(which(DIFF < 0)) + length(which(DIFF > 0)) + 1) 	
  SUMMARY[i,"P.VAL.FSC"] <- P.VAL
  print(round(i/length(STRAIN.LIST)*100,1))
}

write.table(SUMMARY,"RNR1.SUMMARY.DCv34.Pvals_RNR1ctrl.txt",sep="\t",quote=FALSE,row.names=FALSE)

#_____________________________________________________________________________________________________________________________________

#############
#  RNR2
############

########################################
#  CLEANING
########################################

#Clear memory
rm(list=ls())
options(warn=-1)


#Set working directory
parent.dir <- PATH
setwd(parent.dir)

#Load experiment setup
SETUP <- read.csv("Template_RNR2.csv",header=TRUE,as.is=TRUE)
TESTNAMES <- list.files("PATH/RNR2",pattern=".fcs",full.name=TRUE,recursive=TRUE,include.dirs=TRUE)
SETUP<-subset(SETUP, SETUP$FILENAMES %in% TESTNAMES, drop = TRUE)
SETUP[,"COUNTER"] <- c(1:nrow(SETUP))

#Determine Hard Gates
GATES <- GATE.CALIB(SETUP$FILENAMES[1])

#Clean Data
Output <- apply(SETUP,1,CLEANING,GATES=GATES)
OUTPUT <- as.data.frame(Output[[1]])
OUTPUT[2:nrow(SETUP),1:ncol(OUTPUT)] <- NA
for (i in 2:nrow(SETUP))
{
  if (is.null(Output[[i]]))
  {} 
  else {
    OUTPUT[i,] <- Output[[i]]
  }
}

write.table(OUTPUT,"RNR2_Experiment.Output_DCv34.txt",row.names=FALSE,sep="\t")

CLEAN <- cbind.data.frame(SETUP[,1:(ncol(SETUP)-2)],OUTPUT)

write.table(CLEAN,"RNR2_Clean.Data_DCv34.txt",row.names=FALSE,sep="\t")

################################################################
#     PLATE CORRECTION & OUTLIER REMOVAL
################################################################

#Clear memory
rm(list=ls())
options(warn=-1)

#####################
#1-LOADING LIBRARIES#
#####################
library(flowCore)
library(flowClust)
library(flowViz)
library(pcaPP)
library(mixtools)
library(plyr)
library(MASS)
box <- graphics::box
options(warn=-1)

####################################
#2- Quality Control and Corrections#
####################################

parent.dir <- PATH
setwd(parent.dir)

DATA.TYPE <- c(
  "factor","factor","factor","factor","factor","factor","factor","factor","factor","factor","factor","factor","factor","factor","factor",
  "integer","integer","integer","integer",
  "numeric","numeric","numeric","numeric","numeric","numeric","numeric","numeric","numeric","numeric","numeric","numeric","numeric","numeric","numeric","numeric","numeric","numeric","numeric","numeric","numeric","numeric","numeric","numeric","numeric",
  "integer")

###Read in data sets
CLEAN.DATA <- read.table("RNR2_Clean.Data_DCv34.txt",colClasses=DATA.TYPE,header=TRUE, sep="\t")

###Remove samples with NA
CLEAN.DATA <- CLEAN.DATA[complete.cases(CLEAN.DATA),]

##Translate into log scale for appropriate correction
CLEAN.DATA[,"log.RNA.MEDIAN"] <- log((CLEAN.DATA[,"YFP.MEDIAN.FINAL"]) + 0.05)
CLEAN.DATA[,"log.RNA.MAD"] <- CLEAN.DATA[,"YFP.MAD.FINAL"] / (CLEAN.DATA[,"YFP.MEDIAN.FINAL"] + 0.05)
CLEAN.DATA[,"log.RNA.SD"] <- CLEAN.DATA[,"YFP.SD.FINAL"] / (CLEAN.DATA[,"YFP.MEDIAN.FINAL"] + 0.05)

###Remove samples with less than 1000 events
CLEAN.DATA <- subset(CLEAN.DATA,CLEAN.DATA$COUNTS.FINAL > 1000)

####Add variable to ID whether control is own promoter or TDH3
CLEAN.DATA[,"CTRL.GENO"]<-as.factor("TDH3")

###Select desired data / Drop known bad samples


###Seperate controls and remove FSC outliers
CONTROL <- subset(CLEAN.DATA,CLEAN.DATA$CONDITION == "CTRL")
FSC.CORRECT <- lm(CONTROL$FSC.MEDIAN.FINAL ~ 0 + CONTROL$FLOW.RUN)
RESID <- FSC.CORRECT$resid
CLEAN.DATA <- droplevels(CLEAN.DATA[-which(CLEAN.DATA$CONDITION == "CTRL")[abs(RESID) > 3*mad(RESID)],])
CONTROL <- droplevels(subset(CLEAN.DATA,CLEAN.DATA$CONDITION == "CTRL"))

INITIAL.CORRECT <- lm(CONTROL$log.RNA.MEDIAN ~ 0 + CONTROL$FLOW.RUN)
RESID <- INITIAL.CORRECT$resid
CLEAN.DATA<- droplevels(CLEAN.DATA[-which(CLEAN.DATA$CONDITION == "CTRL")[abs(RESID) > 3*mad(RESID)],])
CONTROL <- droplevels(subset(CLEAN.DATA,CLEAN.DATA$CONDITION == "CTRL"))

`%notin%` <- function(x,y) !(x %in% y)
RESULT<-count(CONTROL$FLOW.RUN)
RESULT.MUT<-count(CLEAN.DATA$FLOW.RUN)
EXCLUDE1<-(RESULT.MUT[which(RESULT.MUT[,1] %notin% RESULT[,1]),1])
EXCLUDE2<-(RESULT[which(RESULT[,2] <5),1])
CLEAN.DATA<-droplevels(CLEAN.DATA[CLEAN.DATA$FLOW.RUN %notin% EXCLUDE1,])
CLEAN.DATA<-droplevels(CLEAN.DATA[CLEAN.DATA$FLOW.RUN %notin% EXCLUDE2,])
CONTROL <- subset(CLEAN.DATA,CLEAN.DATA$CONDITION == "CTRL")


###Corrections for YFP median and MAD
#Correct Median
#fit<- lm(log.RNA.MEDIAN~FLOW.RUN+REP+FLOW.RUN+REP+ROW+COLUMN+BLOCK+BLOCK:ROW+ROW:COLUMN+BLOCK,data=CONTROL)
#step <- stepAIC(fit, direction="both")
log.RNA.MEDIAN.CORRECT <- lm(log.RNA.MEDIAN ~ FLOW.RUN + COLUMN + ROW, data=CONTROL)
CLEAN.DATA[,"log.RNA.MEDIAN.CORRECT"] <- CLEAN.DATA$log.RNA.MEDIAN - predict(log.RNA.MEDIAN.CORRECT, CLEAN.DATA) + mean(CONTROL$log.RNA.MEDIAN)
CONTROL <- subset(CLEAN.DATA,CLEAN.DATA$CONDITION == "CTRL")

#Correct sd
#fit<- lm(log.RNA.SD~FLOW.RUN+REP+FLOW.RUN+REP+ROW+COLUMN+BLOCK+BLOCK:ROW+ROW:COLUMN+BLOCK,data=CONTROL)
#step <- step(fit, direction="both")
log.RNA.SD.CORRECT <- lm(log.RNA.SD ~  FLOW.RUN + ROW + COLUMN,data=CONTROL)
CLEAN.DATA[,"log.RNA.SD.CORRECT"] <- CLEAN.DATA$log.RNA.SD - predict(log.RNA.SD.CORRECT , CLEAN.DATA) + mean(CONTROL$log.RNA.SD)
CONTROL <- subset(CLEAN.DATA,CLEAN.DATA$CONDITION == "CTRL")

#Correct mad
#fit<- lm(log.RNA.MAD~FLOW.RUN+REP+FLOW.RUN+REP+ROW+COLUMN+BLOCK+BLOCK:ROW+ROW:COLUMN+BLOCK,data=CONTROL)
#step <- step(fit, direction="both")
log.RNA.MAD.CORRECT <- lm(log.RNA.MAD ~  FLOW.RUN + ROW + COLUMN,data=CONTROL)
CLEAN.DATA[,"log.RNA.MAD.CORRECT"] <- CLEAN.DATA$log.RNA.MAD - predict(log.RNA.MAD.CORRECT , CLEAN.DATA) + mean(CONTROL$log.RNA.MAD)
CONTROL <- subset(CLEAN.DATA,CLEAN.DATA$CONDITION == "CTRL")

#Correct FSC
#fit<- lm(FSC.MEDIAN.FINAL~FLOW.RUN+REP+FLOW.RUN+REP+ROW+COLUMN+BLOCK+BLOCK:ROW+ROW:COLUMN+BLOCK,data=CONTROL)
#step <- stepAIC(fit, direction="both")
FSC.CORRECT <- lm(FSC.MEDIAN.FINAL ~  FLOW.RUN + ROW + COLUMN + ROW:COLUMN,data=CONTROL)
CLEAN.DATA[,"FSC.CORRECT"] <- CLEAN.DATA$FSC.MEDIAN.FINAL - predict(FSC.CORRECT, CLEAN.DATA) + mean(CONTROL$FSC.MEDIAN.FINAL)
CONTROL <- subset(CLEAN.DATA,CLEAN.DATA$CONDITION == "CTRL")


#Back translate to linear scale
CLEAN.DATA[,"YFP.MEDIAN.CORRECT"] <- (exp(CLEAN.DATA[,"log.RNA.MEDIAN.CORRECT"]) - 0.05)
CLEAN.DATA[,"YFP.SD.CORRECT"] <- CLEAN.DATA[,"log.RNA.SD.CORRECT"] * CLEAN.DATA[,"YFP.MEDIAN.CORRECT"]
CLEAN.DATA[,"YFP.MAD.CORRECT"] <- CLEAN.DATA[,"log.RNA.MAD.CORRECT"] * CLEAN.DATA[,"YFP.MEDIAN.CORRECT"]


#PERFORM FABIEN'S OUTLIER CORRECTION FILTERING OUT ANY REPLICATE OUTSIDE OF 4*MAD

#Remove outliers
for (i in 1:nrow(CLEAN.DATA))
{
  CUR <- subset(CLEAN.DATA, STRAIN == CLEAN.DATA[i,"STRAIN"])
  if (CLEAN.DATA[i,"YFP.MEDIAN.CORRECT"] > median(CUR$YFP.MEDIAN.CORRECT) + 4*mad(CUR$YFP.MEDIAN.CORRECT) | CLEAN.DATA[i,"YFP.MEDIAN.CORRECT"] < median(CUR$YFP.MEDIAN.CORRECT) - 4*mad(CUR$YFP.MEDIAN.CORRECT)) {
    CLEAN.DATA[i,"MEDIAN.OUTLIER"] <- "YES"
  } else {
    CLEAN.DATA[i,"MEDIAN.OUTLIER"] <- "NO"
  }
  if (CLEAN.DATA[i,"YFP.SD.CORRECT"] > median(CUR$YFP.SD.CORRECT) + 4*mad(CUR$YFP.SD.CORRECT) | CLEAN.DATA[i,"YFP.SD.CORRECT"] < median(CUR$YFP.SD.CORRECT) - 4*mad(CUR$YFP.SD.CORRECT)) {
    CLEAN.DATA[i,"SD.OUTLIER"] <- "YES"
  } else {
    CLEAN.DATA[i,"SD.OUTLIER"] <- "NO"
  }
  if (CLEAN.DATA[i,"FSC.CORRECT"] > median(CUR$FSC.CORRECT) + 4*mad(CUR$FSC.CORRECT) | CLEAN.DATA[i,"FSC.CORRECT"] < median(CUR$FSC.CORRECT) - 4*mad(CUR$FSC.CORRECT)) {
    CLEAN.DATA[i,"FSC.OUTLIER"] <- "YES"
  } else {
    CLEAN.DATA[i,"FSC.OUTLIER"] <- "NO"
  }	
}

CLEAN.DATA <- subset(CLEAN.DATA, MEDIAN.OUTLIER == "NO" & SD.OUTLIER == "NO" & FSC.OUTLIER == "NO")


#Write processed data to file
write.table(CLEAN.DATA,"RNR2.CLEAN.CORRECT_DCv34.csv",sep=",",quote=FALSE,row.names=FALSE)

#####################
#5 - Control plots  #
#####################

CONTROL <- droplevels(subset(CLEAN.DATA,CLEAN.DATA$CONDITION == "CTRL"))
SHAM <- droplevels(subset(CLEAN.DATA,CLEAN.DATA$CONDITION == "SHAM"))
TDH3.SHAM <- droplevels(subset(CLEAN.DATA,CLEAN.DATA$CONDITION == "TDH3.SHAM"))


# ##############################################################################
#
# #Strain Estimates
#
# #####################################################################################
# #Clear memory
rm(list=ls())
options(warn=-1)

setwd("E:/wittkopp data/processed data/RNA scale analyses/DCv34")

#RNR2
CLEAN.DATA <- read.csv("RNR2.CLEAN.CORRECT_DCv34.csv", header=TRUE)

#Calculate mean and noise for each sample relative to controls
TDH3.SHAM <- subset(CLEAN.DATA, CLEAN.DATA$CONDITION == "TDH3.SHAM")
NEG <- subset(CLEAN.DATA, CLEAN.DATA$CONDITION == "NULL")
NEG<-subset(NEG,NEG$log.RNA.MEDIAN< -0.5)
WT.SHAM <- subset(CLEAN.DATA, CLEAN.DATA$CONDITION == "SHAM")

#ABSOLUTE EXPRESSION SCALE
CLEAN.DATA[,"YFP.MEDIAN.ABS"] <- (CLEAN.DATA$YFP.MEDIAN.CORRECT - median(NEG$YFP.MEDIAN.CORRECT))
CLEAN.DATA[,"YFP.SD.ABS"]   <- CLEAN.DATA$YFP.SD.CORRECT - median(NEG$YFP.SD.CORRECT)
CLEAN.DATA[,"YFP.MAD.ABS"]   <- CLEAN.DATA$YFP.MAD.CORRECT - median(NEG$YFP.SD.CORRECT)

#Re-calculate mean and noise for each sample relative to controls
TDH3.SHAM <- subset(CLEAN.DATA, CLEAN.DATA$CONDITION == "TDH3.SHAM")
NEG <- subset(CLEAN.DATA, CLEAN.DATA$CONDITION == "NULL")
WT.SHAM <- subset(CLEAN.DATA, CLEAN.DATA$CONDITION == "SHAM")

#PROMOTER-SPECIFIC SCALING
CLEAN.DATA[,"YFP.MEDIAN.WT"] <- CLEAN.DATA$YFP.MEDIAN.ABS/median(WT.SHAM$YFP.MEDIAN.ABS)
CLEAN.DATA[,"YFP.SD.WT"]   <- CLEAN.DATA$YFP.SD.ABS/median(WT.SHAM$YFP.MEDIAN.ABS)
CLEAN.DATA[,"YFP.MAD.WT"]   <- CLEAN.DATA$YFP.MAD.ABS/median(WT.SHAM$YFP.MEDIAN.ABS)

#RELATIVE TO TDH3 EXPRESSION
CLEAN.DATA[,"YFP.MEDIAN.TDH3"] <- CLEAN.DATA$YFP.MEDIAN.ABS/median(TDH3.SHAM$YFP.MEDIAN.ABS)
CLEAN.DATA[,"YFP.SD.TDH3"]   <- CLEAN.DATA$YFP.SD.ABS/median(TDH3.SHAM$YFP.MEDIAN.ABS)
CLEAN.DATA[,"YFP.MAD.TDH3"]   <- CLEAN.DATA$YFP.MAD.ABS/median(TDH3.SHAM$YFP.MEDIAN.ABS)

#Re-calculate mean and noise for each sample relative to controls
TDH3.SHAM <- subset(CLEAN.DATA, CLEAN.DATA$CONDITION == "TDH3.SHAM")
NEG <- subset(CLEAN.DATA, CLEAN.DATA$CONDITION == "NULL")
WT.SHAM <- subset(CLEAN.DATA, CLEAN.DATA$CONDITION == "SHAM")

#CALCULATE NOISE METRICS
CLEAN.DATA[,"YFP.CV.WT"]   <- CLEAN.DATA$YFP.SD.WT/CLEAN.DATA$YFP.MEDIAN.WT
CLEAN.DATA[,"YFP.CV.TDH3"]   <- CLEAN.DATA$YFP.SD.TDH3/CLEAN.DATA$YFP.MEDIAN.TDH3
CLEAN.DATA[,"YFP.FANO.WT"]   <- CLEAN.DATA$YFP.SD.WT^2/CLEAN.DATA$YFP.MEDIAN.WT
CLEAN.DATA[,"YFP.FANO.TDH3"]   <- CLEAN.DATA$YFP.SD.TDH3^2/CLEAN.DATA$YFP.MEDIAN.TDH3


#Re-calculate mean and noise for each sample relative to controls
TDH3.SHAM <- subset(CLEAN.DATA, CLEAN.DATA$CONDITION == "TDH3.SHAM")
NEG <- subset(CLEAN.DATA, CLEAN.DATA$CONDITION == "NULL")
WT.SHAM <- subset(CLEAN.DATA, CLEAN.DATA$CONDITION == "SHAM")

#CALCULATE RELATIVE NOISE METRICS
CLEAN.DATA[,"YFP.SD.WT.REL"]   <- CLEAN.DATA$YFP.SD.WT/median(WT.SHAM$YFP.SD.WT)
CLEAN.DATA[,"YFP.SD.TDH3.REL"]   <- CLEAN.DATA$YFP.SD.TDH3/median(WT.SHAM$YFP.SD.TDH3)
CLEAN.DATA[,"YFP.CV.WT.REL"]   <- CLEAN.DATA$YFP.SD.WT.REL/CLEAN.DATA$YFP.MEDIAN.WT
CLEAN.DATA[,"YFP.CV.TDH3.REL"]   <- CLEAN.DATA$YFP.SD.TDH3.REL/CLEAN.DATA$YFP.MEDIAN.TDH3
CLEAN.DATA[,"YFP.FANO.WT.REL"]   <- CLEAN.DATA$YFP.SD.WT.REL^2/CLEAN.DATA$YFP.MEDIAN.WT
CLEAN.DATA[,"YFP.FANO.TDH3.REL"]   <- CLEAN.DATA$YFP.SD.TDH3.REL^2/CLEAN.DATA$YFP.MEDIAN.TDH3


#Add NA for samples where CV or FANO are undefined.
for (i in 1:nrow(CLEAN.DATA))
{
  if (CLEAN.DATA[i,"YFP.MEDIAN.ABS"] < 0 | CLEAN.DATA[i,"YFP.SD.ABS"] < 0) {
    CLEAN.DATA[i,"YFP.MAD.WT"] <- NA
    CLEAN.DATA[i,"YFP.MAD.TDH3"] <- NA
    CLEAN.DATA[i,"YFP.CV.WT"] <- NA
    CLEAN.DATA[i,"YFP.CV.TDH3"] <- NA
    CLEAN.DATA[i,"YFP.CV.WT.REL"] <- NA
    CLEAN.DATA[i,"YFP.CV.TDH3.REL"] <- NA
    CLEAN.DATA[i,"YFP.FANO.WT"] <- NA
    CLEAN.DATA[i,"YFP.FANO.TDH3"] <- NA
    CLEAN.DATA[i,"YFP.FANO.WT.REL"] <- NA
    CLEAN.DATA[i,"YFP.FANO.TDH3.REL"] <- NA
    
  }
}

CLEAN.DATA <- droplevels(subset(CLEAN.DATA,CLEAN.DATA$CONDITION!="CTRL"))

write.table(CLEAN.DATA,"RNR2.CLEAN.ADJUST_DCv34.csv",sep=",",quote=FALSE,row.names=FALSE)

CLEAN.DATA <- read.csv("RNR2.CLEAN.ADJUST_DCv34.csv", header=TRUE)

RNR2.MEAN <- aggregate(cbind(YFP.MEDIAN.ABS,YFP.SD.ABS,YFP.MAD.ABS,YFP.MEDIAN.WT,YFP.SD.WT,YFP.MAD.WT,YFP.MEDIAN.TDH3,YFP.SD.TDH3,YFP.MAD.TDH3,YFP.CV.WT,YFP.FANO.WT,YFP.CV.TDH3,YFP.FANO.TDH3,YFP.SD.WT.REL,YFP.CV.WT.REL,YFP.FANO.WT.REL,YFP.SD.TDH3.REL,YFP.CV.TDH3.REL,YFP.FANO.TDH3.REL,FSC.CORRECT) ~ STRAIN + CONDITION + ASSAY + PLATE + CTRL.GENO, data=CLEAN.DATA, FUN = mean)
RNR2.SD <- aggregate(cbind(YFP.MEDIAN.ABS,YFP.SD.ABS,YFP.MAD.ABS,YFP.MEDIAN.WT,YFP.SD.WT,YFP.MAD.WT,YFP.MEDIAN.TDH3,YFP.SD.TDH3,YFP.MAD.TDH3,YFP.CV.WT,YFP.FANO.WT,YFP.CV.TDH3,YFP.FANO.TDH3,YFP.SD.WT.REL,YFP.CV.WT.REL,YFP.FANO.WT.REL,YFP.SD.TDH3.REL,YFP.CV.TDH3.REL,YFP.FANO.TDH3.REL,FSC.CORRECT) ~ STRAIN + CONDITION + ASSAY + PLATE + CTRL.GENO, data=CLEAN.DATA, FUN = sd)
RNR2.N <- aggregate(cbind(YFP.MEDIAN.ABS,YFP.SD.ABS,YFP.MAD.ABS,YFP.MEDIAN.WT,YFP.SD.WT,YFP.MAD.WT,YFP.MEDIAN.TDH3,YFP.SD.TDH3,YFP.MAD.TDH3,YFP.CV.WT,YFP.FANO.WT,YFP.CV.TDH3,YFP.FANO.TDH3,YFP.SD.WT.REL,YFP.CV.WT.REL,YFP.FANO.WT.REL,YFP.SD.TDH3.REL,YFP.CV.TDH3.REL,YFP.FANO.TDH3.REL,FSC.CORRECT) ~ STRAIN + CONDITION + ASSAY + PLATE + CTRL.GENO, data=CLEAN.DATA, FUN = length)

RNR2.MEAN  <- RNR2.MEAN[which(RNR2.N[,6] >= 3),]
RNR2.SD <- RNR2.SD[which(RNR2.N[,6] >= 3),]
RNR2.N <- RNR2.N[which(RNR2.N[,6] >= 3),]

RNR2 <- cbind.data.frame(RNR2.MEAN,RNR2.SD[,6:25],RNR2.N[,6])

colnames(RNR2) <- c("STRAIN","CONDITION","ASSAY","PLATE","CTRL.GENO",paste0("MEAN.",colnames(RNR2.MEAN[,6:25])),paste0("SD.",colnames(RNR2.SD[,6:25])),"N")

write.table(RNR2,"RNR2.SUMMARY.DATA_DCv34.txt",sep="\t",quote=FALSE,row.names=FALSE)

`%notin%` <- function(x,y) !(x %in% y)

CONTAM.CHECK<-droplevels(subset(CLEAN.DATA,CLEAN.DATA$CONTAM.CHECK!="NONE"))

if(all(CONTAM.CHECK$STRAIN %notin% RNR2$STRAIN)) {
  #All datapoints flagged as suspicious have been removed by outlier removal filters.
  print("Yay! No suspicious values detected. Proceed.")
}

if(any(CONTAM.CHECK$STRAIN %in% RNR2$STRAIN)) {
  CONTAM.CHECK<-CONTAM.CHECK[CONTAM.CHECK$STRAIN %in% CONTAM.CHECK$STRAIN[which(CONTAM.CHECK$STRAIN %in% RNR2$STRAIN)],]
  write.table(CONTAM.CHECK,"RNR2.CONTAM.CHECK_outliersexcl.csv",sep=",",quote=FALSE,row.names=FALSE)  
}


###############################
#HIERARCHICAL PERMUTATION TEST#
###############################

#Clear memory
rm(list=ls())
options(warn=-1)

library(permute)
library(Hmisc)

N.PERM <- 10000

#Permutation function

SHUFFLE <- function(STRAIN.LIST = STRAIN.LIST[[i]],WT.LIST = WT.LIST) {
  SAMPLE <- c(STRAIN.LIST,WT.LIST[[sample(1:length(WT.LIST),1)]])
  OUTPUT <- sample(SAMPLE,length(SAMPLE))
  DIST.SIM <- abs(mean(OUTPUT[1:length(STRAIN.LIST)]) - mean(OUTPUT[(length(STRAIN.LIST)+1):length(SAMPLE)]))
  DIST.OBS <- abs(mean(SAMPLE[1:length(STRAIN.LIST)]) - mean(SAMPLE[(length(STRAIN.LIST)+1):length(SAMPLE)]))
  
  DIFF <- DIST.OBS - DIST.SIM
  
  return(DIFF)
}

parent.dir <- PATH
setwd(parent.dir)

####RNR2 Analysis date 10112016#########################################################################

ALL.REP <- read.csv("RNR2.CLEAN.ADJUST_DCv34.csv",header=TRUE)
ALL.REP <- subset(ALL.REP, MEDIAN.OUTLIER == "NO" & SD.OUTLIER == "NO" & FSC.OUTLIER == "NO")

SUMMARY <- read.table("RNR2.SUMMARY.DATA_DCv34.txt",header=TRUE)

#SUMMARY <- droplevels(subset(SUMMARY, STRAIN != "1139"))

WT <- subset(ALL.REP, CONDITION == "SHAM")
LOW.FSC <- median(WT$FSC.CORRECT) - 4*mad(WT$FSC.CORRECT)
HIGH.FSC <- median(WT$FSC.CORRECT) + 4*mad(WT$FSC.CORRECT)
LOW.FLUO <- median(WT$YFP.MEDIAN.CORRECT) - 4*mad(WT$YFP.MEDIAN.CORRECT)
HIGH.FLUO <- median(WT$YFP.MEDIAN.CORRECT) + 4*mad(WT$YFP.MEDIAN.CORRECT)
WT <- subset(WT, FSC.CORRECT > LOW.FSC & FSC.CORRECT < HIGH.FSC & YFP.MEDIAN.CORRECT > LOW.FLUO & YFP.MEDIAN.CORRECT < HIGH.FLUO)

#MEDIAN

#Create list of WT phenotype grouped by strain

N.WT <- length(unique(WT$STRAIN))
WT.LIST <- vector('list',N.WT)
for (i in 1:N.WT)
{
  CUR.WT <- subset(WT, STRAIN == unique(WT$STRAIN)[i])
  WT.LIST[[i]] <- CUR.WT$YFP.MEDIAN.CORRECT 
}


#Create list of all strain phenotypes grouped by strain

SUMMARY <- SUMMARY[order(SUMMARY$STRAIN),]
ALL.REP <- droplevels(subset(ALL.REP,STRAIN %in% as.character(SUMMARY$STRAIN)))
ALL.REP <- ALL.REP[order(ALL.REP$STRAIN),]
N.STRAIN <- length(unique(ALL.REP$STRAIN))
STRAIN.LIST <- vector('list',N.STRAIN)
for (i in 1:N.STRAIN)
{
  CUR.STRAIN <- subset(ALL.REP, STRAIN == unique(ALL.REP$STRAIN)[i])
  STRAIN.LIST[[i]] <- CUR.STRAIN$YFP.MEDIAN.CORRECT
}


for (i in 1:length(STRAIN.LIST))
{
  DIFF <- sapply(1:N.PERM,FUN=function(x)	SHUFFLE(STRAIN.LIST = STRAIN.LIST[[i]],WT.LIST = WT.LIST))
  P.VAL <- (length(which(DIFF < 0)) + 1) / (length(which(DIFF < 0)) + length(which(DIFF > 0)) + 1) 	
  SUMMARY[i,"P.VAL.MEDIAN"] <- P.VAL
  print(round(i/length(STRAIN.LIST)*100,1))
}

#NOISE: CV

#Create list of WT phenotype grouped by strain

N.WT <- length(unique(WT$STRAIN))
WT.LIST <- vector('list',N.WT)
for (i in 1:N.WT)
{
  CUR.WT <- subset(WT, STRAIN == unique(WT$STRAIN)[i])
  WT.LIST[[i]] <- CUR.WT$YFP.CV.WT
}


#Create list of all strain phenotypes grouped by strain

SUMMARY <- SUMMARY[order(SUMMARY$STRAIN),]
ALL.REP <- droplevels(subset(ALL.REP,STRAIN %in% as.character(SUMMARY$STRAIN)))
ALL.REP <- ALL.REP[order(ALL.REP$STRAIN),]
N.STRAIN <- length(unique(ALL.REP$STRAIN))
STRAIN.LIST <- vector('list',N.STRAIN)
for (i in 1:N.STRAIN)
{
  CUR.STRAIN <- subset(ALL.REP, STRAIN == unique(ALL.REP$STRAIN)[i])
  STRAIN.LIST[[i]] <- CUR.STRAIN$YFP.CV.WT
}


for (i in 1:length(STRAIN.LIST))
{
  DIFF <- sapply(1:N.PERM,FUN=function(x)	SHUFFLE(STRAIN.LIST = STRAIN.LIST[[i]],WT.LIST = WT.LIST))
  P.VAL <- (length(which(DIFF < 0))+1) / (length(which(DIFF < 0)) + length(which(DIFF > 0)) + 1) 	
  SUMMARY[i,"P.VAL.CV.WT"] <- P.VAL
  print(round(i/length(STRAIN.LIST)*100,1))
}

#NOISE: FANO

#Create list of WT phenotype grouped by strain

N.WT <- length(unique(WT$STRAIN))
WT.LIST <- vector('list',N.WT)
for (i in 1:N.WT)
{
  CUR.WT <- subset(WT, STRAIN == unique(WT$STRAIN)[i])
  WT.LIST[[i]] <- CUR.WT$YFP.FANO.WT
}


#Create list of all strain phenotypes grouped by strain

SUMMARY <- SUMMARY[order(SUMMARY$STRAIN),]
ALL.REP <- droplevels(subset(ALL.REP,STRAIN %in% as.character(SUMMARY$STRAIN)))
ALL.REP <- ALL.REP[order(ALL.REP$STRAIN),]
N.STRAIN <- length(unique(ALL.REP$STRAIN))
STRAIN.LIST <- vector('list',N.STRAIN)
for (i in 1:N.STRAIN)
{
  CUR.STRAIN <- subset(ALL.REP, STRAIN == unique(ALL.REP$STRAIN)[i])
  STRAIN.LIST[[i]] <- CUR.STRAIN$YFP.FANO.WT
}


for (i in 1:length(STRAIN.LIST))
{
  DIFF <- sapply(1:N.PERM,FUN=function(x)	SHUFFLE(STRAIN.LIST = STRAIN.LIST[[i]],WT.LIST = WT.LIST))
  P.VAL <- (length(which(DIFF < 0))+1) / (length(which(DIFF < 0)) + length(which(DIFF > 0)) + 1) 	
  SUMMARY[i,"P.VAL.FANO.WT"] <- P.VAL
  print(round(i/length(STRAIN.LIST)*100,1))
}

#FSC MEDIAN

#Create list of WT phenotype grouped by strain

N.WT <- length(unique(WT$STRAIN))
WT.LIST <- vector('list',N.WT)
for (i in 1:N.WT)
{
  CUR.WT <- subset(WT, STRAIN == unique(WT$STRAIN)[i])
  WT.LIST[[i]] <- CUR.WT$FSC.CORRECT
}


#Create list of all strain phenotypes grouped by strain

SUMMARY <- SUMMARY[order(SUMMARY$STRAIN),]
ALL.REP <- droplevels(subset(ALL.REP,STRAIN %in% as.character(SUMMARY$STRAIN)))
ALL.REP <- ALL.REP[order(ALL.REP$STRAIN),]
N.STRAIN <- length(unique(ALL.REP$STRAIN))
STRAIN.LIST <- vector('list',N.STRAIN)
for (i in 1:N.STRAIN)
{
  CUR.STRAIN <- subset(ALL.REP, STRAIN == unique(ALL.REP$STRAIN)[i])
  STRAIN.LIST[[i]] <- CUR.STRAIN$FSC.CORRECT
}


for (i in 1:length(STRAIN.LIST))
{
  DIFF <- sapply(1:N.PERM,FUN=function(x)	SHUFFLE(STRAIN.LIST = STRAIN.LIST[[i]],WT.LIST = WT.LIST))
  P.VAL <- (length(which(DIFF < 0))+1) / (length(which(DIFF < 0)) + length(which(DIFF > 0)) + 1) 	
  SUMMARY[i,"P.VAL.FSC"] <- P.VAL
  print(round(i/length(STRAIN.LIST)*100,1))
}

write.table(SUMMARY,"RNR2.SUMMARY.DCv34.Pvals.txt",sep="\t",quote=FALSE,row.names=FALSE)

#________________________________________________________________________________________________

######################
#    STM1
######################

#Clear memory
rm(list=ls())
options(warn=-1)


#####################
#1-LOADING LIBRARIES#
#####################

library(flowCore)
library(flowClust)
library(flowViz)
library(plotrix)
library(nlme)
library(MethComp)
library(outliers)
library(pcaPP)

library(reshape2)
library(MASS)
library(ggplot2)
library(Hmisc)
library(fBasics)
library(lawstat)
library(fitdistrplus)
library(mixtools)
library(vioplot)
library(gplots)
library(RColorBrewer)
library(calibrate)
library(e1071)

box <- graphics::box


########################################
#  CLEANING
########################################

#Clear memory
rm(list=ls())
options(warn=-1)

#Set working directory
parent.dir <- PATH
setwd(parent.dir)

#Load experiment setup
SETUP <- read.csv("Template_STM1_10102016.csv",header=TRUE,as.is=TRUE)
TESTNAMES <- list.files("PATH/STM1",pattern=".fcs",full.name=TRUE,recursive=TRUE,include.dirs=TRUE)
SETUP<-subset(SETUP, SETUP$FILENAMES %in% TESTNAMES, drop = TRUE)

SETUP[,"COUNTER"] <- c(1:nrow(SETUP))

#Determine Hard Gates
GATES <- GATE.CALIB(SETUP$FILENAMES[1])

#Clean Data
Output <- apply(SETUP,1,CLEANING,GATES=GATES)
OUTPUT <- as.data.frame(Output[[1]])
OUTPUT[2:nrow(SETUP),1:ncol(OUTPUT)] <- NA
for (i in 2:nrow(SETUP))
{
  if (is.null(Output[[i]]))
  {} 
  else {
    OUTPUT[i,] <- Output[[i]]
  }
}

write.table(OUTPUT,"STM1_Experiment.Output_10112016_DCv34.txt",row.names=FALSE,sep="\t")

CLEAN <- cbind.data.frame(SETUP[,1:(ncol(SETUP)-2)],OUTPUT)

write.table(CLEAN,"STM1_Clean.Data_10112016_DCv34.txt",row.names=FALSE,sep="\t")

################################################################
#     PLATE CORRECTION & OUTLIER REMOVAL
################################################################

#Clear memory
rm(list=ls())
options(warn=-1)

#####################
#1-LOADING LIBRARIES#
#####################
library(flowCore)
library(flowClust)
library(flowViz)
library(pcaPP)
library(mixtools)
library(plyr)
library(MASS)
box <- graphics::box
options(warn=-1)

####################################
#2- Quality Control and Corrections#
####################################

parent.dir <- PATH
setwd(parent.dir)

DATA.TYPE <- c(
  "factor","factor","factor","factor","factor","factor","factor","factor","factor","factor","factor","factor","factor","factor","factor",
  "integer","integer","integer","integer",
  "numeric","numeric","numeric","numeric","numeric","numeric","numeric","numeric","numeric","numeric","numeric","numeric","numeric","numeric","numeric","numeric","numeric","numeric","numeric","numeric","numeric","numeric","numeric","numeric","numeric",
  "integer")

###Read in data sets
CLEAN.DATA <- read.table("STM1_Clean.Data_10112016_DCv34.txt",colClasses=DATA.TYPE,header=TRUE, sep="\t")

###Remove samples with NA
CLEAN.DATA <- CLEAN.DATA[complete.cases(CLEAN.DATA),]

##Translate into log scale for appropriate correction
CLEAN.DATA[,"log.RNA.MEDIAN"] <- log((CLEAN.DATA[,"YFP.MEDIAN.FINAL"]) + 0.05)
CLEAN.DATA[,"log.RNA.MAD"] <- CLEAN.DATA[,"YFP.MAD.FINAL"] / (CLEAN.DATA[,"YFP.MEDIAN.FINAL"] + 0.05)
CLEAN.DATA[,"log.RNA.SD"] <- CLEAN.DATA[,"YFP.SD.FINAL"] / (CLEAN.DATA[,"YFP.MEDIAN.FINAL"] + 0.05)

###Remove samples with less than 1000 events
CLEAN.DATA <- subset(CLEAN.DATA,CLEAN.DATA$COUNTS.FINAL > 1000)

####Add variable to ID whether control is own promoter or TDH3
CLEAN.DATA[,"CTRL.GENO"]<-as.factor("STM1")

###Visualize spread of data

CLEAN.DATA$CONTAM.CHECK<-"NONE"
CLEAN.DATA$CONTAM.CHECK[(CLEAN.DATA$CONDITION=="EMS"|CLEAN.DATA$CONDITION=="SHAM"|CLEAN.DATA$CONDITION=="EMS.LOW"|CLEAN.DATA$CONDITION=="EMS.HIGH") & (CLEAN.DATA$YFP.MEDIAN.FINAL<.53)]<-"BACT.CONTAM.CHECK"
CLEAN.DATA$CONTAM.CHECK[(CLEAN.DATA$CONDITION=="EMS"|CLEAN.DATA$CONDITION=="SHAM"|CLEAN.DATA$CONDITION=="EMS.LOW"|CLEAN.DATA$CONDITION=="EMS.HIGH") & (CLEAN.DATA$YFP.MEDIAN.FINAL>0.7)]<-"TDH3.CONTAM.CHECK"
CLEAN.DATA$CONTAM.CHECK[(CLEAN.DATA$CONDITION=="TDH3.SHAM" & CLEAN.DATA$YFP.MEDIAN.FINAL<0.7)]<-"EXP.SHAM.CONTAM.CHECK"
CLEAN.DATA$CONTAM.CHECK[(CLEAN.DATA$CONDITION=="NULL" & CLEAN.DATA$YFP.MEDIAN.FINAL>0.53)]<-"EXP.NULL.CONTAM.CHECK"


###Seperate controls and remove FSC outliers
CONTROL <- subset(CLEAN.DATA,CLEAN.DATA$CONDITION == "CTRL")
FSC.CORRECT <- lm(CONTROL$FSC.MEDIAN.FINAL ~ 0 + CONTROL$FLOW.RUN)
RESID <- FSC.CORRECT$resid
CLEAN.DATA <- droplevels(CLEAN.DATA[-which(CLEAN.DATA$CONDITION == "CTRL")[abs(RESID) > 3*mad(RESID)],])
CONTROL <- droplevels(subset(CLEAN.DATA,CLEAN.DATA$CONDITION == "CTRL"))

INITIAL.CORRECT <- lm(CONTROL$log.RNA.MEDIAN ~ 0 + CONTROL$FLOW.RUN)
RESID <- INITIAL.CORRECT$resid
CLEAN.DATA<- droplevels(CLEAN.DATA[-which(CLEAN.DATA$CONDITION == "CTRL")[abs(RESID) > 3*mad(RESID)],])
CONTROL <- droplevels(subset(CLEAN.DATA,CLEAN.DATA$CONDITION == "CTRL"))

`%notin%` <- function(x,y) !(x %in% y)
RESULT<-count(CONTROL$FLOW.RUN)
RESULT.MUT<-count(CLEAN.DATA$FLOW.RUN)
EXCLUDE1<-(RESULT.MUT[which(RESULT.MUT[,1] %notin% RESULT[,1]),1])
EXCLUDE2<-(RESULT[which(RESULT[,2] <5),1])
CLEAN.DATA<-droplevels(CLEAN.DATA[CLEAN.DATA$FLOW.RUN %notin% EXCLUDE1,])
CLEAN.DATA<-droplevels(CLEAN.DATA[CLEAN.DATA$FLOW.RUN %notin% EXCLUDE2,])
CONTROL <- subset(CLEAN.DATA,CLEAN.DATA$CONDITION == "CTRL")


###Corrections for YFP median and MAD
#Correct Median
#fit<- lm(log.RNA.MEDIAN~FLOW.RUN+REP+STACK.ORDER+STACK.ORDER:FLOW.RUN+REP+ROW+COLUMN+BLOCK+BLOCK:ROW+ROW:COLUMN+STACK.ORDER:BLOCK,data=CONTROL)
#step <- stepAIC(fit, direction="both")
log.RNA.MEDIAN.CORRECT <- lm(log.RNA.MEDIAN ~ FLOW.RUN + COLUMN + ROW, data=CONTROL)
CLEAN.DATA[,"log.RNA.MEDIAN.CORRECT"] <- CLEAN.DATA$log.RNA.MEDIAN - predict(log.RNA.MEDIAN.CORRECT, CLEAN.DATA) + mean(CONTROL$log.RNA.MEDIAN)
CONTROL <- subset(CLEAN.DATA,CLEAN.DATA$CONDITION == "CTRL")

#Correct sd
#fit<- lm(log.RNA.SD~FLOW.RUN+REP+STACK.ORDER+STACK.ORDER:FLOW.RUN+REP+ROW+COLUMN+BLOCK+BLOCK:ROW+ROW:COLUMN+STACK.ORDER:BLOCK,data=CONTROL)
#step <- step(fit, direction="both")
log.RNA.SD.CORRECT <- lm(log.RNA.SD ~  FLOW.RUN + ROW + COLUMN,data=CONTROL)
CLEAN.DATA[,"log.RNA.SD.CORRECT"] <- CLEAN.DATA$log.RNA.SD - predict(log.RNA.SD.CORRECT , CLEAN.DATA) + mean(CONTROL$log.RNA.SD)
CONTROL <- subset(CLEAN.DATA,CLEAN.DATA$CONDITION == "CTRL")

#Correct mad
#fit<- lm(log.RNA.MAD~FLOW.RUN+REP+STACK.ORDER+STACK.ORDER:FLOW.RUN+REP+ROW+COLUMN+BLOCK+BLOCK:ROW+ROW:COLUMN+STACK.ORDER:BLOCK,data=CONTROL)
#step <- step(fit, direction="both")
log.RNA.MAD.CORRECT <- lm(log.RNA.MAD ~  FLOW.RUN + ROW + COLUMN+ ROW:COLUMN,data=CONTROL)
CLEAN.DATA[,"log.RNA.MAD.CORRECT"] <- CLEAN.DATA$log.RNA.MAD - predict(log.RNA.MAD.CORRECT , CLEAN.DATA) + mean(CONTROL$log.RNA.MAD)
CONTROL <- subset(CLEAN.DATA,CLEAN.DATA$CONDITION == "CTRL")

#Correct FSC
#fit<- lm(FSC.MEDIAN.FINAL~FLOW.RUN+REP+STACK.ORDER+STACK.ORDER:FLOW.RUN+REP+ROW+COLUMN+BLOCK+BLOCK:ROW+ROW:COLUMN+STACK.ORDER:BLOCK,data=CONTROL)
#step <- stepAIC(fit, direction="both")
FSC.CORRECT <- lm(FSC.MEDIAN.FINAL ~  FLOW.RUN + ROW + COLUMN,data=CONTROL)
CLEAN.DATA[,"FSC.CORRECT"] <- CLEAN.DATA$FSC.MEDIAN.FINAL - predict(FSC.CORRECT, CLEAN.DATA) + mean(CONTROL$FSC.MEDIAN.FINAL)
CONTROL <- subset(CLEAN.DATA,CLEAN.DATA$CONDITION == "CTRL")


#Back translate to linear scale
CLEAN.DATA[,"YFP.MEDIAN.CORRECT"] <- (exp(CLEAN.DATA[,"log.RNA.MEDIAN.CORRECT"]) - 0.05)
CLEAN.DATA[,"YFP.SD.CORRECT"] <- CLEAN.DATA[,"log.RNA.SD.CORRECT"] * CLEAN.DATA[,"YFP.MEDIAN.CORRECT"]
CLEAN.DATA[,"YFP.MAD.CORRECT"] <- CLEAN.DATA[,"log.RNA.MAD.CORRECT"] * CLEAN.DATA[,"YFP.MEDIAN.CORRECT"]


#PERFORM FABIEN'S OUTLIER CORRECTION FILTERING OUT ANY REPLICATE OUTSIDE OF 4*MAD

levels(CLEAN.DATA$STRAIN)[levels(CLEAN.DATA$STRAIN)=="SHAM"]<-paste("SHAM.",CLEAN.DATA[CLEAN.DATA$STRAIN=="SHAM","PLATE"],".",CLEAN.DATA[CLEAN.DATA$STRAIN=="SHAM","POSITION"],sep="")

#Remove outliers
for (i in 1:nrow(CLEAN.DATA))
{
  CUR <- subset(CLEAN.DATA, STRAIN == CLEAN.DATA[i,"STRAIN"])
  if (CLEAN.DATA[i,"YFP.MEDIAN.CORRECT"] > median(CUR$YFP.MEDIAN.CORRECT) + 4*mad(CUR$YFP.MEDIAN.CORRECT) | CLEAN.DATA[i,"YFP.MEDIAN.CORRECT"] < median(CUR$YFP.MEDIAN.CORRECT) - 4*mad(CUR$YFP.MEDIAN.CORRECT)) {
    CLEAN.DATA[i,"MEDIAN.OUTLIER"] <- "YES"
  } else {
    CLEAN.DATA[i,"MEDIAN.OUTLIER"] <- "NO"
  }
  if (CLEAN.DATA[i,"YFP.SD.CORRECT"] > median(CUR$YFP.SD.CORRECT) + 4*mad(CUR$YFP.SD.CORRECT) | CLEAN.DATA[i,"YFP.SD.CORRECT"] < median(CUR$YFP.SD.CORRECT) - 4*mad(CUR$YFP.SD.CORRECT)) {
    CLEAN.DATA[i,"SD.OUTLIER"] <- "YES"
  } else {
    CLEAN.DATA[i,"SD.OUTLIER"] <- "NO"
  }
  if (CLEAN.DATA[i,"FSC.CORRECT"] > median(CUR$FSC.CORRECT) + 4*mad(CUR$FSC.CORRECT) | CLEAN.DATA[i,"FSC.CORRECT"] < median(CUR$FSC.CORRECT) - 4*mad(CUR$FSC.CORRECT)) {
    CLEAN.DATA[i,"FSC.OUTLIER"] <- "YES"
  } else {
    CLEAN.DATA[i,"FSC.OUTLIER"] <- "NO"
  }	
}

CLEAN.DATA <- subset(CLEAN.DATA, MEDIAN.OUTLIER == "NO" & SD.OUTLIER == "NO" & FSC.OUTLIER == "NO")

#Write processed data to file
write.table(CLEAN.DATA,"STM1.CLEAN.CORRECT_10112016_DCv34.csv",sep=",",quote=FALSE,row.names=FALSE)

#####################
#5 - Control plots  #
#####################

CONTROL <- droplevels(subset(CLEAN.DATA,CLEAN.DATA$CONDITION == "CTRL"))
SHAM <- droplevels(subset(CLEAN.DATA,CLEAN.DATA$CONDITION == "SHAM"))
TDH3.SHAM <- droplevels(subset(CLEAN.DATA,CLEAN.DATA$CONDITION == "TDH3.SHAM"))


###############################################################################
#
#         Strain Estimates
#
###############################################################################

# Clear memory

rm(list=ls())
options(warn=-1)

setwd("E:/wittkopp data/processed data/RNA scale analyses/DCv34")

#STM1
CLEAN.DATA <- read.csv("STM1.CLEAN.CORRECT_10112016_DCv34.csv", header=TRUE)

#Calculate mean and noise for each sample relative to controls
TDH3.SHAM <- subset(CLEAN.DATA, CLEAN.DATA$CONDITION == "TDH3.SHAM")
NEG <- subset(CLEAN.DATA, CLEAN.DATA$CONDITION == "NULL")
WT.SHAM <- subset(CLEAN.DATA, CLEAN.DATA$CONDITION == "SHAM")

#ABSOLUTE EXPRESSION SCALE
CLEAN.DATA[,"YFP.MEDIAN.ABS"] <- (CLEAN.DATA$YFP.MEDIAN.CORRECT - median(NEG$YFP.MEDIAN.CORRECT))
CLEAN.DATA[,"YFP.SD.ABS"]   <- CLEAN.DATA$YFP.SD.CORRECT - median(NEG$YFP.SD.CORRECT)
CLEAN.DATA[,"YFP.MAD.ABS"]   <- CLEAN.DATA$YFP.MAD.CORRECT - median(NEG$YFP.SD.CORRECT)

#Re-calculate mean and noise for each sample relative to controls
TDH3.SHAM <- subset(CLEAN.DATA, CLEAN.DATA$CONDITION == "TDH3.SHAM")
NEG <- subset(CLEAN.DATA, CLEAN.DATA$CONDITION == "NULL")
WT.SHAM <- subset(CLEAN.DATA, CLEAN.DATA$CONDITION == "SHAM")

#PROMOTER-SPECIFIC SCALING
CLEAN.DATA[,"YFP.MEDIAN.WT"] <- CLEAN.DATA$YFP.MEDIAN.ABS/median(WT.SHAM$YFP.MEDIAN.ABS)
CLEAN.DATA[,"YFP.SD.WT"]   <- CLEAN.DATA$YFP.SD.ABS/median(WT.SHAM$YFP.MEDIAN.ABS)
CLEAN.DATA[,"YFP.MAD.WT"]   <- CLEAN.DATA$YFP.MAD.ABS/median(WT.SHAM$YFP.MEDIAN.ABS)

#RELATIVE TO TDH3 EXPRESSION
CLEAN.DATA[,"YFP.MEDIAN.TDH3"] <- CLEAN.DATA$YFP.MEDIAN.ABS/median(TDH3.SHAM$YFP.MEDIAN.ABS)
CLEAN.DATA[,"YFP.SD.TDH3"]   <- CLEAN.DATA$YFP.SD.ABS/median(TDH3.SHAM$YFP.MEDIAN.ABS)
CLEAN.DATA[,"YFP.MAD.TDH3"]   <- CLEAN.DATA$YFP.MAD.ABS/median(TDH3.SHAM$YFP.MEDIAN.ABS)

#Re-calculate mean and noise for each sample relative to controls
TDH3.SHAM <- subset(CLEAN.DATA, CLEAN.DATA$CONDITION == "TDH3.SHAM")
TDH3.SHAM<-subset(TDH3.SHAM,TDH3.SHAM$YFP.MEDIAN.FINAL>0.7)
NEG <- subset(CLEAN.DATA, CLEAN.DATA$CONDITION == "NULL")
WT.SHAM <- subset(CLEAN.DATA, CLEAN.DATA$CONDITION == "SHAM")

#CALCULATE NOISE METRICS
CLEAN.DATA[,"YFP.CV.WT"]   <- CLEAN.DATA$YFP.SD.WT/CLEAN.DATA$YFP.MEDIAN.WT
CLEAN.DATA[,"YFP.CV.TDH3"]   <- CLEAN.DATA$YFP.SD.TDH3/CLEAN.DATA$YFP.MEDIAN.TDH3
CLEAN.DATA[,"YFP.FANO.WT"]   <- CLEAN.DATA$YFP.SD.WT^2/CLEAN.DATA$YFP.MEDIAN.WT
CLEAN.DATA[,"YFP.FANO.TDH3"]   <- CLEAN.DATA$YFP.SD.TDH3^2/CLEAN.DATA$YFP.MEDIAN.TDH3


#Re-calculate mean and noise for each sample relative to controls
TDH3.SHAM <- subset(CLEAN.DATA, CLEAN.DATA$CONDITION == "TDH3.SHAM")
TDH3.SHAM<-subset(TDH3.SHAM,TDH3.SHAM$YFP.MEDIAN.FINAL>0.7)
NEG <- subset(CLEAN.DATA, CLEAN.DATA$CONDITION == "NULL")
WT.SHAM <- subset(CLEAN.DATA, CLEAN.DATA$CONDITION == "SHAM")

#CALCULATE RELATIVE NOISE METRICS
CLEAN.DATA[,"YFP.SD.WT.REL"]   <- CLEAN.DATA$YFP.SD.WT/median(WT.SHAM$YFP.SD.WT)
CLEAN.DATA[,"YFP.SD.TDH3.REL"]   <- CLEAN.DATA$YFP.SD.TDH3/median(WT.SHAM$YFP.SD.TDH3)
CLEAN.DATA[,"YFP.CV.WT.REL"]   <- CLEAN.DATA$YFP.SD.WT.REL/CLEAN.DATA$YFP.MEDIAN.WT
CLEAN.DATA[,"YFP.CV.TDH3.REL"]   <- CLEAN.DATA$YFP.SD.TDH3.REL/CLEAN.DATA$YFP.MEDIAN.TDH3
CLEAN.DATA[,"YFP.FANO.WT.REL"]   <- CLEAN.DATA$YFP.SD.WT.REL^2/CLEAN.DATA$YFP.MEDIAN.WT
CLEAN.DATA[,"YFP.FANO.TDH3.REL"]   <- CLEAN.DATA$YFP.SD.TDH3.REL^2/CLEAN.DATA$YFP.MEDIAN.TDH3


#Add NA for samples where CV or FANO are undefined.
for (i in 1:nrow(CLEAN.DATA))
{
  if (CLEAN.DATA[i,"YFP.MEDIAN.ABS"] < 0 | CLEAN.DATA[i,"YFP.SD.ABS"] < 0) {
    CLEAN.DATA[i,"YFP.MAD.WT"] <- NA
    CLEAN.DATA[i,"YFP.MAD.TDH3"] <- NA
    CLEAN.DATA[i,"YFP.CV.WT"] <- NA
    CLEAN.DATA[i,"YFP.CV.TDH3"] <- NA
    CLEAN.DATA[i,"YFP.CV.WT.REL"] <- NA
    CLEAN.DATA[i,"YFP.CV.TDH3.REL"] <- NA
    CLEAN.DATA[i,"YFP.FANO.WT"] <- NA
    CLEAN.DATA[i,"YFP.FANO.TDH3"] <- NA
    CLEAN.DATA[i,"YFP.FANO.WT.REL"] <- NA
    CLEAN.DATA[i,"YFP.FANO.TDH3.REL"] <- NA
    
  }
}

CLEAN.DATA <- droplevels(subset(CLEAN.DATA,CLEAN.DATA$CONDITION!="CTRL"))

write.table(CLEAN.DATA,"STM1.CLEAN.ADJUST_10112016_DCv34.csv",sep=",",quote=FALSE,row.names=FALSE)

CLEAN.DATA <- read.csv("STM1.CLEAN.ADJUST_10112016_DCv34.csv", header=TRUE)

STM1.MEAN <- aggregate(cbind(YFP.MEDIAN.ABS,YFP.SD.ABS,YFP.MAD.ABS,YFP.MEDIAN.WT,YFP.SD.WT,YFP.MAD.WT,YFP.MEDIAN.TDH3,YFP.SD.TDH3,YFP.MAD.TDH3,YFP.CV.WT,YFP.FANO.WT,YFP.CV.TDH3,YFP.FANO.TDH3,YFP.SD.WT.REL,YFP.CV.WT.REL,YFP.FANO.WT.REL,YFP.SD.TDH3.REL,YFP.CV.TDH3.REL,YFP.FANO.TDH3.REL,FSC.CORRECT) ~ STRAIN + CONDITION + ASSAY + PLATE + CTRL.GENO, data=CLEAN.DATA, FUN = mean)
STM1.SD <- aggregate(cbind(YFP.MEDIAN.ABS,YFP.SD.ABS,YFP.MAD.ABS,YFP.MEDIAN.WT,YFP.SD.WT,YFP.MAD.WT,YFP.MEDIAN.TDH3,YFP.SD.TDH3,YFP.MAD.TDH3,YFP.CV.WT,YFP.FANO.WT,YFP.CV.TDH3,YFP.FANO.TDH3,YFP.SD.WT.REL,YFP.CV.WT.REL,YFP.FANO.WT.REL,YFP.SD.TDH3.REL,YFP.CV.TDH3.REL,YFP.FANO.TDH3.REL,FSC.CORRECT) ~ STRAIN + CONDITION + ASSAY + PLATE + CTRL.GENO, data=CLEAN.DATA, FUN = sd)
STM1.N <- aggregate(cbind(YFP.MEDIAN.ABS,YFP.SD.ABS,YFP.MAD.ABS,YFP.MEDIAN.WT,YFP.SD.WT,YFP.MAD.WT,YFP.MEDIAN.TDH3,YFP.SD.TDH3,YFP.MAD.TDH3,YFP.CV.WT,YFP.FANO.WT,YFP.CV.TDH3,YFP.FANO.TDH3,YFP.SD.WT.REL,YFP.CV.WT.REL,YFP.FANO.WT.REL,YFP.SD.TDH3.REL,YFP.CV.TDH3.REL,YFP.FANO.TDH3.REL,FSC.CORRECT) ~ STRAIN + CONDITION + ASSAY + PLATE + CTRL.GENO, data=CLEAN.DATA, FUN = length)

STM1.MEAN  <- STM1.MEAN[which(STM1.N[,6] >= 3),]
STM1.SD <- STM1.SD[which(STM1.N[,6] >= 3),]
STM1.N <- STM1.N[which(STM1.N[,6] >= 3),]

STM1 <- cbind.data.frame(STM1.MEAN,STM1.SD[,6:25],STM1.N[,6])

colnames(STM1) <- c("STRAIN","CONDITION","ASSAY","PLATE","CTRL.GENO",paste0("MEAN.",colnames(STM1.MEAN[,6:25])),paste0("SD.",colnames(STM1.SD[,6:25])),"N")

write.table(STM1,"STM1.SUMMARY.DATA_10112016_DCv34.txt",sep="\t",quote=FALSE,row.names=FALSE)

`%notin%` <- function(x,y) !(x %in% y)

CONTAM.CHECK<-droplevels(subset(CLEAN.DATA,CLEAN.DATA$CONTAM.CHECK!="NONE"))

if(all(CONTAM.CHECK$STRAIN %notin% STM1$STRAIN)) {
  #All datapoints flagged as suspicious have been removed by outlier removal filters.
  print("Yay! No suspicious values detected. Proceed.")
}

if(any(CONTAM.CHECK$STRAIN %in% STM1$STRAIN)) {
  CONTAM.CHECK<-CONTAM.CHECK[CONTAM.CHECK$STRAIN %in% CONTAM.CHECK$STRAIN[which(CONTAM.CHECK$STRAIN %in% STM1$STRAIN)],]
  write.table(CONTAM.CHECK,"STM1.CONTAM.CHECK_outliersexcl.csv",sep=",",quote=FALSE,row.names=FALSE)  
}


###############################
#HIERARCHICAL PERMUTATION TEST#
###############################

#Clear memory
rm(list=ls())
options(warn=-1)

library(permute)
library(Hmisc)

N.PERM <- 10000

#Permutation function

SHUFFLE <- function(STRAIN.LIST = STRAIN.LIST[[i]],WT.LIST = WT.LIST) {
  SAMPLE <- c(STRAIN.LIST,WT.LIST[[sample(1:length(WT.LIST),1)]])
  OUTPUT <- sample(SAMPLE,length(SAMPLE))
  DIST.SIM <- abs(mean(OUTPUT[1:length(STRAIN.LIST)]) - mean(OUTPUT[(length(STRAIN.LIST)+1):length(SAMPLE)]))
  DIST.OBS <- abs(mean(SAMPLE[1:length(STRAIN.LIST)]) - mean(SAMPLE[(length(STRAIN.LIST)+1):length(SAMPLE)]))
  
  DIFF <- DIST.OBS - DIST.SIM
  
  return(DIFF)
}

parent.dir <- PATH
setwd(parent.dir)

####STM1 Analysis date 10112016#########################################################################

ALL.REP <- read.csv("STM1.CLEAN.ADJUST_10112016_DCv34.csv",header=TRUE)
ALL.REP <- subset(ALL.REP, MEDIAN.OUTLIER == "NO" & SD.OUTLIER == "NO" & FSC.OUTLIER == "NO")

SUMMARY <- read.table("STM1.SUMMARY.DATA_10112016_DCv34.txt",header=TRUE)

#SUMMARY <- droplevels(subset(SUMMARY, STRAIN != "1139"))

WT <- subset(ALL.REP, CONDITION == "SHAM")
LOW.FSC <- median(WT$FSC.CORRECT) - 4*mad(WT$FSC.CORRECT)
HIGH.FSC <- median(WT$FSC.CORRECT) + 4*mad(WT$FSC.CORRECT)
LOW.FLUO <- median(WT$YFP.MEDIAN.CORRECT) - 4*mad(WT$YFP.MEDIAN.CORRECT)
HIGH.FLUO <- median(WT$YFP.MEDIAN.CORRECT) + 4*mad(WT$YFP.MEDIAN.CORRECT)
WT <- subset(WT, FSC.CORRECT > LOW.FSC & FSC.CORRECT < HIGH.FSC & YFP.MEDIAN.CORRECT > LOW.FLUO & YFP.MEDIAN.CORRECT < HIGH.FLUO)

#MEDIAN

#Create list of WT phenotype grouped by strain

N.WT <- length(unique(WT$STRAIN))
WT.LIST <- vector('list',N.WT)
for (i in 1:N.WT)
{
  CUR.WT <- subset(WT, STRAIN == unique(WT$STRAIN)[i])
  WT.LIST[[i]] <- CUR.WT$YFP.MEDIAN.CORRECT 
}


#Create list of all strain phenotypes grouped by strain

SUMMARY <- SUMMARY[order(SUMMARY$STRAIN),]
ALL.REP <- droplevels(subset(ALL.REP,STRAIN %in% as.character(SUMMARY$STRAIN)))
ALL.REP <- ALL.REP[order(ALL.REP$STRAIN),]
N.STRAIN <- length(unique(ALL.REP$STRAIN))
STRAIN.LIST <- vector('list',N.STRAIN)
for (i in 1:N.STRAIN)
{
  CUR.STRAIN <- subset(ALL.REP, STRAIN == unique(ALL.REP$STRAIN)[i])
  STRAIN.LIST[[i]] <- CUR.STRAIN$YFP.MEDIAN.CORRECT
}


for (i in 1:length(STRAIN.LIST))
{
  DIFF <- sapply(1:N.PERM,FUN=function(x)	SHUFFLE(STRAIN.LIST = STRAIN.LIST[[i]],WT.LIST = WT.LIST))
  P.VAL <- (length(which(DIFF < 0)) + 1) / (length(which(DIFF < 0)) + length(which(DIFF > 0)) + 1) 	
  SUMMARY[i,"P.VAL.MEDIAN"] <- P.VAL
  print(round(i/length(STRAIN.LIST)*100,1))
}

#NOISE: CV

#Create list of WT phenotype grouped by strain

N.WT <- length(unique(WT$STRAIN))
WT.LIST <- vector('list',N.WT)
for (i in 1:N.WT)
{
  CUR.WT <- subset(WT, STRAIN == unique(WT$STRAIN)[i])
  WT.LIST[[i]] <- CUR.WT$YFP.CV.WT
}


#Create list of all strain phenotypes grouped by strain

SUMMARY <- SUMMARY[order(SUMMARY$STRAIN),]
ALL.REP <- droplevels(subset(ALL.REP,STRAIN %in% as.character(SUMMARY$STRAIN)))
ALL.REP <- ALL.REP[order(ALL.REP$STRAIN),]
N.STRAIN <- length(unique(ALL.REP$STRAIN))
STRAIN.LIST <- vector('list',N.STRAIN)
for (i in 1:N.STRAIN)
{
  CUR.STRAIN <- subset(ALL.REP, STRAIN == unique(ALL.REP$STRAIN)[i])
  STRAIN.LIST[[i]] <- CUR.STRAIN$YFP.CV.WT
}


for (i in 1:length(STRAIN.LIST))
{
  DIFF <- sapply(1:N.PERM,FUN=function(x)	SHUFFLE(STRAIN.LIST = STRAIN.LIST[[i]],WT.LIST = WT.LIST))
  P.VAL <- (length(which(DIFF < 0))+1) / (length(which(DIFF < 0)) + length(which(DIFF > 0)) + 1) 	
  SUMMARY[i,"P.VAL.CV.WT"] <- P.VAL
  print(round(i/length(STRAIN.LIST)*100,1))
}

#NOISE: FANO

#Create list of WT phenotype grouped by strain

N.WT <- length(unique(WT$STRAIN))
WT.LIST <- vector('list',N.WT)
for (i in 1:N.WT)
{
  CUR.WT <- subset(WT, STRAIN == unique(WT$STRAIN)[i])
  WT.LIST[[i]] <- CUR.WT$YFP.FANO.WT
}


#Create list of all strain phenotypes grouped by strain

SUMMARY <- SUMMARY[order(SUMMARY$STRAIN),]
ALL.REP <- droplevels(subset(ALL.REP,STRAIN %in% as.character(SUMMARY$STRAIN)))
ALL.REP <- ALL.REP[order(ALL.REP$STRAIN),]
N.STRAIN <- length(unique(ALL.REP$STRAIN))
STRAIN.LIST <- vector('list',N.STRAIN)
for (i in 1:N.STRAIN)
{
  CUR.STRAIN <- subset(ALL.REP, STRAIN == unique(ALL.REP$STRAIN)[i])
  STRAIN.LIST[[i]] <- CUR.STRAIN$YFP.FANO.WT
}


for (i in 1:length(STRAIN.LIST))
{
  DIFF <- sapply(1:N.PERM,FUN=function(x)	SHUFFLE(STRAIN.LIST = STRAIN.LIST[[i]],WT.LIST = WT.LIST))
  P.VAL <- (length(which(DIFF < 0))+1) / (length(which(DIFF < 0)) + length(which(DIFF > 0)) + 1) 	
  SUMMARY[i,"P.VAL.FANO.WT"] <- P.VAL
  print(round(i/length(STRAIN.LIST)*100,1))
}

#FSC MEDIAN

#Create list of WT phenotype grouped by strain

N.WT <- length(unique(WT$STRAIN))
WT.LIST <- vector('list',N.WT)
for (i in 1:N.WT)
{
  CUR.WT <- subset(WT, STRAIN == unique(WT$STRAIN)[i])
  WT.LIST[[i]] <- CUR.WT$FSC.CORRECT
}


#Create list of all strain phenotypes grouped by strain

SUMMARY <- SUMMARY[order(SUMMARY$STRAIN),]
ALL.REP <- droplevels(subset(ALL.REP,STRAIN %in% as.character(SUMMARY$STRAIN)))
ALL.REP <- ALL.REP[order(ALL.REP$STRAIN),]
N.STRAIN <- length(unique(ALL.REP$STRAIN))
STRAIN.LIST <- vector('list',N.STRAIN)
for (i in 1:N.STRAIN)
{
  CUR.STRAIN <- subset(ALL.REP, STRAIN == unique(ALL.REP$STRAIN)[i])
  STRAIN.LIST[[i]] <- CUR.STRAIN$FSC.CORRECT
}


for (i in 1:length(STRAIN.LIST))
{
  DIFF <- sapply(1:N.PERM,FUN=function(x)	SHUFFLE(STRAIN.LIST = STRAIN.LIST[[i]],WT.LIST = WT.LIST))
  P.VAL <- (length(which(DIFF < 0))+1) / (length(which(DIFF < 0)) + length(which(DIFF > 0)) + 1) 	
  SUMMARY[i,"P.VAL.FSC"] <- P.VAL
  print(round(i/length(STRAIN.LIST)*100,1))
}

write.table(SUMMARY,"STM1.SUMMARY.DCv34.Pvals.txt",sep="\t",quote=FALSE,row.names=FALSE)

#____________________________________________________________________________________________________

#########################
#  TDH1
#########################


#Clear memory
rm(list=ls())
options(warn=-1)
#Set source directory 
parent.dir <- "C:/Users/andreahd/Dropbox/Wittkopp postdoc/projects/Templates.Analysis/"
setwd(parent.dir)
source("Cleaning.Functions.RNA.v34.R")

#Set working directory
parent.dir <- "E:/wittkopp data/processed data/RNA scale analyses"
setwd(parent.dir)


#TDH1

#Load experiment setup
SETUP <- read.csv("Template_TDH1_V3_10102016_E.csv",header=TRUE,as.is=TRUE)
TESTNAMES <- list.files("PATH/TDH1",pattern=".fcs",full.name=TRUE,recursive=TRUE,include.dirs=TRUE)
SETUP<-subset(SETUP, SETUP$FILENAMES %in% TESTNAMES, drop = TRUE)
SETUP[,"COUNTER"] <- c(1:nrow(SETUP))

#Determine Hard Gates
GATES <- GATE.CALIB(SETUP$FILENAMES[1])

#Clean Data
Output <- apply(SETUP,1,CLEANING,GATES=GATES)
OUTPUT <- as.data.frame(Output[[1]])
OUTPUT[2:nrow(SETUP),1:ncol(OUTPUT)] <- NA
for (i in 2:nrow(SETUP))
{
  if (is.null(Output[[i]]))
  {} 
  else {
    OUTPUT[i,] <- Output[[i]]
  }
}

write.table(OUTPUT,"TDH1_Experiment.Output_V3_10112016_DebrisCleanedv34.txt",row.names=FALSE,sep="\t")

CLEAN <- cbind.data.frame(SETUP[,1:(ncol(SETUP)-2)],OUTPUT)

write.table(CLEAN,"TDH1_Clean.Data_V3_10112016_DebrisCleanedv34.txt",row.names=FALSE,sep="\t")


################################################################
#     PLATE CORRECTION & OUTLIER REMOVAL
################################################################

#Clear memory
rm(list=ls())
options(warn=-1)

#####################
#1-LOADING LIBRARIES#
#####################
library(flowCore)
library(flowClust)
library(flowViz)
library(pcaPP)
library(mixtools)
library(plyr)
library(MASS)
box <- graphics::box
options(warn=-1)

####################################
#2- Quality Control and Corrections#
####################################

parent.dir <- PATH
setwd(parent.dir)

DATA.TYPE <- c(
  "factor","factor","factor","factor","factor","factor","factor","factor","factor","factor","factor","factor","factor","factor","factor",
  "integer","integer","integer","integer",
  "numeric","numeric","numeric","numeric","numeric","numeric","numeric","numeric","numeric","numeric","numeric","numeric","numeric","numeric","numeric","numeric","numeric","numeric","numeric","numeric","numeric","numeric","numeric","numeric","numeric",
  "integer")

###Read in data sets
CLEAN.DATA <- read.table("TDH1_Clean.Data_V3_10112016_DebrisCleanedv34.txt",colClasses=DATA.TYPE,header=TRUE, sep="\t")
CLEAN.DATA$STRAIN<-CLEAN.DATA$ASSAY.PLATE.WELL

###Remove samples with NA
CLEAN.DATA <- CLEAN.DATA[complete.cases(CLEAN.DATA),]

##Translate into log scale for appropriate correction
CLEAN.DATA[,"log.RNA.MEDIAN"] <- log((CLEAN.DATA[,"YFP.MEDIAN.FINAL"]) + 0.05)
CLEAN.DATA[,"log.RNA.MAD"] <- CLEAN.DATA[,"YFP.MAD.FINAL"] / (CLEAN.DATA[,"YFP.MEDIAN.FINAL"] + 0.05)
CLEAN.DATA[,"log.RNA.SD"] <- CLEAN.DATA[,"YFP.SD.FINAL"] / (CLEAN.DATA[,"YFP.MEDIAN.FINAL"] + 0.05)

###Remove samples with less than 1000 events
CLEAN.DATA <- subset(CLEAN.DATA,CLEAN.DATA$COUNTS.FINAL > 1000)

####Add variable to ID whether control is own promoter or TDH3
CLEAN.DATA[,"CTRL.GENO"]<-as.factor("TDH1")


###Visualize data before correction / filtering

CLEAN.DATA$CONTAM.CHECK<-"NONE"
CLEAN.DATA$CONTAM.CHECK[(CLEAN.DATA$CONDITION=="EMS"|CLEAN.DATA$CONDITION=="SHAM"|CLEAN.DATA$CONDITION=="EMS.LOW"|CLEAN.DATA$CONDITION=="EMS.HIGH") & (CLEAN.DATA$YFP.MEDIAN.FINAL<.55)]<-"BACT.CONTAM.CHECK"
CLEAN.DATA$CONTAM.CHECK[(CLEAN.DATA$CONDITION=="EMS"|CLEAN.DATA$CONDITION=="SHAM"|CLEAN.DATA$CONDITION=="EMS.LOW"|CLEAN.DATA$CONDITION=="EMS.HIGH") & (CLEAN.DATA$YFP.MEDIAN.FINAL>0.8)]<-"TDH3.CONTAM.CHECK"
CLEAN.DATA$CONTAM.CHECK[(CLEAN.DATA$CONDITION=="TDH3.SHAM" & CLEAN.DATA$YFP.MEDIAN.FINAL<0.8)]<-"EXP.SHAM.CONTAM.CHECK"
CLEAN.DATA$CONTAM.CHECK[(CLEAN.DATA$CONDITION=="NULL" & CLEAN.DATA$YFP.MEDIAN.FINAL>0.55)]<-"EXP.NULL.CONTAM.CHECK"
CLEAN.DATA$CONTAM.CHECK[CLEAN.DATA$FSC.MEDIAN.FINAL<5.5]<-"BACT.CONTAM.CHECK"


###Seperate controls and remove FSC outliers
CONTROL <- subset(CLEAN.DATA,CLEAN.DATA$CONDITION == "CTRL")
FSC.CORRECT <- lm(CONTROL$FSC.MEDIAN.FINAL ~ 0 + CONTROL$FLOW.RUN)
RESID <- FSC.CORRECT$resid
CLEAN.DATA <- droplevels(CLEAN.DATA[-which(CLEAN.DATA$CONDITION == "CTRL")[abs(RESID) > 3*mad(RESID)],])
CONTROL <- droplevels(subset(CLEAN.DATA,CLEAN.DATA$CONDITION == "CTRL"))

INITIAL.CORRECT <- lm(CONTROL$log.RNA.MEDIAN ~ 0 + CONTROL$FLOW.RUN)
RESID <- INITIAL.CORRECT$resid
CLEAN.DATA<- droplevels(CLEAN.DATA[-which(CLEAN.DATA$CONDITION == "CTRL")[abs(RESID) > 3*mad(RESID)],])
CONTROL <- droplevels(subset(CLEAN.DATA,CLEAN.DATA$CONDITION == "CTRL"))

`%notin%` <- function(x,y) !(x %in% y)
RESULT<-count(CONTROL$FLOW.RUN)
RESULT.MUT<-count(CLEAN.DATA$FLOW.RUN)
EXCLUDE1<-(RESULT.MUT[which(RESULT.MUT[,1] %notin% RESULT[,1]),1])
EXCLUDE2<-(RESULT[which(RESULT[,2] <5),1])
CLEAN.DATA<-droplevels(CLEAN.DATA[CLEAN.DATA$FLOW.RUN %notin% EXCLUDE1,])
CLEAN.DATA<-droplevels(CLEAN.DATA[CLEAN.DATA$FLOW.RUN %notin% EXCLUDE2,])
CONTROL <- subset(CLEAN.DATA,CLEAN.DATA$CONDITION == "CTRL")


###Corrections for YFP median and MAD
#Correct Median
#fit<- lm(log.RNA.MEDIAN~FLOW.RUN+REP+STACK.ORDER+STACK.ORDER:FLOW.RUN+REP+ROW+COLUMN+BLOCK+BLOCK:ROW+ROW:COLUMN+STACK.ORDER:BLOCK,data=CONTROL)
#step <- stepAIC(fit, direction="both")
log.RNA.MEDIAN.CORRECT <- lm(log.RNA.MEDIAN ~ FLOW.RUN + COLUMN + ROW + ROW:COLUMN, data=CONTROL)
CLEAN.DATA[,"log.RNA.MEDIAN.CORRECT"] <- CLEAN.DATA$log.RNA.MEDIAN - predict(log.RNA.MEDIAN.CORRECT, CLEAN.DATA) + mean(CONTROL$log.RNA.MEDIAN)
CONTROL <- subset(CLEAN.DATA,CLEAN.DATA$CONDITION == "CTRL")

#Correct sd
#fit<- lm(log.RNA.SD~FLOW.RUN+REP+STACK.ORDER+STACK.ORDER:FLOW.RUN+REP+ROW+COLUMN+BLOCK+BLOCK:ROW+ROW:COLUMN+STACK.ORDER:BLOCK,data=CONTROL)
#step <- step(fit, direction="both")
log.RNA.SD.CORRECT <- lm(log.RNA.SD ~  FLOW.RUN + ROW + COLUMN,data=CONTROL)
CLEAN.DATA[,"log.RNA.SD.CORRECT"] <- CLEAN.DATA$log.RNA.SD - predict(log.RNA.SD.CORRECT , CLEAN.DATA) + mean(CONTROL$log.RNA.SD)
CONTROL <- subset(CLEAN.DATA,CLEAN.DATA$CONDITION == "CTRL")

#Correct mad
#fit<- lm(log.RNA.MAD~FLOW.RUN+REP+STACK.ORDER+STACK.ORDER:FLOW.RUN+REP+ROW+COLUMN+BLOCK+BLOCK:ROW+ROW:COLUMN+STACK.ORDER:BLOCK,data=CONTROL)
#step <- step(fit, direction="both")
log.RNA.MAD.CORRECT <- lm(log.RNA.MAD ~  FLOW.RUN + ROW + COLUMN,data=CONTROL)
CLEAN.DATA[,"log.RNA.MAD.CORRECT"] <- CLEAN.DATA$log.RNA.MAD - predict(log.RNA.MAD.CORRECT , CLEAN.DATA) + mean(CONTROL$log.RNA.MAD)
CONTROL <- subset(CLEAN.DATA,CLEAN.DATA$CONDITION == "CTRL")

#Correct FSC
#fit<- lm(FSC.MEDIAN.FINAL~FLOW.RUN+REP+STACK.ORDER+STACK.ORDER:FLOW.RUN+REP+ROW+COLUMN+BLOCK+BLOCK:ROW+ROW:COLUMN+STACK.ORDER:BLOCK,data=CONTROL)
#step <- stepAIC(fit, direction="both")
FSC.CORRECT <- lm(FSC.MEDIAN.FINAL ~  FLOW.RUN + ROW + COLUMN,data=CONTROL)
CLEAN.DATA[,"FSC.CORRECT"] <- CLEAN.DATA$FSC.MEDIAN.FINAL - predict(FSC.CORRECT, CLEAN.DATA) + mean(CONTROL$FSC.MEDIAN.FINAL)
CONTROL <- subset(CLEAN.DATA,CLEAN.DATA$CONDITION == "CTRL")

#Back translate to linear scale
CLEAN.DATA[,"YFP.MEDIAN.CORRECT"] <- (exp(CLEAN.DATA[,"log.RNA.MEDIAN.CORRECT"]) - 0.05)
CLEAN.DATA[,"YFP.SD.CORRECT"] <- CLEAN.DATA[,"log.RNA.SD.CORRECT"] * CLEAN.DATA[,"YFP.MEDIAN.CORRECT"]
CLEAN.DATA[,"YFP.MAD.CORRECT"] <- CLEAN.DATA[,"log.RNA.MAD.CORRECT"] * CLEAN.DATA[,"YFP.MEDIAN.CORRECT"]


#PERFORM FABIEN'S OUTLIER CORRECTION FILTERING OUT ANY REPLICATE OUTSIDE OF 4*MAD


#Remove outliers
for (i in 1:nrow(CLEAN.DATA))
{
  CUR <- subset(CLEAN.DATA, STRAIN == CLEAN.DATA[i,"STRAIN"])
  if (CLEAN.DATA[i,"YFP.MEDIAN.CORRECT"] > median(CUR$YFP.MEDIAN.CORRECT) + 4*mad(CUR$YFP.MEDIAN.CORRECT) | CLEAN.DATA[i,"YFP.MEDIAN.CORRECT"] < median(CUR$YFP.MEDIAN.CORRECT) - 4*mad(CUR$YFP.MEDIAN.CORRECT)) {
    CLEAN.DATA[i,"MEDIAN.OUTLIER"] <- "YES"
  } else {
    CLEAN.DATA[i,"MEDIAN.OUTLIER"] <- "NO"
  }
  if (CLEAN.DATA[i,"YFP.SD.CORRECT"] > median(CUR$YFP.SD.CORRECT) + 4*mad(CUR$YFP.SD.CORRECT) | CLEAN.DATA[i,"YFP.SD.CORRECT"] < median(CUR$YFP.SD.CORRECT) - 4*mad(CUR$YFP.SD.CORRECT)) {
    CLEAN.DATA[i,"SD.OUTLIER"] <- "YES"
  } else {
    CLEAN.DATA[i,"SD.OUTLIER"] <- "NO"
  }
  if (CLEAN.DATA[i,"FSC.CORRECT"] > median(CUR$FSC.CORRECT) + 4*mad(CUR$FSC.CORRECT) | CLEAN.DATA[i,"FSC.CORRECT"] < median(CUR$FSC.CORRECT) - 4*mad(CUR$FSC.CORRECT)) {
    CLEAN.DATA[i,"FSC.OUTLIER"] <- "YES"
  } else {
    CLEAN.DATA[i,"FSC.OUTLIER"] <- "NO"
  }	
}

CLEAN.DATA <- subset(CLEAN.DATA, MEDIAN.OUTLIER == "NO" & SD.OUTLIER == "NO" & FSC.OUTLIER == "NO")

#Write processed data to file
write.table(CLEAN.DATA,"TDH1.CLEAN.CORRECT_10112016_DCv34.csv",sep=",",quote=FALSE,row.names=FALSE)

#####################
#5 - Control plots  #
#####################

CONTROL <- droplevels(subset(CLEAN.DATA,CLEAN.DATA$CONDITION == "CTRL"))
SHAM <- droplevels(subset(CLEAN.DATA,CLEAN.DATA$CONDITION == "SHAM"))
TDH3.SHAM <- droplevels(subset(CLEAN.DATA,CLEAN.DATA$CONDITION == "TDH3.SHAM"))


###############################################################################
#
#           Strain Estimates
#
###############################################################################

# Clear memory
rm(list=ls())
options(warn=-1)

setwd(PATH)

#TDH1
CLEAN.DATA <- read.csv("TDH1.CLEAN.CORRECT_10112016_DCv34.csv", header=TRUE)

#Calculate mean and noise for each sample relative to controls
TDH3.SHAM <- subset(CLEAN.DATA, CLEAN.DATA$CONDITION == "TDH3.SHAM")
NEG <- subset(CLEAN.DATA, CLEAN.DATA$CONDITION == "NULL")
WT.SHAM <- subset(CLEAN.DATA, CLEAN.DATA$CONDITION == "SHAM")

#ABSOLUTE EXPRESSION SCALE
CLEAN.DATA[,"YFP.MEDIAN.ABS"] <- (CLEAN.DATA$YFP.MEDIAN.CORRECT - median(NEG$YFP.MEDIAN.CORRECT))
CLEAN.DATA[,"YFP.SD.ABS"]   <- CLEAN.DATA$YFP.SD.CORRECT - median(NEG$YFP.SD.CORRECT)
CLEAN.DATA[,"YFP.MAD.ABS"]   <- CLEAN.DATA$YFP.MAD.CORRECT - median(NEG$YFP.SD.CORRECT)

#Re-calculate mean and noise for each sample relative to controls
TDH3.SHAM <- subset(CLEAN.DATA, CLEAN.DATA$CONDITION == "TDH3.SHAM")
NEG <- subset(CLEAN.DATA, CLEAN.DATA$CONDITION == "NULL")
WT.SHAM <- subset(CLEAN.DATA, CLEAN.DATA$CONDITION == "SHAM")

#PROMOTER-SPECIFIC SCALING
CLEAN.DATA[,"YFP.MEDIAN.WT"] <- CLEAN.DATA$YFP.MEDIAN.ABS/median(WT.SHAM$YFP.MEDIAN.ABS)
CLEAN.DATA[,"YFP.SD.WT"]   <- CLEAN.DATA$YFP.SD.ABS/median(WT.SHAM$YFP.MEDIAN.ABS)
CLEAN.DATA[,"YFP.MAD.WT"]   <- CLEAN.DATA$YFP.MAD.ABS/median(WT.SHAM$YFP.MEDIAN.ABS)

#RELATIVE TO TDH3 EXPRESSION
CLEAN.DATA[,"YFP.MEDIAN.TDH3"] <- CLEAN.DATA$YFP.MEDIAN.ABS/median(TDH3.SHAM$YFP.MEDIAN.ABS)
CLEAN.DATA[,"YFP.SD.TDH3"]   <- CLEAN.DATA$YFP.SD.ABS/median(TDH3.SHAM$YFP.MEDIAN.ABS)
CLEAN.DATA[,"YFP.MAD.TDH3"]   <- CLEAN.DATA$YFP.MAD.ABS/median(TDH3.SHAM$YFP.MEDIAN.ABS)

#Re-calculate mean and noise for each sample relative to controls
TDH3.SHAM <- subset(CLEAN.DATA, CLEAN.DATA$CONDITION == "TDH3.SHAM")
TDH3.SHAM<-subset(TDH3.SHAM,TDH3.SHAM$YFP.MEDIAN.FINAL>0.7)
NEG <- subset(CLEAN.DATA, CLEAN.DATA$CONDITION == "NULL")
WT.SHAM <- subset(CLEAN.DATA, CLEAN.DATA$CONDITION == "SHAM")

#CALCULATE NOISE METRICS
CLEAN.DATA[,"YFP.CV.WT"]   <- CLEAN.DATA$YFP.SD.WT/CLEAN.DATA$YFP.MEDIAN.WT
CLEAN.DATA[,"YFP.CV.TDH3"]   <- CLEAN.DATA$YFP.SD.TDH3/CLEAN.DATA$YFP.MEDIAN.TDH3
CLEAN.DATA[,"YFP.FANO.WT"]   <- CLEAN.DATA$YFP.SD.WT^2/CLEAN.DATA$YFP.MEDIAN.WT
CLEAN.DATA[,"YFP.FANO.TDH3"]   <- CLEAN.DATA$YFP.SD.TDH3^2/CLEAN.DATA$YFP.MEDIAN.TDH3


#Re-calculate mean and noise for each sample relative to controls
TDH3.SHAM <- subset(CLEAN.DATA, CLEAN.DATA$CONDITION == "TDH3.SHAM")
TDH3.SHAM<-subset(TDH3.SHAM,TDH3.SHAM$YFP.MEDIAN.FINAL>0.7)
NEG <- subset(CLEAN.DATA, CLEAN.DATA$CONDITION == "NULL")
WT.SHAM <- subset(CLEAN.DATA, CLEAN.DATA$CONDITION == "SHAM")

#CALCULATE RELATIVE NOISE METRICS
CLEAN.DATA[,"YFP.SD.WT.REL"]   <- CLEAN.DATA$YFP.SD.WT/median(WT.SHAM$YFP.SD.WT)
CLEAN.DATA[,"YFP.SD.TDH3.REL"]   <- CLEAN.DATA$YFP.SD.TDH3/median(WT.SHAM$YFP.SD.TDH3)
CLEAN.DATA[,"YFP.CV.WT.REL"]   <- CLEAN.DATA$YFP.SD.WT.REL/CLEAN.DATA$YFP.MEDIAN.WT
CLEAN.DATA[,"YFP.CV.TDH3.REL"]   <- CLEAN.DATA$YFP.SD.TDH3.REL/CLEAN.DATA$YFP.MEDIAN.TDH3
CLEAN.DATA[,"YFP.FANO.WT.REL"]   <- CLEAN.DATA$YFP.SD.WT.REL^2/CLEAN.DATA$YFP.MEDIAN.WT
CLEAN.DATA[,"YFP.FANO.TDH3.REL"]   <- CLEAN.DATA$YFP.SD.TDH3.REL^2/CLEAN.DATA$YFP.MEDIAN.TDH3


#Add NA for samples where CV or FANO are undefined.
for (i in 1:nrow(CLEAN.DATA))
{
  if (CLEAN.DATA[i,"YFP.MEDIAN.ABS"] < 0 | CLEAN.DATA[i,"YFP.SD.ABS"] < 0) {
    CLEAN.DATA[i,"YFP.MAD.WT"] <- NA
    CLEAN.DATA[i,"YFP.MAD.TDH3"] <- NA
    CLEAN.DATA[i,"YFP.CV.WT"] <- NA
    CLEAN.DATA[i,"YFP.CV.TDH3"] <- NA
    CLEAN.DATA[i,"YFP.CV.WT.REL"] <- NA
    CLEAN.DATA[i,"YFP.CV.TDH3.REL"] <- NA
    CLEAN.DATA[i,"YFP.FANO.WT"] <- NA
    CLEAN.DATA[i,"YFP.FANO.TDH3"] <- NA
    CLEAN.DATA[i,"YFP.FANO.WT.REL"] <- NA
    CLEAN.DATA[i,"YFP.FANO.TDH3.REL"] <- NA
    
  }
}

CLEAN.DATA <- droplevels(subset(CLEAN.DATA,CLEAN.DATA$CONDITION!="CTRL"))

write.table(CLEAN.DATA,"TDH1.CLEAN.ADJUST_10112016_DCv34.csv",sep=",",quote=FALSE,row.names=FALSE)

CLEAN.DATA <- read.csv("TDH1.CLEAN.ADJUST_10112016_DCv34.csv", header=TRUE)

TDH1.MEAN <- aggregate(cbind(YFP.MEDIAN.ABS,YFP.SD.ABS,YFP.MAD.ABS,YFP.MEDIAN.WT,YFP.SD.WT,YFP.MAD.WT,YFP.MEDIAN.TDH3,YFP.SD.TDH3,YFP.MAD.TDH3,YFP.CV.WT,YFP.FANO.WT,YFP.CV.TDH3,YFP.FANO.TDH3,YFP.SD.WT.REL,YFP.CV.WT.REL,YFP.FANO.WT.REL,YFP.SD.TDH3.REL,YFP.CV.TDH3.REL,YFP.FANO.TDH3.REL,FSC.CORRECT) ~ STRAIN + CONDITION + ASSAY + PLATE + CTRL.GENO, data=CLEAN.DATA, FUN = mean)
TDH1.SD <- aggregate(cbind(YFP.MEDIAN.ABS,YFP.SD.ABS,YFP.MAD.ABS,YFP.MEDIAN.WT,YFP.SD.WT,YFP.MAD.WT,YFP.MEDIAN.TDH3,YFP.SD.TDH3,YFP.MAD.TDH3,YFP.CV.WT,YFP.FANO.WT,YFP.CV.TDH3,YFP.FANO.TDH3,YFP.SD.WT.REL,YFP.CV.WT.REL,YFP.FANO.WT.REL,YFP.SD.TDH3.REL,YFP.CV.TDH3.REL,YFP.FANO.TDH3.REL,FSC.CORRECT) ~ STRAIN + CONDITION + ASSAY + PLATE + CTRL.GENO, data=CLEAN.DATA, FUN = sd)
TDH1.N <- aggregate(cbind(YFP.MEDIAN.ABS,YFP.SD.ABS,YFP.MAD.ABS,YFP.MEDIAN.WT,YFP.SD.WT,YFP.MAD.WT,YFP.MEDIAN.TDH3,YFP.SD.TDH3,YFP.MAD.TDH3,YFP.CV.WT,YFP.FANO.WT,YFP.CV.TDH3,YFP.FANO.TDH3,YFP.SD.WT.REL,YFP.CV.WT.REL,YFP.FANO.WT.REL,YFP.SD.TDH3.REL,YFP.CV.TDH3.REL,YFP.FANO.TDH3.REL,FSC.CORRECT) ~ STRAIN + CONDITION + ASSAY + PLATE + CTRL.GENO, data=CLEAN.DATA, FUN = length)

TDH1.MEAN  <- TDH1.MEAN[which(TDH1.N[,6] >= 3),]
TDH1.SD <- TDH1.SD[which(TDH1.N[,6] >= 3),]
TDH1.N <- TDH1.N[which(TDH1.N[,6] >= 3),]

TDH1 <- cbind.data.frame(TDH1.MEAN,TDH1.SD[,6:25],TDH1.N[,6])

colnames(TDH1) <- c("STRAIN","CONDITION","ASSAY","PLATE","CTRL.GENO",paste0("MEAN.",colnames(TDH1.MEAN[,6:25])),paste0("SD.",colnames(TDH1.SD[,6:25])),"N")

write.table(TDH1,"TDH1.SUMMARY.DATA_10112016_DCv34.txt",sep="\t",quote=FALSE,row.names=FALSE)

`%notin%` <- function(x,y) !(x %in% y)

CONTAM.CHECK<-droplevels(subset(CLEAN.DATA,CLEAN.DATA$CONTAM.CHECK!="NONE"))

if(all(CONTAM.CHECK$STRAIN %notin% TDH1$STRAIN)) {
  #All datapoints flagged as suspicious have been removed by outlier removal filters.
  print("Yay! Suspicious values eliminated. Proceed.")
}

if(any(CONTAM.CHECK$STRAIN %in% TDH1$STRAIN)) {
  CONTAM.CHECK<-CONTAM.CHECK[CONTAM.CHECK$STRAIN %in% CONTAM.CHECK$STRAIN[which(CONTAM.CHECK$STRAIN %in% TDH1$STRAIN)],]
  write.table(CONTAM.CHECK,"TDH1.CONTAM.CHECK_outliersexcl.csv",sep=",",quote=FALSE,row.names=FALSE)  
}


###############################
#HIERARCHICAL PERMUTATION TEST#
###############################

#Clear memory
rm(list=ls())
options(warn=-1)

library(permute)
library(Hmisc)

N.PERM <- 10000

#Permutation function

SHUFFLE <- function(STRAIN.LIST = STRAIN.LIST[[i]],WT.LIST = WT.LIST) {
  SAMPLE <- c(STRAIN.LIST,WT.LIST[[sample(1:length(WT.LIST),1)]])
  OUTPUT <- sample(SAMPLE,length(SAMPLE))
  DIST.SIM <- abs(mean(OUTPUT[1:length(STRAIN.LIST)]) - mean(OUTPUT[(length(STRAIN.LIST)+1):length(SAMPLE)]))
  DIST.OBS <- abs(mean(SAMPLE[1:length(STRAIN.LIST)]) - mean(SAMPLE[(length(STRAIN.LIST)+1):length(SAMPLE)]))
  
  DIFF <- DIST.OBS - DIST.SIM
  
  return(DIFF)
}

parent.dir <- PATH
setwd(parent.dir)

####TDH1 Analysis date 10112016#########################################################################

ALL.REP <- read.csv("TDH1.CLEAN.ADJUST_10112016_DCv34.csv",header=TRUE)
ALL.REP <- subset(ALL.REP, MEDIAN.OUTLIER == "NO" & SD.OUTLIER == "NO" & FSC.OUTLIER == "NO")

SUMMARY <- read.table("TDH1.SUMMARY.DATA_10112016_DCv34.txt",header=TRUE)

#SUMMARY <- droplevels(subset(SUMMARY, STRAIN != "1139"))

WT <- subset(ALL.REP, CONDITION == "SHAM")
LOW.FSC <- median(WT$FSC.CORRECT) - 4*mad(WT$FSC.CORRECT)
HIGH.FSC <- median(WT$FSC.CORRECT) + 4*mad(WT$FSC.CORRECT)
LOW.FLUO <- median(WT$YFP.MEDIAN.CORRECT) - 4*mad(WT$YFP.MEDIAN.CORRECT)
HIGH.FLUO <- median(WT$YFP.MEDIAN.CORRECT) + 4*mad(WT$YFP.MEDIAN.CORRECT)
WT <- subset(WT, FSC.CORRECT > LOW.FSC & FSC.CORRECT < HIGH.FSC & YFP.MEDIAN.CORRECT > LOW.FLUO & YFP.MEDIAN.CORRECT < HIGH.FLUO)

#MEDIAN

#Create list of WT phenotype grouped by strain

N.WT <- length(unique(WT$STRAIN))
WT.LIST <- vector('list',N.WT)
for (i in 1:N.WT)
{
  CUR.WT <- subset(WT, STRAIN == unique(WT$STRAIN)[i])
  WT.LIST[[i]] <- CUR.WT$YFP.MEDIAN.CORRECT 
}


#Create list of all strain phenotypes grouped by strain

SUMMARY <- SUMMARY[order(SUMMARY$STRAIN),]
ALL.REP <- droplevels(subset(ALL.REP,STRAIN %in% as.character(SUMMARY$STRAIN)))
ALL.REP <- ALL.REP[order(ALL.REP$STRAIN),]
N.STRAIN <- length(unique(ALL.REP$STRAIN))
STRAIN.LIST <- vector('list',N.STRAIN)
for (i in 1:N.STRAIN)
{
  CUR.STRAIN <- subset(ALL.REP, STRAIN == unique(ALL.REP$STRAIN)[i])
  STRAIN.LIST[[i]] <- CUR.STRAIN$YFP.MEDIAN.CORRECT
}


for (i in 1:length(STRAIN.LIST))
{
  DIFF <- sapply(1:N.PERM,FUN=function(x)	SHUFFLE(STRAIN.LIST = STRAIN.LIST[[i]],WT.LIST = WT.LIST))
  P.VAL <- (length(which(DIFF < 0)) + 1) / (length(which(DIFF < 0)) + length(which(DIFF > 0)) + 1) 	
  SUMMARY[i,"P.VAL.MEDIAN"] <- P.VAL
  print(round(i/length(STRAIN.LIST)*100,1))
}

#NOISE: CV

#Create list of WT phenotype grouped by strain

N.WT <- length(unique(WT$STRAIN))
WT.LIST <- vector('list',N.WT)
for (i in 1:N.WT)
{
  CUR.WT <- subset(WT, STRAIN == unique(WT$STRAIN)[i])
  WT.LIST[[i]] <- CUR.WT$YFP.CV.WT
}


#Create list of all strain phenotypes grouped by strain

SUMMARY <- SUMMARY[order(SUMMARY$STRAIN),]
ALL.REP <- droplevels(subset(ALL.REP,STRAIN %in% as.character(SUMMARY$STRAIN)))
ALL.REP <- ALL.REP[order(ALL.REP$STRAIN),]
N.STRAIN <- length(unique(ALL.REP$STRAIN))
STRAIN.LIST <- vector('list',N.STRAIN)
for (i in 1:N.STRAIN)
{
  CUR.STRAIN <- subset(ALL.REP, STRAIN == unique(ALL.REP$STRAIN)[i])
  STRAIN.LIST[[i]] <- CUR.STRAIN$YFP.CV.WT
}


for (i in 1:length(STRAIN.LIST))
{
  DIFF <- sapply(1:N.PERM,FUN=function(x)	SHUFFLE(STRAIN.LIST = STRAIN.LIST[[i]],WT.LIST = WT.LIST))
  P.VAL <- (length(which(DIFF < 0))+1) / (length(which(DIFF < 0)) + length(which(DIFF > 0)) + 1) 	
  SUMMARY[i,"P.VAL.CV.WT"] <- P.VAL
  print(round(i/length(STRAIN.LIST)*100,1))
}

#NOISE: FANO

#Create list of WT phenotype grouped by strain

N.WT <- length(unique(WT$STRAIN))
WT.LIST <- vector('list',N.WT)
for (i in 1:N.WT)
{
  CUR.WT <- subset(WT, STRAIN == unique(WT$STRAIN)[i])
  WT.LIST[[i]] <- CUR.WT$YFP.FANO.WT
}


#Create list of all strain phenotypes grouped by strain

SUMMARY <- SUMMARY[order(SUMMARY$STRAIN),]
ALL.REP <- droplevels(subset(ALL.REP,STRAIN %in% as.character(SUMMARY$STRAIN)))
ALL.REP <- ALL.REP[order(ALL.REP$STRAIN),]
N.STRAIN <- length(unique(ALL.REP$STRAIN))
STRAIN.LIST <- vector('list',N.STRAIN)
for (i in 1:N.STRAIN)
{
  CUR.STRAIN <- subset(ALL.REP, STRAIN == unique(ALL.REP$STRAIN)[i])
  STRAIN.LIST[[i]] <- CUR.STRAIN$YFP.FANO.WT
}


for (i in 1:length(STRAIN.LIST))
{
  DIFF <- sapply(1:N.PERM,FUN=function(x)	SHUFFLE(STRAIN.LIST = STRAIN.LIST[[i]],WT.LIST = WT.LIST))
  P.VAL <- (length(which(DIFF < 0))+1) / (length(which(DIFF < 0)) + length(which(DIFF > 0)) + 1) 	
  SUMMARY[i,"P.VAL.FANO.WT"] <- P.VAL
  print(round(i/length(STRAIN.LIST)*100,1))
}

#FSC MEDIAN

#Create list of WT phenotype grouped by strain

N.WT <- length(unique(WT$STRAIN))
WT.LIST <- vector('list',N.WT)
for (i in 1:N.WT)
{
  CUR.WT <- subset(WT, STRAIN == unique(WT$STRAIN)[i])
  WT.LIST[[i]] <- CUR.WT$FSC.CORRECT
}


#Create list of all strain phenotypes grouped by strain

SUMMARY <- SUMMARY[order(SUMMARY$STRAIN),]
ALL.REP <- droplevels(subset(ALL.REP,STRAIN %in% as.character(SUMMARY$STRAIN)))
ALL.REP <- ALL.REP[order(ALL.REP$STRAIN),]
N.STRAIN <- length(unique(ALL.REP$STRAIN))
STRAIN.LIST <- vector('list',N.STRAIN)
for (i in 1:N.STRAIN)
{
  CUR.STRAIN <- subset(ALL.REP, STRAIN == unique(ALL.REP$STRAIN)[i])
  STRAIN.LIST[[i]] <- CUR.STRAIN$FSC.CORRECT
}


for (i in 1:length(STRAIN.LIST))
{
  DIFF <- sapply(1:N.PERM,FUN=function(x)	SHUFFLE(STRAIN.LIST = STRAIN.LIST[[i]],WT.LIST = WT.LIST))
  P.VAL <- (length(which(DIFF < 0))+1) / (length(which(DIFF < 0)) + length(which(DIFF > 0)) + 1) 	
  SUMMARY[i,"P.VAL.FSC"] <- P.VAL
  print(round(i/length(STRAIN.LIST)*100,1))
}

write.table(SUMMARY,"TDH1.10112016.SUMMARY.DCv34.Pvals.txt",sep="\t",quote=FALSE,row.names=FALSE)


#___________________________________________________________________________________________________


########################
#   TDH2
########################

#TDH2

#Load experiment setup
SETUP <- read.csv("Template_TDH2_V2_10102016_E.csv",header=TRUE,as.is=TRUE)
TESTNAMES <- list.files("PATH/TDH2",pattern=".fcs",full.name=TRUE,recursive=TRUE,include.dirs=TRUE)
SETUP<-subset(SETUP, SETUP$FILENAMES %in% TESTNAMES, drop = TRUE)

SETUP[,"COUNTER"] <- c(1:nrow(SETUP))

#Determine Hard Gates
GATES <- GATE.CALIB(SETUP$FILENAMES[1])

#Clean Data
Output <- apply(SETUP,1,CLEANING,GATES=GATES)
OUTPUT <- as.data.frame(Output[[1]])
OUTPUT[2:nrow(SETUP),1:ncol(OUTPUT)] <- NA
for (i in 2:nrow(SETUP))
{
  if (is.null(Output[[i]]))
  {} 
  else {
    OUTPUT[i,] <- Output[[i]]
  }
}

write.table(OUTPUT,"TDH2_Experiment.Output_V2_10112016_DebrisCleanedv34.txt",row.names=FALSE,sep="\t")

CLEAN <- cbind.data.frame(SETUP[,1:(ncol(SETUP)-2)],OUTPUT)

write.table(CLEAN,"TDH2_Clean.Data_V2_10112016_DebrisCleanedv34.txt",row.names=FALSE,sep="\t")


################################################################
#     PLATE CORRECTION & OUTLIER REMOVAL
################################################################

#Clear memory
rm(list=ls())
options(warn=-1)

#####################
#1-LOADING LIBRARIES#
#####################
library(flowCore)
library(flowClust)
library(flowViz)
library(pcaPP)
library(mixtools)
library(plyr)
library(MASS)
box <- graphics::box
options(warn=-1)

####################################
#2- Quality Control and Corrections#
####################################

parent.dir <- PATH
setwd(parent.dir)

DATA.TYPE <- c(
  "factor","factor","factor","factor","factor","factor","factor","factor","factor","factor","factor","factor","factor","factor","factor",
  "integer","integer","integer","integer",
  "numeric","numeric","numeric","numeric","numeric","numeric","numeric","numeric","numeric","numeric","numeric","numeric","numeric","numeric","numeric","numeric","numeric","numeric","numeric","numeric","numeric","numeric","numeric","numeric","numeric",
  "integer")

###Read in data sets
CLEAN.DATA <- read.table("TDH2_Clean.Data_V2_10112016_DebrisCleanedv34.txt",colClasses=DATA.TYPE,header=TRUE, sep="\t")

###Remove samples with NA
CLEAN.DATA <- CLEAN.DATA[complete.cases(CLEAN.DATA),]

##Translate into log scale for appropriate correction
CLEAN.DATA[,"log.RNA.MEDIAN"] <- log((CLEAN.DATA[,"YFP.MEDIAN.FINAL"]) + 0.05)
CLEAN.DATA[,"log.RNA.MAD"] <- CLEAN.DATA[,"YFP.MAD.FINAL"] / (CLEAN.DATA[,"YFP.MEDIAN.FINAL"] + 0.05)
CLEAN.DATA[,"log.RNA.SD"] <- CLEAN.DATA[,"YFP.SD.FINAL"] / (CLEAN.DATA[,"YFP.MEDIAN.FINAL"] + 0.05)

###Remove samples with less than 1000 events
CLEAN.DATA <- subset(CLEAN.DATA,CLEAN.DATA$COUNTS.FINAL > 1000)

####Add variable to ID whether control is own promoter or TDH3
CLEAN.DATA[,"CTRL.GENO"]<-as.factor("TDH2")

###Select desired data / Drop known bad samples

CLEAN.DATA$CONTAM.CHECK<-"NONE"
CLEAN.DATA$CONTAM.CHECK[(CLEAN.DATA$CONDITION=="EMS"|CLEAN.DATA$CONDITION=="SHAM"|CLEAN.DATA$CONDITION=="EMS.LOW"|CLEAN.DATA$CONDITION=="EMS.HIGH") & (CLEAN.DATA$YFP.MEDIAN.FINAL<.53)]<-"BACT.CONTAM.CHECK"
CLEAN.DATA$CONTAM.CHECK[(CLEAN.DATA$CONDITION=="EMS"|CLEAN.DATA$CONDITION=="SHAM"|CLEAN.DATA$CONDITION=="EMS.LOW"|CLEAN.DATA$CONDITION=="EMS.HIGH") & (CLEAN.DATA$YFP.MEDIAN.FINAL>0.8)]<-"TDH3.CONTAM.CHECK"
CLEAN.DATA$CONTAM.CHECK[(CLEAN.DATA$CONDITION=="TDH3.SHAM" & CLEAN.DATA$YFP.MEDIAN.FINAL<0.8)]<-"EXP.SHAM.CONTAM.CHECK"
CLEAN.DATA$CONTAM.CHECK[(CLEAN.DATA$CONDITION=="NULL" & CLEAN.DATA$YFP.MEDIAN.FINAL>0.53)]<-"EXP.NULL.CONTAM.CHECK"
CLEAN.DATA$CONTAM.CHECK[CLEAN.DATA$FSC.MEDIAN.FINAL<5.3]<-"BACT.CONTAM.CHECK"


###Seperate controls and remove FSC outliers
CONTROL <- subset(CLEAN.DATA,CLEAN.DATA$CONDITION == "CTRL")
FSC.CORRECT <- lm(CONTROL$FSC.MEDIAN.FINAL ~ 0 + CONTROL$FLOW.RUN)
RESID <- FSC.CORRECT$resid
CLEAN.DATA <- droplevels(CLEAN.DATA[-which(CLEAN.DATA$CONDITION == "CTRL")[abs(RESID) > 3*mad(RESID)],])
CONTROL <- droplevels(subset(CLEAN.DATA,CLEAN.DATA$CONDITION == "CTRL"))

INITIAL.CORRECT <- lm(CONTROL$log.RNA.MEDIAN ~ 0 + CONTROL$FLOW.RUN)
RESID <- INITIAL.CORRECT$resid
CLEAN.DATA<- droplevels(CLEAN.DATA[-which(CLEAN.DATA$CONDITION == "CTRL")[abs(RESID) > 3*mad(RESID)],])
CONTROL <- droplevels(subset(CLEAN.DATA,CLEAN.DATA$CONDITION == "CTRL"))

`%notin%` <- function(x,y) !(x %in% y)
RESULT<-count(CONTROL$FLOW.RUN)
RESULT.MUT<-count(CLEAN.DATA$FLOW.RUN)
EXCLUDE1<-(RESULT.MUT[which(RESULT.MUT[,1] %notin% RESULT[,1]),1])
EXCLUDE2<-(RESULT[which(RESULT[,2] <5),1])
CLEAN.DATA<-droplevels(CLEAN.DATA[CLEAN.DATA$FLOW.RUN %notin% EXCLUDE1,])
CLEAN.DATA<-droplevels(CLEAN.DATA[CLEAN.DATA$FLOW.RUN %notin% EXCLUDE2,])
CONTROL <- subset(CLEAN.DATA,CLEAN.DATA$CONDITION == "CTRL")


###Corrections for YFP median and MAD
#Correct Median
#fit<- lm(log.RNA.MEDIAN~FLOW.RUN+REP+STACK.ORDER+STACK.ORDER:FLOW.RUN+REP+ROW+COLUMN+BLOCK+BLOCK:ROW+ROW:COLUMN+STACK.ORDER:BLOCK,data=CONTROL)
#step <- stepAIC(fit, direction="both")
log.RNA.MEDIAN.CORRECT <- lm(log.RNA.MEDIAN ~ FLOW.RUN + COLUMN + ROW + ROW:COLUMN, data=CONTROL)
CLEAN.DATA[,"log.RNA.MEDIAN.CORRECT"] <- CLEAN.DATA$log.RNA.MEDIAN - predict(log.RNA.MEDIAN.CORRECT, CLEAN.DATA) + mean(CONTROL$log.RNA.MEDIAN)
CONTROL <- subset(CLEAN.DATA,CLEAN.DATA$CONDITION == "CTRL")

#Correct sd
#fit<- lm(log.RNA.SD~FLOW.RUN+REP+STACK.ORDER+STACK.ORDER:FLOW.RUN+REP+ROW+COLUMN+BLOCK+BLOCK:ROW+ROW:COLUMN+STACK.ORDER:BLOCK,data=CONTROL)
#step <- step(fit, direction="both")
log.RNA.SD.CORRECT <- lm(log.RNA.SD ~  FLOW.RUN + ROW + COLUMN+ ROW:COLUMN,data=CONTROL)
CLEAN.DATA[,"log.RNA.SD.CORRECT"] <- CLEAN.DATA$log.RNA.SD - predict(log.RNA.SD.CORRECT , CLEAN.DATA) + mean(CONTROL$log.RNA.SD)
CONTROL <- subset(CLEAN.DATA,CLEAN.DATA$CONDITION == "CTRL")

#Correct mad
#fit<- lm(log.RNA.MAD~FLOW.RUN+REP+STACK.ORDER+STACK.ORDER:FLOW.RUN+REP+ROW+COLUMN+BLOCK+BLOCK:ROW+ROW:COLUMN+STACK.ORDER:BLOCK,data=CONTROL)
#step <- step(fit, direction="both")
log.RNA.MAD.CORRECT <- lm(log.RNA.MAD ~  FLOW.RUN + ROW + COLUMN+ ROW:COLUMN,data=CONTROL)
CLEAN.DATA[,"log.RNA.MAD.CORRECT"] <- CLEAN.DATA$log.RNA.MAD - predict(log.RNA.MAD.CORRECT , CLEAN.DATA) + mean(CONTROL$log.RNA.MAD)
CONTROL <- subset(CLEAN.DATA,CLEAN.DATA$CONDITION == "CTRL")

#Correct FSC
#fit<- lm(FSC.MEDIAN.FINAL~FLOW.RUN+REP+STACK.ORDER+STACK.ORDER:FLOW.RUN+REP+ROW+COLUMN+BLOCK+BLOCK:ROW+ROW:COLUMN+STACK.ORDER:BLOCK,data=CONTROL)
#step <- stepAIC(fit, direction="both")
FSC.CORRECT <- lm(FSC.MEDIAN.FINAL ~  FLOW.RUN + ROW + COLUMN,data=CONTROL)
CLEAN.DATA[,"FSC.CORRECT"] <- CLEAN.DATA$FSC.MEDIAN.FINAL - predict(FSC.CORRECT, CLEAN.DATA) + mean(CONTROL$FSC.MEDIAN.FINAL)
CONTROL <- subset(CLEAN.DATA,CLEAN.DATA$CONDITION == "CTRL")


#Back translate to linear scale
CLEAN.DATA[,"YFP.MEDIAN.CORRECT"] <- (exp(CLEAN.DATA[,"log.RNA.MEDIAN.CORRECT"]) - 0.05)
CLEAN.DATA[,"YFP.SD.CORRECT"] <- CLEAN.DATA[,"log.RNA.SD.CORRECT"] * CLEAN.DATA[,"YFP.MEDIAN.CORRECT"]
CLEAN.DATA[,"YFP.MAD.CORRECT"] <- CLEAN.DATA[,"log.RNA.MAD.CORRECT"] * CLEAN.DATA[,"YFP.MEDIAN.CORRECT"]


#PERFORM FABIEN'S OUTLIER CORRECTION FILTERING OUT ANY REPLICATE OUTSIDE OF 4*MAD


#Remove outliers
for (i in 1:nrow(CLEAN.DATA))
{
  CUR <- subset(CLEAN.DATA, STRAIN == CLEAN.DATA[i,"STRAIN"])
  if (CLEAN.DATA[i,"YFP.MEDIAN.CORRECT"] > median(CUR$YFP.MEDIAN.CORRECT) + 4*mad(CUR$YFP.MEDIAN.CORRECT) | CLEAN.DATA[i,"YFP.MEDIAN.CORRECT"] < median(CUR$YFP.MEDIAN.CORRECT) - 4*mad(CUR$YFP.MEDIAN.CORRECT)) {
    CLEAN.DATA[i,"MEDIAN.OUTLIER"] <- "YES"
  } else {
    CLEAN.DATA[i,"MEDIAN.OUTLIER"] <- "NO"
  }
  if (CLEAN.DATA[i,"YFP.SD.CORRECT"] > median(CUR$YFP.SD.CORRECT) + 4*mad(CUR$YFP.SD.CORRECT) | CLEAN.DATA[i,"YFP.SD.CORRECT"] < median(CUR$YFP.SD.CORRECT) - 4*mad(CUR$YFP.SD.CORRECT)) {
    CLEAN.DATA[i,"SD.OUTLIER"] <- "YES"
  } else {
    CLEAN.DATA[i,"SD.OUTLIER"] <- "NO"
  }
  if (CLEAN.DATA[i,"FSC.CORRECT"] > median(CUR$FSC.CORRECT) + 4*mad(CUR$FSC.CORRECT) | CLEAN.DATA[i,"FSC.CORRECT"] < median(CUR$FSC.CORRECT) - 4*mad(CUR$FSC.CORRECT)) {
    CLEAN.DATA[i,"FSC.OUTLIER"] <- "YES"
  } else {
    CLEAN.DATA[i,"FSC.OUTLIER"] <- "NO"
  }	
}

CLEAN.DATA <- subset(CLEAN.DATA, MEDIAN.OUTLIER == "NO" & SD.OUTLIER == "NO" & FSC.OUTLIER == "NO")

#Write processed data to file
write.table(CLEAN.DATA,"TDH2.CLEAN.CORRECT_10112016_DCv34.csv",sep=",",quote=FALSE,row.names=FALSE)

#####################
#5 - Control plots  #
#####################

CONTROL <- droplevels(subset(CLEAN.DATA,CLEAN.DATA$CONDITION == "CTRL"))
SHAM <- droplevels(subset(CLEAN.DATA,CLEAN.DATA$CONDITION == "SHAM"))
TDH3.SHAM <- droplevels(subset(CLEAN.DATA,CLEAN.DATA$CONDITION == "TDH3.SHAM"))


# ##############################################################################
#
# #Strain Estimates
#
# #####################################################################################
# #Clear memory
rm(list=ls())
options(warn=-1)

setwd("E:/wittkopp data/processed data/RNA scale analyses/DCv34")

#TDH2
CLEAN.DATA <- read.csv("TDH2.CLEAN.CORRECT_10112016_DCv34.csv", header=TRUE)

#Calculate mean and noise for each sample relative to controls
TDH3.SHAM <- subset(CLEAN.DATA, CLEAN.DATA$CONDITION == "TDH3.SHAM")
NEG <- subset(CLEAN.DATA, CLEAN.DATA$CONDITION == "NULL")
WT.SHAM <- subset(CLEAN.DATA, CLEAN.DATA$CONDITION == "SHAM")

#ABSOLUTE EXPRESSION SCALE
CLEAN.DATA[,"YFP.MEDIAN.ABS"] <- (CLEAN.DATA$YFP.MEDIAN.CORRECT - median(NEG$YFP.MEDIAN.CORRECT))
CLEAN.DATA[,"YFP.SD.ABS"]   <- CLEAN.DATA$YFP.SD.CORRECT - median(NEG$YFP.SD.CORRECT)
CLEAN.DATA[,"YFP.MAD.ABS"]   <- CLEAN.DATA$YFP.MAD.CORRECT - median(NEG$YFP.SD.CORRECT)

#Re-calculate mean and noise for each sample relative to controls
TDH3.SHAM <- subset(CLEAN.DATA, CLEAN.DATA$CONDITION == "TDH3.SHAM")
NEG <- subset(CLEAN.DATA, CLEAN.DATA$CONDITION == "NULL")
WT.SHAM <- subset(CLEAN.DATA, CLEAN.DATA$CONDITION == "SHAM")

#PROMOTER-SPECIFIC SCALING
CLEAN.DATA[,"YFP.MEDIAN.WT"] <- CLEAN.DATA$YFP.MEDIAN.ABS/median(WT.SHAM$YFP.MEDIAN.ABS)
CLEAN.DATA[,"YFP.SD.WT"]   <- CLEAN.DATA$YFP.SD.ABS/median(WT.SHAM$YFP.MEDIAN.ABS)
CLEAN.DATA[,"YFP.MAD.WT"]   <- CLEAN.DATA$YFP.MAD.ABS/median(WT.SHAM$YFP.MEDIAN.ABS)

#RELATIVE TO TDH3 EXPRESSION
CLEAN.DATA[,"YFP.MEDIAN.TDH3"] <- CLEAN.DATA$YFP.MEDIAN.ABS/median(TDH3.SHAM$YFP.MEDIAN.ABS)
CLEAN.DATA[,"YFP.SD.TDH3"]   <- CLEAN.DATA$YFP.SD.ABS/median(TDH3.SHAM$YFP.MEDIAN.ABS)
CLEAN.DATA[,"YFP.MAD.TDH3"]   <- CLEAN.DATA$YFP.MAD.ABS/median(TDH3.SHAM$YFP.MEDIAN.ABS)

#Re-calculate mean and noise for each sample relative to controls
TDH3.SHAM <- subset(CLEAN.DATA, CLEAN.DATA$CONDITION == "TDH3.SHAM")
TDH3.SHAM<-subset(TDH3.SHAM,TDH3.SHAM$YFP.MEDIAN.FINAL>0.7)
NEG <- subset(CLEAN.DATA, CLEAN.DATA$CONDITION == "NULL")
WT.SHAM <- subset(CLEAN.DATA, CLEAN.DATA$CONDITION == "SHAM")

#CALCULATE NOISE METRICS
CLEAN.DATA[,"YFP.CV.WT"]   <- CLEAN.DATA$YFP.SD.WT/CLEAN.DATA$YFP.MEDIAN.WT
CLEAN.DATA[,"YFP.CV.TDH3"]   <- CLEAN.DATA$YFP.SD.TDH3/CLEAN.DATA$YFP.MEDIAN.TDH3
CLEAN.DATA[,"YFP.FANO.WT"]   <- CLEAN.DATA$YFP.SD.WT^2/CLEAN.DATA$YFP.MEDIAN.WT
CLEAN.DATA[,"YFP.FANO.TDH3"]   <- CLEAN.DATA$YFP.SD.TDH3^2/CLEAN.DATA$YFP.MEDIAN.TDH3


#Re-calculate mean and noise for each sample relative to controls
TDH3.SHAM <- subset(CLEAN.DATA, CLEAN.DATA$CONDITION == "TDH3.SHAM")
TDH3.SHAM<-subset(TDH3.SHAM,TDH3.SHAM$YFP.MEDIAN.FINAL>0.7)
NEG <- subset(CLEAN.DATA, CLEAN.DATA$CONDITION == "NULL")
WT.SHAM <- subset(CLEAN.DATA, CLEAN.DATA$CONDITION == "SHAM")

#CALCULATE RELATIVE NOISE METRICS
CLEAN.DATA[,"YFP.SD.WT.REL"]   <- CLEAN.DATA$YFP.SD.WT/median(WT.SHAM$YFP.SD.WT)
CLEAN.DATA[,"YFP.SD.TDH3.REL"]   <- CLEAN.DATA$YFP.SD.TDH3/median(WT.SHAM$YFP.SD.TDH3)
CLEAN.DATA[,"YFP.CV.WT.REL"]   <- CLEAN.DATA$YFP.SD.WT.REL/CLEAN.DATA$YFP.MEDIAN.WT
CLEAN.DATA[,"YFP.CV.TDH3.REL"]   <- CLEAN.DATA$YFP.SD.TDH3.REL/CLEAN.DATA$YFP.MEDIAN.TDH3
CLEAN.DATA[,"YFP.FANO.WT.REL"]   <- CLEAN.DATA$YFP.SD.WT.REL^2/CLEAN.DATA$YFP.MEDIAN.WT
CLEAN.DATA[,"YFP.FANO.TDH3.REL"]   <- CLEAN.DATA$YFP.SD.TDH3.REL^2/CLEAN.DATA$YFP.MEDIAN.TDH3


#Add NA for samples where CV or FANO are undefined.
for (i in 1:nrow(CLEAN.DATA))
{
  if (CLEAN.DATA[i,"YFP.MEDIAN.ABS"] < 0 | CLEAN.DATA[i,"YFP.SD.ABS"] < 0) {
    CLEAN.DATA[i,"YFP.MAD.WT"] <- NA
    CLEAN.DATA[i,"YFP.MAD.TDH3"] <- NA
    CLEAN.DATA[i,"YFP.CV.WT"] <- NA
    CLEAN.DATA[i,"YFP.CV.TDH3"] <- NA
    CLEAN.DATA[i,"YFP.CV.WT.REL"] <- NA
    CLEAN.DATA[i,"YFP.CV.TDH3.REL"] <- NA
    CLEAN.DATA[i,"YFP.FANO.WT"] <- NA
    CLEAN.DATA[i,"YFP.FANO.TDH3"] <- NA
    CLEAN.DATA[i,"YFP.FANO.WT.REL"] <- NA
    CLEAN.DATA[i,"YFP.FANO.TDH3.REL"] <- NA
    
  }
}

CLEAN.DATA <- droplevels(subset(CLEAN.DATA,CLEAN.DATA$CONDITION!="CTRL"))

write.table(CLEAN.DATA,"TDH2.CLEAN.ADJUST_10112016_DCv34.csv",sep=",",quote=FALSE,row.names=FALSE)

CLEAN.DATA <- read.csv("TDH2.CLEAN.ADJUST_10112016_DCv34.csv", header=TRUE)

TDH2.MEAN <- aggregate(cbind(YFP.MEDIAN.ABS,YFP.SD.ABS,YFP.MAD.ABS,YFP.MEDIAN.WT,YFP.SD.WT,YFP.MAD.WT,YFP.MEDIAN.TDH3,YFP.SD.TDH3,YFP.MAD.TDH3,YFP.CV.WT,YFP.FANO.WT,YFP.CV.TDH3,YFP.FANO.TDH3,YFP.SD.WT.REL,YFP.CV.WT.REL,YFP.FANO.WT.REL,YFP.SD.TDH3.REL,YFP.CV.TDH3.REL,YFP.FANO.TDH3.REL,FSC.CORRECT) ~ STRAIN + CONDITION + ASSAY + PLATE + CTRL.GENO, data=CLEAN.DATA, FUN = mean)
TDH2.SD <- aggregate(cbind(YFP.MEDIAN.ABS,YFP.SD.ABS,YFP.MAD.ABS,YFP.MEDIAN.WT,YFP.SD.WT,YFP.MAD.WT,YFP.MEDIAN.TDH3,YFP.SD.TDH3,YFP.MAD.TDH3,YFP.CV.WT,YFP.FANO.WT,YFP.CV.TDH3,YFP.FANO.TDH3,YFP.SD.WT.REL,YFP.CV.WT.REL,YFP.FANO.WT.REL,YFP.SD.TDH3.REL,YFP.CV.TDH3.REL,YFP.FANO.TDH3.REL,FSC.CORRECT) ~ STRAIN + CONDITION + ASSAY + PLATE + CTRL.GENO, data=CLEAN.DATA, FUN = sd)
TDH2.N <- aggregate(cbind(YFP.MEDIAN.ABS,YFP.SD.ABS,YFP.MAD.ABS,YFP.MEDIAN.WT,YFP.SD.WT,YFP.MAD.WT,YFP.MEDIAN.TDH3,YFP.SD.TDH3,YFP.MAD.TDH3,YFP.CV.WT,YFP.FANO.WT,YFP.CV.TDH3,YFP.FANO.TDH3,YFP.SD.WT.REL,YFP.CV.WT.REL,YFP.FANO.WT.REL,YFP.SD.TDH3.REL,YFP.CV.TDH3.REL,YFP.FANO.TDH3.REL,FSC.CORRECT) ~ STRAIN + CONDITION + ASSAY + PLATE + CTRL.GENO, data=CLEAN.DATA, FUN = length)

TDH2.MEAN  <- TDH2.MEAN[which(TDH2.N[,6] >= 3),]
TDH2.SD <- TDH2.SD[which(TDH2.N[,6] >= 3),]
TDH2.N <- TDH2.N[which(TDH2.N[,6] >= 3),]

TDH2 <- cbind.data.frame(TDH2.MEAN,TDH2.SD[,6:25],TDH2.N[,6])

colnames(TDH2) <- c("STRAIN","CONDITION","ASSAY","PLATE","CTRL.GENO",paste0("MEAN.",colnames(TDH2.MEAN[,6:25])),paste0("SD.",colnames(TDH2.SD[,6:25])),"N")

write.table(TDH2,"TDH2.SUMMARY.DATA_10112016_DCv34.txt",sep="\t",quote=FALSE,row.names=FALSE)

`%notin%` <- function(x,y) !(x %in% y)

CONTAM.CHECK<-droplevels(subset(CLEAN.DATA,CLEAN.DATA$CONTAM.CHECK!="NONE"))

if(all(CONTAM.CHECK$STRAIN %notin% TDH2$STRAIN)) {
  #All datapoints flagged as suspicious have been removed by outlier removal filters.
  print("Yay! Suspicious values eliminated. Proceed.")
}

if(any(CONTAM.CHECK$STRAIN %in% TDH2$STRAIN)) {
  CONTAM.CHECK<-CONTAM.CHECK[CONTAM.CHECK$STRAIN %in% CONTAM.CHECK$STRAIN[which(CONTAM.CHECK$STRAIN %in% TDH2$STRAIN)],]
  write.table(CONTAM.CHECK,"TDH2.CONTAM.CHECK_outliersexcl.csv",sep=",",quote=FALSE,row.names=FALSE)  
}


###############################
#HIERARCHICAL PERMUTATION TEST#
###############################

#Clear memory
rm(list=ls())
options(warn=-1)

library(permute)
library(Hmisc)

N.PERM <- 10000

#Permutation function

SHUFFLE <- function(STRAIN.LIST = STRAIN.LIST[[i]],WT.LIST = WT.LIST) {
  SAMPLE <- c(STRAIN.LIST,WT.LIST[[sample(1:length(WT.LIST),1)]])
  OUTPUT <- sample(SAMPLE,length(SAMPLE))
  DIST.SIM <- abs(mean(OUTPUT[1:length(STRAIN.LIST)]) - mean(OUTPUT[(length(STRAIN.LIST)+1):length(SAMPLE)]))
  DIST.OBS <- abs(mean(SAMPLE[1:length(STRAIN.LIST)]) - mean(SAMPLE[(length(STRAIN.LIST)+1):length(SAMPLE)]))
  
  DIFF <- DIST.OBS - DIST.SIM
  
  return(DIFF)
}

parent.dir <- PATH
setwd(parent.dir)

####TDH2 Analysis date 10112016#########################################################################

ALL.REP <- read.csv("TDH2.CLEAN.ADJUST_10112016_DCv34.csv",header=TRUE)
ALL.REP <- subset(ALL.REP, MEDIAN.OUTLIER == "NO" & SD.OUTLIER == "NO" & FSC.OUTLIER == "NO")

SUMMARY <- read.table("TDH2.SUMMARY.DATA_10112016_DCv34.txt",header=TRUE)

#SUMMARY <- droplevels(subset(SUMMARY, STRAIN != "1139"))

WT <- subset(ALL.REP, CONDITION == "SHAM")
LOW.FSC <- median(WT$FSC.CORRECT) - 4*mad(WT$FSC.CORRECT)
HIGH.FSC <- median(WT$FSC.CORRECT) + 4*mad(WT$FSC.CORRECT)
LOW.FLUO <- median(WT$YFP.MEDIAN.CORRECT) - 4*mad(WT$YFP.MEDIAN.CORRECT)
HIGH.FLUO <- median(WT$YFP.MEDIAN.CORRECT) + 4*mad(WT$YFP.MEDIAN.CORRECT)
WT <- subset(WT, FSC.CORRECT > LOW.FSC & FSC.CORRECT < HIGH.FSC & YFP.MEDIAN.CORRECT > LOW.FLUO & YFP.MEDIAN.CORRECT < HIGH.FLUO)

#MEDIAN

#Create list of WT phenotype grouped by strain

N.WT <- length(unique(WT$STRAIN))
WT.LIST <- vector('list',N.WT)
for (i in 1:N.WT)
{
  CUR.WT <- subset(WT, STRAIN == unique(WT$STRAIN)[i])
  WT.LIST[[i]] <- CUR.WT$YFP.MEDIAN.CORRECT 
}


#Create list of all strain phenotypes grouped by strain

SUMMARY <- SUMMARY[order(SUMMARY$STRAIN),]
ALL.REP <- droplevels(subset(ALL.REP,STRAIN %in% as.character(SUMMARY$STRAIN)))
ALL.REP <- ALL.REP[order(ALL.REP$STRAIN),]
N.STRAIN <- length(unique(ALL.REP$STRAIN))
STRAIN.LIST <- vector('list',N.STRAIN)
for (i in 1:N.STRAIN)
{
  CUR.STRAIN <- subset(ALL.REP, STRAIN == unique(ALL.REP$STRAIN)[i])
  STRAIN.LIST[[i]] <- CUR.STRAIN$YFP.MEDIAN.CORRECT
}


for (i in 1:length(STRAIN.LIST))
{
  DIFF <- sapply(1:N.PERM,FUN=function(x)	SHUFFLE(STRAIN.LIST = STRAIN.LIST[[i]],WT.LIST = WT.LIST))
  P.VAL <- (length(which(DIFF < 0)) + 1) / (length(which(DIFF < 0)) + length(which(DIFF > 0)) + 1) 	
  SUMMARY[i,"P.VAL.MEDIAN"] <- P.VAL
  print(round(i/length(STRAIN.LIST)*100,1))
}

#NOISE: CV

#Create list of WT phenotype grouped by strain

N.WT <- length(unique(WT$STRAIN))
WT.LIST <- vector('list',N.WT)
for (i in 1:N.WT)
{
  CUR.WT <- subset(WT, STRAIN == unique(WT$STRAIN)[i])
  WT.LIST[[i]] <- CUR.WT$YFP.CV.WT
}


#Create list of all strain phenotypes grouped by strain

SUMMARY <- SUMMARY[order(SUMMARY$STRAIN),]
ALL.REP <- droplevels(subset(ALL.REP,STRAIN %in% as.character(SUMMARY$STRAIN)))
ALL.REP <- ALL.REP[order(ALL.REP$STRAIN),]
N.STRAIN <- length(unique(ALL.REP$STRAIN))
STRAIN.LIST <- vector('list',N.STRAIN)
for (i in 1:N.STRAIN)
{
  CUR.STRAIN <- subset(ALL.REP, STRAIN == unique(ALL.REP$STRAIN)[i])
  STRAIN.LIST[[i]] <- CUR.STRAIN$YFP.CV.WT
}


for (i in 1:length(STRAIN.LIST))
{
  DIFF <- sapply(1:N.PERM,FUN=function(x)	SHUFFLE(STRAIN.LIST = STRAIN.LIST[[i]],WT.LIST = WT.LIST))
  P.VAL <- (length(which(DIFF < 0))+1) / (length(which(DIFF < 0)) + length(which(DIFF > 0)) + 1) 	
  SUMMARY[i,"P.VAL.CV.WT"] <- P.VAL
  print(round(i/length(STRAIN.LIST)*100,1))
}

#NOISE: FANO

#Create list of WT phenotype grouped by strain

N.WT <- length(unique(WT$STRAIN))
WT.LIST <- vector('list',N.WT)
for (i in 1:N.WT)
{
  CUR.WT <- subset(WT, STRAIN == unique(WT$STRAIN)[i])
  WT.LIST[[i]] <- CUR.WT$YFP.FANO.WT
}


#Create list of all strain phenotypes grouped by strain

SUMMARY <- SUMMARY[order(SUMMARY$STRAIN),]
ALL.REP <- droplevels(subset(ALL.REP,STRAIN %in% as.character(SUMMARY$STRAIN)))
ALL.REP <- ALL.REP[order(ALL.REP$STRAIN),]
N.STRAIN <- length(unique(ALL.REP$STRAIN))
STRAIN.LIST <- vector('list',N.STRAIN)
for (i in 1:N.STRAIN)
{
  CUR.STRAIN <- subset(ALL.REP, STRAIN == unique(ALL.REP$STRAIN)[i])
  STRAIN.LIST[[i]] <- CUR.STRAIN$YFP.FANO.WT
}


for (i in 1:length(STRAIN.LIST))
{
  DIFF <- sapply(1:N.PERM,FUN=function(x)	SHUFFLE(STRAIN.LIST = STRAIN.LIST[[i]],WT.LIST = WT.LIST))
  P.VAL <- (length(which(DIFF < 0))+1) / (length(which(DIFF < 0)) + length(which(DIFF > 0)) + 1) 	
  SUMMARY[i,"P.VAL.FANO.WT"] <- P.VAL
  print(round(i/length(STRAIN.LIST)*100,1))
}

#FSC MEDIAN

#Create list of WT phenotype grouped by strain

N.WT <- length(unique(WT$STRAIN))
WT.LIST <- vector('list',N.WT)
for (i in 1:N.WT)
{
  CUR.WT <- subset(WT, STRAIN == unique(WT$STRAIN)[i])
  WT.LIST[[i]] <- CUR.WT$FSC.CORRECT
}


#Create list of all strain phenotypes grouped by strain

SUMMARY <- SUMMARY[order(SUMMARY$STRAIN),]
ALL.REP <- droplevels(subset(ALL.REP,STRAIN %in% as.character(SUMMARY$STRAIN)))
ALL.REP <- ALL.REP[order(ALL.REP$STRAIN),]
N.STRAIN <- length(unique(ALL.REP$STRAIN))
STRAIN.LIST <- vector('list',N.STRAIN)
for (i in 1:N.STRAIN)
{
  CUR.STRAIN <- subset(ALL.REP, STRAIN == unique(ALL.REP$STRAIN)[i])
  STRAIN.LIST[[i]] <- CUR.STRAIN$FSC.CORRECT
}


for (i in 1:length(STRAIN.LIST))
{
  DIFF <- sapply(1:N.PERM,FUN=function(x)	SHUFFLE(STRAIN.LIST = STRAIN.LIST[[i]],WT.LIST = WT.LIST))
  P.VAL <- (length(which(DIFF < 0))+1) / (length(which(DIFF < 0)) + length(which(DIFF > 0)) + 1) 	
  SUMMARY[i,"P.VAL.FSC"] <- P.VAL
  print(round(i/length(STRAIN.LIST)*100,1))
}

write.table(SUMMARY,"TDH2.10112016.SUMMARY.DCv34.Pvals.txt",sep="\t",quote=FALSE,row.names=FALSE)

#_____________________________________________________________________________________________
# #VMA7
#

########################################
#  CLEANING
########################################

#Clear memory
rm(list=ls())
options(warn=-1)


#Set working directory
parent.dir <- PATH
setwd(parent.dir)

#Load experiment setup
SETUP <- read.csv("Template_VMA7_10102016.csv",header=TRUE,as.is=TRUE)
TESTNAMES <- list.files("PATH/VMA7/",pattern=".fcs",full.name=TRUE,recursive=TRUE,include.dirs=TRUE)
SETUP<-subset(SETUP, SETUP$FILENAMES %in% TESTNAMES, drop = TRUE)
SETUP[,"COUNTER"] <- c(1:nrow(SETUP))

#Determine Hard Gates
GATES <- GATE.CALIB(SETUP$FILENAMES[1])

#Clean Data
Output <- apply(SETUP,1,CLEANING,GATES=GATES)
OUTPUT <- as.data.frame(Output[[1]])
OUTPUT[2:nrow(SETUP),1:ncol(OUTPUT)] <- NA
for (i in 2:nrow(SETUP))
{
  if (is.null(Output[[i]]))
  {} 
  else {
    OUTPUT[i,] <- Output[[i]]
  }
}

write.table(OUTPUT,"VMA7_Experiment.Output_10112016_DCv34.txt",row.names=FALSE,sep="\t")

CLEAN <- cbind.data.frame(SETUP[,1:(ncol(SETUP)-2)],OUTPUT)

write.table(CLEAN,"VMA7_Clean.Data_10112016_DCv34.txt",row.names=FALSE,sep="\t")

################################################################
#     PLATE CORRECTION & OUTLIER REMOVAL
################################################################

#Clear memory
rm(list=ls())
options(warn=-1)

#####################
#1-LOADING LIBRARIES#
#####################
library(flowCore)
library(flowClust)
library(flowViz)
library(pcaPP)
library(mixtools)
library(plyr)
library(MASS)
box <- graphics::box
options(warn=-1)

####################################
#2- Quality Control and Corrections#
####################################

parent.dir <- PATH

setwd(parent.dir)

DATA.TYPE <- c(
  "factor","factor","factor","factor","factor","factor","factor","factor","factor","factor","factor","factor","factor","factor","factor",
  "integer","integer","integer","integer",
  "numeric","numeric","numeric","numeric","numeric","numeric","numeric","numeric","numeric","numeric","numeric","numeric","numeric","numeric","numeric","numeric","numeric","numeric","numeric","numeric","numeric","numeric","numeric","numeric","numeric",
  "integer")

###Read in data sets
CLEAN.DATA <- read.table("VMA7_Clean.Data_10112016_DCv34.txt",colClasses=DATA.TYPE,header=TRUE, sep="\t")

###Remove samples with NA
CLEAN.DATA <- CLEAN.DATA[complete.cases(CLEAN.DATA),]

##Translate into log scale for appropriate correction
CLEAN.DATA[,"log.RNA.MEDIAN"] <- log((CLEAN.DATA[,"YFP.MEDIAN.FINAL"]) + 0.05)
CLEAN.DATA[,"log.RNA.MAD"] <- CLEAN.DATA[,"YFP.MAD.FINAL"] / (CLEAN.DATA[,"YFP.MEDIAN.FINAL"] + 0.05)
CLEAN.DATA[,"log.RNA.SD"] <- CLEAN.DATA[,"YFP.SD.FINAL"] / (CLEAN.DATA[,"YFP.MEDIAN.FINAL"] + 0.05)

###Remove samples with less than 1000 events
CLEAN.DATA <- subset(CLEAN.DATA,CLEAN.DATA$COUNTS.FINAL > 1000)
CLEAN.DATA <- subset(CLEAN.DATA,CLEAN.DATA$PLATE != "P1")

####Add variable to ID whether control is own promoter or TDH3
CLEAN.DATA[,"CTRL.GENO"]<-as.factor("VMA7")


###Visualize spread before correction / filtering

CLEAN.DATA$CONTAM.CHECK<-"NONE"
CLEAN.DATA$CONTAM.CHECK[(CLEAN.DATA$CONDITION=="EMS"|CLEAN.DATA$CONDITION=="SHAM"|CLEAN.DATA$CONDITION=="EMS.LOW"|CLEAN.DATA$CONDITION=="EMS.HIGH") & (CLEAN.DATA$YFP.MEDIAN.FINAL<.53)]<-"BACT.CONTAM.CHECK"
CLEAN.DATA$CONTAM.CHECK[(CLEAN.DATA$CONDITION=="EMS"|CLEAN.DATA$CONDITION=="SHAM"|CLEAN.DATA$CONDITION=="EMS.LOW"|CLEAN.DATA$CONDITION=="EMS.HIGH") & (CLEAN.DATA$YFP.MEDIAN.FINAL>0.7)]<-"TDH3.CONTAM.CHECK"
CLEAN.DATA$CONTAM.CHECK[(CLEAN.DATA$CONDITION=="TDH3.SHAM" & CLEAN.DATA$YFP.MEDIAN.FINAL<0.7)]<-"EXP.SHAM.CONTAM.CHECK"
CLEAN.DATA$CONTAM.CHECK[(CLEAN.DATA$CONDITION=="NULL" & CLEAN.DATA$YFP.MEDIAN.FINAL>0.53)]<-"EXP.NULL.CONTAM.CHECK"


###Seperate controls and remove FSC outliers
CONTROL <- subset(CLEAN.DATA,CLEAN.DATA$CONDITION == "CTRL")
FSC.CORRECT <- lm(CONTROL$FSC.MEDIAN.FINAL ~ 0 + CONTROL$FLOW.RUN)
RESID <- FSC.CORRECT$resid
CLEAN.DATA <- droplevels(CLEAN.DATA[-which(CLEAN.DATA$CONDITION == "CTRL")[abs(RESID) > 3*mad(RESID)],])
CONTROL <- droplevels(subset(CLEAN.DATA,CLEAN.DATA$CONDITION == "CTRL"))

INITIAL.CORRECT <- lm(CONTROL$log.RNA.MEDIAN ~ 0 + CONTROL$FLOW.RUN)
RESID <- INITIAL.CORRECT$resid
CLEAN.DATA<- droplevels(CLEAN.DATA[-which(CLEAN.DATA$CONDITION == "CTRL")[abs(RESID) > 3*mad(RESID)],])
CONTROL <- droplevels(subset(CLEAN.DATA,CLEAN.DATA$CONDITION == "CTRL"))

`%notin%` <- function(x,y) !(x %in% y)
RESULT<-count(CONTROL$FLOW.RUN)
RESULT.MUT<-count(CLEAN.DATA$FLOW.RUN)
EXCLUDE1<-(RESULT.MUT[which(RESULT.MUT[,1] %notin% RESULT[,1]),1])
EXCLUDE2<-(RESULT[which(RESULT[,2] <5),1])
CLEAN.DATA<-droplevels(CLEAN.DATA[CLEAN.DATA$FLOW.RUN %notin% EXCLUDE1,])
CLEAN.DATA<-droplevels(CLEAN.DATA[CLEAN.DATA$FLOW.RUN %notin% EXCLUDE2,])
CONTROL <- subset(CLEAN.DATA,CLEAN.DATA$CONDITION == "CTRL")


###Corrections for YFP median and MAD
#Correct Median
#fit<- lm(log.RNA.MEDIAN~FLOW.RUN+REP+STACK.ORDER+STACK.ORDER:FLOW.RUN+REP+ROW+COLUMN+BLOCK+BLOCK:ROW+ROW:COLUMN+STACK.ORDER:BLOCK,data=CONTROL)
#step <- stepAIC(fit, direction="both")
log.RNA.MEDIAN.CORRECT <- lm(log.RNA.MEDIAN ~ FLOW.RUN + COLUMN + ROW + ROW:COLUMN, data=CONTROL)
CLEAN.DATA[,"log.RNA.MEDIAN.CORRECT"] <- CLEAN.DATA$log.RNA.MEDIAN - predict(log.RNA.MEDIAN.CORRECT, CLEAN.DATA) + mean(CONTROL$log.RNA.MEDIAN)
CONTROL <- subset(CLEAN.DATA,CLEAN.DATA$CONDITION == "CTRL")

#Correct sd
#fit<- lm(log.RNA.SD~FLOW.RUN+REP+STACK.ORDER+STACK.ORDER:FLOW.RUN+REP+ROW+COLUMN+BLOCK+BLOCK:ROW+ROW:COLUMN+STACK.ORDER:BLOCK,data=CONTROL)
#step <- step(fit, direction="both")
log.RNA.SD.CORRECT <- lm(log.RNA.SD ~  FLOW.RUN + ROW + COLUMN+ ROW:COLUMN,data=CONTROL)
CLEAN.DATA[,"log.RNA.SD.CORRECT"] <- CLEAN.DATA$log.RNA.SD - predict(log.RNA.SD.CORRECT , CLEAN.DATA) + mean(CONTROL$log.RNA.SD)
CONTROL <- subset(CLEAN.DATA,CLEAN.DATA$CONDITION == "CTRL")

#Correct mad
#fit<- lm(log.RNA.MAD~FLOW.RUN+REP+STACK.ORDER+STACK.ORDER:FLOW.RUN+REP+ROW+COLUMN+BLOCK+BLOCK:ROW+ROW:COLUMN+STACK.ORDER:BLOCK,data=CONTROL)
#step <- step(fit, direction="both")
log.RNA.MAD.CORRECT <- lm(log.RNA.MAD ~  FLOW.RUN + ROW + COLUMN+ ROW:COLUMN,data=CONTROL)
CLEAN.DATA[,"log.RNA.MAD.CORRECT"] <- CLEAN.DATA$log.RNA.MAD - predict(log.RNA.MAD.CORRECT , CLEAN.DATA) + mean(CONTROL$log.RNA.MAD)
CONTROL <- subset(CLEAN.DATA,CLEAN.DATA$CONDITION == "CTRL")

#Correct FSC
#fit<- lm(FSC.MEDIAN.FINAL~FLOW.RUN+REP+STACK.ORDER+STACK.ORDER:FLOW.RUN+REP+ROW+COLUMN+BLOCK+BLOCK:ROW+ROW:COLUMN+STACK.ORDER:BLOCK,data=CONTROL)
#step <- stepAIC(fit, direction="both")
FSC.CORRECT <- lm(FSC.MEDIAN.FINAL ~  FLOW.RUN + ROW + COLUMN,data=CONTROL)
CLEAN.DATA[,"FSC.CORRECT"] <- CLEAN.DATA$FSC.MEDIAN.FINAL - predict(FSC.CORRECT, CLEAN.DATA) + mean(CONTROL$FSC.MEDIAN.FINAL)
CONTROL <- subset(CLEAN.DATA,CLEAN.DATA$CONDITION == "CTRL")

#Back translate to linear scale
CLEAN.DATA[,"YFP.MEDIAN.CORRECT"] <- (exp(CLEAN.DATA[,"log.RNA.MEDIAN.CORRECT"]) - 0.05)
CLEAN.DATA[,"YFP.SD.CORRECT"] <- CLEAN.DATA[,"log.RNA.SD.CORRECT"] * CLEAN.DATA[,"YFP.MEDIAN.CORRECT"]
CLEAN.DATA[,"YFP.MAD.CORRECT"] <- CLEAN.DATA[,"log.RNA.MAD.CORRECT"] * CLEAN.DATA[,"YFP.MEDIAN.CORRECT"]


#PERFORM FABIEN'S OUTLIER CORRECTION FILTERING OUT ANY REPLICATE OUTSIDE OF 4*MAD

#Remove outliers
for (i in 1:nrow(CLEAN.DATA))
{
  CUR <- subset(CLEAN.DATA, STRAIN == CLEAN.DATA[i,"STRAIN"])
  if (CLEAN.DATA[i,"YFP.MEDIAN.CORRECT"] > median(CUR$YFP.MEDIAN.CORRECT) + 4*mad(CUR$YFP.MEDIAN.CORRECT) | CLEAN.DATA[i,"YFP.MEDIAN.CORRECT"] < median(CUR$YFP.MEDIAN.CORRECT) - 4*mad(CUR$YFP.MEDIAN.CORRECT)) {
    CLEAN.DATA[i,"MEDIAN.OUTLIER"] <- "YES"
  } else {
    CLEAN.DATA[i,"MEDIAN.OUTLIER"] <- "NO"
  }
  if (CLEAN.DATA[i,"YFP.SD.CORRECT"] > median(CUR$YFP.SD.CORRECT) + 4*mad(CUR$YFP.SD.CORRECT) | CLEAN.DATA[i,"YFP.SD.CORRECT"] < median(CUR$YFP.SD.CORRECT) - 4*mad(CUR$YFP.SD.CORRECT)) {
    CLEAN.DATA[i,"SD.OUTLIER"] <- "YES"
  } else {
    CLEAN.DATA[i,"SD.OUTLIER"] <- "NO"
  }
  if (CLEAN.DATA[i,"FSC.CORRECT"] > median(CUR$FSC.CORRECT) + 4*mad(CUR$FSC.CORRECT) | CLEAN.DATA[i,"FSC.CORRECT"] < median(CUR$FSC.CORRECT) - 4*mad(CUR$FSC.CORRECT)) {
    CLEAN.DATA[i,"FSC.OUTLIER"] <- "YES"
  } else {
    CLEAN.DATA[i,"FSC.OUTLIER"] <- "NO"
  }	
}

CLEAN.DATA <- subset(CLEAN.DATA, MEDIAN.OUTLIER == "NO" & SD.OUTLIER == "NO" & FSC.OUTLIER == "NO")

#Write processed data to file
write.table(CLEAN.DATA,"VMA7.CLEAN.CORRECT_10112016_DCv34.csv",sep=",",quote=FALSE,row.names=FALSE)

#####################
#5 - Control plots  #
#####################

CONTROL <- droplevels(subset(CLEAN.DATA,CLEAN.DATA$CONDITION == "CTRL"))
SHAM <- droplevels(subset(CLEAN.DATA,CLEAN.DATA$CONDITION == "SHAM"))
TDH3.SHAM <- droplevels(subset(CLEAN.DATA,CLEAN.DATA$CONDITION == "TDH3.SHAM"))


###############################################################################
#
#       Strain Estimates
#
################################################################################

# Clear memory
rm(list=ls())
options(warn=-1)

setwd("E:/wittkopp data/processed data/RNA scale analyses/DCv34")

#VMA7
CLEAN.DATA <- read.csv("VMA7.CLEAN.CORRECT_10112016_DCv34.csv", header=TRUE)

#Calculate mean and noise for each sample relative to controls
TDH3.SHAM <- subset(CLEAN.DATA, CLEAN.DATA$CONDITION == "TDH3.SHAM")
NEG <- subset(CLEAN.DATA, CLEAN.DATA$CONDITION == "NULL")
WT.SHAM <- subset(CLEAN.DATA, CLEAN.DATA$CONDITION == "SHAM")

#ABSOLUTE EXPRESSION SCALE
CLEAN.DATA[,"YFP.MEDIAN.ABS"] <- (CLEAN.DATA$YFP.MEDIAN.CORRECT - median(NEG$YFP.MEDIAN.CORRECT))
CLEAN.DATA[,"YFP.SD.ABS"]   <- CLEAN.DATA$YFP.SD.CORRECT - median(NEG$YFP.SD.CORRECT)
CLEAN.DATA[,"YFP.MAD.ABS"]   <- CLEAN.DATA$YFP.MAD.CORRECT - median(NEG$YFP.SD.CORRECT)

#Re-calculate mean and noise for each sample relative to controls
TDH3.SHAM <- subset(CLEAN.DATA, CLEAN.DATA$CONDITION == "TDH3.SHAM")
NEG <- subset(CLEAN.DATA, CLEAN.DATA$CONDITION == "NULL")
WT.SHAM <- subset(CLEAN.DATA, CLEAN.DATA$CONDITION == "SHAM")

#PROMOTER-SPECIFIC SCALING
CLEAN.DATA[,"YFP.MEDIAN.WT"] <- CLEAN.DATA$YFP.MEDIAN.ABS/median(WT.SHAM$YFP.MEDIAN.ABS)
CLEAN.DATA[,"YFP.SD.WT"]   <- CLEAN.DATA$YFP.SD.ABS/median(WT.SHAM$YFP.MEDIAN.ABS)
CLEAN.DATA[,"YFP.MAD.WT"]   <- CLEAN.DATA$YFP.MAD.ABS/median(WT.SHAM$YFP.MEDIAN.ABS)

#RELATIVE TO TDH3 EXPRESSION
CLEAN.DATA[,"YFP.MEDIAN.TDH3"] <- CLEAN.DATA$YFP.MEDIAN.ABS/median(TDH3.SHAM$YFP.MEDIAN.ABS)
CLEAN.DATA[,"YFP.SD.TDH3"]   <- CLEAN.DATA$YFP.SD.ABS/median(TDH3.SHAM$YFP.MEDIAN.ABS)
CLEAN.DATA[,"YFP.MAD.TDH3"]   <- CLEAN.DATA$YFP.MAD.ABS/median(TDH3.SHAM$YFP.MEDIAN.ABS)

#Re-calculate mean and noise for each sample relative to controls
TDH3.SHAM <- subset(CLEAN.DATA, CLEAN.DATA$CONDITION == "TDH3.SHAM")
TDH3.SHAM<-subset(TDH3.SHAM,TDH3.SHAM$YFP.MEDIAN.FINAL>0.7)
NEG <- subset(CLEAN.DATA, CLEAN.DATA$CONDITION == "NULL")
WT.SHAM <- subset(CLEAN.DATA, CLEAN.DATA$CONDITION == "SHAM")

#CALCULATE NOISE METRICS
CLEAN.DATA[,"YFP.CV.WT"]   <- CLEAN.DATA$YFP.SD.WT/CLEAN.DATA$YFP.MEDIAN.WT
CLEAN.DATA[,"YFP.CV.TDH3"]   <- CLEAN.DATA$YFP.SD.TDH3/CLEAN.DATA$YFP.MEDIAN.TDH3
CLEAN.DATA[,"YFP.FANO.WT"]   <- CLEAN.DATA$YFP.SD.WT^2/CLEAN.DATA$YFP.MEDIAN.WT
CLEAN.DATA[,"YFP.FANO.TDH3"]   <- CLEAN.DATA$YFP.SD.TDH3^2/CLEAN.DATA$YFP.MEDIAN.TDH3


#Re-calculate mean and noise for each sample relative to controls
TDH3.SHAM <- subset(CLEAN.DATA, CLEAN.DATA$CONDITION == "TDH3.SHAM")
TDH3.SHAM<-subset(TDH3.SHAM,TDH3.SHAM$YFP.MEDIAN.FINAL>0.7)
NEG <- subset(CLEAN.DATA, CLEAN.DATA$CONDITION == "NULL")
WT.SHAM <- subset(CLEAN.DATA, CLEAN.DATA$CONDITION == "SHAM")

#CALCULATE RELATIVE NOISE METRICS
CLEAN.DATA[,"YFP.SD.WT.REL"]   <- CLEAN.DATA$YFP.SD.WT/median(WT.SHAM$YFP.SD.WT)
CLEAN.DATA[,"YFP.SD.TDH3.REL"]   <- CLEAN.DATA$YFP.SD.TDH3/median(WT.SHAM$YFP.SD.TDH3)
CLEAN.DATA[,"YFP.CV.WT.REL"]   <- CLEAN.DATA$YFP.SD.WT.REL/CLEAN.DATA$YFP.MEDIAN.WT
CLEAN.DATA[,"YFP.CV.TDH3.REL"]   <- CLEAN.DATA$YFP.SD.TDH3.REL/CLEAN.DATA$YFP.MEDIAN.TDH3
CLEAN.DATA[,"YFP.FANO.WT.REL"]   <- CLEAN.DATA$YFP.SD.WT.REL^2/CLEAN.DATA$YFP.MEDIAN.WT
CLEAN.DATA[,"YFP.FANO.TDH3.REL"]   <- CLEAN.DATA$YFP.SD.TDH3.REL^2/CLEAN.DATA$YFP.MEDIAN.TDH3


#Add NA for samples where CV or FANO are undefined.
for (i in 1:nrow(CLEAN.DATA))
{
  if (CLEAN.DATA[i,"YFP.MEDIAN.ABS"] < 0 | CLEAN.DATA[i,"YFP.SD.ABS"] < 0) {
    CLEAN.DATA[i,"YFP.MAD.WT"] <- NA
    CLEAN.DATA[i,"YFP.MAD.TDH3"] <- NA
    CLEAN.DATA[i,"YFP.CV.WT"] <- NA
    CLEAN.DATA[i,"YFP.CV.TDH3"] <- NA
    CLEAN.DATA[i,"YFP.CV.WT.REL"] <- NA
    CLEAN.DATA[i,"YFP.CV.TDH3.REL"] <- NA
    CLEAN.DATA[i,"YFP.FANO.WT"] <- NA
    CLEAN.DATA[i,"YFP.FANO.TDH3"] <- NA
    CLEAN.DATA[i,"YFP.FANO.WT.REL"] <- NA
    CLEAN.DATA[i,"YFP.FANO.TDH3.REL"] <- NA
    
  }
}

CLEAN.DATA <- droplevels(subset(CLEAN.DATA,CLEAN.DATA$CONDITION!="CTRL"))

write.table(CLEAN.DATA,"VMA7.CLEAN.ADJUST_10112016_DCv34.csv",sep=",",quote=FALSE,row.names=FALSE)

CLEAN.DATA <- read.csv("VMA7.CLEAN.ADJUST_10112016_DCv34.csv", header=TRUE)

VMA7.MEAN <- aggregate(cbind(YFP.MEDIAN.ABS,YFP.SD.ABS,YFP.MAD.ABS,YFP.MEDIAN.WT,YFP.SD.WT,YFP.MAD.WT,YFP.MEDIAN.TDH3,YFP.SD.TDH3,YFP.MAD.TDH3,YFP.CV.WT,YFP.FANO.WT,YFP.CV.TDH3,YFP.FANO.TDH3,YFP.SD.WT.REL,YFP.CV.WT.REL,YFP.FANO.WT.REL,YFP.SD.TDH3.REL,YFP.CV.TDH3.REL,YFP.FANO.TDH3.REL,FSC.CORRECT) ~ STRAIN + CONDITION + ASSAY + PLATE + CTRL.GENO + CONTAM.CHECK, data=CLEAN.DATA, FUN = mean)
VMA7.SD <- aggregate(cbind(YFP.MEDIAN.ABS,YFP.SD.ABS,YFP.MAD.ABS,YFP.MEDIAN.WT,YFP.SD.WT,YFP.MAD.WT,YFP.MEDIAN.TDH3,YFP.SD.TDH3,YFP.MAD.TDH3,YFP.CV.WT,YFP.FANO.WT,YFP.CV.TDH3,YFP.FANO.TDH3,YFP.SD.WT.REL,YFP.CV.WT.REL,YFP.FANO.WT.REL,YFP.SD.TDH3.REL,YFP.CV.TDH3.REL,YFP.FANO.TDH3.REL,FSC.CORRECT) ~ STRAIN + CONDITION + ASSAY + PLATE + CTRL.GENO + CONTAM.CHECK, data=CLEAN.DATA, FUN = sd)
VMA7.N <- aggregate(cbind(YFP.MEDIAN.ABS,YFP.SD.ABS,YFP.MAD.ABS,YFP.MEDIAN.WT,YFP.SD.WT,YFP.MAD.WT,YFP.MEDIAN.TDH3,YFP.SD.TDH3,YFP.MAD.TDH3,YFP.CV.WT,YFP.FANO.WT,YFP.CV.TDH3,YFP.FANO.TDH3,YFP.SD.WT.REL,YFP.CV.WT.REL,YFP.FANO.WT.REL,YFP.SD.TDH3.REL,YFP.CV.TDH3.REL,YFP.FANO.TDH3.REL,FSC.CORRECT) ~ STRAIN + CONDITION + ASSAY + PLATE + CTRL.GENO + CONTAM.CHECK, data=CLEAN.DATA, FUN = length)

VMA7.MEAN  <- VMA7.MEAN[which(VMA7.N[,7] >= 3),]
VMA7.SD <- VMA7.SD[which(VMA7.N[,7] >= 3),]
VMA7.N <- VMA7.N[which(VMA7.N[,7] >= 3),]

VMA7 <- cbind.data.frame(VMA7.MEAN,VMA7.SD[,7:26],VMA7.N[,7])

colnames(VMA7) <- c("STRAIN","CONDITION","ASSAY","PLATE","CTRL.GENO","CONTAM.CHECK",paste0("MEAN.",colnames(VMA7.MEAN[,6:25])),paste0("SD.",colnames(VMA7.SD[,6:25])),"N")

if(sum(VMA7$CONTAM.CHECK=="NONE")==length(VMA7$CONTAM.CHECK)) {
  #All datapoints flagged as suspicious have been removed by outlier removal filters.
  print("Yay! Suspicious values eliminated. Proceed.")
}

if(sum(VMA7$CONTAM.CHECK=="NONE")!=length(VMA7$CONTAM.CHECK)) {
  EXCLUDE.LIST<-read.table("CONTAM.CHECK.EXCLUDE.STRAIN.NAMES.txt")
  CONTAM.CHECK<-CONTAM.CHECK[CONTAM.CHECK$STRAIN %in% CONTAM.CHECK$STRAIN[which(CONTAM.CHECK$STRAIN %notin% EXCLUDE)],]
  
  write.table(CONTAM.CHECK,"VMA7.CONTAM.CHECK_outliersexcl.csv",sep=",",quote=FALSE,row.names=FALSE)  
}
write.table(VMA7,"VMA7.SUMMARY.DATA_10112016_DCv34.txt",sep="\t",quote=FALSE,row.names=FALSE)


###############################
#HIERARCHICAL PERMUTATION TEST#
###############################

#Clear memory
rm(list=ls())
options(warn=-1)

library(permute)
library(Hmisc)

N.PERM <- 10000

#Permutation function

SHUFFLE <- function(STRAIN.LIST = STRAIN.LIST[[i]],WT.LIST = WT.LIST) {
  SAMPLE <- c(STRAIN.LIST,WT.LIST[[sample(1:length(WT.LIST),1)]])
  OUTPUT <- sample(SAMPLE,length(SAMPLE))
  DIST.SIM <- abs(mean(OUTPUT[1:length(STRAIN.LIST)]) - mean(OUTPUT[(length(STRAIN.LIST)+1):length(SAMPLE)]))
  DIST.OBS <- abs(mean(SAMPLE[1:length(STRAIN.LIST)]) - mean(SAMPLE[(length(STRAIN.LIST)+1):length(SAMPLE)]))
  
  DIFF <- DIST.OBS - DIST.SIM
  
  return(DIFF)
}

 parent.dir <- "PATH"
 
 setwd(parent.dir)
 
 ####VMA7 Analysis date 10112016#########################################################################
 
ALL.REP <- read.csv("VMA7.CLEAN.ADJUST_10112016_DCv34.csv",header=TRUE)
ALL.REP <- subset(ALL.REP, MEDIAN.OUTLIER == "NO" & SD.OUTLIER == "NO" & FSC.OUTLIER == "NO")

SUMMARY <- read.table("VMA7.SUMMARY.DATA_10112016_DCv34.txt",header=TRUE)

#SUMMARY <- droplevels(subset(SUMMARY, STRAIN != "1139"))

WT <- subset(ALL.REP, CONDITION == "SHAM")
LOW.FSC <- median(WT$FSC.CORRECT) - 4*mad(WT$FSC.CORRECT)
HIGH.FSC <- median(WT$FSC.CORRECT) + 4*mad(WT$FSC.CORRECT)
LOW.FLUO <- median(WT$YFP.MEDIAN.CORRECT) - 4*mad(WT$YFP.MEDIAN.CORRECT)
HIGH.FLUO <- median(WT$YFP.MEDIAN.CORRECT) + 4*mad(WT$YFP.MEDIAN.CORRECT)
WT <- subset(WT, FSC.CORRECT > LOW.FSC & FSC.CORRECT < HIGH.FSC & YFP.MEDIAN.CORRECT > LOW.FLUO & YFP.MEDIAN.CORRECT < HIGH.FLUO)

#MEDIAN

#Create list of WT phenotype grouped by strain

N.WT <- length(unique(WT$STRAIN))
WT.LIST <- vector('list',N.WT)
for (i in 1:N.WT)
{
  CUR.WT <- subset(WT, STRAIN == unique(WT$STRAIN)[i])
  WT.LIST[[i]] <- CUR.WT$YFP.MEDIAN.CORRECT
}

 
#Create list of all strain phenotypes grouped by strain
 
SUMMARY <- SUMMARY[order(SUMMARY$STRAIN),]
ALL.REP <- droplevels(subset(ALL.REP,STRAIN %in% as.character(SUMMARY$STRAIN)))
ALL.REP <- ALL.REP[order(ALL.REP$STRAIN),]
N.STRAIN <- length(unique(ALL.REP$STRAIN))
STRAIN.LIST <- vector('list',N.STRAIN)
for (i in 1:N.STRAIN)
{
  CUR.STRAIN <- subset(ALL.REP, STRAIN == unique(ALL.REP$STRAIN)[i])
  STRAIN.LIST[[i]] <- CUR.STRAIN$YFP.MEDIAN.CORRECT
}


for (i in 1:length(STRAIN.LIST))
{
  DIFF <- sapply(1:N.PERM,FUN=function(x)	SHUFFLE(STRAIN.LIST = STRAIN.LIST[[i]],WT.LIST = WT.LIST))
  P.VAL <- (length(which(DIFF < 0)) + 1) / (length(which(DIFF < 0)) + length(which(DIFF > 0)) + 1)
  SUMMARY[i,"P.VAL.MEDIAN"] <- P.VAL
  print(round(i/length(STRAIN.LIST)*100,1))
}

#NOISE: CV

#Create list of WT phenotype grouped by strain

N.WT <- length(unique(WT$STRAIN))
WT.LIST <- vector('list',N.WT)
for (i in 1:N.WT)
{
  CUR.WT <- subset(WT, STRAIN == unique(WT$STRAIN)[i])
  WT.LIST[[i]] <- CUR.WT$YFP.CV.WT
}


#Create list of all strain phenotypes grouped by strain

SUMMARY <- SUMMARY[order(SUMMARY$STRAIN),]
ALL.REP <- droplevels(subset(ALL.REP,STRAIN %in% as.character(SUMMARY$STRAIN)))
ALL.REP <- ALL.REP[order(ALL.REP$STRAIN),]
N.STRAIN <- length(unique(ALL.REP$STRAIN))
STRAIN.LIST <- vector('list',N.STRAIN)
for (i in 1:N.STRAIN)
{
  CUR.STRAIN <- subset(ALL.REP, STRAIN == unique(ALL.REP$STRAIN)[i])
  STRAIN.LIST[[i]] <- CUR.STRAIN$YFP.CV.WT
}


for (i in 1:length(STRAIN.LIST))
{
  DIFF <- sapply(1:N.PERM,FUN=function(x)	SHUFFLE(STRAIN.LIST = STRAIN.LIST[[i]],WT.LIST = WT.LIST))
  P.VAL <- (length(which(DIFF < 0))+1) / (length(which(DIFF < 0)) + length(which(DIFF > 0)) + 1)
  SUMMARY[i,"P.VAL.CV.WT"] <- P.VAL
  print(round(i/length(STRAIN.LIST)*100,1))
}

#NOISE: FANO

#Create list of WT phenotype grouped by strain

N.WT <- length(unique(WT$STRAIN))
WT.LIST <- vector('list',N.WT)
for (i in 1:N.WT)
{
  CUR.WT <- subset(WT, STRAIN == unique(WT$STRAIN)[i])
  WT.LIST[[i]] <- CUR.WT$YFP.FANO.WT
}


#Create list of all strain phenotypes grouped by strain

SUMMARY <- SUMMARY[order(SUMMARY$STRAIN),]
ALL.REP <- droplevels(subset(ALL.REP,STRAIN %in% as.character(SUMMARY$STRAIN)))
ALL.REP <- ALL.REP[order(ALL.REP$STRAIN),]
N.STRAIN <- length(unique(ALL.REP$STRAIN))
STRAIN.LIST <- vector('list',N.STRAIN)
for (i in 1:N.STRAIN)
{
  CUR.STRAIN <- subset(ALL.REP, STRAIN == unique(ALL.REP$STRAIN)[i])
  STRAIN.LIST[[i]] <- CUR.STRAIN$YFP.FANO.WT
}


for (i in 1:length(STRAIN.LIST))
{
  DIFF <- sapply(1:N.PERM,FUN=function(x)	SHUFFLE(STRAIN.LIST = STRAIN.LIST[[i]],WT.LIST = WT.LIST))
  P.VAL <- (length(which(DIFF < 0))+1) / (length(which(DIFF < 0)) + length(which(DIFF > 0)) + 1)
  SUMMARY[i,"P.VAL.FANO.WT"] <- P.VAL
  print(round(i/length(STRAIN.LIST)*100,1))
}

#FSC MEDIAN

#Create list of WT phenotype grouped by strain

N.WT <- length(unique(WT$STRAIN))
WT.LIST <- vector('list',N.WT)
for (i in 1:N.WT)
{
  CUR.WT <- subset(WT, STRAIN == unique(WT$STRAIN)[i])
  WT.LIST[[i]] <- CUR.WT$FSC.CORRECT
}


#Create list of all strain phenotypes grouped by strain

SUMMARY <- SUMMARY[order(SUMMARY$STRAIN),]
ALL.REP <- droplevels(subset(ALL.REP,STRAIN %in% as.character(SUMMARY$STRAIN)))
ALL.REP <- ALL.REP[order(ALL.REP$STRAIN),]
N.STRAIN <- length(unique(ALL.REP$STRAIN))
STRAIN.LIST <- vector('list',N.STRAIN)
for (i in 1:N.STRAIN)
{
  CUR.STRAIN <- subset(ALL.REP, STRAIN == unique(ALL.REP$STRAIN)[i])
  STRAIN.LIST[[i]] <- CUR.STRAIN$FSC.CORRECT
}


for (i in 1:length(STRAIN.LIST))
{
  DIFF <- sapply(1:N.PERM,FUN=function(x)	SHUFFLE(STRAIN.LIST = STRAIN.LIST[[i]],WT.LIST = WT.LIST))
  P.VAL <- (length(which(DIFF < 0))+1) / (length(which(DIFF < 0)) + length(which(DIFF > 0)) + 1)
  SUMMARY[i,"P.VAL.FSC"] <- P.VAL
  print(round(i/length(STRAIN.LIST)*100,1))
}

write.table(SUMMARY,"VMA7.SUMMARY.DCv34.Pvals.txt",sep="\t",quote=FALSE,row.names=FALSE)
```
