## Supplemental tables, figures, files, and methods for "Empirical measures of mutational effects define neutral models of regulatory evolution in *Saccharomyces cerevisiae*": Supplementary File 4.R

```
#######################################################################
# Hodgins-Davis, Duveau, Walker, and Wittkopp. 2019.  
#
# Supplementary File 4
# includes R functions for analyzing processed flow cytometry data 
# to compare mutational effects among different promoters
#
# To reproduce results presented in the manuscript, you can simply load the processed data
# provided with the manuscript, or you can use Supplementary Files 2 and 3 to generate that
# processed data and then assemble it in a common format by uncommenting the first section 
# of code below. 
#
#This code was written for R version 3.3.3.

#Clear memory
rm(list=ls())
options(warn=-1)

setwd("PATH")


library(RColorBrewer)
library(vioplot)
library(plyr)
library(kSamples)
library(ggplot2)
require(gridExtra)
library(forcats)
library(grid)
library(e1071)
library(robustbase)
library(FactoMineR)

col_list<-c("red3", "darkorange1", "gold", "#63BF4D", "springgreen4", "#4DBFBF", "#4D7ABF", "royalblue4", "#A84DBF", "maroon")

#Assemble different strain data into a common data frame

#To generate the dataset
# #Oct 2016 DATA
# 
# GPD1 <- read.table("GPD1.SUMMARY.DCv34.Pvals.txt", header=TRUE); GPD1$SORT<-"SORT1"; GPD1$SCORE<-"SC1";
# OST1 <- read.table("OST1.SUMMARY.DCv34.Pvals.txt", header=TRUE); OST1$SORT<-"SORT1"; OST1$SCORE<-"SC1";
# PFY1 <- read.table("PFY1.SUMMARY.10112016.DCv34.Pvals.txt", header=TRUE); PFY1$SORT<-"SORT1"; PFY1$SCORE<-"SC1";
# RNR1 <- read.table("RNR1.SUMMARY.DCv34.Pvals_RNR1ctrl.txt", header=TRUE); RNR1$SORT<-"SORT1"; RNR1$SCORE<-"SC1";
# RNR2 <- read.table("RNR2.SUMMARY.DCv34.Pvals.txt", header=TRUE); RNR2$SORT<-"SORT1"; RNR2$SCORE<-"SC1";
# STM1 <- read.table("STM1.SUMMARY.DCv34.Pvals.txt", header=TRUE); STM1$SORT<-"SORT1"; STM1$SCORE<-"SC1";
# TDH1 <- read.table("TDH1.10112016.SUMMARY.DCv34.Pvals.txt", header=TRUE);TDH1$ASSAY<-"TDH1";TDH1$SORT<-"SORT1";TDH1$SCORE<-"SC2";
# TDH2 <- read.table("TDH2.10112016.SUMMARY.DCv34.Pvals.txt", header=TRUE); TDH2$ASSAY<-"TDH2";TDH2$SORT<-"SORT1";TDH2$SCORE<-"SC2"
# VMA7 <- read.table("VMA7.SUMMARY.DCv34.Pvals.txt", header=TRUE); VMA7$SORT<-"SORT1"; VMA7$SCORE<-"SC1";
# 
# # Remove known contaminated samples
# RNR1 <- droplevels(subset(RNR1, !(RNR1$CONDITION=="SHAM"&RNR1$MEAN.YFP.MEDIAN.TDH3>0.9)))
# TDH1<-droplevels(subset(TDH1,!(TDH1$MEAN.YFP.MEDIAN.WT>5 & TDH1$CONDITION=="EMS")))
# 
# ALL.PROMOTERS<-rbind(TDH1,TDH2,PFY1,VMA7,STM1,RNR1,RNR2,GPD1,OST1)
# 
# ALL.PROMOTERS$ASSAY<-as.factor(ALL.PROMOTERS$ASSAY);ALL.PROMOTERS$SORT<-as.factor(ALL.PROMOTERS$SORT);ALL.PROMOTERS$SCORE<-as.factor(ALL.PROMOTERS$SCORE);
# ALL.PROMOTERS$EXP<-as.factor(paste(ALL.PROMOTERS$ASSAY,ALL.PROMOTERS$SORT,ALL.PROMOTERS$SCORE,sep="."))
# 
# ##Select whether to include CONTAM CHECK flagged samples
# ALL.PROMOTERS<-droplevels(subset(ALL.PROMOTERS,ALL.PROMOTERS$CONTAM.CHECK=="NONE"))
# 
# #Read in TDH3 data and put it in a common format
# TDH3.SORT1 <- read.table("TDH3.DATA.FROM.FABIEN/SUMMARY.DATA.XP1.txt", header=TRUE)
# TDH3.SORT1$ASSAY<-"TDH3"; TDH3.SORT1$SORT<-"SORT1";TDH3.SORT1$SCORE<-"SC1";TDH3.SORT1$CONTAM.CHECK<-"NONE";TDH3.SORT1$CTRL.GENO<-"TDH3";
# TDH3.SORT1$CONDITION<-as.factor(paste(TDH3.SORT1$TREATMENT,TDH3.SORT1$CLASS,sep="."))
# TDH3.SORT1$PLATE<-as.factor(paste0("P",TDH3.SORT1$PLATE))
# TDH3.SORT1<-droplevels(subset(TDH3.SORT1,TDH3.SORT1$CONDITION=="EMS.ALL"|TDH3.SORT1$CONDITION=="EMS.LOW"|TDH3.SORT1$CONDITION=="EMS.HIGH"|TDH3.SORT1$CONDITION=="SHAM.ALL"|TDH3.SORT1$CONDITION=="SHAM.WT"))
# TDH3.SORT1$CONDITION<-revalue(TDH3.SORT1$CONDITION, c("EMS.ALL"="EMS", "SHAM.ALL"="SHAM","SHAM.WT"="WT"))
# TDH3.SORT1<-TDH3.SORT1[ ,c("STRAIN","CONDITION","ASSAY","PLATE","CTRL.GENO","CONTAM.CHECK","SORT","SCORE","YFP.MEDIAN.CORRECT.MEAN","YFP.SD.CORRECT.MEAN","YFP.MEDIAN.RELATIVE.MEAN","YFP.MEDIAN.RELATIVE.MEAN",
#                            "FSC.MEDIAN.RELATIVE.MEAN","YFP.SD.SCALED.MEAN","YFP.SD.SCALED.MEAN", "YFP.CV.MEAN","YFP.CV.MEAN","YFP.CV.RELATIVE.MEAN",  "YFP.CV.RELATIVE.MEAN", 
#                            "YFP.SD.RELATIVE.MEAN","YFP.SD.RELATIVE.MEAN", "YFP.MEDIAN.CORRECT.SD",  
#                            "YFP.SD.CORRECT.SD","YFP.MEDIAN.RELATIVE.SD","YFP.MEDIAN.RELATIVE.SD","FSC.MEDIAN.RELATIVE.SD","YFP.SD.SCALED.SD",  
#                            "YFP.CV.SD","YFP.CV.SD","YFP.CV.RELATIVE.SD","YFP.CV.RELATIVE.SD","YFP.SD.RELATIVE.SD","YFP.SD.RELATIVE.SD", "N", "P.VAL.MEDIAN","P.VAL.NOISE")]
# colnames(TDH3.SORT1)<-c("STRAIN","CONDITION","ASSAY","PLATE","CTRL.GENO","CONTAM.CHECK","SORT","SCORE","MEAN.YFP.MEDIAN.ABS","MEAN.YFP.SD.ABS","MEAN.YFP.MEDIAN.WT","MEAN.YFP.MEDIAN.TDH3",
#                         "MEAN.FSC.CORRECT","MEAN.YFP.SD.WT.REL","MEAN.YFP.SD.TDH3.REL", "MEAN.YFP.CV.WT","MEAN.YFP.CV.TDH3","MEAN.YFP.CV.WT.REL","MEAN.YFP.CV.TDH3.REL",  
#                         "MEAN.YFP.SD.WT.REL","MEAN.YFP.SD.TDH3.REL","SD.YFP.MEDIAN.ABS",   
#                         "SD.YFP.SD.ABS","SD.YFP.MEDIAN.WT","SD.YFP.MEDIAN.TDH3","SD.FSC.CORRECT","SD.YFP.SD.SCALED",  
#                         "SD.YFP.CV.WT","SD.YFP.CV.TDH3","SD.YFP.CV.WT.REL","SD.YFP.CV.TDH3.REL","SD.YFP.SD.WT.REL","SD.YFP.SD.TDH3.REL", "N","P.VAL.MEDIAN","P.VAL.NOISE")
# 
# TDH3.SORT2 <- read.table("C:/Users/andreahd/My Documents/working analyses/2017 analyses/DCv34/TDH3.XP2.SUMMARY.Pvals.txt", header=TRUE)
# TDH3.SORT2$ASSAY<-"TDH3";TDH3.SORT2$SORT<-"SORT2";TDH3.SORT2$SCORE<-"SC1";TDH3.SORT2$CONTAM.CHECK<-"NONE";TDH3.SORT2$CTRL.GENO<-"TDH3"
# TDH3.SORT2$CONDITION<-as.factor(TDH3.SORT2$CLASS)
# TDH3.SORT2$PLATE<-as.factor(paste0("P",TDH3.SORT2$PLATE))
# TDH3.SORT2<-droplevels(subset(TDH3.SORT2,TDH3.SORT2$CONDITION=="EMS"|TDH3.SORT2$CONDITION=="SHAM"))
# TDH3.SORT2<-TDH3.SORT2[ ,c("STRAIN","CONDITION","ASSAY","PLATE","CTRL.GENO","CONTAM.CHECK","SORT","SCORE","YFP.MEDIAN.CORRECT.MEAN","YFP.SD.CORRECT.MEAN","YFP.MEDIAN.RELATIVE.MEAN","YFP.MEDIAN.RELATIVE.MEAN",
#                            "FSC.MEDIAN.RELATIVE.MEAN","YFP.SD.SCALED.MEAN","YFP.SD.SCALED.MEAN", "YFP.CV.MEAN","YFP.CV.MEAN","YFP.CV.RELATIVE.MEAN",  "YFP.CV.RELATIVE.MEAN", 
#                            "YFP.SD.RELATIVE.MEAN","YFP.SD.RELATIVE.MEAN", "YFP.MEDIAN.CORRECT.SD", 
#                            "YFP.SD.CORRECT.SD","YFP.MEDIAN.RELATIVE.SD","YFP.MEDIAN.RELATIVE.SD","FSC.MEDIAN.RELATIVE.SD","YFP.SD.SCALED.SD",  
#                            "YFP.CV.SD","YFP.CV.SD","YFP.CV.RELATIVE.SD","YFP.CV.RELATIVE.SD","YFP.SD.RELATIVE.SD","YFP.SD.RELATIVE.SD", "N","P.VAL.MEDIAN","P.VAL.CV.WT","P.VAL.FANO.WT","P.VAL.FSC")]
# colnames(TDH3.SORT2)<-c("STRAIN","CONDITION","ASSAY","PLATE","CTRL.GENO","CONTAM.CHECK","SORT","SCORE","MEAN.YFP.MEDIAN.ABS","MEAN.YFP.SD.ABS","MEAN.YFP.MEDIAN.WT","MEAN.YFP.MEDIAN.TDH3",
#                         "MEAN.FSC.CORRECT","MEAN.YFP.SD.WT.REL","MEAN.YFP.SD.TDH3.REL", "MEAN.YFP.CV.WT","MEAN.YFP.CV.TDH3","MEAN.YFP.CV.WT.REL","MEAN.YFP.CV.TDH3.REL",  
#                         "MEAN.YFP.SD.WT.REL","MEAN.YFP.SD.TDH3.REL","SD.YFP.MEDIAN.ABS",   
#                         "SD.YFP.SD.ABS","SD.YFP.MEDIAN.WT","SD.YFP.MEDIAN.TDH3","SD.FSC.CORRECT","SD.YFP.SD.SCALED",  
#                         "SD.YFP.CV.WT","SD.YFP.CV.TDH3","SD.YFP.CV.WT.REL","SD.YFP.CV.TDH3.REL","SD.YFP.SD.WT.REL","SD.YFP.SD.TDH3.REL", "N","P.VAL.MEDIAN","P.VAL.CV.WT","P.VAL.FANO.WT","P.VAL.FSC")
# 
# ##JOIN ALL DATASETS
# ALLP.WTDH3.COLNAMES<-c("STRAIN","CONDITION","ASSAY","PLATE","CTRL.GENO","CONTAM.CHECK","SORT","SCORE","MEAN.YFP.MEDIAN.ABS","MEAN.YFP.SD.ABS","MEAN.YFP.MEDIAN.WT","MEAN.YFP.MEDIAN.TDH3",
#                        "MEAN.FSC.CORRECT","MEAN.YFP.SD.WT.REL","MEAN.YFP.SD.TDH3.REL", "MEAN.YFP.CV.WT","MEAN.YFP.CV.TDH3","MEAN.YFP.CV.WT.REL","MEAN.YFP.CV.TDH3.REL",  
#                        "MEAN.YFP.SD.WT.REL","MEAN.YFP.SD.TDH3.REL","SD.YFP.MEDIAN.ABS",   
#                        "SD.YFP.SD.ABS","SD.YFP.MEDIAN.WT","SD.YFP.MEDIAN.TDH3","SD.FSC.CORRECT", 
#                        "SD.YFP.CV.WT","SD.YFP.CV.TDH3","SD.YFP.CV.WT.REL","SD.YFP.CV.TDH3.REL","SD.YFP.SD.WT.REL","SD.YFP.SD.TDH3.REL", "N","P.VAL.MEDIAN")
# 
# 
# #subsampling TDH3.SORT1 sham to match depth of sampling for other promoters
# TDH3.SORT1.PRUNED<-subset(TDH3.SORT1,TDH3.SORT1$CONDITION!="SHAM")
# ALLSHAM<-droplevels(subset(TDH3.SORT1,TDH3.SORT1$CONDITION=="SHAM"))
# CHOOSE.SHAM<-levels(ALLSHAM$STRAIN)[seq(1,260,4)]
# CHOOSE.TDH3.SHAM<-levels(ALLSHAM$STRAIN)[seq(20,277,4)]
# SHAM_subset<-ALLSHAM[ALLSHAM$STRAIN %in% CHOOSE.SHAM,]
# TDH3.SHAM_subset<-ALLSHAM[ALLSHAM$STRAIN %in% CHOOSE.TDH3.SHAM,]
# TDH3.SHAM_subset$CONDITION<-"TDH3.SHAM"
# TDH3.SORT1.PRUNED<-droplevels(rbind(TDH3.SORT1.PRUNED,SHAM_subset,TDH3.SHAM_subset))
# 
# ALL.PROMOTERS.WTDH3<-rbind(ALL.PROMOTERS[,ALLP.WTDH3.COLNAMES],TDH3.SORT1.PRUNED[,ALLP.WTDH3.COLNAMES])
# ALL.PROMOTERS.WTDH3$ASSAY<-factor(ALL.PROMOTERS.WTDH3$ASSAY)
# ALL.PROMOTERS.WTDH3$EXP<-as.factor(paste(ALL.PROMOTERS.WTDH3$ASSAY,ALL.PROMOTERS.WTDH3$SORT,ALL.PROMOTERS.WTDH3$SCORE,sep="."))
# 
# write.table(ALL.PROMOTERS.WTDH3,file="PROCESSED.DATA.txt",quote=FALSE,row.names=FALSE,col.names=TRUE,sep="\t")
# write.table(TDH3.SORT1,file="TDH3.EXP1.DATA.txt",row.names=FALSE,col.names=TRUE,sep="\t",quote=FALSE)
# write.table(TDH3.SORT2,file="TDH3.EXP2.DATA.txt",row.names=FALSE,col.names=TRUE,sep="\t",quote=FALSE)

###################################################################
#Load data files provided with manuscript

rm(list=ls())
options(warn=-1)

col_list<-c("red3", "darkorange1", "gold", "#63BF4D", "springgreen4", "#4DBFBF", "#4D7ABF", "royalblue4", "#A84DBF", "maroon")

ALL.PROMOTERS.WTDH3<-read.table("PROCESSED.DATA.txt", header = TRUE,sep="\t")
TDH3.SORT1<-read.table("TDH3.EXP1.DATA.txt", header = TRUE,sep="\t")
TDH3.SORT2<-read.table("TDH3.EXP2.DATA.txt", header = TRUE,sep="\t")


###################################################################
#Figure 1
#To assess the repeatabiliy of independent mutagenesis experiments:

#QQplot of 2 TDH3 samples
windows()
par(mfrow=c(1,2))
qqplot(TDH3.SORT1[TDH3.SORT1$CONDITION=="EMS","MEAN.YFP.MEDIAN.WT"],TDH3.SORT2[TDH3.SORT2$CONDITION=="EMS","MEAN.YFP.MEDIAN.WT"],
       xlim=c(0.1,1.9),ylim=c(0.1,1.9),bg="red",xlab="Experiment 1 (n=254)",ylab="Experiment 2 (n=1213)",pch=20,col="red",bty="L")
abline(a=0,b=1,lty=2,lwd=2)
#add gray shading at 25-75% density
XQUART<-quantile(TDH3.SORT1[TDH3.SORT1$CONDITION=="EMS","MEAN.YFP.MEDIAN.WT"], c(0.25,0.75))
YQUART<-quantile(TDH3.SORT2[TDH3.SORT2$CONDITION=="EMS","MEAN.YFP.MEDIAN.WT"], c(0.25,0.75))
polygon(c(XQUART, rev(XQUART)), c(c(1.91,1.91),c(0,0)), col = alpha("gray",0.3),border="NA")
polygon( c(c(0,0),c(1.91,1.91)),c(YQUART, rev(YQUART)), col = alpha("gray",0.3),border="NA")
#add box around 95% density
XQUANT<-quantile(TDH3.SORT1[TDH3.SORT1$CONDITION=="EMS","MEAN.YFP.MEDIAN.WT"], c(0.025,0.975))
YQUANT<-quantile(TDH3.SORT2[TDH3.SORT2$CONDITION=="EMS","MEAN.YFP.MEDIAN.WT"], c(0.025,0.975))
polygon(c(XQUANT, rev(XQUANT)), c(c(YQUANT[2],YQUANT[2]),c(YQUANT[1],YQUANT[1])), col = "NA",border="black",lty=1,lwd=1)
#plot 95% inset panel
#windows()
QUANT<-c(min(XQUANT[1],YQUANT[1]),max(XQUANT[2],YQUANT[2]))
qqplot(TDH3.SORT1[TDH3.SORT1$CONDITION=="EMS","MEAN.YFP.MEDIAN.WT"],TDH3.SORT2[TDH3.SORT2$CONDITION=="EMS","MEAN.YFP.MEDIAN.WT"],xlim=c(XQUANT[1],XQUANT[2]),ylim=c(YQUANT[1],YQUANT[2]),col="red",bg="red",xlab="Experiment 1 (n=254)",ylab="Experiment 2 (n=1213)",pch=21)
abline(a=0,b=1,lty=2,lwd=2)
#add dotted lines at quartiles
XQUART<-quantile(TDH3.SORT1[TDH3.SORT1$CONDITION=="EMS","MEAN.YFP.MEDIAN.WT"], c(0.25,0.75))
YQUART<-quantile(TDH3.SORT2[TDH3.SORT2$CONDITION=="EMS","MEAN.YFP.MEDIAN.WT"], c(0.25,0.75))
abline(v=XQUART,lty=3,col="gray")
abline(h=YQUART,lty=3,col="gray")
#add solid lines at 10 & 90% density
XQUANT<-quantile(TDH3.SORT1[TDH3.SORT1$CONDITION=="EMS","MEAN.YFP.MEDIAN.WT"], c(0.1,0.9))
YQUANT<-quantile(TDH3.SORT2[TDH3.SORT2$CONDITION=="EMS","MEAN.YFP.MEDIAN.WT"], c(0.1,0.9))
abline(v=XQUANT,lty=1,col="gray")
abline(h=YQUANT,lty=1,col="gray")

#test for sig diff between 2 independnent TDH3 experiments
wilcox.test(TDH3.SORT1[TDH3.SORT1$CONDITION=="EMS","MEAN.YFP.MEDIAN.WT"],TDH3.SORT2[TDH3.SORT2$CONDITION=="EMS","MEAN.YFP.MEDIAN.WT"],conf.int=TRUE,correct=TRUE,conf.level=0.95)
ks.test(TDH3.SORT1[TDH3.SORT1$CONDITION=="EMS","MEAN.YFP.MEDIAN.WT"],TDH3.SORT2[TDH3.SORT2$CONDITION=="EMS","MEAN.YFP.MEDIAN.WT"])
ad.test(TDH3.SORT1[TDH3.SORT1$CONDITION=="EMS","MEAN.YFP.MEDIAN.WT"],TDH3.SORT2[TDH3.SORT2$CONDITION=="EMS","MEAN.YFP.MEDIAN.WT"])

#######################################################################################

###########################
#                         #
#   Comparing Symmetry    #
#                         #
###########################

#Figure 2A-J:
#Generate density distributions of median expression values by promoter and condition

require(gridExtra)
GPD1<-ggplot(ALL.PROMOTERS.WTDH3[ALL.PROMOTERS.WTDH3$ASSAY=="GPD1"& (ALL.PROMOTERS.WTDH3$CONDITION=="EMS"|ALL.PROMOTERS.WTDH3$CONDITION=="SHAM"),],aes(x=MEAN.YFP.MEDIAN.WT,group=CONDITION,fill=CONDITION)) + scale_fill_manual(values=c(col_list[1],alpha("gray",0.5)))+
  geom_density(adjust=3,show.legend=FALSE)+ geom_vline(xintercept=1,color="black",lty=3,lwd=0.6) +annotate("text", label = "GPD1", x=-Inf,y=Inf,hjust=-0.2,vjust=1.2, size = 5, colour = "black")+xlim(0.4,1.6) +
  xlab("") + ylab("Density")+
  theme_bw()+theme(panel.grid.major = element_blank(), panel.grid.minor = element_blank())

OST1<-ggplot(ALL.PROMOTERS.WTDH3[ALL.PROMOTERS.WTDH3$ASSAY=="OST1"& (ALL.PROMOTERS.WTDH3$CONDITION=="EMS"|ALL.PROMOTERS.WTDH3$CONDITION=="SHAM"),],aes(x=MEAN.YFP.MEDIAN.WT,group=CONDITION,fill=CONDITION)) + scale_fill_manual(values=c(col_list[2],alpha("gray",0.5)))+
  geom_density(adjust=3,show.legend=FALSE) +  geom_vline(xintercept=1,color="black",lty=3,lwd=0.6) +annotate("text", label = " OST1", x=-Inf,y=Inf,hjust=-0.2,vjust=1.2, size = 5, colour = "black")+ xlim(0.4,1.6) + 
  xlab("") + ylab("")+
  theme_bw()+theme(panel.grid.major = element_blank(), panel.grid.minor = element_blank())

PFY1<-ggplot(ALL.PROMOTERS.WTDH3[ALL.PROMOTERS.WTDH3$ASSAY=="PFY1"& (ALL.PROMOTERS.WTDH3$CONDITION=="EMS"|ALL.PROMOTERS.WTDH3$CONDITION=="SHAM"),],aes(x=MEAN.YFP.MEDIAN.WT,group=CONDITION,fill=CONDITION)) + scale_fill_manual(values=c(col_list[3],alpha("gray",0.5)))+
  geom_density(adjust=3,show.legend=FALSE) +  geom_vline(xintercept=1,color="black",lty=3,lwd=0.6) +annotate("text", label = "PFY1", x=-Inf,y=Inf,hjust=-0.2,vjust=1.2, size = 5, colour = "black")+xlim(0.4,1.6) + 
  xlab("") + ylab("")+
  theme_bw()+theme(panel.grid.major = element_blank(), panel.grid.minor = element_blank())

RNR1<-ggplot(ALL.PROMOTERS.WTDH3[ALL.PROMOTERS.WTDH3$ASSAY=="RNR1"& (ALL.PROMOTERS.WTDH3$CONDITION=="EMS"|ALL.PROMOTERS.WTDH3$CONDITION=="SHAM"),],aes(x=MEAN.YFP.MEDIAN.WT,group=CONDITION,fill=CONDITION)) + scale_fill_manual(values=c(col_list[4],alpha("gray",0.5)))+
  geom_density(adjust=3,show.legend=FALSE) +  geom_vline(xintercept=1,color="black",lty=3,lwd=0.6) +annotate("text", label = "RNR1", x=-Inf,y=Inf,hjust=-0.2,vjust=1.2, size = 5, colour = "black")+ xlim(0.4,1.6) + 
  xlab("") + ylab("")+
  theme_bw()+theme(panel.grid.major = element_blank(), panel.grid.minor = element_blank())

RNR2<-ggplot(ALL.PROMOTERS.WTDH3[ALL.PROMOTERS.WTDH3$ASSAY=="RNR2"& (ALL.PROMOTERS.WTDH3$CONDITION=="EMS"|ALL.PROMOTERS.WTDH3$CONDITION=="SHAM"),],aes(x=MEAN.YFP.MEDIAN.WT,group=CONDITION,fill=CONDITION)) + scale_fill_manual(values=c(col_list[5],alpha("gray",0.5)))+
  geom_density(adjust=3,show.legend=FALSE) +  geom_vline(xintercept=1,color="black",lty=3,lwd=0.6) +annotate("text", label = "RNR2", x=-Inf,y=Inf,hjust=-0.2,vjust=1.2, size = 5, colour = "black")+ xlim(0.4,1.6) + 
  xlab("") + ylab("")+
  theme_bw()+theme(panel.grid.major = element_blank(), panel.grid.minor = element_blank())

STM1<-ggplot(ALL.PROMOTERS.WTDH3[ALL.PROMOTERS.WTDH3$ASSAY=="STM1"& (ALL.PROMOTERS.WTDH3$CONDITION=="EMS"|ALL.PROMOTERS.WTDH3$CONDITION=="SHAM"),],aes(x=MEAN.YFP.MEDIAN.WT,group=CONDITION,fill=CONDITION)) + scale_fill_manual(values=c(col_list[6],alpha("gray",0.5)))+
  geom_density(adjust=3,show.legend=FALSE) +  geom_vline(xintercept=1,color="black",lty=3,lwd=0.6) +annotate("text", label = "STM1", x=-Inf,y=Inf,hjust=-0.2,vjust=1.2, size = 5, colour = "black")+ xlim(0.4,1.6) +  
  xlab("Median Expression") + ylab("Density")+theme_bw()+theme(panel.grid.major = element_blank(), panel.grid.minor = element_blank())

TDH1<-ggplot(ALL.PROMOTERS.WTDH3[ALL.PROMOTERS.WTDH3$ASSAY=="TDH1"& (ALL.PROMOTERS.WTDH3$CONDITION=="EMS"|ALL.PROMOTERS.WTDH3$CONDITION=="SHAM"),],aes(x=MEAN.YFP.MEDIAN.WT,group=CONDITION,fill=CONDITION)) + scale_fill_manual(values=c(col_list[7],alpha("gray",0.5)))+
  geom_density(adjust=3,show.legend=FALSE) +  geom_vline(xintercept=1,color="black",lty=3,lwd=0.6)+annotate("text", label = "TDH1", x=-Inf,y=Inf,hjust=-0.2,vjust=1.2, size = 5, colour = "black") + xlim(0.4,1.6) + 
  xlab("Median Expression") + ylab("")+theme_bw()+theme(panel.grid.major = element_blank(), panel.grid.minor = element_blank())

TDH2<-ggplot(ALL.PROMOTERS.WTDH3[ALL.PROMOTERS.WTDH3$ASSAY=="TDH2"& (ALL.PROMOTERS.WTDH3$CONDITION=="EMS"|ALL.PROMOTERS.WTDH3$CONDITION=="SHAM"),],aes(x=MEAN.YFP.MEDIAN.WT,group=CONDITION,fill=CONDITION)) + scale_fill_manual(values=c(col_list[8],alpha("gray",0.5)))+
  geom_density(adjust=3,show.legend=FALSE) +  geom_vline(xintercept=1,color="black",lty=3,lwd=0.6) +annotate("text", label = "TDH2", x=-Inf,y=Inf,hjust=-0.2,vjust=1.2, size = 5, colour = "black")+ xlim(0.4,1.6) + 
  xlab("Median Expression") + ylab("")+theme_bw()+theme(panel.grid.major = element_blank(), panel.grid.minor = element_blank())

TDH3<-ggplot(ALL.PROMOTERS.WTDH3[ALL.PROMOTERS.WTDH3$ASSAY=="TDH3"& (ALL.PROMOTERS.WTDH3$CONDITION=="EMS"|ALL.PROMOTERS.WTDH3$CONDITION=="SHAM"),],aes(x=MEAN.YFP.MEDIAN.WT,group=CONDITION,fill=CONDITION)) + scale_fill_manual(values=c(col_list[9],alpha("gray",0.5)))+
  geom_density(adjust=3,show.legend=FALSE) +  geom_vline(xintercept=1,color="black",lty=3,lwd=0.6) +annotate("text", label = "TDH3", x=-Inf,y=Inf,hjust=-0.2,vjust=1.2, size = 5, colour = "black")+ xlim(0.4,1.6) + 
  xlab("Median Expression") + ylab("")+theme_bw()+theme(panel.grid.major = element_blank(), panel.grid.minor = element_blank())

VMA7<-ggplot(ALL.PROMOTERS.WTDH3[ALL.PROMOTERS.WTDH3$ASSAY=="VMA7"& (ALL.PROMOTERS.WTDH3$CONDITION=="EMS"|ALL.PROMOTERS.WTDH3$CONDITION=="SHAM"),],aes(x=MEAN.YFP.MEDIAN.WT,group=CONDITION,fill=CONDITION)) + scale_fill_manual(values=c(col_list[10],alpha("gray",0.5)))+
  geom_density(adjust=3,show.legend=FALSE) +  geom_vline(xintercept=1,color="black",lty=3,lwd=0.6) +annotate("text", label = "VMA7", x=-Inf,y=Inf,hjust=-0.2,vjust=1.2, size = 5, colour = "black")+
  xlim(0.4,1.6) + xlab("Median Expression") + ylab("")+theme_bw()+theme(panel.grid.major = element_blank(), panel.grid.minor = element_blank())

windows()
grid.arrange(GPD1,OST1,PFY1,RNR1,RNR2,STM1,TDH1,TDH2,TDH3,VMA7,ncol=5)


#Binomial test
BINOM.RESULTS<-data.frame(matrix(ncol = 7, nrow = 10))
names(BINOM.RESULTS)<-c("ASSAY","SHAM.PROB.DECREASE","SHAM.PVAL","SHAM.FDR.PVAL","EMS.PROB.DECREASE","EMS.PVAL","EMS.FDR.PVAL")
for (j in 1:length(levels(ALL.PROMOTERS.WTDH3$ASSAY))){
  EMS<-ALL.PROMOTERS.WTDH3[(ALL.PROMOTERS.WTDH3$CONDITION=="EMS") & (ALL.PROMOTERS.WTDH3$ASSAY==levels(ALL.PROMOTERS.WTDH3$ASSAY)[j]),"MEAN.YFP.MEDIAN.WT"]  
  SHAM<-ALL.PROMOTERS.WTDH3[(ALL.PROMOTERS.WTDH3$CONDITION=="SHAM") & (ALL.PROMOTERS.WTDH3$ASSAY==levels(ALL.PROMOTERS.WTDH3$ASSAY)[j]),"MEAN.YFP.MEDIAN.WT"]  
  SHAM.TEST<-binom.test(count(SHAM>1)[,2],0.5,alternative="two.sided",conf.level=0.95)
  BINOM.RESULTS[j,1]<-levels(ALL.PROMOTERS.WTDH3$ASSAY)[j]
  BINOM.RESULTS[j,2]<-SHAM.TEST$estimate
  BINOM.RESULTS[j,3]<-SHAM.TEST$p.value
  EMS.TEST<-binom.test(count(EMS>1)[,2],0.5,alternative="two.sided",conf.level=0.95)
  BINOM.RESULTS[j,5]<-EMS.TEST$estimate
  BINOM.RESULTS[j,6]<-EMS.TEST$p.value
  
}
BINOM.RESULTS[,4]<-p.adjust(BINOM.RESULTS[,3],method="BH")
BINOM.RESULTS[,7]<-p.adjust(BINOM.RESULTS[,6],method="BH")


#Permutation test - increases vs decreases

#custom function pasted at the end of this file

#permutes mean differences between groups and also downsamples to match group sizes
DOWNSAMPLE.DIST.PERMUTE.MEAN <- function(GROUP.1,GROUP.2) {
  N.DOWNSAMPLE <-min(length(GROUP.1),length(GROUP.2))
  N.PERM <- 10000
  OBS.DIFF <- rep(NA,N.PERM)
  PERM.DIFF <- rep(NA,N.PERM)
  for (i in 1:N.PERM)
  {
    SAMPLE.1 <-sample(GROUP.1,size=N.DOWNSAMPLE,replace=TRUE)
    SAMPLE.2 <-sample(GROUP.2,size=N.DOWNSAMPLE,replace=TRUE)
    OBS.DIFF[i] <- mean(SAMPLE.1)-mean(SAMPLE.2)
    POOL <- c(SAMPLE.1,SAMPLE.2)
    POS.1 <- sample(1:length(POOL),size=N.DOWNSAMPLE,replace=TRUE)
    PERM.1 <- POOL[POS.1]
    PERM.2 <- POOL[-POS.1]
    PERM.DIFF[i] <- mean(PERM.1)-mean(PERM.2)
  }
  EFFECT.SIZE <-mean(OBS.DIFF)
  P.VAL <- (length(which(abs(PERM.DIFF) > abs(OBS.DIFF))) + 1)/(N.PERM + 1)
  RESULTS <- list(EFFECT.SIZE,P.VAL)
  return(RESULTS)}

#Calculate permutation result on mean magnitude of increases vs decreases
PERMUT.MEANMAG.RESULTS<-data.frame(matrix(ncol = 7, nrow = 10))
names(PERMUT.MEANMAG.RESULTS)<-c("ASSAY","SHAM.STATISTIC","SHAM.PVAL","EMS.STATISTIC","EMS.PVAL")
for (j in 1:length(levels(ALL.PROMOTERS.WTDH3$ASSAY))){
  EMS<-ALL.PROMOTERS.WTDH3[(ALL.PROMOTERS.WTDH3$CONDITION=="EMS") & (ALL.PROMOTERS.WTDH3$ASSAY==levels(ALL.PROMOTERS.WTDH3$ASSAY)[j]),"MEAN.YFP.MEDIAN.WT"]  
  SHAM<-ALL.PROMOTERS.WTDH3[(ALL.PROMOTERS.WTDH3$CONDITION=="SHAM") & (ALL.PROMOTERS.WTDH3$ASSAY==levels(ALL.PROMOTERS.WTDH3$ASSAY)[j]),"MEAN.YFP.MEDIAN.WT"]  
  SHAM.TEST<-DOWNSAMPLE.DIST.PERMUTE.MEAN(SHAM[SHAM>1]-1,1-SHAM[SHAM<1])
  PERMUT.MEANMAG.RESULTS[j,1]<-levels(ALL.PROMOTERS.WTDH3$ASSAY)[j]
  PERMUT.MEANMAG.RESULTS[j,2]<-SHAM.TEST[1]
  PERMUT.MEANMAG.RESULTS[j,3]<-SHAM.TEST[2]
  EMS.TEST<-DOWNSAMPLE.DIST.PERMUTE.MEAN(EMS[EMS>1]-1,1-EMS[EMS<1])
  PERMUT.MEANMAG.RESULTS[j,4]<-EMS.TEST[1]
  PERMUT.MEANMAG.RESULTS[j,5]<-EMS.TEST[2]
  
}
PERMUT.MEANMAG.RESULTS[,3]<-p.adjust( PERMUT.MEANMAG.RESULTS[,3],method="BH")
PERMUT.MEANMAG.RESULTS[,6]<-p.adjust( PERMUT.MEANMAG.RESULTS[,6],method="BH")


###########################
#                         #
# Comparing Distributions #
#                         #
###########################


#Calculate Zscores for data 

for (i in 1:nrow(ALL.PROMOTERS.WTDH3))
{
  CUR <- subset(ALL.PROMOTERS.WTDH3, ALL.PROMOTERS.WTDH3$EXP == ALL.PROMOTERS.WTDH3[i,"EXP"])
  CUR.SHAM<-subset(CUR, CONDITION == "SHAM"|CONDITION == "SHAM.ALL")
  ALL.PROMOTERS.WTDH3[i,"ZSCORE.YFP.MEDIAN"] <- (ALL.PROMOTERS.WTDH3[i,"MEAN.YFP.MEDIAN.WT"] - median(CUR.SHAM$MEAN.YFP.MEDIAN.WT))/mad(CUR.SHAM$MEAN.YFP.MEDIAN.WT)
  ALL.PROMOTERS.WTDH3[i,"ZSCORE.YFP.CV"] <- (ALL.PROMOTERS.WTDH3[i,"MEAN.YFP.CV.WT"] - median(CUR.SHAM$MEAN.YFP.CV.WT))/mad(CUR.SHAM$MEAN.YFP.CV.WT)
}

##################################################################
#Figure S1: Violin plots of sham - supplemental figure

windows()
SHAM<-droplevels(subset(ALL.PROMOTERS.WTDH3, ALL.PROMOTERS.WTDH3$CONDITION=="SHAM"))
SHAM$ASSAY = factor(SHAM$ASSAY,levels(SHAM$ASSAY)[c(1:8,10,9)])
col_list<-col_list[c(1:8,10,9)]
BEFORE<-ggplot(SHAM,aes(x=ASSAY, y=MEAN.YFP.MEDIAN.WT,group=ASSAY,fill=ASSAY)) + scale_fill_manual(values=col_list)+
  geom_violin(scale="area",trim="FALSE",show.legend="FALSE",width=1)+ ylab("Median Expression")+xlab("") + 
  theme_bw()+theme(text=element_text(size=20),axis.text.x = element_text(angle=90,size=18, hjust=1),panel.grid.major = element_blank(), panel.grid.minor = element_blank())
AFTER<-ggplot(SHAM,aes(x=ASSAY, y=ZSCORE.YFP.MEDIAN,group=ASSAY,fill=ASSAY)) + scale_fill_manual(values=col_list)+
  geom_violin(scale="area",trim="FALSE",show.legend="FALSE",width=0.8)+ ylab("Zscore")+xlab("")+ylim(-12,12)+
  theme_bw()+theme(text=element_text(size=20),axis.text.x = element_text(angle=90,size=18, hjust=1),panel.grid.major = element_blank(), panel.grid.minor = element_blank())
grid.arrange(BEFORE,AFTER,ncol=2)

NORMAL.Z<-data.frame(matrix(ncol = 5, nrow = 10))
names(NORMAL.Z)<-c("ASSAY","SHAM.SHAPIRO.W","SHAM.SHAPIRO.PVAL","EMS.SHAPIRO.W","EMS.SHAPIRO.PVAL")

#Testing for normality
for (j in 1:length(levels(ALL.PROMOTERS.WTDH3$ASSAY))){
  CUR<-ALL.PROMOTERS.WTDH3[(ALL.PROMOTERS.WTDH3$CONDITION=="EMS"|ALL.PROMOTERS.WTDH3$CONDITION=="SHAM") & (ALL.PROMOTERS.WTDH3$ASSAY==levels(ALL.PROMOTERS.WTDH3$ASSAY)[j]),]  
  
  TEST<-shapiro.test(CUR[CUR$CONDITION=="SHAM","ZSCORE.YFP.MEDIAN"])
  NORMAL.Z[j,1]<-levels(ALL.PROMOTERS.WTDH3$ASSAY)[j]
  NORMAL.Z[j,2]<-TEST$statistic
  NORMAL.Z[j,3]<-TEST$p.value
  
  TEST<-shapiro.test(CUR[CUR$CONDITION=="EMS","ZSCORE.YFP.MEDIAN"])
  NORMAL.Z[j,4]<-TEST$statistic
  NORMAL.Z[j,5]<-TEST$p.value
}
#Applying multiple test corrections
NORMAL.Z$SHAM.PADJ<-p.adjust(NORMAL.Z$SHAM.SHAPIRO.PVAL,method="BH",n=length(levels(ALL.PROMOTERS.WTDH3$ASSAY)))
NORMAL.Z$EMS.PADJ<-p.adjust(NORMAL.Z$EMS.SHAPIRO.PVAL,method="BH",n=length(levels(ALL.PROMOTERS.WTDH3$ASSAY)))
  

#Identify the number of EMS-treated genotypes with mean medians within 2 MAD of the distribution median
SD.Z<-data.frame(matrix(ncol = 2, nrow = 10))
names(NORMAL.Z)<-c("ASSAY","PCT.2MAD")

for (j in 1:length(levels(ALL.PROMOTERS.WTDH3$ASSAY))){
  CUR<-ALL.PROMOTERS.WTDH3[(ALL.PROMOTERS.WTDH3$CONDITION=="EMS") & (ALL.PROMOTERS.WTDH3$ASSAY==levels(ALL.PROMOTERS.WTDH3$ASSAY)[j]),]  
  SD.Z[j,1]<-levels(ALL.PROMOTERS.WTDH3$ASSAY)[j]
  IN<-CUR[CUR$ZSCORE.YFP.MEDIAN<2&CUR$ZSCORE.YFP.MEDIAN>-2,"STRAIN"]
  SD.Z[j,2]<-length(IN)/nrow(CUR)
}


#What does the distribution of mutational effects look like on a scale
#that takes into account differences in non-genetic variability?

#Figure 3: Zscore ggridges plot with Q-Q plots to right

pdf("Figure 3 draft median zscore densities qqplot insets fixed.pdf")
#jpeg(filename="Figure 5 draft median zscore densities qqplot insets.jpg")
#windows()
EMS<-droplevels(subset(ALL.PROMOTERS.WTDH3, ALL.PROMOTERS.WTDH3$CONDITION=="EMS"|ALL.PROMOTERS.WTDH3$CONDITION=="EMS.ALL"))

EMS$ASSAY = factor(EMS$ASSAY,levels(EMS$ASSAY)[c(1:8,10,9)])
col_list<-col_list[c(1:8,10,9)]

LEFTPANEL<-ggplot(EMS, aes(x = ZSCORE.YFP.MEDIAN, y = fct_rev(ASSAY), fill = ASSAY)) + 
  geom_density_ridges(scale = 1.5) + geom_density_ridges(rel_min_height = 0.01)  +
  scale_fill_cyclical(values = col_list)  + 
  geom_vline(xintercept = 0, color = "white",linetype = "dashed") +
  geom_vline(xintercept = -2, color = "gray",linetype = "dashed") +
  geom_vline(xintercept = 2, color = "gray",linetype = "dashed") +
  theme_ridges(center_axis_labels = TRUE)+
  scale_x_continuous(limits = c(-10,10),expand = c(0.01, 0)) +
  scale_y_discrete(expand = c(0, 0)) +
  xlab("Expression Zscore") + ylab("") 

INCREASES<-ALL.PROMOTERS.WTDH3[(ALL.PROMOTERS.WTDH3$CONDITION=="EMS")&(ALL.PROMOTERS.WTDH3$ZSCORE.YFP.MEDIAN>0),c("ASSAY","ZSCORE.YFP.MEDIAN")]
INCREASES$DIR<-"INC"
DECREASES<-ALL.PROMOTERS.WTDH3[(ALL.PROMOTERS.WTDH3$CONDITION=="EMS")&(ALL.PROMOTERS.WTDH3$ZSCORE.YFP.MEDIAN<0),c("ASSAY","ZSCORE.YFP.MEDIAN")]
DECREASES$ZSCORE.YFP.MEDIAN<-(-DECREASES$ZSCORE.YFP.MEDIAN)
DECREASES$DIR<-"DEC"  
QQDATA<-rbind(INCREASES,DECREASES)

#How do promoters differ in the frequency and magnitude of continuous 
#increases vs decreases?

vp.Right <- viewport(height=unit(1, "npc"), width=unit(0.8, "npc"), 
                     just=c("left","top"), 
                     y=1, x=0)
par(mfcol=c(11,5))
par(mar=c(1,1,1,1))
par(mgp=c(0.5,.1,0))
plot.new();plot.new();plot.new();plot.new();plot.new();plot.new();plot.new();plot.new();plot.new();plot.new();plot.new();
plot.new();plot.new();plot.new();plot.new();plot.new();plot.new();plot.new();plot.new();plot.new();plot.new();plot.new();
plot.new();plot.new();plot.new();plot.new();plot.new();plot.new();plot.new();plot.new();plot.new();plot.new();plot.new();
plot.new();plot.new();plot.new();plot.new();plot.new();plot.new();plot.new();plot.new();plot.new();plot.new();plot.new();


#maxlim<-ceiling(max(c(INCREASES[INCREASES$ASSAY=="GPD1","ZSCORE.YFP.MEDIAN"],DECREASES[DECREASES$ASSAY=="GPD1","ZSCORE.YFP.MEDIAN"])))
Q95_U<-max(c(quantile(INCREASES[INCREASES$ASSAY=="GPD1","ZSCORE.YFP.MEDIAN"],0.95),quantile(DECREASES[DECREASES$ASSAY=="GPD1","ZSCORE.YFP.MEDIAN"],0.95)))
GPD1<-qqplot(INCREASES[INCREASES$ASSAY=="GPD1","ZSCORE.YFP.MEDIAN"],DECREASES[DECREASES$ASSAY=="GPD1","ZSCORE.YFP.MEDIAN"],pch=16, col=col_list[1],xlab="",ylab="",xlim=c(0,Q95_U),ylim=c(0,Q95_U), cex=1,tck=0)
abline(a=0,b=1,lty=3)
#maxlim<-ceiling(max(c(INCREASES[INCREASES$ASSAY=="OST1","ZSCORE.YFP.MEDIAN"],DECREASES[DECREASES$ASSAY=="OST1","ZSCORE.YFP.MEDIAN"])))
Q95_U<-max(c(quantile(INCREASES[INCREASES$ASSAY=="OST1","ZSCORE.YFP.MEDIAN"],0.95),quantile(DECREASES[DECREASES$ASSAY=="OST1","ZSCORE.YFP.MEDIAN"],0.95)))
OST1<-qqplot(INCREASES[INCREASES$ASSAY=="OST1","ZSCORE.YFP.MEDIAN"],DECREASES[DECREASES$ASSAY=="OST1","ZSCORE.YFP.MEDIAN"],pch=16, col=col_list[2],xlab="",ylab="",xlim=c(0,Q95_U),ylim=c(0,Q95_U),cex.lab=1.2, cex=1,tck=0)
abline(a=0,b=1,lty=3)
maxlim<-ceiling(max(c(INCREASES[INCREASES$ASSAY=="PFY1","ZSCORE.YFP.MEDIAN"],DECREASES[DECREASES$ASSAY=="PFY1","ZSCORE.YFP.MEDIAN"])))
Q95_U<-max(c(quantile(INCREASES[INCREASES$ASSAY=="PFY1","ZSCORE.YFP.MEDIAN"],0.95),quantile(DECREASES[DECREASES$ASSAY=="PFY1","ZSCORE.YFP.MEDIAN"],0.95)))
PFY1<-qqplot(INCREASES[INCREASES$ASSAY=="PFY1","ZSCORE.YFP.MEDIAN"],DECREASES[DECREASES$ASSAY=="PFY1","ZSCORE.YFP.MEDIAN"],pch=16, col=col_list[3],xlab="",ylab="",xlim=c(0,Q95_U),ylim=c(0,Q95_U),cex.lab=1.2, cex=1,tck=0)
abline(a=0,b=1,lty=3)
maxlim<-ceiling(max(c(INCREASES[INCREASES$ASSAY=="RNR1","ZSCORE.YFP.MEDIAN"],DECREASES[DECREASES$ASSAY=="RNR1","ZSCORE.YFP.MEDIAN"])))
Q95_U<-max(c(quantile(INCREASES[INCREASES$ASSAY=="RNR1","ZSCORE.YFP.MEDIAN"],0.95),quantile(DECREASES[DECREASES$ASSAY=="RNR1","ZSCORE.YFP.MEDIAN"],0.95)))
RNR1<-qqplot(INCREASES[INCREASES$ASSAY=="RNR1","ZSCORE.YFP.MEDIAN"],DECREASES[DECREASES$ASSAY=="RNR1","ZSCORE.YFP.MEDIAN"],pch=16, col=col_list[4],xlab="",ylab="",xlim=c(0,Q95_U),ylim=c(0,Q95_U),cex.lab=1.2, cex=1,tck=0)
abline(a=0,b=1,lty=3)
maxlim<-ceiling(max(c(INCREASES[INCREASES$ASSAY=="RNR2","ZSCORE.YFP.MEDIAN"],DECREASES[DECREASES$ASSAY=="RNR2","ZSCORE.YFP.MEDIAN"])))
Q95_U<-max(c(quantile(INCREASES[INCREASES$ASSAY=="RNR2","ZSCORE.YFP.MEDIAN"],0.95),quantile(DECREASES[DECREASES$ASSAY=="RNR2","ZSCORE.YFP.MEDIAN"],0.95)))
RNR2<-qqplot(INCREASES[INCREASES$ASSAY=="RNR2","ZSCORE.YFP.MEDIAN"],DECREASES[DECREASES$ASSAY=="RNR2","ZSCORE.YFP.MEDIAN"],pch=16, col=col_list[5],xlab="",ylab="",xlim=c(0,Q95_U),ylim=c(0,Q95_U),cex.lab=1.2, cex=1,tck=0)
abline(a=0,b=1,lty=3)
maxlim<-ceiling(max(c(INCREASES[INCREASES$ASSAY=="STM1","ZSCORE.YFP.MEDIAN"],DECREASES[DECREASES$ASSAY=="STM1","ZSCORE.YFP.MEDIAN"])))
Q95_U<-max(c(quantile(INCREASES[INCREASES$ASSAY=="STM1","ZSCORE.YFP.MEDIAN"],0.95),quantile(DECREASES[DECREASES$ASSAY=="STM1","ZSCORE.YFP.MEDIAN"],0.95)))
STM1<-qqplot(INCREASES[INCREASES$ASSAY=="STM1","ZSCORE.YFP.MEDIAN"],DECREASES[DECREASES$ASSAY=="STM1","ZSCORE.YFP.MEDIAN"],pch=16, col=col_list[6],xlab="",ylab="",xlim=c(0,Q95_U),ylim=c(0,Q95_U),cex.lab=1.2, cex=1,tck=0)
abline(a=0,b=1,lty=3)
maxlim<-ceiling(max(c(INCREASES[INCREASES$ASSAY=="TDH1","ZSCORE.YFP.MEDIAN"],DECREASES[DECREASES$ASSAY=="TDH1","ZSCORE.YFP.MEDIAN"])))
Q95_U<-max(c(quantile(INCREASES[INCREASES$ASSAY=="TDH1","ZSCORE.YFP.MEDIAN"],0.95),quantile(DECREASES[DECREASES$ASSAY=="TDH1","ZSCORE.YFP.MEDIAN"],0.95)))
TDH1<-qqplot(INCREASES[INCREASES$ASSAY=="TDH1","ZSCORE.YFP.MEDIAN"],DECREASES[DECREASES$ASSAY=="TDH1","ZSCORE.YFP.MEDIAN"],pch=16, col=col_list[7],xlab="",ylab="",xlim=c(0,Q95_U),ylim=c(0,Q95_U),cex.lab=1.2, cex=1,tck=0)
abline(a=0,b=1,lty=3)
maxlim<-ceiling(max(c(INCREASES[INCREASES$ASSAY=="TDH2","ZSCORE.YFP.MEDIAN"],DECREASES[DECREASES$ASSAY=="TDH2","ZSCORE.YFP.MEDIAN"])))
Q95_U<-max(c(quantile(INCREASES[INCREASES$ASSAY=="TDH2","ZSCORE.YFP.MEDIAN"],0.95),quantile(DECREASES[DECREASES$ASSAY=="TDH2","ZSCORE.YFP.MEDIAN"],0.95)))
TDH2<-qqplot(INCREASES[INCREASES$ASSAY=="TDH2","ZSCORE.YFP.MEDIAN"],DECREASES[DECREASES$ASSAY=="TDH2","ZSCORE.YFP.MEDIAN"],pch=16, col=col_list[8],xlab="",ylab="",xlim=c(0,Q95_U),ylim=c(0,Q95_U),cex.lab=1.2, cex=1,tck=0)
abline(a=0,b=1,lty=3)
maxlim<-ceiling(max(c(INCREASES[INCREASES$ASSAY=="TDH3","ZSCORE.YFP.MEDIAN"],DECREASES[DECREASES$ASSAY=="TDH3","ZSCORE.YFP.MEDIAN"])))
Q95_U<-max(c(quantile(INCREASES[INCREASES$ASSAY=="TDH3","ZSCORE.YFP.MEDIAN"],0.95),quantile(DECREASES[DECREASES$ASSAY=="TDH3","ZSCORE.YFP.MEDIAN"],0.95)))
TDH3<-qqplot(INCREASES[INCREASES$ASSAY=="TDH3","ZSCORE.YFP.MEDIAN"],DECREASES[DECREASES$ASSAY=="TDH3","ZSCORE.YFP.MEDIAN"],pch=16, col=col_list[9],xlab="",ylab="",xlim=c(0,Q95_U),ylim=c(0,Q95_U),cex.lab=1.2, cex=1,tck=0)
abline(a=0,b=1,lty=3)
maxlim<-ceiling(max(c(INCREASES[INCREASES$ASSAY=="VMA7","ZSCORE.YFP.MEDIAN"],DECREASES[DECREASES$ASSAY=="VMA7","ZSCORE.YFP.MEDIAN"])))
Q95_U<-max(c(quantile(INCREASES[INCREASES$ASSAY=="VMA7","ZSCORE.YFP.MEDIAN"],0.95),quantile(DECREASES[DECREASES$ASSAY=="VMA7","ZSCORE.YFP.MEDIAN"],0.95)))
VMA7<-qqplot(INCREASES[INCREASES$ASSAY=="VMA7","ZSCORE.YFP.MEDIAN"],DECREASES[DECREASES$ASSAY=="VMA7","ZSCORE.YFP.MEDIAN"],pch=16, col=col_list[10],xlab="",ylab="",xlim=c(0,Q95_U),ylim=c(0,Q95_U),cex.lab=1.2, cex=1,tck=0)
abline(a=0,b=1,lty=3)

print(LEFTPANEL, vp=vp.Right)
dev.off()


#Are mutational distributions statistically different from one another?

#Pairwise AD.TEST (contributes to coloring in Figure S6)
AD.RESULTS.EMS.SHAM.Z<-data.frame(matrix(ncol = 3, nrow = 10))
AD.RESULTS.EMS.EMS.Z<-data.frame(matrix(ncol = 4, nrow = 100))

for (j in 1:length(levels(ALL.PROMOTERS.WTDH3$ASSAY))){
  CUR<-ALL.PROMOTERS.WTDH3[(ALL.PROMOTERS.WTDH3$CONDITION=="EMS"|ALL.PROMOTERS.WTDH3$CONDITION=="SHAM") & (ALL.PROMOTERS.WTDH3$ASSAY==levels(ALL.PROMOTERS.WTDH3$ASSAY)[j]),]  
  
  TEST<-ad.test(CUR[CUR$CONDITION=="SHAM","ZSCORE.YFP.MEDIAN"],CUR[CUR$CONDITION=="EMS","ZSCORE.YFP.MEDIAN"])
  AD.RESULTS.EMS.SHAM.Z[j,1]<-levels(ALL.PROMOTERS.WTDH3$ASSAY)[j]
  AD.RESULTS.EMS.SHAM.Z[j,2]<-TEST$ad[2,1]
  AD.RESULTS.EMS.SHAM.Z[j,3]<-TEST$ad[2,3]
  
  for (i in 1:length(levels(ALL.PROMOTERS.WTDH3$ASSAY))){
    ALT<-ALL.PROMOTERS.WTDH3[(ALL.PROMOTERS.WTDH3$CONDITION=="EMS" & ALL.PROMOTERS.WTDH3$ASSAY==levels(ALL.PROMOTERS.WTDH3$ASSAY)[i]),]  
    
    TEST<-ad.test(CUR[CUR$CONDITION=="EMS","ZSCORE.YFP.MEDIAN"],ALT[,"ZSCORE.YFP.MEDIAN"])
    AD.RESULTS.EMS.EMS.Z[(j-1)*10+i,1]<-levels(ALL.PROMOTERS.WTDH3$ASSAY)[j]
    AD.RESULTS.EMS.EMS.Z[(j-1)*10+i,2]<-levels(ALL.PROMOTERS.WTDH3$ASSAY)[i]
    AD.RESULTS.EMS.EMS.Z[(j-1)*10+i,3]<-TEST$ad[2,1]
    AD.RESULTS.EMS.EMS.Z[(j-1)*10+i,4]<-TEST$ad[2,3]
  }
}
#Remove duplicate pairwise contrasts
keep<-c(seq(2,10,1),seq(13,20,1),seq(24,30,1),seq(35,40,1),seq(46,50,1),seq(57,60,1),seq(68,70,1),seq(79,80,1),90)
AD.RESULTS.EMS.EMS.Z<-AD.RESULTS.EMS.EMS.Z[keep,]

#Applying multiple test corrections
AD.RESULTS.EMS.SHAM.Z$PADJ<-p.adjust(AD.RESULTS.EMS.SHAM.Z$X3,method="BH",n=length(levels(ALL.PROMOTERS.WTDH3$ASSAY)))
AD.RESULTS.EMS.EMS.Z$PADJ<-p.adjust(AD.RESULTS.EMS.EMS.Z$X4,method="BH",n=nrow(AD.RESULTS.EMS.EMS.Z))
DIST.SIGDIFF<-AD.RESULTS.EMS.EMS.Z[AD.RESULTS.EMS.EMS.Z$PADJ<0.05,]


#Visualizing pairwise differences among distributions of mutational effects with QQ plots

# Figure S6:Generating matix of pairwise qqplots among promoter distributions 

ALL.PAIRWISE<-data.frame(P1=c(rep("GPD1",10),rep("OST1",10),rep("PFY1",10),rep("RNR1",10),rep("RNR2",10),rep("STM1",10),rep("TDH1",10),rep("TDH2",10),rep("VMA7",10),rep("TDH3",10)),P2=c(rep(levels(ALL.PROMOTERS.WTDH3$ASSAY),10)))
keep<-c(seq(2,10,1),seq(13,20,1),seq(24,30,1),seq(35,40,1),seq(46,50,1),seq(57,60,1),seq(68,70,1),seq(79,80,1),90)
counter<-1

EMS<-ALL.PROMOTERS.WTDH3[ALL.PROMOTERS.WTDH3$CONDITION=="EMS",]
windows()
par(mfcol=c(9,9)) #multiple panels down column
par(mar = c(0, 0, 0, 0), oma = c(4, 5, 4, 0.5)) #set margins
par(tcl = -0.25) #tick length
par(mgp = c(2, 0.6, 0)) # margin line
counter<-1


for (j in 1:9){
  PROM1<-levels(ALL.PROMOTERS.WTDH3$ASSAY)[j]
  for (k in 2:10){
    PROM2<-levels(ALL.PROMOTERS.WTDH3$ASSAY)[k]
    contrast<-which((ALL.PAIRWISE$P1==PROM1&ALL.PAIRWISE$P2==PROM2)==1)
    if(contrast%in%keep){
      if(any(DIST.SIGDIFF$X1==PROM1&DIST.SIGDIFF$X2==PROM2)){
        qqplot(EMS[EMS$ASSAY==PROM1,"ZSCORE.YFP.MEDIAN"],EMS[EMS$ASSAY==PROM2,"ZSCORE.YFP.MEDIAN"],xlim=c(-20,20),ylim=c(-20,20),main="",col="red",pch=18,axes=F)
        box(col = "black")
        
      }
      if(!any(DIST.SIGDIFF$X1==PROM1&DIST.SIGDIFF$X2==PROM2)){
        qqplot(EMS[EMS$ASSAY==PROM1,"ZSCORE.YFP.MEDIAN"],EMS[EMS$ASSAY==PROM2,"ZSCORE.YFP.MEDIAN"],xlim=c(-20,20),ylim=c(-20,20), main="",col="black",pch=20,axes=F)
        box(col = "black")
      }
      abline(0,1,lty=3)
    }
    if(!contrast%in%keep){
      plot.new()
    }
    if (counter %in% c(seq(9,81,9))){
      axis(1, col = "grey40", col.axis = "grey20", at = seq(-20,20,10))
    }
    if (counter %in% c(1,11,21,31,41,51,61,71,81)){
      mtext(PROM1, side = 3, line = 0.5, adj=1,cex = 0.8, col = "black")
    }
    
    if (counter %in% c(1:9)){
      axis(2, col = "grey40", col.axis = "grey20", at = seq(-20,20,10),las=2)
      mtext(PROM2, side = 2, line = 1.5, las=2,cex = 0.8,col = "black")
      
    }
    counter<-counter+1                                                           
  }
}


#What is the nature of the differences among distributions of mutational effects?


###
# PCA on robust summary stats
###

SUMMARY.STATS.ROBUST<-data.frame(matrix(ncol = 10, nrow = 10))
names(SUMMARY.STATS.ROBUST)<-c("ASSAY","MEDIAN","MAD","MC","LMC","RMC","N","MIN","MAX","VAR")

SHAM.SUMMARY.STATS.ROBUST<-data.frame(matrix(ncol = 10, nrow = 10))
names(SHAM.SUMMARY.STATS.ROBUST)<-c("ASSAY","MEDIAN","MAD","MC","LMC","RMC","N","MIN","MAX","VAR")

SUMMARY.STATS.ROBUST.Z<-data.frame(matrix(ncol = 10, nrow = 10))
names(SUMMARY.STATS.ROBUST.Z)<-c("ASSAY","MEDIAN","MAD","MC","LMC","RMC","N","MIN","MAX","VAR")

for (j in 1:length(levels(ALL.PROMOTERS.WTDH3$ASSAY))){
  CUR<-ALL.PROMOTERS.WTDH3[(ALL.PROMOTERS.WTDH3$CONDITION=="EMS") & (ALL.PROMOTERS.WTDH3$ASSAY==levels(ALL.PROMOTERS.WTDH3$ASSAY)[j]),]  
  SUMMARY.STATS.ROBUST[j,1]<-levels(ALL.PROMOTERS.WTDH3$ASSAY)[j]
  SUMMARY.STATS.ROBUST[j,2]<-median(CUR[,"MEAN.YFP.MEDIAN.WT"])
  SUMMARY.STATS.ROBUST[j,3]<-mad(CUR[,"MEAN.YFP.MEDIAN.WT"])
  SUMMARY.STATS.ROBUST[j,4]<-mc(CUR[,"MEAN.YFP.MEDIAN.WT"])
  SUMMARY.STATS.ROBUST[j,5]<-mc(CUR[CUR$MEAN.YFP.MEDIAN.WT<SUMMARY.STATS.ROBUST[j,2],"MEAN.YFP.MEDIAN.WT"])
  SUMMARY.STATS.ROBUST[j,6]<-mc(CUR[CUR$MEAN.YFP.MEDIAN.WT>SUMMARY.STATS.ROBUST[j,2],"MEAN.YFP.MEDIAN.WT"])
  SUMMARY.STATS.ROBUST[j,7]<-nrow(CUR)
  SUMMARY.STATS.ROBUST[j,8]<-min(CUR[,"MEAN.YFP.MEDIAN.WT"])
  SUMMARY.STATS.ROBUST[j,9]<-max(CUR[,"MEAN.YFP.MEDIAN.WT"])
  SUMMARY.STATS.ROBUST[j,10]<-var(CUR[,"MEAN.YFP.MEDIAN.WT"])
  
  SUMMARY.STATS.ROBUST.Z[j,1]<-levels(ALL.PROMOTERS.WTDH3$ASSAY)[j]
  SUMMARY.STATS.ROBUST.Z[j,2]<-median(CUR[,"ZSCORE.YFP.MEDIAN"])
  SUMMARY.STATS.ROBUST.Z[j,3]<-mad(CUR[,"ZSCORE.YFP.MEDIAN"])
  SUMMARY.STATS.ROBUST.Z[j,4]<-mc(CUR[,"ZSCORE.YFP.MEDIAN"])
  SUMMARY.STATS.ROBUST.Z[j,5]<-mc(CUR[CUR$ZSCORE.YFP.MEDIAN<SUMMARY.STATS.ROBUST.Z[j,2],"ZSCORE.YFP.MEDIAN"])
  SUMMARY.STATS.ROBUST.Z[j,6]<-mc(CUR[CUR$ZSCORE.YFP.MEDIAN>SUMMARY.STATS.ROBUST.Z[j,2],"ZSCORE.YFP.MEDIAN"])
  SUMMARY.STATS.ROBUST.Z[j,7]<-nrow(CUR)
  SUMMARY.STATS.ROBUST.Z[j,8]<-min(CUR[,"ZSCORE.YFP.MEDIAN"])
  SUMMARY.STATS.ROBUST.Z[j,9]<-max(CUR[,"ZSCORE.YFP.MEDIAN"])
  SUMMARY.STATS.ROBUST.Z[j,10]<-var(CUR[,"ZSCORE.YFP.MEDIAN"])
  
  CUR<-ALL.PROMOTERS.WTDH3[(ALL.PROMOTERS.WTDH3$CONDITION=="SHAM") & (ALL.PROMOTERS.WTDH3$ASSAY==levels(ALL.PROMOTERS.WTDH3$ASSAY)[j]),]  
  SHAM.SUMMARY.STATS.ROBUST[j,1]<-levels(ALL.PROMOTERS.WTDH3$ASSAY)[j]
  SHAM.SUMMARY.STATS.ROBUST[j,2]<-median(CUR[,"MEAN.YFP.MEDIAN.WT"])
  SHAM.SUMMARY.STATS.ROBUST[j,3]<-mad(CUR[,"MEAN.YFP.MEDIAN.WT"])
  SHAM.SUMMARY.STATS.ROBUST[j,4]<-mc(CUR[,"MEAN.YFP.MEDIAN.WT"])
  SHAM.SUMMARY.STATS.ROBUST[j,5]<-mc(CUR[CUR$MEAN.YFP.MEDIAN.WT<SUMMARY.STATS.ROBUST[j,2],"MEAN.YFP.MEDIAN.WT"])
  SHAM.SUMMARY.STATS.ROBUST[j,6]<-mc(CUR[CUR$MEAN.YFP.MEDIAN.WT>SUMMARY.STATS.ROBUST[j,2],"MEAN.YFP.MEDIAN.WT"])
  SHAM.SUMMARY.STATS.ROBUST[j,7]<-nrow(CUR)
  SHAM.SUMMARY.STATS.ROBUST[j,8]<-min(CUR[,"MEAN.YFP.MEDIAN.WT"])
  SHAM.SUMMARY.STATS.ROBUST[j,9]<-max(CUR[,"MEAN.YFP.MEDIAN.WT"])
  SHAM.SUMMARY.STATS.ROBUST[j,10]<-var(CUR[,"MEAN.YFP.MEDIAN.WT"])
  
  }
row.names(SUMMARY.STATS.ROBUST)<-SUMMARY.STATS.ROBUST[,1]
row.names(SUMMARY.STATS.ROBUST.Z)<-SUMMARY.STATS.ROBUST.Z[,1]

#Is proportion of EMS-treated cells within 2 MAD of median on either side correlated with sham MAD?
SD.Z<-cbind(SD.Z,SHAM.SUMMARY.STATS.ROBUST[,2:3])
names(SD.Z)<-c("ASSAY","PROP.2MAD.Z","SHAM.MEDIAN","SHAM.MAD")

cor.test(SD.Z$PROP.2MAD.Z,SD.Z$SHAM.MEDIAN)
cor.test(SD.Z$PROP.2MAD.Z,SD.Z$SHAM.MAD)
cor.test(SD.Z$PROP.2MAD.Z,SD.Z$SHAM.MEDIAN, method="spearman")
cor.test(SD.Z$PROP.2MAD.Z,SD.Z$SHAM.MAD,method="spearman")
#nope
cor.test(SHAM.SUMMARY.STATS.ROBUST$VAR,SUMMARY.STATS.ROBUST.Z$VAR)
cor.test(SHAM.SUMMARY.STATS.ROBUST$VAR,SUMMARY.STATS.ROBUST.Z$VAR,method="spearman")


cor.test(SHAM.SUMMARY.STATS.ROBUST$MAD,SUMMARY.STATS.ROBUST.Z$MAD)
cor.test(SHAM.SUMMARY.STATS.ROBUST$MAD,SUMMARY.STATS.ROBUST.Z$MAD,method="spearman")


#Check for correlation among variables
cor(SUMMARY.STATS.ROBUST.Z[,c(2,3,5,6)])

#Perform PCA

pca.Z = PCA(SUMMARY.STATS.ROBUST.Z[,c(2:6)], graph = FALSE)

pca.Z$eig
pca.Z$var$cor
pca.Z$var$contrib

windows()
par(mfrow=c(2,3))
plot.PCA(pca.Z,c(1,2),habillage="ind",col.hab = col_list,cex.lab=1.5,cex.axis=1.5)
plot.PCA(pca.Z,c(2,3),habillage="ind",col.hab = col_list,cex.lab=1.5,cex.axis=1.5)
plot.PCA(pca.Z,c(1,3),habillage="ind",col.hab = col_list,cex.lab=1.5,cex.axis=1.5)
plot.PCA(pca.Z,c(1,2),choix="var",cex.lab=1.5,cex.axis=1.5)
plot.PCA(pca.Z,c(2,3),choix="var",cex.lab=1.5,cex.axis=1.5)
plot.PCA(pca.Z,c(1,3),choix="var",cex.lab=1.5,cex.axis=1.5)

PCA.ORDER.DIM1<-names(sort(pca.Z$ind$coord[,1]))
PCA.ORDER.DIM2<-names(sort(pca.Z$ind$coord[,2]))
PCA.ORDER.DIM3<-names(sort(pca.Z$ind$coord[,3]))


#Reorder promoters by Principle Components 1-3
EMS<-droplevels(subset(ALL.PROMOTERS.WTDH3, ALL.PROMOTERS.WTDH3$CONDITION=="EMS"|ALL.PROMOTERS.WTDH3$CONDITION=="EMS.ALL"))
PCA.ORDER.DIM1.DATA<-EMS[,c("ASSAY","ZSCORE.YFP.MEDIAN")]
PCA.ORDER.DIM1.DATA<-PCA.ORDER.DIM1.DATA[order(match(EMS$ASSAY, PCA.ORDER.DIM1)),]
PCA.ORDER.DIM1.DATA$ASSAY<-factor(PCA.ORDER.DIM1.DATA$ASSAY,levels = as.character(PCA.ORDER.DIM1))

PCA.ORDER.DIM2.DATA<-EMS[,c("ASSAY","ZSCORE.YFP.MEDIAN")]
PCA.ORDER.DIM2.DATA<-PCA.ORDER.DIM2.DATA[order(match(EMS$ASSAY, PCA.ORDER.DIM2)),]
PCA.ORDER.DIM2.DATA$ASSAY<-factor(PCA.ORDER.DIM2.DATA$ASSAY,levels = as.character(PCA.ORDER.DIM2))

PCA.ORDER.DIM3.DATA<-EMS[,c("ASSAY","ZSCORE.YFP.MEDIAN")]
PCA.ORDER.DIM3.DATA<-PCA.ORDER.DIM3.DATA[order(match(EMS$ASSAY, PCA.ORDER.DIM3)),]
PCA.ORDER.DIM3.DATA$ASSAY<-factor(PCA.ORDER.DIM3.DATA$ASSAY,levels = as.character(PCA.ORDER.DIM3))

start.order<-levels(ALL.PROMOTERS.WTDH3$ASSAY)
DIM1.reord<-match(PCA.ORDER.DIM1,start.order)
DIM2.reord<-match(PCA.ORDER.DIM2,start.order)
DIM3.reord<-match(PCA.ORDER.DIM3,start.order)

#Examine as stacked histograms
DIM1<-ggplot(PCA.ORDER.DIM1.DATA, aes(x = ZSCORE.YFP.MEDIAN, y = fct_rev(ASSAY), fill = ASSAY)) + 
  geom_density_ridges(scale = 1.5) + geom_density_ridges(rel_min_height = 0.01)  +
  scale_fill_cyclical(values = col_list[DIM1.reord])  + 
  geom_vline(xintercept = 0, color = "white",linetype = "dashed") +
  theme_ridges(center_axis_labels = TRUE)+
  scale_x_continuous(limits = c(-10,10),expand = c(0.01, 0)) +
  scale_y_discrete(expand = c(0, 0)) +
  xlab("Expression Zscore") + ylab("") 

DIM2<-ggplot(PCA.ORDER.DIM2.DATA, aes(x = ZSCORE.YFP.MEDIAN, y = fct_rev(ASSAY), fill = ASSAY)) + 
  geom_density_ridges(scale = 1.5) + geom_density_ridges(rel_min_height = 0.01)  +
  scale_fill_cyclical(values = col_list[DIM2.reord])  + 
  geom_vline(xintercept = 0, color = "white",linetype = "dashed") +
  theme_ridges(center_axis_labels = TRUE)+
  scale_x_continuous(limits = c(-10,10),expand = c(0.01, 0)) +
  scale_y_discrete(expand = c(0, 0)) +
  xlab("Expression Zscore") + ylab("") 

DIM3<-ggplot(PCA.ORDER.DIM3.DATA, aes(x = ZSCORE.YFP.MEDIAN, y = fct_rev(ASSAY), fill = ASSAY)) + 
  geom_density_ridges(scale = 1.5) + geom_density_ridges(rel_min_height = 0.01)  +
  scale_fill_cyclical(values = col_list[DIM3.reord])  + 
  geom_vline(xintercept = 0, color = "white",linetype = "dashed") +
  theme_ridges(center_axis_labels = TRUE)+
  scale_x_continuous(limits = c(-10,10),expand = c(0.01, 0)) +
  scale_y_discrete(expand = c(0, 0)) +
  xlab("Expression Zscore") + ylab("") 

windows()
grid.arrange(DIM1,DIM2,DIM3, ncol=3)

#Look at summary stats ranked along PCA dimensions
SUMMARY.STATS.ROBUST.Z[PCA.ORDER.DIM1,]
SUMMARY.STATS.ROBUST.Z[PCA.ORDER.DIM2,]
SUMMARY.STATS.ROBUST.Z[PCA.ORDER.DIM3,]


###########################################
#                                         #
#                                         #
#           Neutral Simulations           #
#                                         #
#                                         #
###########################################


#See Supplementary File 5 for the code for the simulations themselves. 
#Routines below are used to plot data summarizing mean population phenotype 
#each generation extracted from the simulations.

#Plotting Neutral Results
setwd("C:/Users/andreahd/My Documents/working analyses/2017 analyses/DCv34/neutral predictions/excl contam/50Kgen_mult")
GENE<-c("GPD1","OST1","PFY1","RNR1","RNR2","STM1","TDH1","TDH2","TDH3","VMA7")
GENE_COL<-c("red3", "darkorange1", "gold", "#63BF4D", "springgreen4", "#4DBFBF", "#4D7ABF", "royalblue4", "maroon", "#A84DBF")
FIGSTR<-c("a","b","c","d","e","f","g","h","i","j","k")
#Figure 4:
# Evolutionary trajectories for full empirical model
# Population size: N = 1000

windows()
par(mfcol = c(5,2),
    oma = c(0,4,3,0) + 0.1,
    mar = c(0,4,1,1) + 0.1,
    omi = c(0.5,0,0,0) + 0.1)
for(j in c(1:10)){
  #for(j in c(1:10)){
  filename<-paste0("SUMMARY_",GENE[j],"_UNWEIGHTED_MULT_MUTRATE_SAMPLE_GEN50K_POPSIZE1000_REP500.txt")
  EVOL.DATA<-read.table(filename, sep="\t",header=FALSE)
  
  #plot(1:50000,log2(EVOL.DATA[,7],xlim=c(0,50000),xlab="Generations",ylim=c(0,max(log2(EVOL.DATA[,12])*1.1),ylab="Median Divergence",col=GENE_COL[j],type="l",lwd=1,cex.lab=1.2,cex=1.2)#main=paste0(GENE[j]," N=1000"))
  plot(1:50000,log2(EVOL.DATA[,7]),xlim=c(0,50000),ylim=c(0,max(log2(EVOL.DATA[,12])*1.05)),xaxt="n",xlab="",ylab="",col=GENE_COL[j],type="l",lwd=1,cex.lab=1.2,cex=1.2,las=1)#main=paste0(GENE[j]," N=1000"))
  #abline(h=100,lty=3)
  
  polygon(c(1:50000, rev(1:50000)), c(log2(EVOL.DATA[,2]), rev(log2(EVOL.DATA[,12]))),
          col = alpha(GENE_COL[j],0.2), border = NA)
  polygon(c(1:50000, rev(1:50000)), c(log2(EVOL.DATA[,3]), rev(log2(EVOL.DATA[,11]))),
          col = alpha(GENE_COL[j],0.2), border = NA)
  polygon(c(1:50000, rev(1:50000)), c(log2(EVOL.DATA[,4]), rev(log2(EVOL.DATA[,10]))),
          col = alpha(GENE_COL[j],0.2), border = NA)
  polygon(c(1:50000, rev(1:50000)), c(log2(EVOL.DATA[,5]), rev(log2(EVOL.DATA[,9]))),
          col = alpha(GENE_COL[j],0.2), border = NA)
  polygon(c(1:50000, rev(1:50000)), c(log2(EVOL.DATA[,6]), rev(log2(EVOL.DATA[,8]))),
          col = alpha(GENE_COL[j],0.2), border = NA)
  legend("bottomleft",GENE[j],cex=1.4,bty="n")
  
  #mtext(paste0(FIGSTR[j],"."),side=2,line=3, cex = 1.1,at=max(log2(EVOL.DATA[,12])*1.1), col = "black",las=2)
 
   if(j==1){
    mtext("Figure 4", side = 3, line = 0.7,at=0,cex=1.1,col="black")
  }
  
  if(j==5|j==10){
    mtext(seq(0,50000,10000), side = 1, line = 1, at=seq(0,50000,10000),cex = 0.8, col = "black")
    mtext("Generations",side = 1, line = 2.5,cex=1,col="black")
  }
  if(j<6){
    mtext("Divergence", side = 2, line = 3, cex=0.8,col="black")
  }
}

#Supporting figure:
# Evolutionary trajectories for full empirical model
# Population size: N = 100

windows()
par(mfcol = c(5,2),
    oma = c(0,4,2,0) + 0.1,
    mar = c(0,4,1,1) + 0.1,
    omi = c(0.5,0,0,0) + 0.1)
for(j in c(1:10)){

  filename<-paste0("SUMMARY_",GENE[j],"_UNWEIGHTED_MULT_MUTRATE_SAMPLE_GEN50K_POPSIZE100_REP500.txt")
  EVOL.DATA<-read.table(filename, sep="\t",header=FALSE)
  
  plot(1:50000,log2(EVOL.DATA[,7]),xlim=c(0,50000),ylim=c(0,max(log2(EVOL.DATA[,12])*1.1)),xaxt="n",xlab="",ylab="",col=GENE_COL[j],type="l",lwd=1,cex.lab=1.2,cex=1.2,las=1)#main=paste0(GENE[j]," N=1000"))

  polygon(c(1:50000, rev(1:50000)), c(log2(EVOL.DATA[,2]), rev(log2(EVOL.DATA[,12]))),
          col = alpha(GENE_COL[j],0.2), border = NA)
  polygon(c(1:50000, rev(1:50000)), c(log2(EVOL.DATA[,3]), rev(log2(EVOL.DATA[,11]))),
          col = alpha(GENE_COL[j],0.2), border = NA)
  polygon(c(1:50000, rev(1:50000)), c(log2(EVOL.DATA[,4]), rev(log2(EVOL.DATA[,10]))),
          col = alpha(GENE_COL[j],0.2), border = NA)
  polygon(c(1:50000, rev(1:50000)), c(log2(EVOL.DATA[,5]), rev(log2(EVOL.DATA[,9]))),
          col = alpha(GENE_COL[j],0.2), border = NA)
  polygon(c(1:50000, rev(1:50000)), c(log2(EVOL.DATA[,6]), rev(log2(EVOL.DATA[,8]))),
          col = alpha(GENE_COL[j],0.2), border = NA)
  legend("bottomleft",GENE[j],cex=1.4,bty="n")

  mtext(paste0(FIGSTR[j],"."),side=2,line=3, cex = 1.1,at=max(log2(EVOL.DATA[,12])*1.1), col = "black",las=2)
  
  
    if(j==1){
    mtext("Supplementary Figure 9", side = 3, line = 0.7,at=10000,cex=1.1,col="black")
  }
  
  if(j==5|j==10){
    mtext(seq(0,50000,10000), side = 1, line = 1, at=seq(0,50000,10000),cex = 0.8, col = "black")
    mtext("Generations",side = 1, line = 2.5,cex=1,col="black")
    }
  if(j<6){
    mtext("Divergence", side = 2, line = 3, cex=0.8,col="black")
  }
}

#Extract information about full distributions of population medians and sd for each replicate
#Then test for differences in model predictions and relatinoship to distribution parameters

library(data.table)

GENE.DESIG<-c("GPD1.SORT1.SC1","OST1.SORT1.SC1","PFY1.SORT1.SC1","RNR1.SORT1.SC1","RNR2.SORT1.SC1","STM1.SORT1.SC1","TDH1.SORT1.SC2","TDH2.SORT1.SC2","TDH3.SORT1.SC1","VMA7.SORT1.SC1")

DIVERGENCE<-data.frame(matrix(ncol = 13, nrow = 500*2*10))

names(DIVERGENCE)<-c("ASSAY","MODEL","ITER","N","EXP_50K","EXP_1K_MEDIAN","EXP_1K_SD","EXP_1K_MIN","EXP_1K_MAX","DIST","DIST_SD")
counter<-1

for(j in c(1:10)){
  CUR<-ALL.PROMOTERS.WTDH3[(ALL.PROMOTERS.WTDH3$CONDITION=="SHAM" & ALL.PROMOTERS.WTDH3$EXP==GENE.DESIG[j]),]
  DENS<-density(CUR$MEAN.YFP.MEDIAN.WT,from=0,to=4,adjust=2)
  ANCESTRAL_SHAM<-as.data.frame(replicate(500,(100*(sample(CUR[,"MEAN.YFP.MEDIAN.WT"],100, replace=TRUE)+rnorm(100, 0, DENS$bw)))))
  
  filename<-paste0(GENE.DESIG[j],"_MEDIAN.WT_PROJECTION.MEANS_UNWEIGHTED_MULT_MUTRATE_PARENTSAMPLE_GEN50K_POPSIZE1000_REP500.txt")
  DATA<-fread(filename,select=50000)
  DIVERGENCE[counter:(counter-1+nrow(DATA)),"ASSAY"]<-GENE[j]
  DIVERGENCE[counter:(counter-1+nrow(DATA)),"MODEL"]<-"EMPIRICAL"
  DIVERGENCE[counter:(counter-1+nrow(DATA)),"ITER"]<-1:nrow(DATA)
  DIVERGENCE[counter:(counter-1+nrow(DATA)),"N"]<-"100"
  DIVERGENCE[counter:(counter-1+nrow(DATA)),"EXP_50K"]<-DATA
  DIVERGENCE[counter:(counter-1+nrow(DATA)),"EXP_1K_MEDIAN"]<-sapply(ANCESTRAL_SHAM,median)
  DIVERGENCE[counter:(counter-1+nrow(DATA)),"EXP_1K_SD"]<-sapply(ANCESTRAL_SHAM,sd)
  DIVERGENCE[counter:(counter-1+nrow(DATA)),"EXP_1K_MIN"]<-sapply(ANCESTRAL_SHAM,range)[,1]
  DIVERGENCE[counter:(counter-1+nrow(DATA)),"EXP_1K_MAX"]<-sapply(ANCESTRAL_SHAM,range)[,2]
  DIVERGENCE[counter:(counter-1+nrow(DATA)),"DIST"] <- DIVERGENCE[counter:(counter-1+nrow(DATA)),"EXP_50K"]-DIVERGENCE[counter:(counter-1+nrow(DATA)),"EXP_1K_MEDIAN"]
  DIVERGENCE[counter:(counter-1+nrow(DATA)),"DIST_SD"] <- DIVERGENCE[counter:(counter-1+nrow(DATA)),"DIST"]/median(DIVERGENCE[counter:(counter-1+nrow(DATA)),"EXP_1K_SD"])
  DIVERGENCE[counter:(counter-1+nrow(DATA)),"MAD"] <-SUMMARY.STATS.ROBUST[SUMMARY.STATS.ROBUST$ASSAY==GENE[j],"MAD"]
  DIVERGENCE[counter:(counter-1+nrow(DATA)),"MC"] <-SUMMARY.STATS.ROBUST[SUMMARY.STATS.ROBUST$ASSAY==GENE[j],"MC"]  
  DIVERGENCE[counter:(counter-1+nrow(DATA)),"LMC"] <-SUMMARY.STATS.ROBUST[SUMMARY.STATS.ROBUST$ASSAY==GENE[j],"LMC"] 
  DIVERGENCE[counter:(counter-1+nrow(DATA)),"RMC"] <-SUMMARY.STATS.ROBUST[SUMMARY.STATS.ROBUST$ASSAY==GENE[j],"RMC"] 
  
  counter<-counter+nrow(DATA)
  filename<-paste0(GENE.DESIG[j],"_MEDIAN.WT_PROJECTION.MEANS_UNWEIGHTED_MULT_BROWNIAN_PARENTSAMPLE_GEN50K_POPSIZE1000_REP500.txt")
  DATA<-fread(filename,select=50000)
  DIVERGENCE[counter:(counter-1+nrow(DATA)),"ASSAY"]<-GENE[j]
  DIVERGENCE[counter:(counter-1+nrow(DATA)),"MODEL"]<-"BROWN"
  DIVERGENCE[counter:(counter-1+nrow(DATA)),"ITER"]<-1:nrow(DATA)
  DIVERGENCE[counter:(counter-1+nrow(DATA)),"N"]<-"100"
  DIVERGENCE[counter:(counter-1+nrow(DATA)),"EXP_50K"]<-DATA
  DIVERGENCE[counter:(counter-1+nrow(DATA)),"EXP_1K_MEDIAN"]<-sapply(ANCESTRAL_SHAM,median)
  DIVERGENCE[counter:(counter-1+nrow(DATA)),"EXP_1K_SD"]<-sapply(ANCESTRAL_SHAM,sd)
  DIVERGENCE[counter:(counter-1+nrow(DATA)),"EXP_1K_MIN"]<-sapply(ANCESTRAL_SHAM,range)[,1]
  DIVERGENCE[counter:(counter-1+nrow(DATA)),"EXP_1K_MAX"]<-sapply(ANCESTRAL_SHAM,range)[,2]
  DIVERGENCE[counter:(counter-1+nrow(DATA)),"DIST"] <- DIVERGENCE[counter:(counter-1+nrow(DATA)),"EXP_50K"]-DIVERGENCE[counter:(counter-1+nrow(DATA)),"EXP_1K_MEDIAN"]
  DIVERGENCE[counter:(counter-1+nrow(DATA)),"DIST_SD"] <- DIVERGENCE[counter:(counter-1+nrow(DATA)),"DIST"]/median(DIVERGENCE[counter:(counter-1+nrow(DATA)),"EXP_1K_SD"])
  DIVERGENCE[counter:(counter-1+nrow(DATA)),"MAD"] <-SUMMARY.STATS.ROBUST[SUMMARY.STATS.ROBUST$ASSAY==GENE[j],"MAD"]
  DIVERGENCE[counter:(counter-1+nrow(DATA)),"MC"] <-SUMMARY.STATS.ROBUST[SUMMARY.STATS.ROBUST$ASSAY==GENE[j],"MC"]
  DIVERGENCE[counter:(counter-1+nrow(DATA)),"LMC"] <-SUMMARY.STATS.ROBUST[SUMMARY.STATS.ROBUST$ASSAY==GENE[j],"LMC"]
  DIVERGENCE[counter:(counter-1+nrow(DATA)),"RMC"] <-SUMMARY.STATS.ROBUST[SUMMARY.STATS.ROBUST$ASSAY==GENE[j],"RMC"]
  
  counter<-counter+nrow(DATA)
}

DIVERGENCE$ASSAY<-as.factor(DIVERGENCE$ASSAY)
DIVERGENCE$MODEL<-as.factor(DIVERGENCE$MODEL)
write.table(DIVERGENCE,"DivergenceNeutral2Models_N1000wSHAM.txt",sep="\t",quote=FALSE,row.names=FALSE)
#DIVERGENCE<-read.table("DivergenceNeutral2Models_N1000wSHAM.txt",header=TRUE,sep="\t")

SUMMARY.EXP_50K<-aggregate(EXP_50K ~ MODEL+ASSAY,data=DIVERGENCE[DIVERGENCE$MODEL=="BROWN",],FUN = median)
SUMMARY.EXP_50K<-merge(SUMMARY.EXP_50K,aggregate(EXP_50K ~ MODEL+ASSAY,data=DIVERGENCE[DIVERGENCE$MODEL=="EMPIRICAL",],FUN = median),by="ASSAY",suffixes=c(".BROWN",".EMPIRICAL"))

SUMMARY.EXP_50K.SD<-aggregate(EXP_50K ~ MODEL+ASSAY,data=DIVERGENCE[DIVERGENCE$MODEL=="BROWN",],FUN = sd)
SUMMARY.EXP_50K.SD<-merge(SUMMARY.EXP_50K.SD,aggregate(EXP_50K ~ MODEL+ASSAY,data=DIVERGENCE[DIVERGENCE$MODEL=="EMPIRICAL",],FUN = sd),by="ASSAY",suffixes=c(".BROWN",".EMPIRICAL"))
SUMMARY.EXP_50K.SD<-SUMMARY.EXP_50K.SD[,c(1,3,5)]
colnames(SUMMARY.EXP_50K.SD)<-c("ASSAY","EXP_50K.SD.BROWN","EXP_50K.SD.EMPIRICAL")

SUMMARY.COMBINED<-merge(SUMMARY.STATS.ROBUST,SUMMARY.EXP_50K[,c("ASSAY","EXP_50K.BROWN","EXP_50K.EMPIRICAL")],by="ASSAY")
SUMMARY.COMBINED<-merge(SUMMARY.COMBINED,SUMMARY.EXP_50K.SD,by="ASSAY")


library(plotrix)

#Do Brownian motion and empirical mutational predictions differ significantly after 50K gen evolution?
LM<-lm(EXP_50K~MODEL*ASSAY,data=DIVERGENCE)
windows()
par(mfrow=c(2,2))
plot(LM) # QQ plot suggests departure from normality, try log transforming
LM<-lm(log(EXP_50K)~MODEL*ASSAY,data=DIVERGENCE)
plot(LM)
LM2<-update(LM,~.-MODEL:ASSAY)
anova(LM, LM2, test = "F")  # interaction significant, useful to explain variability in divergence


#Checking a non-parametric test: for each promoter, do Brownian and empirical distributions impact divergence?
MODEL.FITS<-data.frame(matrix(ncol = 5, nrow = 10))
names(MODEL.FITS)<-c("ASSAY","Normality","W","P","PADJ")

for(j in c(1:10)){
  WIL<-wilcox.test(DIVERGENCE[DIVERGENCE$ASSAY==GENE[j]&DIVERGENCE$MODEL=="EMPIRICAL","EXP_50K"],DIVERGENCE[DIVERGENCE$ASSAY==GENE[j]&DIVERGENCE$MODEL=="BROWN","EXP_50K"])
  MODEL.FITS[j,"ASSAY"]<-GENE[j]
  MODEL.FITS[j,"Normality"]<-shapiro.test(DIVERGENCE[DIVERGENCE$ASSAY==GENE[j],"EXP_50K"])
  MODEL.FITS[j,"W"]<-WIL$statistic
  MODEL.FITS[j,"P"]<-WIL$p.value
}
MODEL.FITS[,"PADJ"]<-p.adjust(MODEL.FITS$P,method="BH",n=nrow(MODEL.FITS))
#all satisfy normality so parametric model is fine, but results largely similar (only GPD1 not pairwise sig)


#Which distribution shape metrics best predict magnitude of divergence for each model?
#Using summaries of median behavior by promoters since distribution statistics are fixed per promoter
LM<-lm(log(SUMMARY.COMBINED$EXP_50K.EMPIRICAL)~(SUMMARY.COMBINED$MAD+SUMMARY.COMBINED$MC+SUMMARY.COMBINED$LMC+SUMMARY.COMBINED$RMC))
#plot(LM)
LM2<-lm(log(SUMMARY.COMBINED$EXP_50K.EMPIRICAL)~(SUMMARY.COMBINED$MC+SUMMARY.COMBINED$LMC))
summary(LM2)

p.adjust(0.001247,method="BH",n=3)

LM<-lm(log(SUMMARY.COMBINED$EXP_50K.BROWN)~(SUMMARY.COMBINED$MAD+SUMMARY.COMBINED$MC+SUMMARY.COMBINED$LMC+SUMMARY.COMBINED$RMC))
#plot(LM)
LM2<-lm(log(SUMMARY.COMBINED$EXP_50K.BROWN)~(SUMMARY.COMBINED$MAD+SUMMARY.COMBINED$MC))
#plot(LM2)
summary(LM2)
p.adjust(0.0004599,method="BH",n=2)


#Plot results 
#Figure 5A
windows()
par(mfrow=c(1,2),
    oma = c(5,4,4,2) + 0.1)
#Summary of median phenotypes after 50K gen evolution with 2 neutral models
plot(log(SUMMARY.COMBINED$EXP_50K.EMPIRICAL),log(SUMMARY.COMBINED$EXP_50K.BROWN),
     xlim=c(0,9.1),ylim=c(1.5,7.5),
     pch=20,col=GENE_COL,
     xlab="Empirical Gen 50K (Log Phenotype))",ylab="Brownian Gen 50K (Log Phenotype)")
abline(v=log(100),lty=3)
abline(h=log(100),lty=3)
plotCI(log(SUMMARY.COMBINED$EXP_50K.EMPIRICAL),y=log(SUMMARY.COMBINED$EXP_50K.BROWN),
       uiw=2*(1.96*log(SUMMARY.COMBINED$EXP_50K.SD.EMPIRICAL)/(sqrt(500))),
       err="x",add=TRUE,scol=GENE_COL,pch=21)
plotCI(log(SUMMARY.COMBINED$EXP_50K.EMPIRICAL),y=log(SUMMARY.COMBINED$EXP_50K.BROWN),
       uiw=2*(1.96*log(SUMMARY.COMBINED$EXP_50K.SD.BROWN)/(sqrt(500))),
       err="y",add=TRUE,scol=GENE_COL,pch=21)
mtext("a.",side=2,line=1, cex = 1.2,at=8.2, col = "black",las=2)
mtext("Figure 6",side=2,line=1, cex = 1.2,at=9, col = "black",las=2)


#Figure 5B
#Divergence as predicted by skewness for 2 models

plot(SUMMARY.COMBINED$MC,log(SUMMARY.COMBINED$EXP_50K.BROWN),
     pch=21,col=GENE_COL,xlim=c(-0.28,0.28),ylim=c(0,9.8),xlab=" Skewness (MC)",ylab="Gen 50K (Log Phenotype)")
LM<-lm(log(SUMMARY.COMBINED$EXP_50K.BROWN)~SUMMARY.COMBINED$MC)
summary(LM)
abline(LM,lty=3,col="lightgrey")
points(SUMMARY.COMBINED$MC,log(SUMMARY.COMBINED$EXP_50K.EMPIRICAL),
       pch=19,col=GENE_COL)
LM<-lm(log(SUMMARY.COMBINED$EXP_50K.EMPIRICAL)~SUMMARY.COMBINED$MC)
summary(LM)
abline(LM,lty=1,col="lightgrey")
arrows(SUMMARY.COMBINED$MC, log(SUMMARY.COMBINED$EXP_50K.BROWN), 
       x1 = SUMMARY.COMBINED$MC, y1 = log(SUMMARY.COMBINED$EXP_50K.EMPIRICAL),
       col=GENE_COL,length=0)
points(SUMMARY.COMBINED$MC,log(SUMMARY.COMBINED$EXP_50K.BROWN),
       pch=21,col=GENE_COL,bg="white")
legend("topleft",c("Brownian","Empirical"),pch=c(21,19),col=c("black","black"),bty="n")
mtext("b.",side=2,line=1, cex = 1.2,at=11.1, col = "black",las=2)


###############################
#                             #
##   Promoter Properties      #
#                             #
###############################


FULL.PROPERTIES<-read.table("PATH/GeneListAnalysis_Mutagenized.txt",header=TRUE,sep="\t")
FULL.PROPERTIES$ASSAY<-substr(FULL.PROPERTIES$SHORT,1,4)
FULL.PROPERTIES$NUCLEOSOME.CLASS<-as.character(FULL.PROPERTIES$NUCLEOSOME.CLASS)
FULL.PROPERTIES[FULL.PROPERTIES$ASSAY=="OST1","NUCLEOSOME.CLASS"]<-"NA"
FULL.PROPERTIES[FULL.PROPERTIES$ASSAY=="GPD1","NUCLEOSOME.CLASS"]<-"OPN"
FULL.PROPERTIES[FULL.PROPERTIES$ASSAY=="GPD1","NUCLEOSOME.CLASS"]<-"OPN"
FULL.PROPERTIES[FULL.PROPERTIES$ASSAY=="RNR1","NUCLEOSOME.CLASS"]<-"OPN"
FULL.PROPERTIES[FULL.PROPERTIES$ASSAY=="RNR2","NUCLEOSOME.CLASS"]<-"OPN"
FULL.PROPERTIES[FULL.PROPERTIES$ASSAY=="STM1","NUCLEOSOME.CLASS"]<-"NA"
FULL.PROPERTIES[FULL.PROPERTIES$ASSAY=="TDH1","NUCLEOSOME.CLASS"]<-"OPN"
FULL.PROPERTIES[FULL.PROPERTIES$ASSAY=="TDH2","NUCLEOSOME.CLASS"]<-"NA"
FULL.PROPERTIES[FULL.PROPERTIES$ASSAY=="TDH3","NUCLEOSOME.CLASS"]<-"NA"
FULL.PROPERTIES$NUCLEOSOME.CLASS<-as.factor(FULL.PROPERTIES$NUCLEOSOME.CLASS)
FULL.PROPERTIES$TATA.MISMATCH<-as.character(FULL.PROPERTIES$TATA.MISMATCH)
FULL.PROPERTIES[FULL.PROPERTIES$ASSAY=="TDH3","TATA.MISMATCH"]<-"0"
FULL.PROPERTIES[FULL.PROPERTIES$ASSAY=="VMA7","TATA.MISMATCH"]<-"2" #">2"
FULL.PROPERTIES[FULL.PROPERTIES$ASSAY=="TDH1","TATA.MISMATCH"]<-"0" #">2"
FULL.PROPERTIES[FULL.PROPERTIES$ASSAY=="TDH2","TATA.MISMATCH"]<-"0" #">2"
FULL.PROPERTIES[FULL.PROPERTIES$ASSAY=="TDH3","TATA.BIS"]<-as.factor("TATA-containing")

BOXPLOT.DATA<-merge(SUMMARY.STATS.ROBUST,FULL.PROPERTIES[,c("ASSAY","MEAN.GFP","CV.GFP","NUCLEOSOME.CLASS","DENSITY.NUCLEOSOME.PROMOTER","N.NUCLEOSOME.WINDOW","TATA.MISMATCH","FITNESS.HOMO","DUPLICATE","N.TRANS.FACTORS","MEAN.EXPR.SKELLY","VAR.EXPR.SKELLY","VM.MED")],by="ASSAY")
BOXPLOT.DATA$MEAN.GFP.FACTOR<-ifelse(BOXPLOT.DATA$MEAN.GFP < median(BOXPLOT.DATA$MEAN.GFP),"LOW","HIGH")
BOXPLOT.DATA$CV.GFP.FACTOR<-ifelse(BOXPLOT.DATA$CV.GFP < median(BOXPLOT.DATA$CV.GFP),"LOW","HIGH")
BOXPLOT.DATA$TATA.FACTOR<-as.factor(ifelse(BOXPLOT.DATA$TATA.MISMATCH=="0","TATA+","TATA-"))
BOXPLOT.DATA$FITNESS.FACTOR<-ifelse(BOXPLOT.DATA$FITNESS.HOMO<0.85,"LOW","HIGH")
BOXPLOT.DATA$N.TRANS.FACTORS.FACTOR<-as.factor(ifelse(BOXPLOT.DATA$N.TRANS.FACTORS < median(BOXPLOT.DATA$N.TRANS.FACTORS),"LOW","HIGH"))

BOXPLOT.DATA$NUCLEOSOME.OCC.MEASURE<-"NA"
#Hornung et al definition of nucleosome occupancy based on 3 datasets
BOXPLOT.DATA[BOXPLOT.DATA$ASSAY=="PFY1","NUCLEOSOME.OCC.MEASURE"]<-(-1.53)
BOXPLOT.DATA[BOXPLOT.DATA$ASSAY=="OST1","NUCLEOSOME.OCC.MEASURE"]<-(-1.00)
BOXPLOT.DATA[BOXPLOT.DATA$ASSAY=="GPD1","NUCLEOSOME.OCC.MEASURE"]<-(1.13)
BOXPLOT.DATA[BOXPLOT.DATA$ASSAY=="VMA7","NUCLEOSOME.OCC.MEASURE"]<-(-0.911)
BOXPLOT.DATA[BOXPLOT.DATA$ASSAY=="RNR2","NUCLEOSOME.OCC.MEASURE"]<-(0.959)
BOXPLOT.DATA$NUCLEOSOME.OCC.MEASURE<-as.numeric(BOXPLOT.DATA$NUCLEOSOME.OCC.MEASURE)

#Tirosh and Barkai 2008 definition of nucleosome occupancy based on 1 dataset
BOXPLOT.DATA$NUCLEOSOME.OCC.MEASURE2<-"NA"
BOXPLOT.DATA[BOXPLOT.DATA$ASSAY=="RNR1","NUCLEOSOME.OCC.MEASURE2"]<-(2.1829)
BOXPLOT.DATA[BOXPLOT.DATA$ASSAY=="STM1","NUCLEOSOME.OCC.MEASURE2"]<-(2.1787)
BOXPLOT.DATA[BOXPLOT.DATA$ASSAY=="TDH1","NUCLEOSOME.OCC.MEASURE2"]<-(1.2742)
BOXPLOT.DATA[BOXPLOT.DATA$ASSAY=="PFY1","NUCLEOSOME.OCC.MEASURE2"]<-(-1.2348)
BOXPLOT.DATA[BOXPLOT.DATA$ASSAY=="OST1","NUCLEOSOME.OCC.MEASURE2"]<-(-0.7840)
BOXPLOT.DATA[BOXPLOT.DATA$ASSAY=="GPD1","NUCLEOSOME.OCC.MEASURE2"]<-(1.7503)
BOXPLOT.DATA[BOXPLOT.DATA$ASSAY=="VMA7","NUCLEOSOME.OCC.MEASURE2"]<-(-0.7144)
BOXPLOT.DATA[BOXPLOT.DATA$ASSAY=="RNR2","NUCLEOSOME.OCC.MEASURE2"]<-(1.2379)

BOXPLOT.DATA$NUCLEOSOME.OCC.MEASURE2<-as.numeric(BOXPLOT.DATA$NUCLEOSOME.OCC.MEASURE2)

BOXPLOT.DATA$NUCLEOSOME.FACTOR<-ifelse(BOXPLOT.DATA$NUCLEOSOME.OCC.MEASURE2 < median(BOXPLOT.DATA[which(BOXPLOT.DATA$NUCLEOSOME.OCC.MEASURE2!="NA"),"NUCLEOSOME.OCC.MEASURE2"]),"LOW","HIGH")

cols <-c("GPD1" ="red3","OST1"="darkorange1","PFY1"="gold","RNR1"="#63BF4D","RNR2"="springgreen4",
         "STM1"="#4DBFBF","TDH1"="#4D7ABF","TDH2"="royalblue4","TDH3"="maroon","VMA7"="#A84DBF")

A1<-ggplot(BOXPLOT.DATA, aes(x=MEAN.GFP.FACTOR, y=MAD)) + 
  geom_boxplot(fill="gray")+ geom_point(aes(x=MEAN.GFP.FACTOR, y=MAD),col=cols)+#geom_smooth(method=lm)+
  ylim(min(BOXPLOT.DATA$MAD)*0.95,max(BOXPLOT.DATA$MAD)*1.05)+ 
  labs(x="Mean Expression", y = "MAD")+theme_bw()+theme(axis.title.x=element_blank(),panel.grid.major = element_blank(), panel.grid.minor = element_blank())
B1<-ggplot(BOXPLOT.DATA, aes(x=CV.GFP.FACTOR, y=MAD)) + 
  geom_boxplot(fill="gray")+ geom_point(aes(x=CV.GFP.FACTOR, y=MAD),col=cols)+
  ylim(min(BOXPLOT.DATA$MAD)*0.95,max(BOXPLOT.DATA$MAD)*1.05)+
  labs(x="Expression CV", y = "MAD")+theme_bw()+
  theme(panel.grid.major = element_blank(), panel.grid.minor = element_blank(),axis.title.x=element_blank(),axis.title.y=element_blank(),axis.text.y=element_blank(),axis.ticks.y=element_blank())
C1<-ggplot(BOXPLOT.DATA, aes(x=TATA.FACTOR, y=MAD)) + 
  geom_boxplot(fill="gray")+ geom_point(aes(x=TATA.FACTOR, y=MAD),col=cols)+
  ylim(min(BOXPLOT.DATA$MAD)*0.95,max(BOXPLOT.DATA$MAD)*1.05)+
  labs(x="Canonical TATA", y = "MAD")+
  theme_bw()+theme(panel.grid.major = element_blank(), panel.grid.minor = element_blank())+
  theme(axis.title.y=element_blank(),axis.title.x=element_blank(),axis.text.y=element_blank(),axis.ticks.y=element_blank())
D1<-ggplot(BOXPLOT.DATA, aes(x=NUCLEOSOME.OCC.MEASURE2, y=MAD)) + 
  geom_point(aes(x=NUCLEOSOME.OCC.MEASURE2,y=MAD),col=cols)+geom_smooth(method=lm, se=FALSE,lty=3)+
  ylim(min(BOXPLOT.DATA$MAD)*0.95,max(BOXPLOT.DATA$MAD)*1.05)+
  labs(x="Nucleosome Occupancy", y = "MAD")+
  theme_bw()+theme(panel.grid.major = element_blank(), panel.grid.minor = element_blank())+
  theme(axis.title.y=element_blank(),axis.title.x=element_blank(),axis.text.y=element_blank(),axis.ticks.y=element_blank())
E1<-ggplot(BOXPLOT.DATA, aes(x=N.TRANS.FACTORS.FACTOR, y=MAD)) + 
  geom_boxplot(fill="gray")+ geom_point(aes(x=N.TRANS.FACTORS.FACTOR,y=MAD),col=cols)+
  ylim(min(BOXPLOT.DATA$MAD)*0.95,max(BOXPLOT.DATA$MAD)*1.05)+
  labs(x="N Trans Factors", y = "MAD")+
  theme_bw()+theme(panel.grid.major = element_blank(), panel.grid.minor = element_blank())+
  theme(axis.title.y=element_blank(),axis.title.x=element_blank(),axis.text.y=element_blank(),axis.ticks.y=element_blank())
F1<-ggplot(BOXPLOT.DATA, aes(x=DUPLICATE, y=MAD)) + 
  geom_boxplot(fill="gray")+ geom_point(aes(x=DUPLICATE, y=MAD),col=cols)+
  ylim(min(BOXPLOT.DATA$MAD)*0.95,max(BOXPLOT.DATA$MAD)*1.05)+
  labs(x="Duplicated Gene", y = "MAD")+theme_bw()+theme(panel.grid.major = element_blank(), panel.grid.minor = element_blank())+
  theme(axis.title.y=element_blank(),axis.title.x=element_blank(),axis.text.y=element_blank(),axis.ticks.y=element_blank())
G1<-ggplot(BOXPLOT.DATA, aes(x=FITNESS.HOMO, y=MAD)) + 
  geom_point(aes(x=FITNESS.HOMO, y=MAD),col=cols)+geom_smooth(method=lm, se=FALSE,lty=3)+
  ylim(min(BOXPLOT.DATA$MAD)*0.95,max(BOXPLOT.DATA$MAD)*1.05)+
  labs(x="Deletion Fitness", y = "MAD")+
  theme_bw()+theme(panel.grid.major = element_blank(), panel.grid.minor = element_blank())+
  theme(axis.title.y=element_blank(),axis.title.x=element_blank(),axis.text.y=element_blank(),axis.ticks.y=element_blank())
A2<-ggplot(BOXPLOT.DATA, aes(x=MEAN.GFP.FACTOR, y=MC)) + 
  geom_boxplot(fill="gray")+ geom_point(aes(x=MEAN.GFP.FACTOR, y=MC),col=cols)+#geom_smooth(method=lm)+
  ylim(min(BOXPLOT.DATA$MC)*0.95,max(BOXPLOT.DATA$MC)*1.05)+ 
  labs(x="Mean Expression", y = "MC")+theme_bw()+theme(panel.grid.major = element_blank(), panel.grid.minor = element_blank())
B2<-ggplot(BOXPLOT.DATA, aes(x=CV.GFP.FACTOR, y=MC)) + 
  geom_boxplot(fill="gray")+ geom_point(aes(x=CV.GFP.FACTOR, y=MC),col=cols)+
  ylim(min(BOXPLOT.DATA$MC)*0.95,max(BOXPLOT.DATA$MC)*1.05)+
  labs(x="Expression CV", y = "MC")+theme_bw()+
  theme(panel.grid.major = element_blank(), panel.grid.minor = element_blank(),axis.title.y=element_blank(),axis.text.y=element_blank(),axis.ticks.y=element_blank())
C2<-ggplot(BOXPLOT.DATA, aes(x=TATA.FACTOR, y=MC)) + 
  geom_boxplot(fill="gray")+ geom_point(aes(x=TATA.FACTOR, y=MC),col=cols)+
  ylim(min(BOXPLOT.DATA$MC)*0.95,max(BOXPLOT.DATA$MC)*1.05)+
  labs(x="Canonical TATA", y = "MC")+
  theme_bw()+theme(panel.grid.major = element_blank(), panel.grid.minor = element_blank())+
  theme(axis.title.y=element_blank(),axis.text.y=element_blank(),axis.ticks.y=element_blank())
D2<-ggplot(BOXPLOT.DATA, aes(x=NUCLEOSOME.OCC.MEASURE2, y=MC)) + 
  geom_point(aes(x=NUCLEOSOME.OCC.MEASURE2,y=MC),col=cols)+geom_smooth(method=lm, se=FALSE,lty=3)+
  ylim(min(BOXPLOT.DATA$MC)*0.95,max(BOXPLOT.DATA$MC)*1.05)+
  labs(x="Nucleosome Occupancy", y = "MC")+
  theme_bw()+theme(panel.grid.major = element_blank(), panel.grid.minor = element_blank())+
  theme(axis.title.y=element_blank(),axis.text.y=element_blank(),axis.ticks.y=element_blank())
E2<-ggplot(BOXPLOT.DATA, aes(x=N.TRANS.FACTORS.FACTOR, y=MC)) + 
  geom_boxplot(fill="gray")+ geom_point(aes(x=N.TRANS.FACTORS.FACTOR,y=MC),col=cols)+
  ylim(min(BOXPLOT.DATA$MC)*0.95,max(BOXPLOT.DATA$MC)*1.05)+
  labs(x="N Trans Factors", y = "MC")+
  theme_bw()+theme(panel.grid.major = element_blank(), panel.grid.minor = element_blank())+
  theme(axis.title.y=element_blank(),axis.text.y=element_blank(),axis.ticks.y=element_blank())
F2<-ggplot(BOXPLOT.DATA, aes(x=DUPLICATE, y=MC)) + 
  geom_boxplot(fill="gray")+ geom_point(aes(x=DUPLICATE, y=MC),col=cols)+
  ylim(min(BOXPLOT.DATA$MC)*0.95,max(BOXPLOT.DATA$MC)*1.05)+
  labs(x="Duplicated Gene", y = "MC")+theme_bw()+theme(panel.grid.major = element_blank(), panel.grid.minor = element_blank())+
  theme(axis.title.y=element_blank(),axis.text.y=element_blank(),axis.ticks.y=element_blank())
G2<-ggplot(BOXPLOT.DATA, aes(x=FITNESS.HOMO, y=MC)) + 
  geom_point(aes(x=FITNESS.HOMO, y=MC),col=cols)+geom_smooth(method=lm, se=FALSE,lty=3)+
  ylim(min(BOXPLOT.DATA$MC)*0.95,max(BOXPLOT.DATA$MC)*1.05)+
  labs(x="Deletion Fitness", y = "MC")+
  theme_bw()+theme(panel.grid.major = element_blank(), panel.grid.minor = element_blank())+
  theme(axis.title.y=element_blank(),axis.text.y=element_blank(),axis.ticks.y=element_blank())

grid.arrange(A1,B1,C1,E1,F1,D1,G1,A2,B2,C2,E2,F2,D2,G2,nrow=2)


#Model simplification with multiple test correction to ID any well-supported relationships

#MAD
anova(lm(MAD~FITNESS.HOMO+CV.GFP.FACTOR+NUCLEOSOME.OCC.MEASURE2+N.TRANS.FACTORS.FACTOR+MEAN.GFP.FACTOR+TATA.FACTOR,data=BOXPLOT.DATA))
#no significant result at all

#MC
anova(lm(MC~FITNESS.HOMO+CV.GFP.FACTOR+NUCLEOSOME.OCC.MEASURE2+N.TRANS.FACTORS.FACTOR+MEAN.GFP.FACTOR+TATA.FACTOR,data=BOXPLOT.DATA))
anova(lm(MC~FITNESS.HOMO+NUCLEOSOME.OCC.MEASURE2+N.TRANS.FACTORS.FACTOR+MEAN.GFP.FACTOR+TATA.FACTOR,data=BOXPLOT.DATA))
anova(lm(MC~FITNESS.HOMO+NUCLEOSOME.OCC.MEASURE2+N.TRANS.FACTORS.FACTOR+TATA.FACTOR,data=BOXPLOT.DATA))
anova(lm(MC~FITNESS.HOMO+NUCLEOSOME.OCC.MEASURE2+N.TRANS.FACTORS.FACTOR,data=BOXPLOT.DATA))
anova(lm(MC~FITNESS.HOMO+NUCLEOSOME.OCC.MEASURE2,data=BOXPLOT.DATA))
anova(lm(MC~FITNESS.HOMO,data=BOXPLOT.DATA))
p.adjust(0.02319,method="BH",n=6)
#no significant result after multiple correction

windows()
ggplot(BOXPLOT.DATA, aes(x=FITNESS.HOMO, y=MC)) + 
  geom_point(aes(x=FITNESS.HOMO, y=MC),col=cols)+ geom_smooth(method=lm)+
  ylim(min(BOXPLOT.DATA$MC)*0.95,max(BOXPLOT.DATA$MC)*1.05)+
  labs(x="Deletion Fitness", y = "Skewness (MC)")+
  theme_bw()+theme(panel.grid.major = element_blank(), panel.grid.minor = element_blank())

#Explore Polymorphism

A<-ggplot(BOXPLOT.DATA, aes(x=MAD, y=VAR.EXPR.SKELLY)) + 
 geom_point(col=cols)+ylim(-0.05,0.2)+
  geom_smooth(method='lm',formula=y~x,se=FALSE)+
  geom_text(aes(label=ifelse(MAD>0.09,as.character(ASSAY),""),hjust=1, vjust=0),nudge_x = -0.001)+
  labs(x="Mutational Variance", y = "Polymorphism")+
  theme_bw()+theme(panel.grid.major = element_blank(), panel.grid.minor = element_blank())

A + geom_smooth(data=BOXPLOT.DATA[BOXPLOT.DATA$ASSAY!="TDH1",], aes(x=MAD, y=VAR.EXPR.SKELLY),method='lm',formula=y~x, se=FALSE,lty=3) 
  
B<-ggplot(BOXPLOT.DATA[BOXPLOT.DATA$ASSAY!="TDH1",], aes(x=MAD, y=VAR.EXPR.SKELLY)) +
  geom_point(col=cols[c(1:6,8:10)])+ylim(-0.05,0.2)+
  geom_smooth(method='lm',formula=y~x,lty=3,se=FALSE)+
  ggtitle("Without TDH1")+
  labs(x="Mutational Variance", y = "Polymorphism")+
  theme_bw()+theme(panel.grid.major = element_blank(), panel.grid.minor = element_blank())
C<-ggplot(BOXPLOT.DATA, aes(x=MC, y=VAR.EXPR.SKELLY)) +
  geom_point(col=cols)+ylim(-0.02,0.18)+
  geom_smooth(method='lm',formula=y~x,lty=3,se=FALSE)+
  labs(x="Mutational Skew", y = "Polymorphism")+
  theme_bw()+theme(panel.grid.major = element_blank(), panel.grid.minor = element_blank())

D<-ggplot(BOXPLOT.DATA[BOXPLOT.DATA$ASSAY!="TDH1",], aes(x=MC, y=VAR.EXPR.SKELLY)) +
  geom_point(col=cols[c(1:6,8:10)])+ylim(-0.02,0.18)+
  geom_smooth(method='lm',formula=y~x,lty=3,se=FALSE)+
  labs(x="Mutational Skew", y = "Polymorphism")+
  theme_bw()+theme(panel.grid.major = element_blank(), panel.grid.minor = element_blank())

#windows()
grid.arrange(A,B,C,D,nrow=2)
grid.arrange(A,B,nrow=1)


summary(lm(MAD~VAR.EXPR.SKELLY,data=BOXPLOT.DATA))
summary(lm(MAD~VAR.EXPR.SKELLY,data=BOXPLOT.DATA[BOXPLOT.DATA$ASSAY!="TDH1",]))
summary(lm(MC~VAR.EXPR.SKELLY,data=BOXPLOT.DATA))
summary(lm(MC~VAR.EXPR.SKELLY,data=BOXPLOT.DATA[BOXPLOT.DATA$ASSAY!="TDH1",]))


#Custom functions
#############################################################################
DOWNSAMPLE.DIST.PERMUTE.MEAN <- function(GROUP.1,GROUP.2) {
  
  N.DOWNSAMPLE <-min(length(GROUP.1),length(GROUP.2))
  
  N.PERM <- 10000
  
  OBS.DIFF <- rep(NA,N.PERM)
  PERM.DIFF <- rep(NA,N.PERM)
  
  for (i in 1:N.PERM)
  {
    SAMPLE.1 <-sample(GROUP.1,size=N.DOWNSAMPLE,replace=TRUE)
    SAMPLE.2 <-sample(GROUP.2,size=N.DOWNSAMPLE,replace=TRUE)
    
    OBS.DIFF[i] <- mean(SAMPLE.1)-mean(SAMPLE.2)
    
    POOL <- c(SAMPLE.1,SAMPLE.2)
    
    POS.1 <- sample(1:length(POOL),size=N.DOWNSAMPLE,replace=TRUE)
    PERM.1 <- POOL[POS.1]
    PERM.2 <- POOL[-POS.1]
    
    PERM.DIFF[i] <- mean(PERM.1)-mean(PERM.2)
  }
  
  EFFECT.SIZE <-mean(OBS.DIFF)
  
  P.VAL <- (length(which(abs(PERM.DIFF) > abs(OBS.DIFF))) + 1)/(N.PERM + 1)
  
  RESULTS <- list(EFFECT.SIZE,P.VAL)
  
  return(RESULTS)}
```
