## Supplemental tables, figures, files, and methods for "Empirical measures of mutational effects define neutral models of regulatory evolution in *Saccharomyces cerevisiae*": Supplementary File 5.R

```
#######################################################################
# Hodgins-Davis, Duveau, Walker, and Wittkopp. 2019.  
#
# Supplementary File 5
# includes R functions for evolutionary simulations modeling evolution
# from different distributions of mutational effects
#
# Simulations were run on a cluster iterating different jobs over j, which 
# specified the choice of promoter distribution.
#
# Two varieties of simulations are provided below, one drawing mutational
# effects directly from empirical data sampled for each promoter ("Empirical") and the
# other using the empirical data to identify variance in mutational effects and drawing
# mutational effects from a normal distribution with the empirical variance ("Brownian").
#
# Input to the script includes the distribution of expression values
# estimated for sham and mutant populations for each promoter.
#
# Output includes one file with the complete summary of population means
# at every generation of each independent simulation and one file summarizing
# the proportion of independent simulations captured at a range of quantiles.
#
#
# This code was written for R version 3.3.3.


############################
#   Empirical Simulation   #
############################


load("INPUT.RData")

REP<-500
GEN<-50000
N_POP<-1000

genome<-12495682
mutrate<-1.67*10^-10

j<-as.numeric(Sys.getenv("PBS_ARRAYID"))


#Multiplicative
#for(j in 1:length(levels(ALL.PROMOTERS.WTDH3$EXP))){
 
  CUR<-ALL.PROMOTERS.WTDH3[(ALL.PROMOTERS.WTDH3$CONDITION=="EMS"|ALL.PROMOTERS.WTDH3$CONDITION=="SHAM") & ALL.PROMOTERS.WTDH3$EXP==levels(ALL.PROMOTERS.WTDH3$EXP)[j],]
  SHAM.BW<-DENSITY.WTDH3.MEDIAN[(DENSITY.WTDH3.MEDIAN$CONDITION=="SHAM" & DENSITY.WTDH3.MEDIAN$EXP==levels(ALL.PROMOTERS.WTDH3$EXP)[j]),"BW"]
  EMS.BW<-DENSITY.WTDH3.MEDIAN[(DENSITY.WTDH3.MEDIAN$EXP==levels(ALL.PROMOTERS.WTDH3$EXP)[j]),"BW"]
  MUT<-CUR[CUR$CONDITION=="EMS",]
  
  SIM <- data.frame(matrix(ncol = GEN, nrow = REP))
  
  for(k in 1:REP){
    ANCESTRAL_SHAM<-sample(CUR[CUR$CONDITION=="SHAM","MEAN.YFP.MEDIAN.WT"],N_POP, replace=TRUE)+rnorm(N_POP, 0, SHAM.BW)
    POP <- data.frame(matrix(ncol = GEN, nrow = N_POP))
    POP[1:N_POP,1]<-ANCESTRAL_SHAM*100
    #POP[1:N_POP,1]<-100
    
    for(i in 2:GEN){
      POP[which(POP[,i-1]<0),i-1]<-0
      POP[,i]<-sample(POP[,(i-1)],replace=TRUE)
      N_MUT<-sum(rbinom(N_POP,size=1,prob=genome*mutrate))
      if (N_MUT>0){
        NEW_EXP<-sample(MUT[,"MEAN.YFP.MEDIAN.WT"],N_MUT, replace=TRUE)+rnorm(N_MUT, 0, EMS.BW)
        IND<-sample.int(100,size=N_MUT,replace=TRUE)
        POP[IND,i]<- POP[IND,i]*NEW_EXP
      }
    }
    SIM[k,]<-colMeans(POP)
    
    
  }
  
  write.table(SIM,paste0(levels(ALL.PROMOTERS.WTDH3$EXP)[j],"_MEDIAN.WT","_PROJECTION.MEANS_UNWEIGHTED_MULT_MUTRATE_PARENTSAMPLE_GEN",GEN/1000,"K_POPSIZE",N_POP,"_REP",REP,".txt"),sep="\t",quote=FALSE,row.names=FALSE)

  #Extract results

  GENE<-c("GPD1.SORT1.SC1","OST1.SORT1.SC1","PFY1.SORT1.SC1","RNR1.SORT1.SC1","RNR2.SORT1.SC1","STM1.SORT1.SC1","TDH1.SORT1.SC2","TDH2.SORT1.SC2","TDH3.SORT1.SC1","VMA7.SORT1.SC1")

  OUTPUT<-data.frame(matrix(ncol = 13, nrow = GEN))
  
  for (i in 1:GEN){
    OUTPUT[i,]<- quantile(SIM[2:REP,],probs=c(0,0.025,0.1,0.2,0.3,0.4,0.5,0.6,0.7,0.8,0.9,0.975,1))
  }
  
  write.table(OUTPUT,paste0("SUMMARY_",substr(GENE[j],start = 1, stop = 4),"_UNWEIGHTED_MULT_MUTRATE_SAMPLE_GEN50K_POPSIZE",N_POP,"_REP500.txt"),quote=FALSE, sep="\t",row.names = FALSE,col.names = FALSE)
  
  
  
##########################
#   Brownian Simulation  #
##########################
  
  load("INPUT.RData")
  
  REP<-500
  GEN<-50000
  N_POP<-100
  
  genome<-12495682
  mutrate<-1.67*10^-10
  
  j<-as.numeric(Sys.getenv("PBS_ARRAYID"))
  
  
  #Multiplicative
  #for(j in 1:length(levels(ALL.PROMOTERS.WTDH3$EXP))){
  
  CUR<-ALL.PROMOTERS.WTDH3[(ALL.PROMOTERS.WTDH3$CONDITION=="EMS"|ALL.PROMOTERS.WTDH3$CONDITION=="SHAM") & ALL.PROMOTERS.WTDH3$EXP==levels(ALL.PROMOTERS.WTDH3$EXP)[j],]
  SHAM.BW<-DENSITY.WTDH3.MEDIAN[(DENSITY.WTDH3.MEDIAN$CONDITION=="SHAM" & DENSITY.WTDH3.MEDIAN$EXP==levels(ALL.PROMOTERS.WTDH3$EXP)[j]),"BW"]
  EMS.BW<-DENSITY.WTDH3.MEDIAN[(DENSITY.WTDH3.MEDIAN$EXP==levels(ALL.PROMOTERS.WTDH3$EXP)[j]),"BW"]
  MUT<-CUR[CUR$CONDITION=="EMS",]
  MUT.SD<-sd(CUR[CUR$CONDITION=="EMS","MEAN.YFP.MEDIAN.WT"])
  
  SIM <- data.frame(matrix(ncol = GEN, nrow = REP))
  
  for(k in 1:REP){
    ANCESTRAL_SHAM<-sample(CUR[CUR$CONDITION=="SHAM","MEAN.YFP.MEDIAN.WT"],N_POP, replace=TRUE)+rnorm(N_POP, 0, SHAM.BW)
    POP <- data.frame(matrix(ncol = GEN, nrow = N_POP))
    POP[1:N_POP,1]<-ANCESTRAL_SHAM*100
    #POP[1:N_POP,1]<-100
    
    for(i in 2:GEN){
      POP[which(POP[,i-1]<0),i-1]<-0
      POP[,i]<-sample(POP[,(i-1)],replace=TRUE)
      N_MUT<-sum(rbinom(N_POP,size=1,prob=genome*mutrate))
      if (N_MUT>0){
        NEW_EXP<-rnorm(N_MUT, mean=1, sd = MUT.SD)
        IND<-sample.int(100,size=N_MUT,replace=TRUE)
        POP[IND,i]<- POP[IND,i]*NEW_EXP
      }
    }
    SIM[k,]<-colMeans(POP)
    
    
  }
  
  write.table(SIM,paste0(levels(ALL.PROMOTERS.WTDH3$EXP)[j],"_MEDIAN.WT","_PROJECTION.MEANS_UNWEIGHTED_MULT_BROWNIAN_PARENTSAMPLE_GEN",GEN/1000,"K_POPSIZE",N_POP,"_REP",REP,".txt"),sep="\t",quote=FALSE,row.names=FALSE)
  
  #Extract results
  OUTPUT<-data.frame(matrix(ncol = 13, nrow = GEN))
  
  for (i in 1:GEN){
    OUTPUT[i,]<- quantile(SIM[2:REP,],probs=c(0,0.025,0.1,0.2,0.3,0.4,0.5,0.6,0.7,0.8,0.9,0.975,1))
  }
  
  write.table(OUTPUT,paste0("SUMMARY_",substr(GENE[j],start = 1, stop = 4),"_UNWEIGHTED_MULT_BROWNIAN_SAMPLE_GEN",GEN/1000,"K_POPSIZE",N_POP,"_REP",REP,".txt"),quote=FALSE, sep="\t",row.names = FALSE,col.names = FALSE)
  
  
```
