## Supplemental tables, figures, files, and methods for "Empirical measures of mutational effects define neutral models of regulatory evolution in *Saccharomyces cerevisiae*": Supplementary Tables 01302019.pdf

Supplementary Table 1

| Gene Name | ORF | Mean Expression [a] | Expression CV [a] | Canonical TATA[b] | Nucleosome Occupancy Score [c] | Homozygote Deletion Fitness [d] |
|---|---|---|---|---|---|---|
| OST1 | YJL002C | 1075 | 12.2 | absent | -0.78 | 0.00 |
| VMA7 | YGR020C | 1428 | 12.1 | present | -0.71 | 0.66 |
| PFY1 | YOR122C | 1678 | 12.1 | absent | -1.23 | 0.00 |
| TDH1 | YJL052W | 1870 | 36.3 | present | 1.27 | 1.01 |
| GPD1 | YDL022W | 2206 | 21.7 | present | 1.75 | 1.03 |
| STM1 | YLR150W | 6741 | 15.1 | absent | 2.18 | 0.98 |
| RNR2 | YJL026W | 11336 | 11.4 | present | 1.24 | 0.00 |
| TDH2 | YJR009C | 15358 | 23.4 | present | - | 1.01 |
| RNR1 | YER070W | 27856 | 35.5 | present | 2.18 | 0.00 |
| TDH3 | YGR192C | 61318 | 12.1 | present | - | 0.97 |

[a] Mean and CV calculated based on GFP tagged fluorescence from Newman et al 2006

[b] Evidence for canonical TATA extracted from Basehoar et al 2004

[c] Nucleosome occupancy data provided from Tirosh and Barkai 2008

[d] Homozygote deletion fitness data from Steinmetz et al 2002

Supplementary Table 2

A.

| Assay | Condition | Median Initial Counts/ Sample | Initial Lower 95% Bound | Initial Upper 95% Bound | Median Final Counts/Sample | Final Lower 95% Bound | Final Upper 95% Bound | Minimum # Events/ Sample Included |
|---|---|---|---|---|---|---|---|---|
| GPD1 | EMS | 17626 | 7290 | 21278 | 4939 | 2501 | 7206 | 1426 |
| GPD1 | SHAM | 17841 | 13644 | 21482 | 5115 | 3353 | 6674 | 2465 |
| OST1 | EMS | 16706 | 623 | 20809 | 4273 | 2333 | 6240 | 1087 |
| OST1 | SHAM | 16790 | 603 | 21349 | 4225 | 2356 | 6338 | 1954 |
| PFY1 | EMS | 17306 | 562 | 22475 | 3841 | 2093 | 5135 | 1040 |
| PFY1 | SHAM | 17667 | 574 | 22266 | 3914 | 2997 | 5002 | 1637 |
| RNR1 | EMS | 17903 | 11332 | 22471 | 3646 | 2269 | 5014 | 1354 |
| RNR1 | SHAM | 18247 | 13652 | 21975 | 3621 | 2785 | 4630 | 1818 |
| RNR2 | EMS | 17921 | 11736 | 21008 | 4517 | 2876 | 6722 | 1033 |
| RNR2 | SHAM | 17976 | 15165 | 20949 | 4454 | 2951 | 6115 | 1841 |
| STM1 | EMS | 18445 | 4963 | 23190 | 4278 | 2270 | 6902 | 1062 |
| STM1 | SHAM | 18974 | 12853 | 22946 | 4376 | 2672 | 6556 | 1353 |
| TDH1 | EMS | 19310 | 1259 | 21158 | 4440 | 2252 | 6161 | 1053 |
| TDH1 | SHAM | 19369 | 13586 | 21105 | 4275 | 2426 | 5553 | 1518 |
| TDH2 | EMS | 18815 | 1998 | 22780 | 4428 | 2213 | 6549 | 1024 |
| TDH2 | SHAM | 19297 | 9963 | 22530 | 4237 | 2268 | 6044 | 1599 |
| VMA7 | EMS | 17917 | 12909 | 20695 | 4267 | 2659 | 6025 | 1096 |
| VMA7 | SHAM | 18091 | 14569 | 20891 | 4318 | 3062 | 5791 | 1867 |

B.

| Assay | EMS-treated | Promoter Sham | TDH3 Sham |
|---|---|---|---|
| GPD1 | 227 | 62 | 59 |
| OST1 | 217 | 60 | 53 |
| PFY1 | 196 | 52 | 54 |
| RNR1 | 225 | 60 | 57 |
| RNR2 | 210 | 59 | 58 |
| STM1 | 221 | 58 | 52 |
| TDH1 | 171 | 49 | 48 |
| TDH2 | 148 | 44 | 44 |
| TDH3 | 254 | 65 | 65 |
| VMA7 | 202 | 52 | 50 |

Supplementary Table 3

A.

| Assay | Sham Probability Decrease | Sham P-value BH adjusted | EMS Probability Decrease | EMS P-value BH adjusted |
|---|---|---|---|---|
| GPD1 | 0.45 | 0.90 | 0.44 | 0.19 |
| OST1 | 0.42 | 0.73 | 0.45 | 0.24 |
| PFY1 | 0.52 | 0.90 | 0.44 | 0.22 |
| RNR1 | 0.53 | 0.90 | 0.59 | 0.05 |
| RNR2 | 0.63 | 0.38 | 0.55 | 0.24 |
| STM1 | 0.48 | 0.90 | 0.38 | 0.00 |
| TDH1 | 0.59 | 0.38 | 0.50 | 1.00 |
| TDH2 | 0.47 | 0.90 | 0.53 | 0.30 |
| VMA7 | 0.54 | 0.90 | 0.55 | 0.24 |
| TDH3 | 0.48 | 0.99 | 0.54 | 0.29 |

B.

| Assay | Sham Permutation Statistic | Sham P-value | EMS Permutation Statistic | EMS P-value |
|---|---|---|---|---|
| GPD1 | -0.013 | 0.18 | -0.009 | 0.45 |
| OST1 | -0.010 | 0.16 | 0.001 | 0.51 |
| PFY1 | 0.004 | 0.45 | -0.004 | 0.50 |
| RNR1 | 0.001 | 0.50 | -0.009 | 0.15 |
| RNR2 | -0.005 | 0.50 | -0.036 | 0.01 |
| STM1 | 0.001 | 0.51 | 0.022 | 0.03 |
| TDH1 | -0.014 | 0.30 | 0.084 | 0.01 |
| TDH2 | -0.012 | 0.09 | 0.014 | 0.40 |
| VMA7 | -0.007 | 0.31 | -0.003 | 0.48 |

| | | | | |
|---|---|---|---|---|
| TDH3 | 0.003 | 0.487 | -0.013 | 0.335 |

Supplementary Table 4

| Promoter | Sample sizes | | Median | | MAD | | Medcouple | | Left Medcouple | | Right Medcouple | | % of mutants within: | | |
|---|---|---|---|---|---|---|---|---|---|---|---|---|---|---|---|
| | EMS | Sham | Native | Zscore | Native | Zscore | Native | Zscore | Native | Zscore | Native | Zscore | 1 SD | 2 SD | 3 SD |
| **GPD1** | 227 | 62 | 1.01 | 0.079 | 0.04 | 1.47 | 0.00 | 0.00 | -0.26 | -0.27 | 0.29 | 0.42 | 44% | 71% | 84% |
| **OST1** | 217 | 60 | 1.00 | 0.009 | 0.04 | 1.59 | -0.03 | -0.03 | -0.17 | -0.27 | 0.39 | 0.39 | 50% | 76% | 88% |
| **PFY1** | 196 | 52 | 1.00 | 0.177 | 0.05 | 1.44 | 0.03 | 0.03 | -0.29 | -0.18 | 0.28 | 0.47 | 46% | 74% | 86% |
| **RNR1** | 225 | 60 | 1.00 | -0.280 | 0.03 | 2.20 | -0.09 | -0.09 | -0.24 | -0.28 | 0.25 | 0.14 | 58% | 86% | 94% |
| **RNR2** | 210 | 59 | 1.00 | 0.250 | 0.08 | 1.27 | -0.23 | -0.23 | -0.27 | -0.04 | 0.26 | 0.03 | 39% | 68% | 83% |
| **STM1** | 221 | 58 | 1.01 | 0.399 | 0.03 | 1.65 | 0.08 | 0.08 | -0.12 | -0.18 | 0.55 | 0.64 | 48% | 78% | 89% |
| **TDH1** | 171 | 49 | 1.00 | -0.055 | 0.10 | 1.88 | 0.13 | 0.13 | -0.20 | -0.05 | 0.45 | 0.50 | 49% | 74% | 84% |
| **TDH2** | 148 | 44 | 0.99 | -0.081 | 0.04 | 1.52 | 0.15 | 0.15 | -0.44 | -0.26 | 0.51 | 0.61 | 57% | 85% | 93% |
| **TDH3** | 254 | 65 | 1.00 | 0.054 | 0.02 | 2.97 | -0.05 | -0.04 | -0.48 | -0.47 | 0.43 | 0.52 | 35% | 60% | 79% |
| **VMA7** | 202 | 52 | 0.99 | -0.124 | 0.04 | 1.26 | 0.01 | 0.01 | -0.25 | -0.10 | 0.32 | 0.38 | 53% | 79% | 87% |

Supplementary Table 5

| Assay | EMS Shapiro's W | EMS P-value BH adjusted |
|---|---|---|
| GPD1 | 0.69 | 8.96E-20 |
| OST1 | 0.90 | 7.14E-11 |
| PFY1 | 0.86 | 2.73E-12 |
| RNR1 | 0.88 | 3.51E-12 |
| RNR2 | 0.90 | 7.14E-11 |
| STM1 | 0.68 | 8.96E-20 |
| TDH1 | 0.71 | 1.31E-16 |
| TDH2 | 0.77 | 8.92E-14 |
| TDH3 | 0.31 | 7.29E-29 |
| VMA7 | 0.95 | 4.06E-06 |

Supplementary Table 6

| Promoter 1 | Promoter 2 | Anderson-Darling statistic | EMS P-value BH adjusted | |
|---|---|---|---|---|
| GPD1 | OST1 | 0.461 | 0.788 | |
| GPD1 | PFY1 | 0.527 | 0.736 | |
| GPD1 | RNR1 | 4.62 | 0.022 | * |
| GPD1 | RNR2 | 1.6 | 0.209 | |
| GPD1 | STM1 | 3.1 | 0.064 | |
| GPD1 | TDH1 | 2.22 | 0.116 | |
| GPD1 | TDH2 | 0.555 | 0.725 | |
| GPD1 | VMA7 | 1.7 | 0.196 | |
| GPD1 | TDH3 | 1.39 | 0.271 | |
| OST1 | PFY1 | 1.35 | 0.272 | |
| OST1 | RNR1 | 3.77 | 0.039 | * |
| OST1 | RNR2 | 2.23 | 0.116 | |
| OST1 | STM1 | 4.51 | 0.022 | * |
| OST1 | TDH1 | 1.75 | 0.188 | |
| OST1 | TDH2 | 0.632 | 0.679 | |
| OST1 | VMA7 | 1.34 | 0.272 | |
| OST1 | TDH3 | 0.584 | 0.711 | |
| PFY1 | RNR1 | 6.31 | 0.005 | ** |
| PFY1 | RNR2 | 1.65 | 0.203 | |
| PFY1 | STM1 | 1.98 | 0.150 | |
| PFY1 | TDH1 | 3.34 | 0.055 | |
| PFY1 | TDH2 | 1.27 | 0.296 | |
| PFY1 | VMA7 | 2.9 | 0.073 | |
| PFY1 | TDH3 | 2.84 | 0.074 | |
| RNR1 | RNR2 | 8.35 | 0.002 | ** |

| | | | | |
|---|---|---|---|---|
| RNR1 | STM1 | 10.6 | 0.000 | *** |
| RNR1 | TDH1 | 3.96 | 0.034 | * |
| RNR1 | TDH2 | 2.55 | 0.091 | |
| RNR1 | VMA7 | 6.11 | 0.005 | ** |
| RNR1 | TDH3 | 2.72 | 0.081 | |
| RNR2 | STM1 | 5.63 | 0.008 | ** |
| RNR2 | TDH1 | 6.1 | 0.005 | ** |
| RNR2 | TDH2 | 2.92 | 0.073 | |
| RNR2 | VMA7 | 1.86 | 0.169 | |
| RNR2 | TDH3 | 3.2 | 0.061 | |
| STM1 | TDH1 | 3.35 | 0.055 | |
| STM1 | TDH2 | 2.55 | 0.091 | |
| STM1 | VMA7 | 7.81 | 0.002 | ** |
| STM1 | TDH3 | 7.6 | 0.002 | ** |
| TDH1 | TDH2 | 0.813 | 0.531 | |
| TDH1 | VMA7 | 4.04 | 0.034 | * |
| TDH1 | TDH3 | 2.42 | 0.101 | |
| TDH2 | VMA7 | 2.33 | 0.109 | |
| TDH2 | TDH3 | 1.03 | 0.394 | |
| VMA7 | TDH3 | 1.08 | 0.376 | |

Supplementary Table 7

**A. Multiple regression coefficients for model: Log Mean Population Phenotype after 50K Generations ~ Promoter * Model Type**

Adj R2 = 0.5389, $F_{19,9980}$ = 616.2, p-value < 2.2 * 10-16

| Term | Estimate | Standard Error | t value | P value | Significance |
|---|---|---|---|---|---|
| (Intercept) | 4.10 | 0.05 | 80.2 | <2e-16 | *** |
| MODEL-EMPIRICAL | 0.06 | 0.07 | 0.8 | 0.414 | |
| ASSAY-OST1 | 0.35 | 0.07 | 4.8 | 0.000 | *** |
| ASSAY-PFY1 | 0.19 | 0.07 | 2.7 | 0.007 | ** |
| ASSAY-RNR1 | 0.46 | 0.07 | 6.4 | 0.000 | *** |
| ASSAY-RNR2 | -0.07 | 0.07 | -0.9 | 0.343 | |
| ASSAY-STM1 | 0.25 | 0.07 | 3.5 | 0.000 | *** |
| ASSAY-TDH1 | -1.66 | 0.07 | -23.0 | <2e-16 | *** |
| ASSAY-TDH2 | -0.14 | 0.07 | -1.9 | 0.061 | . |
| ASSAY-TDH3 | 0.09 | 0.07 | 1.3 | 0.204 | |
| ASSAY-VMA7 | 0.35 | 0.07 | 4.8 | 0.000 | *** |
| MODEL-EMPIRICAL:ASSAY-OST1 | 0.30 | 0.10 | 3.0 | 0.003 | ** |
| MODEL-EMPIRICAL:ASSAY-PFY1 | 0.28 | 0.10 | 2.7 | 0.007 | ** |
| MODEL-EMPIRICAL:ASSAY-RNR1 | -1.12 | 0.10 | -10.9 | <2e-16 | *** |
| MODEL-EMPIRICAL:ASSAY-RNR2 | -2.71 | 0.10 | -26.5 | <2e-16 | *** |
| MODEL-EMPIRICAL:ASSAY-STM1 | 1.98 | 0.10 | 19.4 | <2e-16 | *** |
| MODEL-EMPIRICAL:ASSAY-TDH1 | 4.81 | 0.10 | 47.0 | <2e-16 | *** |
| MODEL-EMPIRICAL:ASSAY-TDH2 | 0.14 | 0.10 | 1.4 | 0.160 | |
| MODEL-EMPIRICAL:ASSAY-TDH3 | -1.94 | 0.10 | -19.0 | <2e-16 | *** |
| MODEL-EMPIRICAL:ASSAY-VMA7 | -0.66 | 0.10 | -6.5 | 0.000 | *** |

**B.**

**Brownian Motion: Log Grand Median Population Phenotype at Generation 50K ~ MAD + MC**

| | Estimate | Standard Error | t value | P value | Signicance |
|---|---|---|---|---|---|
| (Intercept) | 5.0755 | 0.1594 | 31.843 | 7.79E-09 | *** |
| MAD | -20.923 | 3.093 | -6.765 | 0.000261 | *** |
| MC | -2.4822 | 0.681 | -3.645 | 0.008231 | ** |

Adj R2 = 0.8569, $F_{2,7}$ = 27.95, FDR corrected p-value= 0.0004599

**Full Empirical: Log Grand Median Population Phenotype at Generation 50K ~ MC +LMC**

| | Estimate | Standard Error | t value | P value | Signicance |
|---|---|---|---|---|---|
| (Intercept) | 7.1011 | 0.7715 | 9.205 | 3.68E-05 | *** |
| MC | 12.1226 | 2.2507 | 5.386 | 0.00102 | ** |
| LMC | 11.4364 | 2.8824 | 3.968 | 0.00541 | ** |

Adj R2 = 0.81, $F_{2,7}$ = 20.15, FDR corrected p-value= 0.0037

Supplementary Table 8

| Promoter | Gaussian ΔBIC | LaPlace Symm ΔBIC | LaPlace Asymm ΔBIC | Best BIC | MLL | P1 | P2 | P3 | BIC |
|---|---|---|---|---|---|---|---|---|---|
| All Data | 1963 | 874 | 0 | LP Asymm | -1239 | 0.00 | 0.46 | 0.50 | 9469 |
| GPD1 | 173.3 | 94.1 | 0 | LP Asymm | -518 | 0.14 | 0.90 | 0.52 | 1052 |
| OST1 | 53.3 | 39.1 | 0 | LP Asymm | -459 | 0.01 | 0.76 | 0.50 | 934 |
| PFY1 | 73.6 | 35.3 | 0 | LP Asymm | -409 | 0.19 | 0.74 | 0.51 | 834 |
| RNR1 | 65.9 | 194.9 | 0 | LP Asymm | -555 | 0.11 | 1.05 | 0.59 | 1127 |
| RNR2 | 46.8 | 24.1 | 0 | LP Asymm | -375 | 0.65 | 0.49 | 0.66 | 767 |
| STM1 | 172.5 | 163.3 | 0 | LP Asymm | -510 | -0.40 | 0.79 | 0.31 | 1036 |
| TDH1 | 131.0 | 174.1 | 0 | LP Asymm | -403 | -1.28 | 0.71 | 0.24 | 821 |
| TDH2 | 92.2 | 113.1 | 0 | LP Asymm | -364 | -0.21 | 1.06 | 0.45 | 742 |
| VMA7 | 19.8 | 1.6 | 0 | LP Asymm*/Symm | -373 | -0.12 | 0.58 | 0.50 | 762 |
| TDH3 | 457.1 | 243.2 | 0 | LP Asymm | -631 | 0.54 | 1.03 | 0.63 | 1278 |
